# The N-Terminus of *Sophora tonkinensis* Cytochrome P450s Evolves Neutrally yet Encodes Rich Functional Information: A Protein Language Model Analysis

**DOI:** 10.64898/2026.03.06.710024

**Authors:** Zhu Qiao, Jing Wang, Ben Qin, Fan Wei, Ying Liang

**Affiliations:** Guangxi Key Laboratory of Medicinal Resources Protection and Genetic Improvement, Guangxi Engineering Research Center of TCM Resource Intelligent Creation, National Center for TCM Inheritance and Innovation, Guangxi Botanical Garden of Medicinal Plants, Nanning 530023, China

**Keywords:** Cytochrome P450, N-terminal membrane anchor, protein language model, relaxed purifying selection, protein embeddings, membrane topology, *Sophora tonkinensis*

## Abstract

The N-terminal membrane anchor of cytochrome P450s is essential for function yet exceptionally variable—a paradox resistant to alignment-based analysis. Using an alignment-free protein language model (ESM2) pipeline on 345 *Sophora tonkinensis* P450s, we show the N-terminal 50 residues evolve under pervasive neutral relaxation (mean dispersion 0.62), punctuated by a single constrained island: a PxxG structural hinge (P^21^, G^24^ conserved at 98%/99%). Despite lacking sequence-level constraint, embedding-space decomposition resolves this scaffold into two separable channels. DIVA (unsupervised) captures a topological template encoding membrane-anchoring architecture, validated by physicochemical correlations recapitulating ER signal-anchor insertion and confirmed by DeepLoc- 2.1. PIVOT (supervised ablation) captures a family-associated signal whose peak position distinguishes CYP families (ε^2^ = 0.44, permutation p < 0.0001) despite label-free training. The channels are largely separable: PIVOT peaks lie downstream of DIVA peaks in 84.4% of proteins, with mutually exclusive signal-loss sets. The P450 N-terminus thus integrates membrane-topological and family-identity information within a single relaxed region. Neutral evolution is reframed not as absence of information, but as the condition enabling multiplexed functional encoding—a mechanistic rationale for the functional importance of this variable anchor.

## 1. Introduction

The structural diversity and bioactivity of plant specialized metabolites, such as the anti-malarial artemisinin and the chemotherapeutic paclitaxel, underpin both ecological resilience and modern pharmacopeia (Elshafie et al., 2023; Ramawat & Mérillon, 2025; Tissier et al., 2014). This chemical ingenuity is largely orchestrated by cytochrome P450 monooxygenases (P450s), which catalyze regio- and stereo-specific oxidations, dramatically diversifying metabolic scaffolds (Banerjee & Hamberger, 2018; Zhao et al., 2025). As one of nature’s largest and most widespread enzyme superfamilies, P450s have undergone continuous expansion and functional divergence, making them an exceptional model for studying evolutionary innovation in plant metabolism (Fang et al., 2024; Hamberger & Bak, 2013).

However, the very features that make P450s evolutionarily fascinating—high sequence diversity and limited conservation in non-core regions—pose fundamental challenges for traditional sequence-based analysis. Phylogenetic methods reliant on homology often struggle to resolve fine-scale functional shifts and are particularly ill-suited for detecting convergent evolution (Zallot et al., 2016). More critically, methods that detect positive selection or functional constraint from multiple sequence alignments (MSAs)—including codon-based selection models and conservation scoring algorithms—share a common prerequisite: they require unambiguous positional homology across all sequences. When this prerequisite is violated, as it is in regions with extensive insertions and deletions (indels), downstream inferences become systematically unreliable (Fletcher & Yang, 2010).

A particularly instructive case is the N-terminal signal sequence of plant P450s. The N-terminal ∼30-50 residues form a hydrophobic transmembrane helix that anchors the enzyme to the cytosolic face of the endoplasmic reticulum—a function essential for correct subcellular localization, membrane topology determination, and access to cognate NADPH-cytochrome P450 reductases (Werck-Reichhart & Feyereisen, 2000; Schuler & Werck-Reichhart, 2003). Yet this same region is among the most sequence-divergent segments across the P450 superfamily, tolerating extensive indels and substitutions that would be incompatible with the structurally constrained catalytic core (Hansen et al., 2021). Indeed, Schuler & Werck-Reichhart (2003) explicitly acknowledged that the “high degrees of variation and spacing” in the N-terminus “prevent unambiguous alignment.” Two decades later, no method has overcome this fundamental limitation: MSA-based analyses either exclude the N-terminus or proceed with alignments whose columns contain functionally non-homologous residues, producing results that are difficult to interpret biologically.

This MSA-dependent blind spot is not merely a technical inconvenience—it obscures a potentially important biological question. The N-terminus presents an apparent paradox: it is simultaneously functionally indispensable and highly variable. Does N-terminal sequence variation merely reflect relaxed constraint on a passive membrane anchor—as expected under the neutral theory of molecular evolution (Kimura, 1968)—or does it encode functional information that contributes actively to evolutionary diversification? Individual case studies support the latter possibility. Renault et al. (2017) demonstrated that a gene duplication event in the CYP98A family produced two paralogs with markedly different N-terminal hydrophobicities, resulting in a membrane topology inversion accompanied by functional divergence in substrate preference. Similarly, members of the CYP81A subfamily in grasses metabolize herbicides from at least seven distinct chemical classes (Dimaano & Iwakami, 2021), and their N-terminal regions show signatures of adaptive evolution that may contribute to their exceptional substrate promiscuity. However, whether topology-modifying N-terminal variation is a general driver of P450 functional divergence—and through what molecular encoding strategy it operates—has not been systematically investigated at superfamily scale, in large part because the required analytical tools have been unavailable.

The recent advent of Protein Language Models (PLMs), such as ESM2, offers a transformative approach that fundamentally bypasses the MSA requirement (Lin et al., 2023). Trained on evolutionary-scale sequence data, ESM2 learns the biophysical “grammar” of proteins, projecting sequences into a functional semantic space where proximity reflects functional similarity independent of direct homology (Yeung et al., 2023; Wiatrak et al., 2025). Because ESM2 embeddings capture sequence features across 1,280 dimensions—including higher-order co- evolutionary couplings and long-range context dependencies—they enable, in principle, the decomposition of a polypeptide segment into multiple information components that may represent distinct functional constraints. Critically, these embeddings can be analyzed geometrically without requiring positional alignment: one can ask whether the embedding vectors at a given position cluster tightly across sequences (indicating strong functional constraint) or scatter randomly (indicating relaxed evolution), without ever needing to establish which residue in one sequence “corresponds to” which residue in another.

Recent theoretical developments further motivate a residue-level dissection of the P450 N-terminus. Werck-Reichhart (2023) proposed that catalytic promiscuity—the ability of a single P450 to convert multiple structurally diverse substrates—is a widespread driver of plant P450 family expansion and functional diversification. This model, however, focuses primarily on the catalytic domain. An open question is whether the N-terminal signal sequence harbors an independent evolutionary channel—potentially operating through altered subcellular localization rather than altered substrate specificity—that complements catalytic promiscuity in generating functional diversity. More fundamentally, the conventional view that a contiguous polypeptide segment encodes a single functional signal may reflect the limitations of the low-dimensional physicochemical frameworks (hydrophobicity scales, helical propensity indices, charge distribution) that have dominated protein analysis for decades (Kyte & Doolittle, 1982; Chou & Fasman, 1978). In a high-dimensional sequence-feature space, two functional constraints that would compete for the same physicochemical axes may occupy approximately separable subspaces, enabling simultaneous encoding of distinct information types on the same sequence segment. Whether such dimensionality-driven constraint decomposition operates in natural proteins—and whether it contributes to evolvability—has not, to our knowledge, been empirically tested in any protein family.

Leveraging these advances, this study presents an integrative analysis of the P450 superfamily from *Sophora tonkinensis*—a medicinal plant valued for its anti-inflammatory alkaloids and flavonoids (Zeng et al., 2025). We employ a three-pronged computational strategy that proceeds from constraint detection to signal decomposition to independent validation. First, we introduce PEACE (PLM Embedding Analysis for Constraint Estimation), an alignment-free framework that measures per-position evolutionary constraint directly from ESM2 embedding geometry, bypassing the MSA requirement entirely and operating in two complementary modes for cross-validation. PEACE establishes the neutral-relaxation baseline of the N-terminus and identifies the sole constrained element— the PxxG structural hinge. Second, we decompose the constraint-free N-terminal segment into two information channels using complementary per-residue importance metrics: DIVA (Differential Vector Attribution), an unsupervised measure capturing each residue’s alignment with the protein’s intrinsic functional specificity vector, which recovers a conserved topological template; and PIVOT (Perturbation-based Importance Vector Orientation via Transformers), a supervised ablation-based measure quantifying each residue’s marginal contribution to functional cluster discrimination, which captures a family-associated semantic signal. Third, we validate the separable-channel architecture against an independent localization predictor (DeepLoc-2.1), demonstrating that DIVA and PIVOT signals are differentially recognized by a state-of-the-art tool that was agnostic to our decomposition. Together, these methods enable us to investigate a long-standing question in plant P450 biology: whether the N-terminal signal sequence, whose sequence variability has precluded traditional MSA-based analysis, encodes functionally meaningful information that contributes to the evolutionary diversification of this enzyme superfamily. The results reveal that the N-terminus simultaneously accommodates multiple functional constraints through a mechanism that has not, to our knowledge, been previously described.

## 2. Materials and Methods

### 2.1 Identification and sequence analysis of P450 genes in *S. tonkinensis*

The identification of the P450 gene family was initiated with all annotated protein sequences derived from our previously sequenced and assembled genome of *S. tonkinensis*. The initial candidate proteins were retrieved by searching the proteome with the hidden Markov model (HMM) profile for the conserved P450 domain (Pfam: PF00067), obtained from the InterPro database (https://www.ebi.ac.uk/interpro/entry/pfam). Subsequently, an iterative search was performed using a custom, species-specific HMM model built from these initial candidates [HMMER 3.3.2; http://hmmer.org/; (Eddy, 2011)]. All resulting sequences were subjected to verification against the Conserved Domain Database (CDD) (https://www.ncbi.nlm.nih.gov/Structure/bwrpsb/bwrpsb.cgi) to remove those with incomplete domains.

### 2.2 Construction of phylogenetic tree for P450 protein in *S. tonkinensis*

A phylogenetic tree was constructed to elucidate the evolutionary relationships among the identified P450 proteins in *S. tonkinensis*. The full-length amino acid sequences of all identified P450 proteins were first aligned using MUSCLE with default parameters [MUSCLE v5.1; https://drive5.com/muscle/index.htm; (Edgar, 2022)]. The resulting multiple sequence alignment was subsequently refined using trimAl to remove poorly aligned positions and divergent regions [trimAl v1.4.rev15; https://trimal.cgenomics.org/; (Capella-Gutiérrez et al., 2009)], thereby enhancing the phylogenetic signal. A maximum likelihood phylogenetic tree was then inferred from the trimmed alignment using IQ-TREE 2 with the best-fit substitution model selected automatically based on the Bayesian Information Criterion. Branch support was assessed with 1000 ultrafast bootstrap replicates [IQ-TREE multicore version 2.2.2.2; http://www.iqtree.org/; (Minh et al., 2020)]. The final tree was visualized and annotated using the Interactive Tree of Life online tool (iTOL: https://itol.embl.de/).

### 2.3 Conservative motif and chromosome location analysis of *S. tonkinensis* P450 protein

To identify conserved motifs, the P450 protein sequences were analyzed with MEME [MEME version 5.5.1; https://meme-suite.org/meme/; (Bailey & Elkan, 1994)], configured to discover 10 motifs with widths ranging from 6 to 100 amino acids. The resulting motifs were subsequently processed using FIMO (FIMO version 5.5.1; https://meme-suite.org/meme/; (Grant et al., 2011)), and the output was visualized with a custom R script [R version 4.5.0; specifically the ggplot2 (Wickham, 2016) and Biostrings packages (Pagès et al., 2025)]. For chromosomal localization, the genomic coordinates of the P450 genes were determined and their distribution across chromosomes was plotted using another R script [R version 4.5.0; specifically the ggplot2 (Wickham, 2016) and scales packages (Wickham et al., 2025)].

### 2.4 Collinearity analysis within the genome

Based on the genome annotation of the *S. tonkinensis* and the BLAST [BLAST 2.6.0+; https://ftp.ncbi.nlm.nih.gov/blast/executables/blast+/; (Altschul et al., 1990)] comparison results of all encoded protein sequences within the genome, MCscanX was employed to identify collinear gene blocks [https://github.com/wyp1125/MCScanX; (Wang et al., 2012)]. Gene pairs resulting from large-scale duplication events specific to the P450 gene family were subsequently extracted. The results of this intra-species gene duplication analysis were visualized using Circos software [circos v0.69-8; http://www.circos.ca; (Krzywinski et al., 2009)].

Furthermore, to investigate interspecies synteny, we performed a comparative genomic analysis against three reference species: *Arabidopsis thaliana*, *Arachis hypogaea* (peanut), and *Glycine max* (soybean). Given that peanut and soybean belong to the same family as the species under study (though representing different genera), this comparison provides valuable evolutionary insights. The analysis and visualization of syntenic relationships for both the whole genome and specific gene families were conducted using the Python-based jcvi tool kit [https://github.com/tanghaibao/jcvi; (Tang et al., 2024)].

### 2.5 Functional Landscape Analysis Using ESM2

All P450 sequences were projected into a functional embedding space using the ESM2 model (ESM2-650m-t33) (Lin et al., 2023). A pairwise cosine distance matrix was calculated from the mean-pooled embeddings, and hierarchical clustering was applied to define functional clusters. Phylogenetic clusters were independently identified as monophyletic groups from a maximum-likelihood tree. The congruence between functional and phylogenetic clustering was quantified using the V-measure (Rosenberg & Hirschberg, 2007) and Adjusted Mutual Information (AMI) (Vinh et al., 2010). Resulting discrepancies were visualized using Sankey diagrams.

These analyses were performed using a modular set of custom Python scripts, designed to facilitate maintenance and iterative development. The complete analysis pipeline, including scripts, is available on GitHub (https://github.com/qiaoweizhuo-pixel/P450/tree/main/ESM_EVOL_ALS2).

Notably, we identified 13 phylogenetically closely related protein pairs that were assigned to distinct ESM2-based functional clusters. To investigate this divergence, we systematically compared the sequence differences within each pair. This analysis revealed that substantial variation—predominantly involving deletions and substitutions—was concentrated in the N-terminal regions, whereas the remaining sequences exhibited only sporadic amino acid differences. Consequently, subsequent investigations focused on the role of N-terminal sequence variation in functional differentiation.

### 2.6 PEACE: A dual-mode alignment-free framework for positional constraint estimation and directional divergence analysis using protein language models

Traditional MSA-based evolutionary analyses are often unreliable for divergent protein families like plant CYP450s, due to extensive indels at the N-terminus (Schuler & Werck-Reichhart, 2003; Nelson, 2006; Sezutsu et al., 2013).

Briefly, to illustrate this problem, a multiple sequence alignment of the first 58 N-terminal positions was generated using MUSCLE v5.1 (Edgar, 2022) and subjected to the Mixed Effects Model of Evolution (MEME; Murrell et al., 2012) as implemented in HyPhy v2.5 (Kosakovsky Pond et al., 2020), with default parameters and a significance threshold of p ≤ 0.01. MEME was selected because it is specifically designed to detect episodic positive selection affecting a subset of lineages at individual sites—the pattern expected if N-terminal variation reflects adaptive functional diversification. Applying HyPhy MEME to the first 58 N-terminal positions identified a proportion of positively selected sites that notably surpassed that of even well-known positive selection hotspots (17/58=29.31%, Supplementary Figure S1), such as antibody CDRs (Chang & Casali, 1994; Bose & Sinha, 2005). Such an abnormally high signal is inconsistent with genuine biological selection in CYP450s and instead aligns with alignment-induced dN inflation in indel-rich regions (Wong et al., 2008; Fletcher & Yang, 2010; Markova-Raina & Petrov, 2011; Blackburne & Whelan, 2013). Therefore, an alignment-free approach is essential to circumvent these artifacts.

PEACE (Protein Language Model Embedding Analysis for Constraint Estimation) is a computational framework designed to infer positional evolutionary constraints and directional divergence across a protein family without the requirement of multiple sequence alignment (MSA). The algorithm leverages embeddings from a pre-trained protein language model and provides two complementary matching modes for robust analysis of divergent protein families (Figure 1).

**Figure 1.**
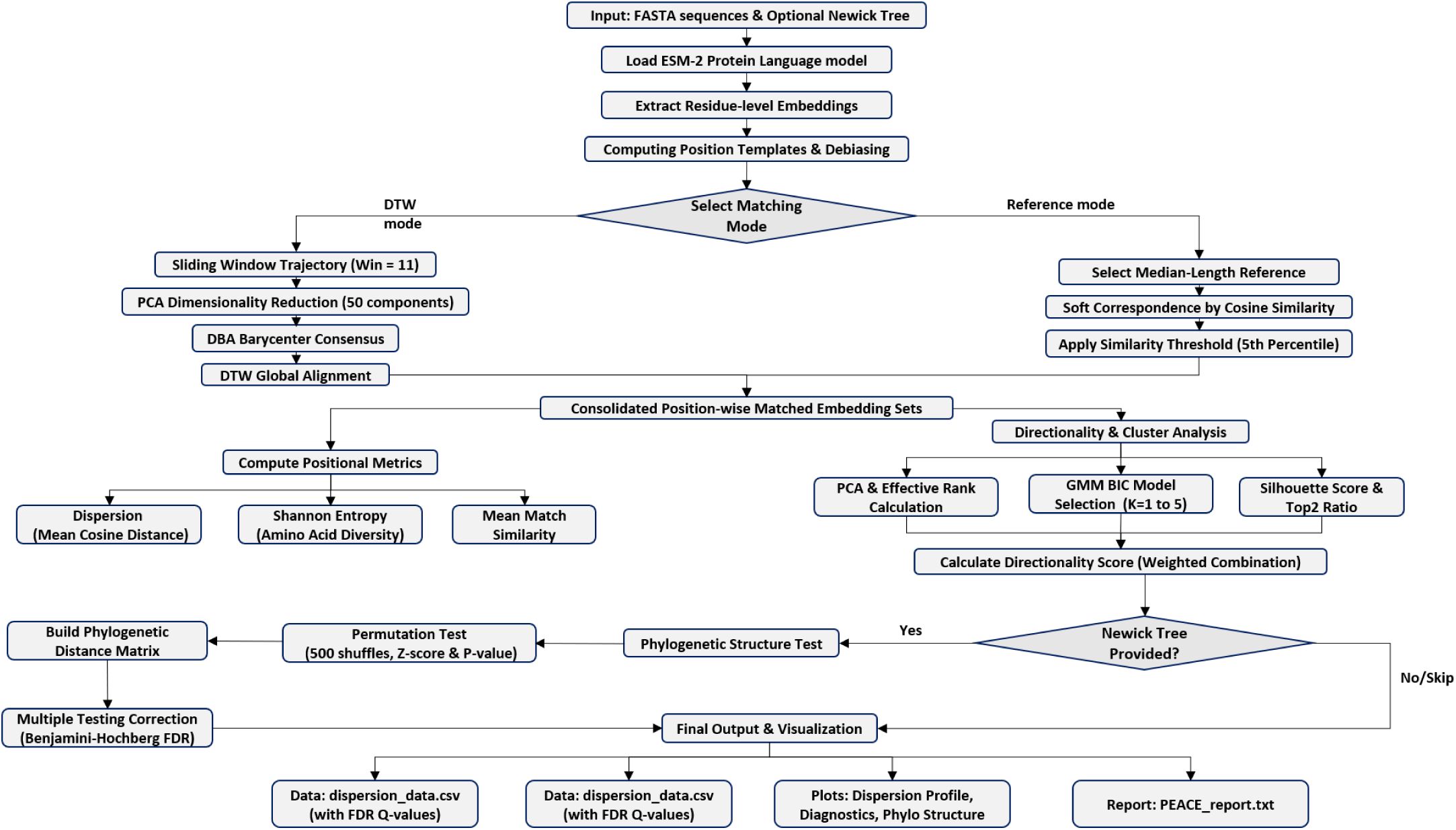
Overview of the PEACE algorithmic pipeline. The flowchart illustrates the dual-mode architecture for analyzing evolutionary constraints. (1) Sequences and optional phylogenetic trees are input, and embeddings are extracted via ESM2 followed by position-specific debiasing. (2) The analysis branches into two modes: DTW mode (using sliding windows, PCA, and DBA alignment) for robust regional consensus, and Reference mode (using median-length reference and soft correspondence) for residue-level precision. (3) Positional metrics (dispersion, Shannon entropy, match similarity) are derived from the matched embedding sets. (4) Directional divergence is quantified using effective rank, GMM clustering, and a composite Directionality Score. (5) If a phylogenetic tree is provided, a permutation-based test coupled with Benjamini-Hochberg FDR correction assesses clade-specific selection signals. (6) The pipeline generates comprehensive data tables (CSV), diagnostic plots, and a summary report.

#### Embedding extraction and debiasing

The input protein sequences were processed by the ESM2 protein language model (esm2_t33_650M_UR50D). Residue-level embeddings were extracted from the last hidden layer representations. To eliminate global positional background biases introduced by the overall amino acid composition of the family, a “position template” was constructed for every distinct absolute alignment coordinate by averaging the embeddings of sequences occupying that coordinate. Debiased embeddings were then generated by subtracting these templates from the corresponding original residue vectors. All downstream analyses were performed using these debiased embeddings.

#### Dual-mode matching strategies

Given the high sequence divergence and frequent indels inherent in ancient gene families, PEACE operates via two complementary modes to construct a per-position matched embedding dataset:

##### *Reference mode* (residue-level resolution)

The median-length sequence within the family was selected as the reference. For each residue position *i* on the reference sequence, the cosine similarity was computed between its debiased embedding and every embedding in each target sequence. For each target sequence, the residue with the maximum cosine similarity (soft correspondence) was identified as the matched position for reference position *i*. A matching threshold was determined as the 5th percentile of the maximum cosine similarities observed from a random subsample of positions and target sequences, which was used to filter out low-confidence alignments.

##### *DTW mode* (window-level regional resolution)

A sliding window (default window size = 11) was applied to each debiased sequence, and the embeddings within the window were averaged to construct a “trajectory.” To mitigate high-dimensional noise, Principal Component Analysis (PCA) was performed to reduce the dimensionality of all window-mean embeddings to 50 components. A family-wide consensus trajectory was then generated using Dynamic Time Warping Barycenter Averaging (DBA; Petitjean et al., 2011). Subsequently, each sequence’s reduced trajectory was aligned to the consensus trajectory using the DTW algorithm (Giorgino, 2009), and the corresponding window-mean embeddings in the original high-dimensional space were collected for each consensus position.

#### Position-specific evolutionary constraint metrics

Based on the collection of matched embeddings for a given position, the following metrics were calculated:

##### Dispersion

Defined as the mean pairwise cosine distance among all matched embeddings. Low dispersion indicates strong purifying selection, whereas high dispersion suggests relaxed or diversified selection.

##### Mean Match Similarity

In Reference mode, this metric represented the direct average of the original cosine similarities. In DTW mode, due to the nature of window-means, this metric mathematically equals 1 - dispersion, serving as a consistent indicator.

##### Shannon Entropy

Calculated based on the frequency of amino acid residues within the matched sets to evaluate sequence-level amino acid diversity.

#### Directionality score and GMM-based clustering

To distinguish whether divergence at a position is stochastically isotropic or follows directional/adaptive patterns, a Directionality Score (DS) was implemented. PCA was applied to the matched embeddings, and Gaussian Mixture Models (GMMs) with BIC optimization were utilized to automatically determine the optimal number of clusters (ranging from 1 to 5). The DS was calculated as a weighted combination of three features: Effective Rank (derived from the PCA eigenvalues), Structure Score (silhouette score if ≥2 clusters, otherwise the Top-2 ratio of explained variance), and a Dispersion Term (non-linear hyperbolic tangent transformation). Positions with high DS (>0.6) and low effective rank were identified as high-priority candidates for directional divergence.

#### Phylogenetic structure test and FDR correction

To determine whether the GMM-inferred clusters are significantly associated with the underlying species phylogeny, a permutation-based phylogenetic structure test was incorporated into PEACE. Using a user-provided Newick phylogenetic tree, a pairwise phylogenetic distance matrix was computed. For each evaluated position, the observed mean within-cluster phylogenetic distance was calculated. To build a null distribution, the cluster labels were randomly shuffled 500 times, and the within-cluster distances were recalculated. A Z-score and an empirical p-value were computed based on this null distribution. Because hundreds of positions are tested simultaneously across the protein, the Benjamini-Hochberg (BH) procedure was applied to calculate false discovery rate (FDR)-adjusted q-values, significantly mitigating the risk of false positives.

#### Visualization and statistical analysis

To intuitively illustrate the structural divergence at the top-ranked directional positions, the matched embedding sets were projected onto a two-dimensional plane. While PCA was used as the default method due to computational efficiency, PEACE also supports non-linear dimensionality reduction algorithms, specifically t-SNE (van der Maaten & Hinton, 2008) and UMAP (McInnes et al., 2018), which can be enabled by user-defined parameters. Additionally, the framework automatically performs a comparative analysis between the N-terminal region (by default, the first 50 residues) and the core region, including Mann-Whitney U tests to identify regional biases in constraint regimes.

#### Implementation

PEACE was implemented in Python 3.12 and leverages the PyTorch and HuggingFace Transformers libraries for deep learning inference. The scikit-learn library was utilized for dimensionality reduction (PCA, t-SNE), GMM clustering, and statistical assessments, while the statsmodels library handled the multiple testing correction (BH-FDR). The complete script is available on GitHub (https://github.com/qiaoweizhuo-pixel/P450/tree/main/PEACE)

#### Cross-species PxxG hinge scan

P450 sequences for eight land plant species were retrieved from UniProtKB (The UniProt Consortium, 2025) via the REST API (rest.uniprot.org/uniprotkb/search) on 2026-08-06, using taxonomy-restricted queries with the “cytochrome p450” protein family annotation (fallback: EC 1.14.-.- or gene symbol CYP*). For *Sophora tonkinensis*, the 345 P450 proteins from Section 2.1 were used. All sequences were scanned for the PxxG core motif (P-X-X-G) within the first 60 residues; match positions were recorded and summarized by median position and the proportion of matches within 50 residues, to distinguish genuine N-terminal hinges from spurious internal hits. The complete script is available on GitHub (https://github.com/qiaoweizhuo-pixel/P450/tree/main/PXXG_scan).

### 2.7 DeepLoc-2.1 subcellular localization prediction

Subcellular localization predictions were generated using the DeepLoc-2.1 online service (https://services.healthtech.dtu.dk/services/DeepLoc-2.1/; Nielsen, 2025). The full-length amino acid sequences of all 345 *Sophora tonkinensis* P450s were submitted via the web interface in FASTA format with default parameters. Output included a predicted localization class and a probability score for each category (cytoplasm, extracellular, membrane, mitochondrion, nucleus, plastid, lysosome/vacuole, peroxisome, Golgi apparatus, endoplasmic reticulum). For subsequent analysis, proteins classified as “endoplasmic reticulum” were designated ER-predicted; all other categories were grouped as non-ER. The ER probability score was used as a continuous metric to assess correlation with N-terminal residue-importance features.

### 2.8 Per-Residue Importance Decomposition of the N-Terminus via ESM2 Embeddings

To identify which regions of the P450 protein sequence carry functional information relevant to cluster discrimination, two complementary per-residue importance metrics—**PIVOT** (Perturbation-based Importance Vector Orientation via Transformers) and **DIVA** (Differential Vector Attribution)—were first computed across the full protein length for all 345 P450s. This genome-wide scan revealed a pronounced importance hotspot concentrated in the N-terminal region (section 3. 5). Based on this finding, and informed by the PEACE analysis (Section 2.6) establishing that the N-terminal 50 amino acids constitute a constraint-free scaffold, and by preliminary PIVOT/DIVA hotspot profiling identifying this region as carrying the strongest functional signal, subsequent high-resolution analyses were focused on positions 1-50.

#### 2.8.1 PIVOT: Perturbation-based Importance Vector Orientation via Transformers

If a single residue carries information that helps distinguish one P450 functional group from another, removing that residue’s contribution from the protein’s ESM2 representation should shift the representation in a way that confuses a classifier trained to recognize functional groups. PIVOT measures exactly this—how much a classifier’s confidence changes when one residue is “hidden” from the model. The method proceeds in two stages: a coarse-grained population-level scan to identify which broad regions of the P450 sequence carry functional information (Phase 1), followed by a fine-grained single-residue analysis to rank individual positions across the entire protein (Phase 2) (Figure 2). The complete analysis script is available on GitHub (https://github.com/qiaoweizhuo-pixel/P450/tree/main/PIVOT)

**Figure 2.**
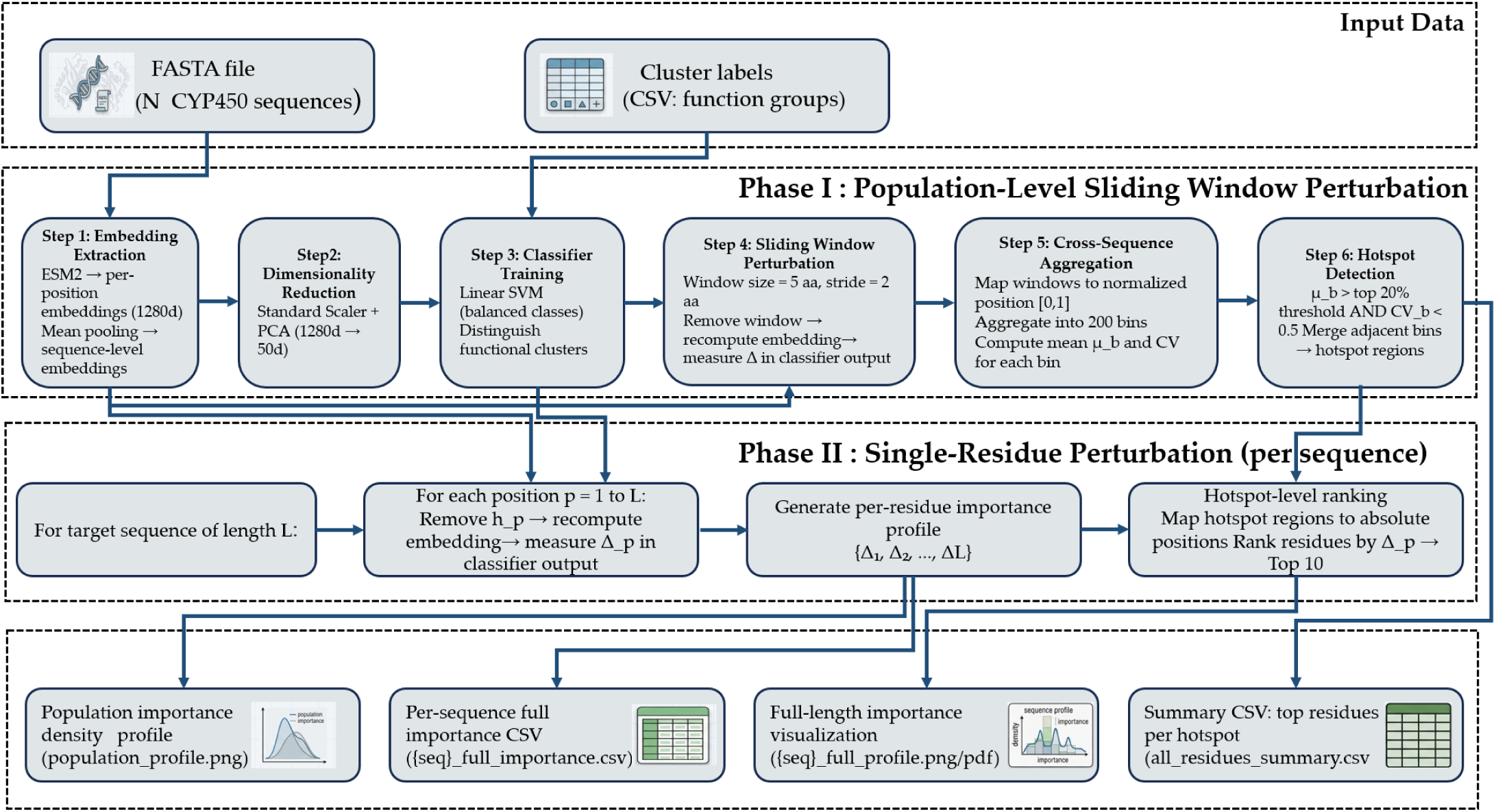
Schematic of the PIVOT pipeline. Phase 1: a length-normalized sliding-window perturbation scan across all 345 full-length P450 sequences identifies population-level importance hotspots (top 20% mean Δ1, CV < 0.5). Phase 2: single-residue omission is performed across the full-length sequences to quantify each position’s marginal contribution to SVM cluster discrimination, yielding per-residue importance scores. Key residues are then identified by ranking scores.

##### Phase 1 — Population-level hotspot identification

For each of the N = 345 full-length P450 sequences Si of length L, per-residue embeddings were extracted from the 33rd transformer layer of ESM2-650M (embedding dimension d = 1280). A global representation of each sequence, Vi(mean) è R^d^, was obtained by averaging the embeddings of all Li residues. After standardizing the per-protein representations (StandardScaler) and reducing dimensionality via PCA (k = 50 components), a linear SVM (LinearSVC, C = 1.0) was trained to discriminate the 12 ESM2-defined functional clusters (Section 2.5), producing a decision function f(v).

A sliding window of fixed size w = 5 residues (stride s = 2) was passed across each sequence. For each window covering residues j through j + w - 1, a perturbed representation was computed by averaging the embeddings of all residues except those in the window. The importance of the window was scored as the absolute change in the SVM decision function. Δ1 is large when the five omitted residues collectively shift the representation in a direction that changes the classifier’s decision.

To aggregate across proteins of different lengths, each window was assigned a normalized position p_norm_ è [0, 1] based on the midpoint of the window (0 = N-terminus, 1 = C-terminus). All windowed ΔÌ values from all 345 sequences were pooled into n = 200 equally spaced bins along this normalized axis, and the mean Δ1 and its coefficient of variation (CV) were computed per bin.

Hotspot regions were defined as contiguous bins satisfying two criteria: (i) the mean Δ1 ranked in the top 20% across all non-empty bins, and (ii) the CV was below 0.5, ensuring that high importance was consistently observed across the population rather than driven by a few outlier proteins.

##### Phase 2 — Single-residue importance quantification across the full protein

Phase 1 identified where the population-level signal is concentrated. Phase 2 asked: within each individual protein, which specific residues contribute most to functional cluster discrimination? To answer this, single-residue perturbation was performed across the entire length of each protein (all L positions), producing a complete per-residue importance profile.

For each position p in a protein of length L, the residue’s embedding was omitted from the mean-pooled representation. The importance of position p is the Li change in the SVM decision function.

###### Scaling behavior

Because the representations are mean-pooled, omitting a single residue shifts the mean. For a linear SVM with decision normal w (in the PCA-reduced space), the importance therefore scales with the factor 1/(L - 1), which is shared by all residues within a protein and therefore does not affect their relative ranking. The numerator is the biologically meaningful term—it measures how strongly residue p’s embedding deviates from the protein’s average embedding in the specific direction relevant to cluster discrimination. A residue receives a high PIVOT score not because it is intrinsically “important” in any absolute sense, but because its embedding is unusual in the respects that matter for distinguishing functional groups, and hiding it from the classifier therefore shifts the decision boundary by a larger amount than hiding a residue whose embedding is closer to the protein mean in the classification-relevant subspace.

This interpretation has an important practical consequence: PIVOT is a first-order sensitivity metric that identifies residues whose embeddings are unusually informative for functional classification in context, rather than residues that are universally conserved or catalytic. A residue that is highly similar in embedding space to the protein’s average (and therefore receives a low PIVOT score) may still be essential for structure or catalysis—it simply does not carry cluster-discriminating information within the ESM2 feature space.

To facilitate within-protein comparisons and cross-protein aggregation, raw Ip scores were min-max normalized to [0, 1] independently for each protein. A PIVOT score near 1 indicates that omitting that residue causes the largest perturbation to the classifier among all positions in that protein—i.e., the residue carries information that the classifier relies on to assign the protein to its correct functional cluster. Scores near 0 indicate residues whose omission has negligible effect.

#### 2.8.2 DIVA: Differential Vector Attribution

Every protein sequence, when processed by ESM2, is assigned a global representation (the CLS token) that summarizes its overall functional identity. This global representation can be thought of as the protein’s “address” in functional space. At the same time, the average of all per-residue embeddings captures the typical local features of that protein—its compositional baseline. The direction from this baseline to the global address—the specificity vector—points toward what makes this particular protein functionally distinct. DIVA asks: for each residue, how strongly does its local embedding align with the protein’s own specificity direction? A residue whose embedding points in the same direction as the specificity vector is likely to carry features that contribute to the protein’s functional identity.

DIVA is fully unsupervised: it requires no labelled data, no predefined functional clusters, and no classifier training. It is computed independently for each protein sequence using only the embeddings produced by ESM2 (Fig. 3). The complete analysis script is available on GitHub (https://github.com/qiaoweizhuo-pixel/P450/tree/main/DIVA)

**Figure 3.**
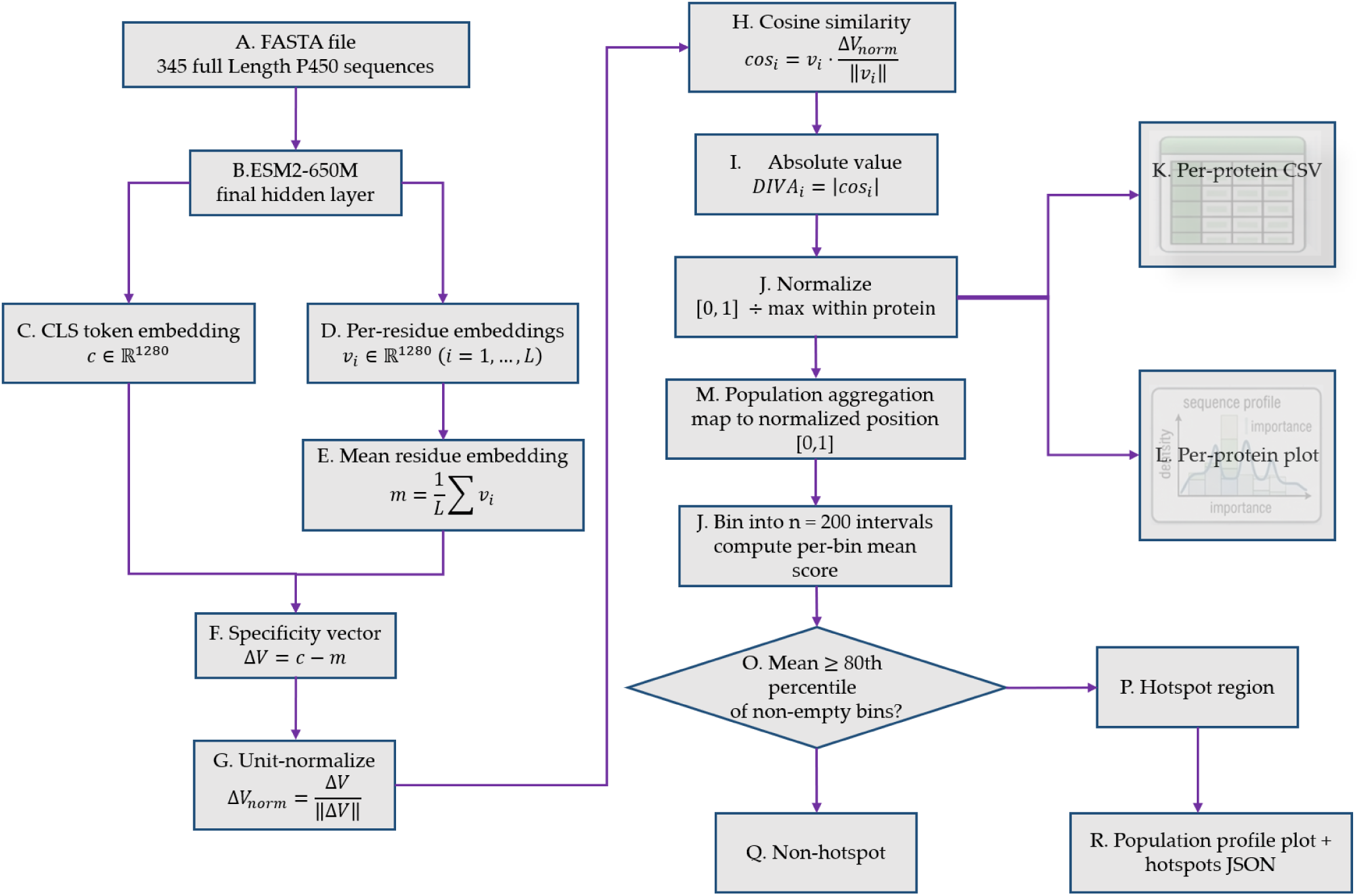
Schematic of the DIVA pipeline. For each protein sequence independently, ESM2-650M extracts the CLS token embedding (c), mean residue embedding (m̅), and per-residue embeddings (v*i*). The specificity vector is unit-normalized, and per-residue DIVA scores are computed as the absolute cosine similarity between each 巧 and the specificity direction. Scores are normalized to [0, 1] by dividing by the within-protein maximum.

##### Feature extraction

For each of the N = 345 full-length P450 sequences, embeddings were extracted from the final hidden layer of ESM2-650M (embedding dimension d = 1280; Lin et al., 2023). Three quantities were obtained per sequence: (i) CLS token embedding, c ∈ ℝ^d^ representing the global functional semantics of the entire P450 sequence; (ii) mean residue embedding, m̅ ∈ ℝ^d^, the average of all per-residue embeddings; and (iii) per-residue embeddings *v_i_* ∈ ℝ^d^ for all positions i = 1,…, L.

##### Specificity vector and DIVA score

The specificity vector, Δv, is defined as the difference between the global semantic representation and the average residue background: Δv = c - m̅. This vector points from “what this protein is made of, on average” toward “what this protein is, functionally.” After unit-normalizing the specificity vector, the DIVA score for residue i is computed as the absolute cosine similarity between its embedding and the specificity direction.

A high DIVA score indicates that residue i’s embedding is oriented along the same direction as the protein’s functional specificity vector—its local sequence features are aligned with what makes this protein functionally distinct. A low DIVA score indicates that the residue’s embedding is approximately orthogonal to the specificity direction—its features are not particularly relevant to this protein’s functional identity, at least as captured by ESM2.

Critically, DIVA does not measure whether a residue is conserved, hydrophobic, or catalytically important. It measures a more abstract property: whether the residue’s embedding—which encodes the residue and its sequence context—points in the same high-dimensional direction as the protein’s functional signature.

##### Normalization

For each protein, raw DIVA scores were normalized to [0, 1] by dividing by the maximum score within that protein. This per-protein normalization removes differences in the absolute magnitude of the specificity vector across proteins and focuses the analysis on the relative importance of residues within each protein.

##### Population-level hotspot detection

To identify regions of consistently high importance across the population, per-residue normalized DIVA scores from all sequences were mapped to normalized positions and binned into n = 200 equally spaced intervals. For each bin, the mean importance score was computed across all residue-position observations falling within that bin. Hotspot regions were defined as contiguous bins where the mean score exceeded the 80th percentile of all non-empty bin means. The resulting hotspot map identifies which regions of the P450 sequence most consistently carry features aligned with each protein’s functional specificity direction across the superfamily.

#### 2.8.3 Physicochemical Validation of DIVA Scores

To determine what biological signal DIVA scores capture—beyond their operational definition as cosine similarity between per-residue embeddings and the per-protein specificity vector—per-residue DIVA scores were tested for systematic associations with experimentally derived amino acid physicochemical properties. The biological signal resides not in the magnitude of any single correlation, but in the pattern of sign reversals across the three segments defined a priori from the known biophysics of ER signal-anchor insertion (von Heijne, 1989): a genuine topology template should produce, for example, positive DIVA-hydropathy correlations in the transmembrane helix that reverse to negative correlations at the cytosolic exit, whereas unstructured noise would show no systematic relationship to segment boundaries.

The N-terminal 50 residues were partitioned into positions 1-8 (pre-insertion), 9-40 (transmembrane helix core), and 41-50 (cytosolic exit bend). Four experimentally derived physicochemical properties were obtained from the AAindex database (Kawashima & Kanehisa, 2000; Kawashima et al., 2008): hydropathy (Kyte & Doolittle, 1982), helical propensity (Chou & Fasman, 1978), side-chain volume (Creighton, 1993), and aromaticity (binary: F, Y, W = 1; others = 0).

For each segment, Spearman rank correlations were computed between per-residue DIVA scores and each property across all residue-position observations (345 proteins × 50 positions = 17,250 observations). Bonferroni correction accounted for four simultaneous tests per segment. To evaluate whether the observed across- segment sign pattern could arise by chance, a permutation test (10,000 shuffles of position labels, preserving marginal distributions) assessed the joint configuration of correlation signs across all three segments and four properties.

Amino acids were classified by charge at pH 7.0 as positive (R, K), negative (D, E), polar uncharged (H, N, Q, S, T, C, Y), or nonpolar (A, V, L, I, M, F, P, G). Kruskal-Wallis tests compared DIVA score distributions across charge categories within each segment, with post-hoc one-sided Mann-Whitney U tests comparing charged versus uncharged residues.

To rule out the interpretation that DIVA merely recapitulates alignment variability, Shannon entropy H(p) was computed at each position across all 345 proteins, and its Spearman correlation with mean per-site DIVA scores was calculated within each segment.

#### 2.8.4 N-Terminal Importance Profile Clustering and Cross-Metric Comparison

To characterize how individual proteins differ in their N-terminal importance architecture, per-protein DIVA and PIVOT importance profiles were independently subjected to the N-terminal Importance Profile Clustering (NIPC) pipeline. The complete analysis script is available on GitHub (https://github.com/qiaoweizhuo-pixel/P450/tree/main/NIPC).

##### N-terminal window

For each protein of length L, the N-terminal window was defined as L_N = max(30, min(100, [L × 0.15])).

##### Classification: N-terminal-driven vs. Weak-N

Each protein was classified as “N-terminal-driven” or “Weak-N” (weak N-terminal dependency) based on two criteria. (i) Concentration: µ*N_tem_* / *µ*_full_ ≥ 1.0. (ii) Peak (mode-dependent): DIVA mode: max(l_N-term_) ≥ 0.85 × max(I_full_); PIVOT mode: max(I_N-term_) ≥ 1.5 × µ_N-term_. Proteins failing either criterion were designated Weak-N.

##### Profile resampling and clustering

N-terminal-driven profiles underwent Gaussian smoothing (σ = 1.0) and cubic interpolation to n = 50 points, followed by min-max normalization to [0, 1]. Pairwise distances were computed using correlation distance (d = 1 - r), and hierarchical clustering was performed with average linkage. The number of clusters k was determined automatically from the largest jump in successive merge distances among the last 10 merge steps, bounded between k_min_ = 3 and k_max_ = 6. Clusters were labeled Early-N, Mid-N, or Late-N according to mean peak position. Clustering was performed independently for DIVA and PIVOT profiles.

##### Cross-metric comparison

The sets of genes classified as Weak-N by DIVA and by PIVOT were compared for overlap. For each protein, the N-terminal peak position (positions 1-50) was defined as the residue with the maximum importance score, normalized to [0, 1]. DIVA and PIVOT peak positions were compared across all shared genes via Pearson (r) and Spearman (ρ) correlation; proteins with | Δ | ≤ 0.15 were classified as on-diagonal. For genes classified as N-terminal-driven by both metrics, DIVA and PIVOT cluster assignments were cross-tabulated.

#### 2.8.5 Statistical validation of PIVOT peak association with CYP families and bimodality analysis

To determine whether the N-terminal PIVOT peak position carries a genuine family-associated semantic signal, we systematically tested its association with CYP family identities. For each P450 sequence, the PIVOT peak position was defined as the residue with the maximum normalized score within the N-terminal 50 amino acids. Families with fewer than five members (n < 5) were excluded to ensure statistical robustness (minimum family size).

A Kruskal-Wallis (KW) test was performed to assess overall differences in peak positions across CYP families. The effect size was quantified using epsilon-squared (*ε*^2^), which estimates the proportion of variance in peak position explained by family identity. To rule out the possibility that the observed KW statistic (H) arose from random label assignment, a permutation test was conducted with 10,000 shuffles of the family labels. The empirical p-value was calculated as the proportion of permutations producing an *H_perm_* value greater than or equal to the observed *H_obs_*.

Given the significant omnibus result, Dunn’s post-hoc test with Holm correction for multiple comparisons was applied to identify pairwise family differences. The z-statistics and adjusted p-values were calculated using the rank-transformed peak positions. To further characterize the profiles, we introduced a bimodality detection algorithm. Based on the family-averaged PIVOT profiles, local maxima were identified using a sliding window (window size = 3). Secondary peaks were defined as local maxima separated by at least 15 residues from the global peak and exceeding 80% of the global peak’s magnitude (*Strict bimodal*), or between 70% and 80% (*Relaxed bimodal*). Families failing these criteria were classified as *Unimodal*. This classification allowed us to visualize representative family profiles across different architectural tiers. All statistical analyses were performed using custom Python scripts leveraging the scipy.stats and statsmodels libraries.

### 2.9 Validation against DeepLoc-2.1 subcellular localization

To test whether the N-terminal features decomposed by DIVA and PIVOT are recognized by an independent localization predictor, the DeepLoc-2.1 predictions from Section 2.7 (ER vs. Non-ER classification, plus continuous probability scores) were compared against DIVA and PIVOT metrics across all 345 proteins.

For each protein, mean DIVA and PIVOT scores were computed over three N-terminal segments (positions 1-8, 9-40, and 41-50) plus the full window (1-50). DIVA and PIVOT peak positions were defined as the residue within positions 1-50 bearing the maximum importance score. Segment means and peak positions were compared between ER (n = 328) and Non-ER (n = 17) proteins via two-sided Mann-Whitney U tests. Spearman rank correlations (ρ) were computed between twelve DIVA and PIVOT metrics—peak positions, segment means, the DIVA 1-8/9-40 ratio, and the PIVOT coefficient of variation over positions 1-50—and the continuous DeepLoc-2.1 probability scores for Endoplasmic reticulum, Transmembrane, Peripheral, and Soluble.

Proteins classified as DIVA Early-N and DIVA Weak-N (Section 2.8.4) were tested for differential enrichment in ER versus Non-ER via Fisher’s exact test (two-sided). Kruskal-Wallis tests assessed whether the continuous ER probability score differed across DIVA and PIVOT cluster assignments, restricted to clusters containing ≥3 proteins. The DeepLoc-2.1 “Signals” annotation was cross-tabulated against cluster assignments, and proteins lacking any predicted signal peptide or transmembrane domain were examined individually.

Three summary metrics quantified the degree to which each protein’s DIVA profile conforms to the tripartite topological template of an N-terminal signal-anchor:(i) TM ratio = mean DIVA(9-40)/ mean DIVA(1-50); (ii) TM → Exit drop = mean DIVA(9—40)- mean DIVA(41-50); and (iii) Charge ratio = mean DIVA(41-50) / mean DIVA(9-40). These three metrics, together with DIVA and PIVOT peak positions and segment means, were compared between ER and Non-ER proteins via two-sided Mann-Whitney U tests.

## 3. Results

### 3.1 A Landscape of Functional Diversity: Identification and Phylogeny of the P450 Superfamily in *S. tonkinensis*

Comprehensive analysis of the *S. tonkinensis* genome led to the initial identification of 381 cytochrome P450 proteins. After removing completely duplicated sequences, a final set of 345 non-redundant cytochrome P450 proteins was obtained for further analysis. A maximum-likelihood phylogenetic tree reconstructed from their sequences revealed considerable diversity, classifying these proteins into 8 clans, 35 families, and 57 subfamilies (Fig. 4; Supplementary Table S1), indicating a complex evolutionary history conducive to functional diversification.

**Figure 4.**
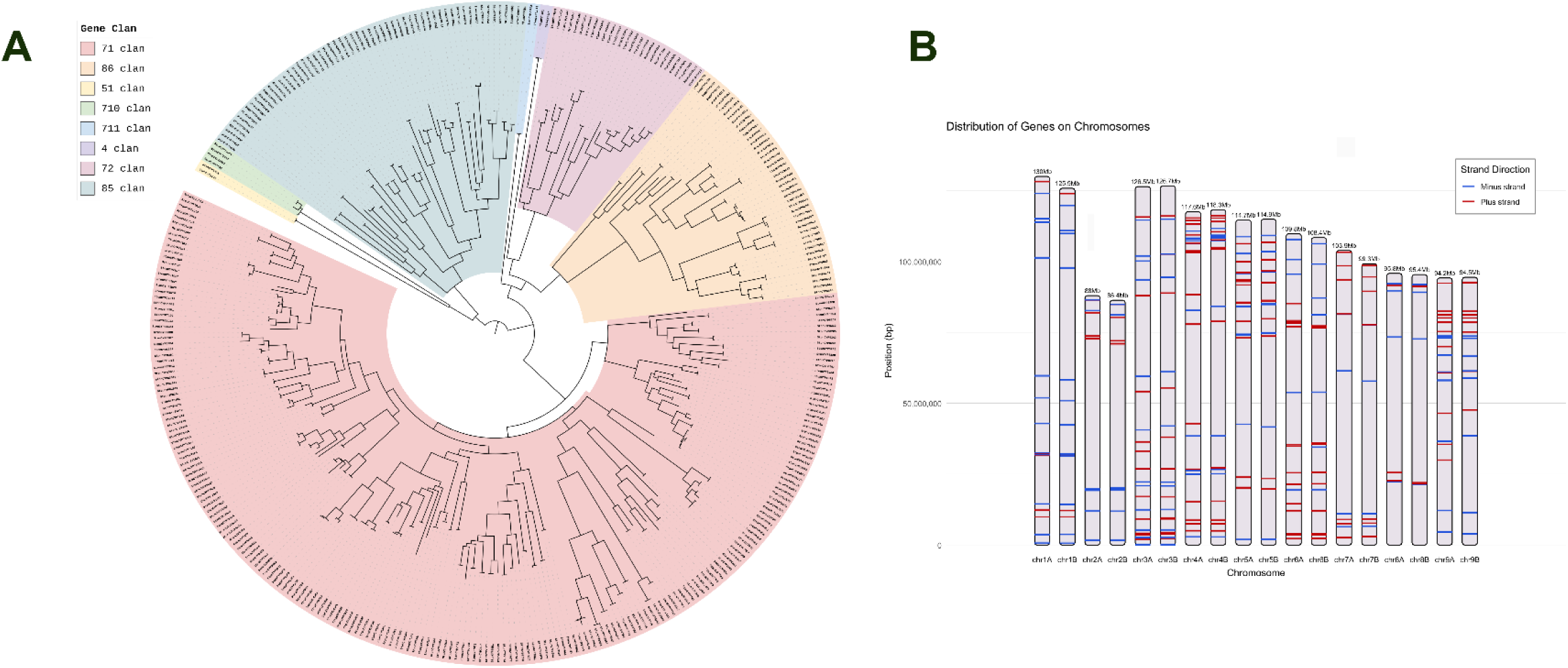
Comprehensive analysis of the P450 gene family. **A.** Phylogenetic tree of the P450 supergene family. Different colors represent distinct clans (major evolutionary branches), illustrating the evolutionary relationships among members. **B.** Chromosomal distribution of P450 genes. Gene positions are mapped across 18 chromosomes, with red bars indicating genes on the positive strand and blue bars indicating genes on the negative strand.

Conserved motif analysis using MEME identified ten significant motifs across the family: five 21-amino acid motifs, three 15-amino acid motifs, and two 11- amino acid motifs (Supplementary Figure S2). Gene structure and promoter cis-element analyses are provided in Supplementary Figure. S3 and S4, respectively. Promoter analysis indicated an enrichment of light-responsive elements, followed by elements related to methyl jasmonate (Me-JA) and abscisic acid (ABA) signaling (see Supplementary Table T1).

Genomic distribution analysis showed that P450 genes are located on all 18 chromosomes (9 homologous pairs) of *S. tonkinensis*, though their number is not strictly correlated with chromosome length (Fig. 4). Chromosome 5 harbored the highest number (31 genes), including several tightly clustered tandem arrays, whereas chromosome 2 and the 8B homoeolog contained the fewest (11 and 10 genes, respectively). Notably, the 8B chromosome lacked two P450 genes present on its 8A homoeolog.

Intra-genomic synteny analysis revealed that collinear P450 gene pairs are predominantly located on homologous chromosomes, with a minor fraction involved in cross-homoeolog duplications (Fig. 5A). Comparative analysis with *Arabidopsis thaliana* (a distantly related species) (Fig. 5B) and the closely related legumes *Arachis hypogaea* (peanut) and *Glycine max* (soybean) provided evolutionary context (Fig. 5B). While only two homologous chromosomal regions with limited synteny (containing one P450 homolog) were detected with *A. thaliana*, more extensive conservation was observed with peanut (three syntenic blocks) and soybean (approximately 3.5 syntenic blocks, containing at least five P450 homologs).

**Figure 5.**
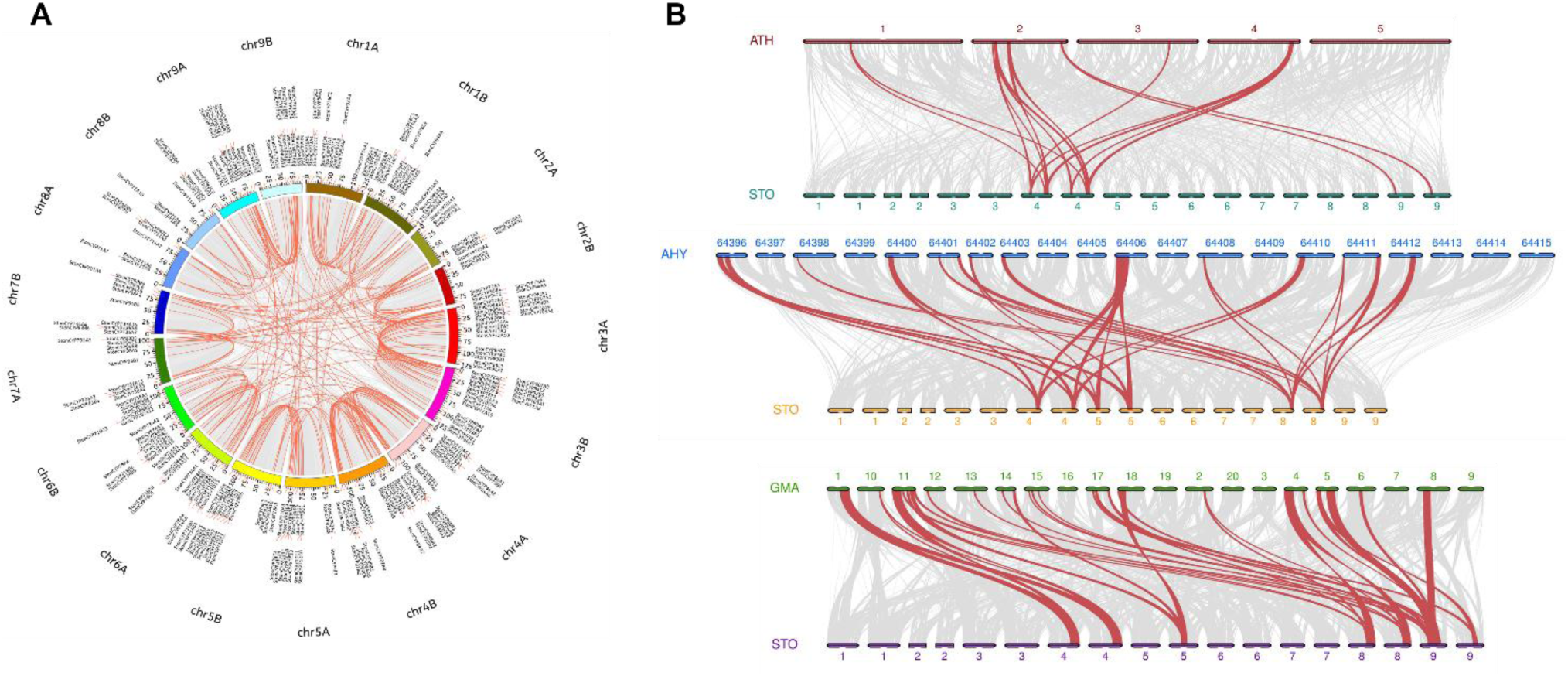
Synteny analysis of the P450 gene family within *S. tonkinensis* and across related species. **A.** Intra-genomic synteny analysis of *S. tonkinensis*. The Circos plot illustrates collinear blocks of P450 genes across different chromosomes within the *S. tonkinensis* genome. **B.** Comparative genomic synteny analysis between *S. tonkinensis* and three reference species. The dot plots, generated using JCVI, visualize collinear regions between the genomes of *S. tonkinensis* (STO) and *Arabidopsis thaliana* (ATH), *Glycine max* (soybean,GMA), and *Arachis hypogaea* (peanut, AHY). Gray lines connect syntenic blocks derived from whole-genome alignment, while red lines specifically highlight syntenic blocks containing genes from the P450 family.

### 3.2 Decoupling of Function from Phylogeny Revealed by Protein Language Modeling

To independently assess functional relationships beyond sequence homology, we projected the entire P450 repertoire into a semantic feature space using the ESM2 protein language model. Subsequent UMAP visualization revealed distinct clusters based on inferred functional similarity (Fig. 6). Quantitative comparison between these ESM2-derived functional clusters and clusters defined by monophyletic phylogenetic groups showed only moderate congruence (V-measure = 0.520, Adjusted Mutual Information = 0.480). A Sankey diagram further illustrated substantial functional reassignment of sequences between the two clustering schemes, highlighting that phylogenetic proximity is not a reliable predictor of functional similarity (Fig. 7A). Most strikingly, this analysis identified 13 phylogenetically adjacent sister gene pairs that were assigned to distinct functional clusters by ESM2 (Fig. 7B, C), providing concrete examples of recent functional divergence.

**Figure 6.**
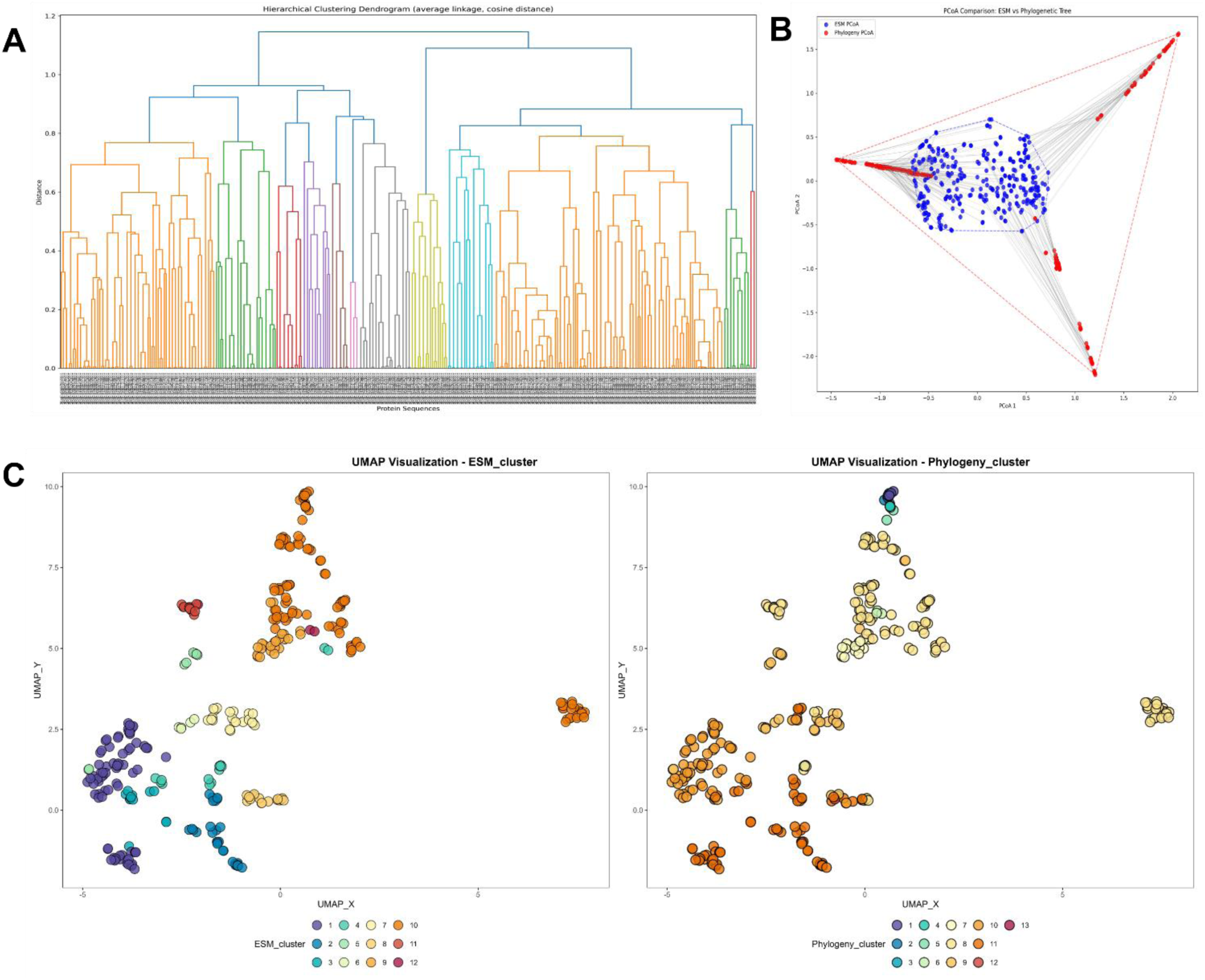
Comparative analysis of protein classification using evolutionary tree clustering and ESM2 embeddings. **A.** Hierarchical clustering dendrogram of P450 proteins based on ESM2 embeddings. The tree is colored to represent 12 distinct clusters derived from the clustering analysis. **B.** Principal Coordinates Analysis (PCoA) comparing the distribution of two classification schemes: red points represent clusters derived from the full-length sequence phylogenetic tree, and blue points represent clusters derived from the hierarchical clustering of ESM2 embeddings. **C, D.** UMAP visualization of the high-dimensional ESM2 embeddings for all P450 proteins. In (C), the same points are colored according to the 12 clusters defined by hierarchical clustering of the ESM2 embeddings. In (D), points are colored according to the 13 monophyletic clades defined by phylogenetic tree analysis, facilitating a direct visual comparison of the two classification approaches.

**Figure 7.**
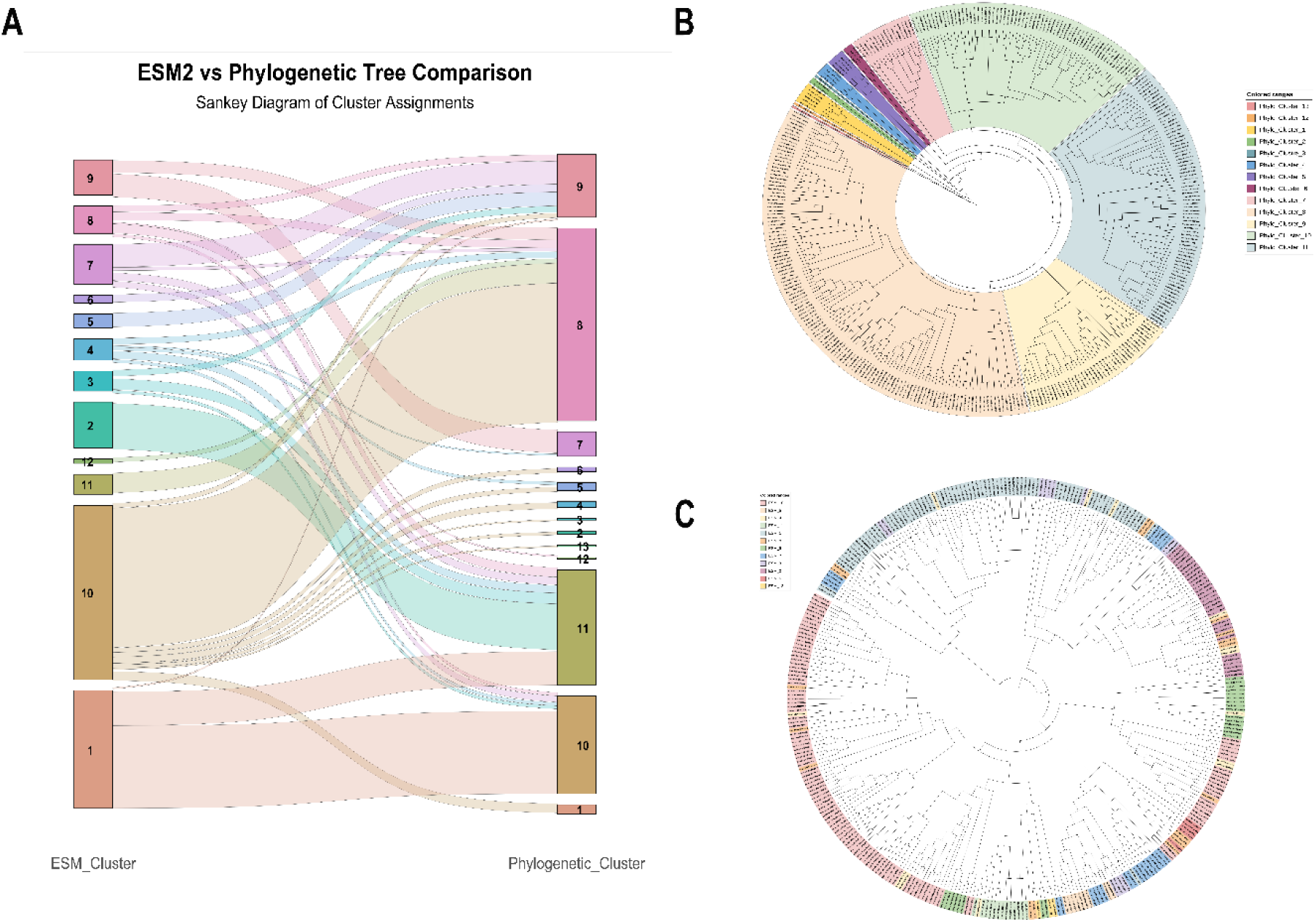
Correspondence and divergence between phylogenetic clustering and ESM2 embedding-based clustering. **A.** Sankey diagram illustrating the mapping of protein members between the 12 ESM2 clusters and the 13 phylogenetic clusters. The width of each flow represents the number of proteins shared between a specific pair of clusters from the two classification schemes. **B.** Phylogenetic tree of P450 proteins, with branches colored according to the 13 defined phylogenetic clusters. Each colored group corresponds to a distinct, monophyletic clade. **C.** The same phylogenetic tree as in (B), but with branches colored according to their assignment to the 12 ESM2 clusters. This visualization reveals instances where evolutionarily proximate branches in the tree are assigned to different functional clusters by the ESM2 embedding analysis, highlighting potential functional divergence within closely related clades.

### 3.3 N-terminal Sequence Variation as a Primary Driver of Functional Reassignment

To investigate the molecular basis for the functional divergence observed in the 13 sister pairs (Figure 7C), we performed detailed multiple sequence alignments for each pair. This analysis revealed a consistent and striking pattern: the most prominent sequence differences were heavily concentrated in the N-terminal region, frequently manifesting as extensive deletions, insertions, or a high density of amino acid substitutions. In contrast, the core catalytic domains remained relatively conserved (the complete alignment files are provided as text files in the Full Alignments within the Supplementary Materials). This strong association between dramatic N-terminal sequence variation and ESM2-based functional reassignment suggests that alterations in this region are a primary driver of functional diversification among recently duplicated P450s, providing a clear target for further mechanistic investigation.

### 3.4 PEACE: Alignment-free constraint estimation reveals a neutral-relaxation landscape punctuated by a single PxxG structural hinge

#### 3.4.1 Motivation and dual-mode design

Multiple sequence alignment (MSA) is the standard substrate for residue-level constraint inference via dN/dS or Shannon entropy. However, the P450 N-terminus is densely populated with insertions and deletions (indels), making column-wise MSA alignment in this region intrinsically unreliable. Positions that appear variable by entropy may merely reflect alignment ambiguity rather than genuine evolutionary relaxation; conversely, columns that appear conserved may reflect alignment compression rather than purifying selection. This problem is particularly acute for the first ∼50 amino acids, where >60% of sequences carry length polymorphisms relative to any single reference.

To circumvent MSA artifacts, we developed PEACE (Per-position Embedding Alignment-free Constraint Estimation), which replaces column-wise alignment with soft correspondence in ESM2 embedding space. Concretely, for each reference position *i*, PEACE identifies the query sequence position whose embedding vector has maximal cosine similarity to the reference embedding at position *i*. This yields a soft mapping that respects local sequence context without enforcing rigid columnar alignment. Per-position constraint is then quantified as *dispersion*—the complement of mean match similarity (1 - mean cosine similarity), ranging from 0 (perfectly constrained) to 1 (maximally relaxed).

PEACE was implemented in two complementary modes:

##### PEACE-REFERENCE

All query sequences are soft-corresponded against a single median reference sequence (StonChr6AG199490.1, chosen as the embedding-space centroid of the *S. tonkinensis* P450 family). This mode provides a fixed coordinate system for comparing constraint magnitudes across positions and is the primary mode for island detection.

##### PEACE-DTW

A consensus sequence is constructed via DTW Barycenter Averaging (DBA, window size = 11; Petitjean et al., 2011) over all 345 sequences, iteratively refined (max 10 iterations). All sequences are then soft-corresponded to this consensus. This mode is fully reference-free and captures constraint patterns that are robust to the choice of reference.

Both modes operate on ESM2 (t33_650M_UR50D) embeddings projected to 50 PCA dimensions for computational efficiency, and both incorporate FDR correction (Benjamini-Hochberg) for phylogenetic permutation tests of directional selection.

#### 3.4.2 Global landscape: The N-terminus is under pervasive neutral relaxation

In PEACE-REFERENCE mode, the reference sequence StonChr6AG199490.1 (509 residues) was automatically selected as the median-length representative of the family (Figure 8A). The mean dispersion across all 509 aligned positions was 0.542 ± 0.122. Strikingly, the N-terminal region (reference positions with mean soft-correspondence position ≤ 50; effectively the first ∼51 residues) showed significantly higher dispersion than the catalytic core: 0.622 ± 0.089 versus 0.533 ± 0.115 (Δ = +0.089, Mann-Whitney p = 1.12 × 10^-7^; Figure 8). Approximately 84% of N-terminal residues exhibited dispersion values above the global mean, consistent with a regime of pervasive neutral relaxation. In contrast, the core catalytic domain was dominated by low-dispersion positions—the six most constrained residues (dispersion range 0.142-0.311) all mapped to functionally essential catalytic motifs (EXXR, FxxGxxxCxG, PERF) in the heme-binding region (positions 441-448)(Werck-Reichhart & Feyereisen, 2000), confirming that PEACE correctly recovers known structural constraints (Supplementary Table T2).

**Figure 8.**
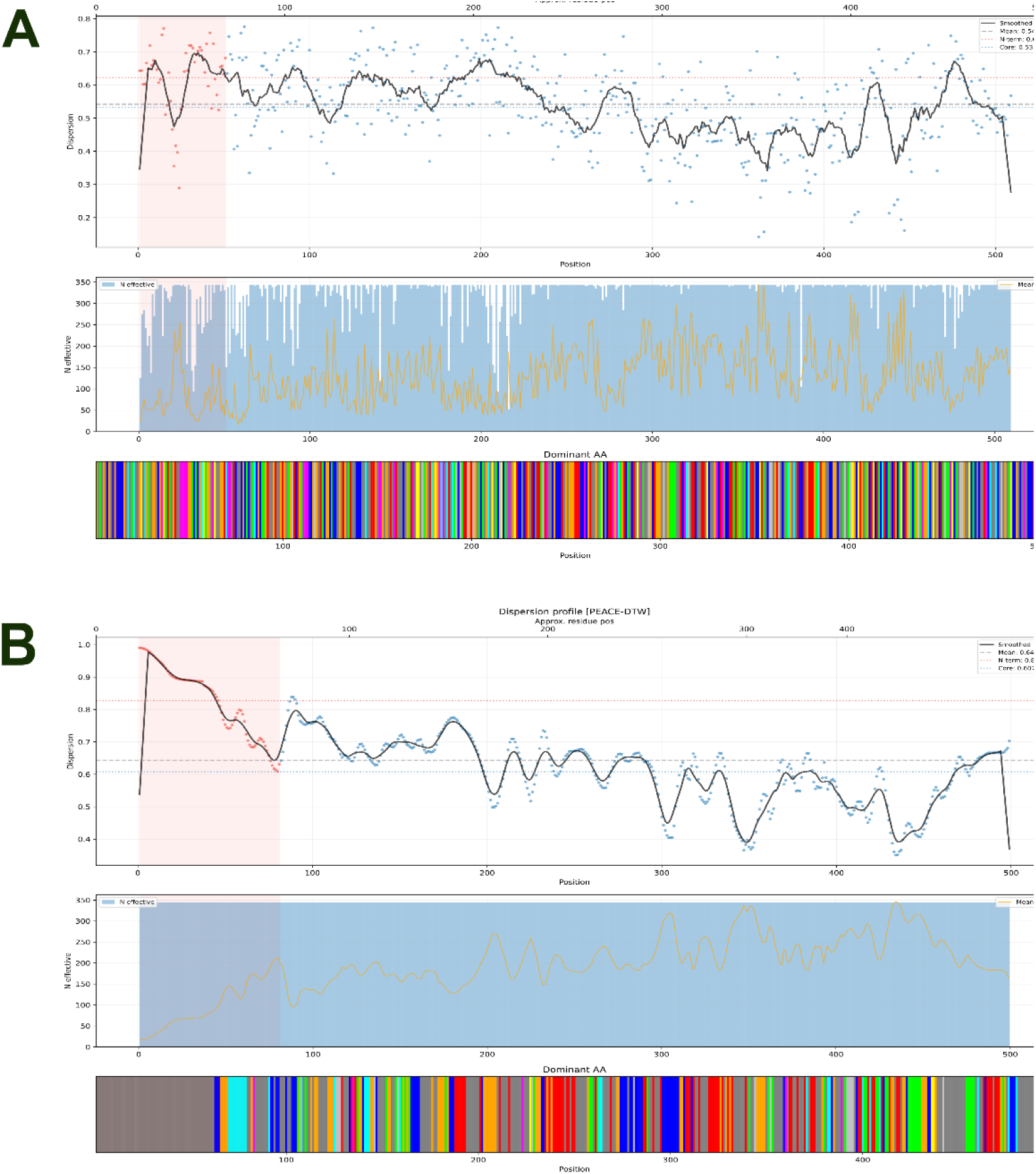
Dispersion profiles of protein sequences generated by the PEACE algorithm. **A.** Dispersion profile in Reference mode, which employs soft correspondence against a median-length reference sequence. **B.** Dispersion profile in DTW (Dynamic Time Warping) mode, which constructs a consensus sequence via DBA (DTW Barycenter Averaging) from windowed embeddings, with the N-terminal region highlighted in the red shaded area. Both panels consist of three aligned subplots from top to bottom: (1) Dispersion vs. Position: Raw dispersion scores (blue dots) along with a smoothed trend line (black solid). Horizontal dashed lines indicate mean and regional thresholds. The red shaded area highlights the N-terminal or active residue region (defined by the n_term_boundary parameter, default ≤ 50). (2) N_effective vs. Position: Displays the effective number of sequences aligning to each position (blue area), overlaid with the mean match similarity (orange line). (3) Dominant AA: A color-coded bar plot showing the most frequent amino acid (dominant_aa) at each aligned position. Algorithmic differences: Reference mode aligns all target sequences to a single reference sequence using thresholded cosine similarity, whereas DTW mode uses a window-sliding trajectory and PCA-reduced vectors to compute a DTW consensus, followed by position-wise embedding aggregation.

PEACE-DTW amplified this signal. With the DTW consensus as reference, mean N-terminal dispersion rose to 0.827 versus 0.608 for the core (Δ = +0.219, Mann-Whitney p = 2.53 × 10^-33^). The extreme N-terminus (consensus positions 1-10) was essentially unalignable—dispersion exceeded 0.96 at every position, with dominant amino acids defaulting to leucine as the algorithmic fallback when no genuine consensus exists (Figure 8B). Eighteen N-terminal positions in DTW mode exceeded the mean + 2σ threshold for extreme variability, compared to zero in REFERENCE mode, reflecting DTW’s greater sensitivity to length heterogeneity at the very N-terminal tip. Critically, both modes converged on the same qualitative conclusion: the N-terminal 50 amino acids constitute a neutral-relaxation landscape, sharply distinct from the strongly constrained catalytic domain (Figure 8).

Validation against sequence-level Shannon entropy confirmed that PEACE dispersion captures constraint information independent of amino acid frequency distributions. Dispersion and Shannon entropy were only weakly correlated (Spearman ρ = 0.124 in REFERENCE mode; ρ = 0.187 in DTW mode), and partial correlation analysis showed that N-terminal elevated dispersion persisted after controlling for Shannon entropy (REFERENCE: N-term residual = +0.086, core residual = -0.010, Mann-Whitney p = 2.06 × 10^−8^; DTW: N-term residual = +0.192, core residual = -0.037, p = 7.48 × 10^-37^). Thus, PEACE dispersion captures embedding-level structural constraint that is not reducible to amino acid conservation alone (Supplementary Figure S5).

#### 3.4.3 A single constrained island: The PxxG structural hinge at positions 20-27

Against this neutral-relaxation background, PEACE-REFERENCE revealed exactly one constrained island within the N-terminal 50 amino acids: positions 20-27, with the reference sequence motif **W^20^PVVGMLP^27^**(Figure 8A). The constraint profile of this octapeptide defines a characteristic “PxxG” pattern (Table 1).

**Table 1.** Residue-level dispersion and conservation metrics for the PxxG structural hinge (positions 20-27). Note: Pro^21^ (Dispersion = 0.356, near-absolute conservation) and Gly^24^ (Dispersion = 0.289, near-absolute conservation) act as the rigid “constrained spine” of the hinge, while the flanking residues (positions 22-23 and 25-27) display moderate-to-high variability, characteristic of the flexible flanks surrounding a structural hinge.

| Position | Dispersion | Shannon | Dominant residue | Conservation |
| --- | --- | --- | --- | --- |
| 20 | 0.466 | 2.069 | W (41%), L (38%) | Moderate |
| **21** | **0.356** | **0.149** | **P (98%)** | **Near-absolute** |
| 22 | 0.417 | 2.051 | I (46%), V (20%), L (19%) | Low–moderate |
| 23 | 0.397 | 1.674 | I (57%), V (24%) | Low–moderate |
| **24** | **0.289** | **0.101** | **G (99%)** | **Near-absolute** |
| 25 | 0.526 | 2.467 | N (46%), H (14%), M (14%) | Low |
| 26 | 0.534 | 0.675 | L (89%) | High |
| 27 | 0.641 | 2.521 | P (35%), H (31%) | Low |

The two anchoring residues—Pro^21^ (dispersion = 0.356, 98% Pro) and Gly24 (dispersion = 0.289, 99% Gly)—are the most constrained residues in the entire N-terminal 50 amino acids and among the most constrained anywhere outside the catalytic core. Their near-absolute conservation, embedded within a flanking context of moderate-to-high variability (positions 22-23 and 25-27), creates a distinctive “constrained spine within flexible flanks” architecture characteristic of a structural hinge: the Pro introduces a kink via backbone φ/ψ restrictions, the Gly provides the conformational flexibility that defines the hinge point, and the intervening hydrophobic residues (V/I^22^, V/I^23^) anchor the motif within the membrane-proximal environment. Leu^26^ (89% conserved) extends the hydrophobic register. The PxxG pattern matches the canonical Pro-rich hinge motif (PxxP/G) found at the junction between the N-terminal transmembrane anchor and the globular catalytic domain in eukaryotic P450s (Williams et al., 2000).

Importantly, no other residue in the N-terminal 50 amino acids approached this level of constraint. The next most constrained N-terminal position after the PxxG island was position 38 (Tyr, dispersion = 0.681), nearly twice the dispersion of Gly^24^. This binary landscape—a single constrained island in an otherwise relaxed N-terminus—is the central finding of the PEACE analysis.

#### 3.4.4 Soft correspondence validation: The PxxG hinge maps to positions 44-51 in most sequences

A defining feature of PEACE is that constraint is measured in embedding space, not sequence space. The “positions 20-27” refer to the reference sequence StonChr6AG199490.1. To determine where the PxxG motif actually resides in each query sequence, we extracted the soft-correspondence mapping for each of the eight island residues across all 344 non-reference sequences (Supplementary Table T3).

The results confirmed that the PxxG island is genuinely N-terminal in the overwhelming majority of sequences, but its absolute sequence position is systematically shifted downstream relative to the reference (Figure 9):

**Figure 9.**
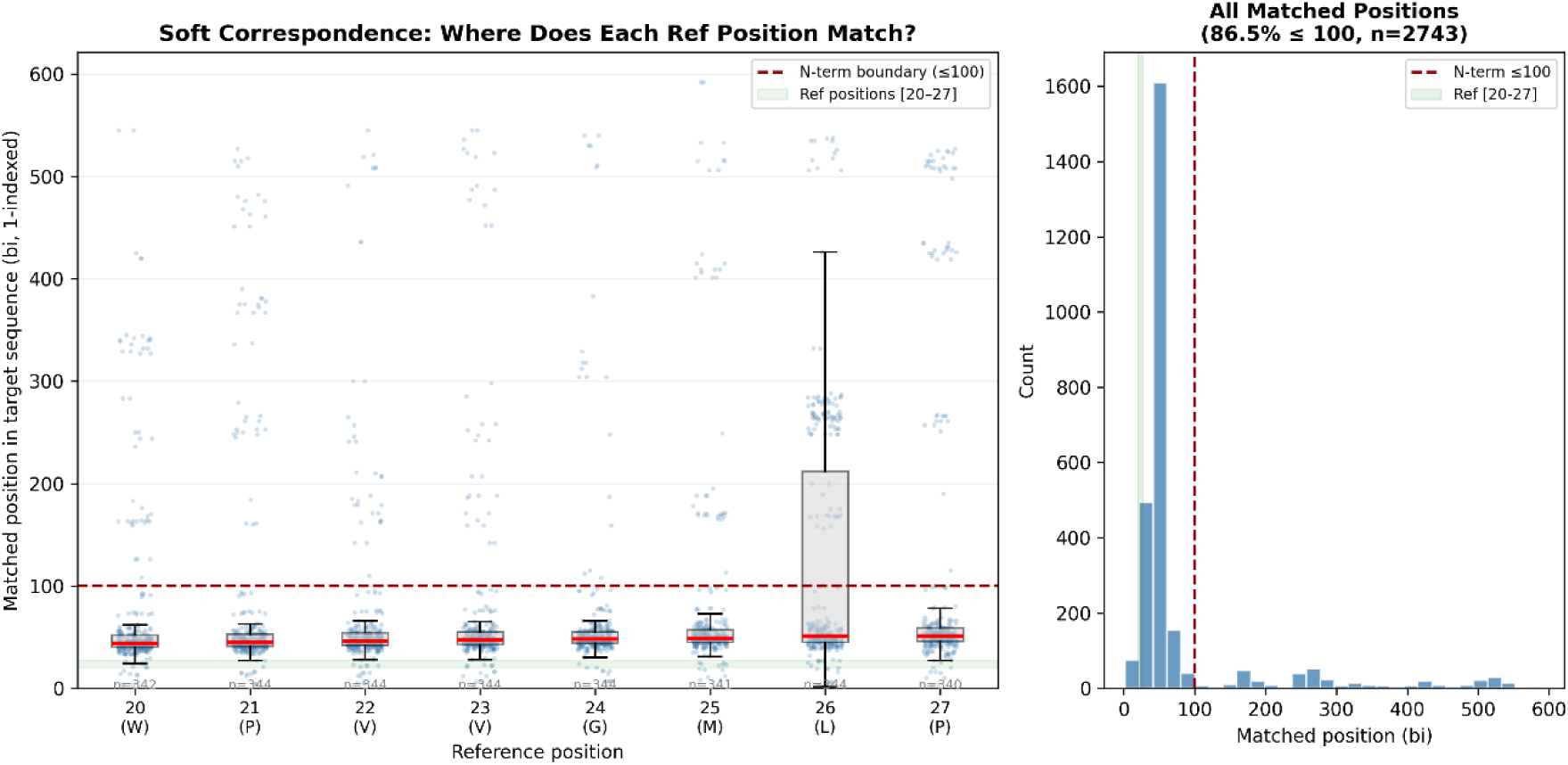
Validation of the N-terminal localization for the PxxG constrained island (positions 20 - 27) via soft correspondence mapping. A. Strip and box plot showing the distribution of matched positions (bi, 1-indexed) in target sequences for each reference position (20-27). Individual soft-matched positions are shown as steelblue dots, boxplots indicate the interquartile range, and red lines mark the median. The red dashed line indicates the defined N-terminal boundary (≤100), while the green shaded band marks the reference positions [20-27]. The number (n) of matched target sequences for each reference position is annotated above the x-axis. B. Overall histogram of all matched positions across the island (total n = 2,743). The red dashed line marks the N-terminal boundary. The inset text indicates that 86.5% of the matched positions fall within the first 100 residues of the target sequences. These results demonstrate that the PEACE-REFERENCE soft correspondence reliably maps the 8-residue PxxG hinge to the corresponding N-terminal region of all paralogous sequences, despite the presence of indel variations.

Several features of this distribution are notable. First, the median match positions span 44-51, representing a systematic downstream shift of approximately 24 residues relative to the reference positions 20-27 (Table 2). This shift reflects the cumulative effect of N-terminal indels: the reference sequence has a relatively compact N-terminus, whereas most query sequences carry insertions in the region upstream of the PxxG hinge. Second, Gly^24^—the most constrained residue—is also the most reliably positioned: 94.2% of its matches fall within the N-terminal 100 amino acids, and its median position (48) has the tightest interquartile range. This is consistent with Gly24 serving as the structural anchor of the hinge: its embedding signature is sufficiently distinctive that PEACE can localize it with high confidence even across substantial sequence divergence. Third, Leu^26^ shows the lowest N-terminal localization rate (70.3%) and the highest match position variance (SD = 123.5), reflecting the fact that hydrophobic residues have more ambiguous embedding signatures and generate more false-positive soft correspondences at downstream positions.

**Table 2.** Soft-correspondence mapping of the PxxG hinge residues from the reference sequence to non-reference sequences.

| Reference position | Reference AA | Median match position | Dominant AA frequency (%) | % in N-terminal (≤100 aa) |
| --- | --- | --- | --- | --- |
| 20 | W | 44 | 41% | 86.8% |
| **21** | **P** | **45** | **98%** | **87.2%** |
| 22 | V | 46 | 46% | 89.8% |
| 23 | V | 47 | 57% | 89.8% |
| **24** | **G** | **48** | **99%** | **94.2%** |
| 25 | M | 49 | 46% | 88.6% |
| 26 | L | 51 | 89% | 70.3% |
| 27 | P | 51 | 35% | 85.6% |

At the sequence level, 89.0% of the 344 non-reference sequences (306/344) had their PxxG core (P^21^ and G^24^) both soft-corresponded to positions ≤ 100, confirming that the constrained island is a genuine N-terminal feature in the vast majority of *S. tonkinensis* P450s. The 11% of sequences where the PxxG was not recovered in the N-terminus predominantly involved Pro^21^ mismatches: Pro has a relatively generic embedding signature, making its soft correspondence intrinsically noisier than that of Gly24.

#### 3.4.5 Cross-species conservation of the PxxG N-terminal hinge

To assess whether the PxxG N-terminal hinge is a general feature of plant P450s or a S. tonkinensis-specific phenomenon, we performed a computational hinge scan across nine land plant species spanning bryophytes to eudicots. For each species, P450 sequences were retrieved from UniProtKB (downloaded 2026-08-06) via the REST API using taxonomy-restricted queries with the “cytochrome P450” protein family annotation (fallback: EC 1.14.-.- or gene symbol CYP*; The UniProt Consortium, 2025); for *S. tonkinensis*, the 345 P450 proteins identified in Section 2.1 were used. All sequences were searched for the core PxxG motif (P-X-X-G, where the first position must be Pro and the fourth must be Gly) within the N-terminal 60 residues, with matches further validated by the strict PxxGxL extended motif where possible. The proportion of P450 sequences carrying an N-terminal PxxG hinge, and the position distribution of matched hinges, were quantified for each species (Table 3; Supplementary Tables T4 and T5).

**Table 3.**
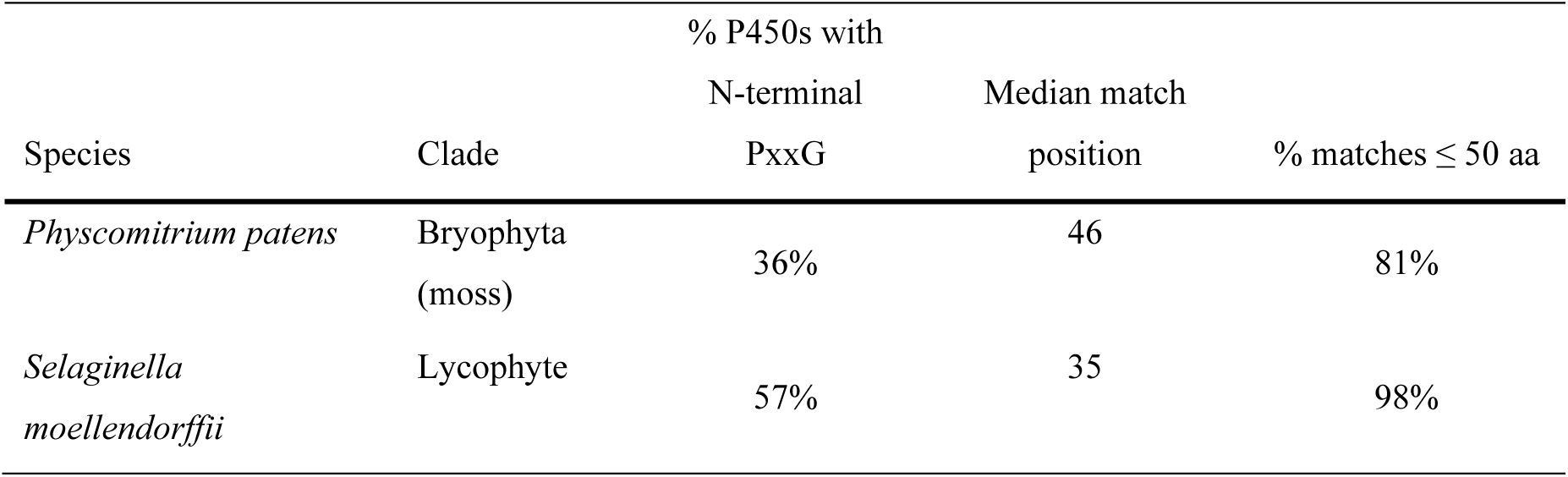
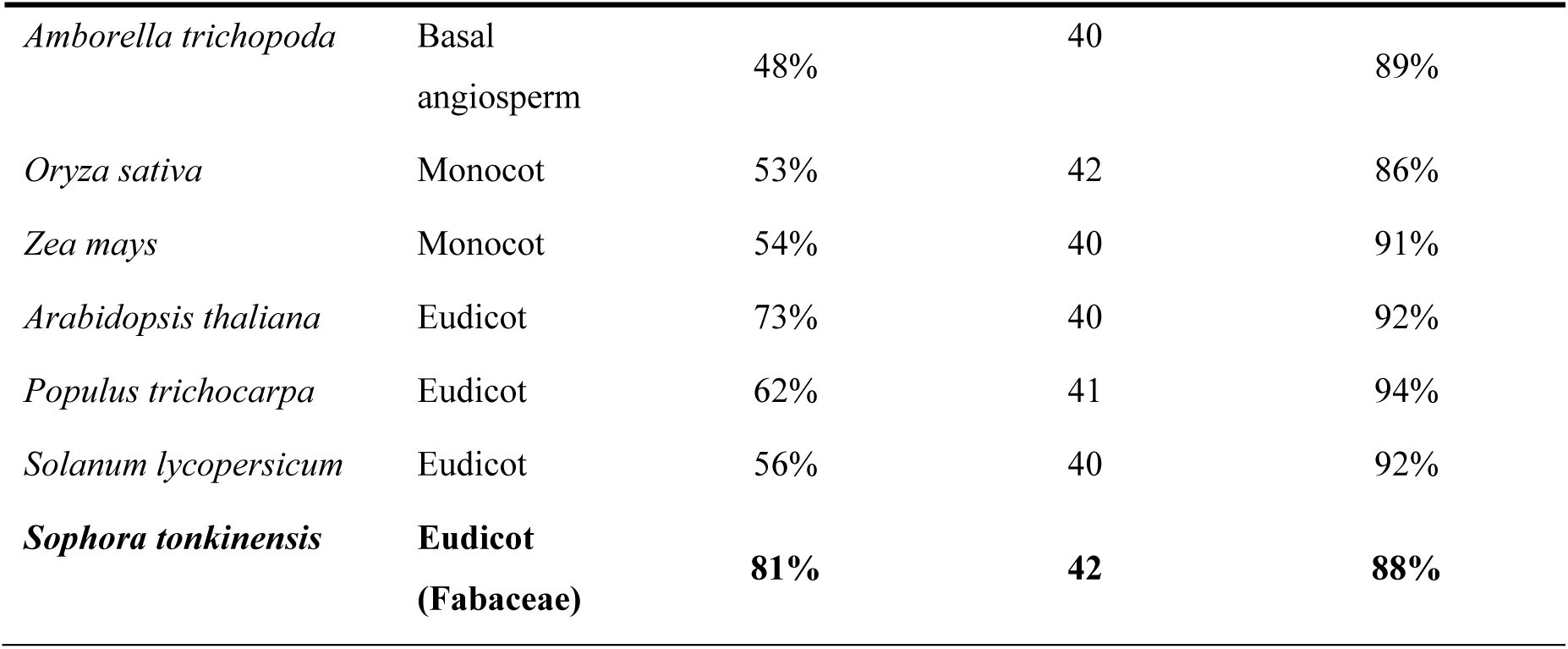
Prevalence and median match positions of the N-terminal PxxG motif across plant species. Match positions are 0-based; “N-term PxxG” denotes a P-X-X-G match within the first 60 residues of the native sequence, with the distribution of positions (median and %≤50) reported to exclude spurious internal hits

In Physcomitrium patens (moss), only 36% of P450s carried an N-terminal PxxG motif within the first 60 residues (median match position 46), indicating that the hinge has not been fixed as an obligate N-terminal feature at the base of land plants. In vascular plants, fixation rose markedly, to 48-73%, with tightly clustered median match positions (35-42) and the large majority (81-98%) of matches falling within the N-terminal 50 amino acids—consistent with a high-quality N-terminal hinge in most vascular plant P450s. Notably, *Selaginella moellendorffii* (lycophyte) showed a higher fixation rate (57%) than the basalmost angiosperm *Amborella trichopoda* (48%). This pattern is most parsimoniously explained by the extensive P450 family expansion in *Selaginella* (417 P450s vs. 220 in *Amborella*): the hinge appears to have become widely fixed early in vascular plant evolution, with subsequent diversification of eudicot families showing variable fixation (56-73%) rather than a continuous increase.

*S. tonkinensis* was the clear outlier: 81% of its P450s carried an N-terminal PxxG hinge by direct sequence-level motif scanning within the first 60 residues— substantially higher than any other species examined—consistent with the 89% recovery rate obtained by PEACE embedding-level soft correspondence (Section 3.4.4). These two estimates are complementary rather than conflicting, as they derive from distinct detection criteria: PEACE soft correspondence localizes the PxxG core through embedding similarity without imposing a fixed positional window, with recovery assessed within the N-terminal 100 residues (Section 3.4.4), whereas the sequence-level estimate requires strict P-X-X-G pattern matching restricted to the first 60 residues. The ∼8-percentage-point difference therefore reflects two factors: hinges located between positions 61-100 that are recovered by embedding-based matching but excluded from the stricter sequence-level window, and embedding-confirmed hinges that lack the exact P-X-X-G pattern. Despite these methodological differences, both estimates place *S. tonkinensis* substantially above all other eudicots examined (56-73%), independently supporting a family-wide fixation of the N-terminal hinge. The median match position in *S. tonkinensis* (42) fell within the range observed in other angiosperms (40-42), indicating that the elevated fixation reflects a higher prevalence of the hinge across the family rather than a positional shift.

#### 3.4.6 The PxxG hinge delimits a natural boundary for N-terminal analysis

The soft-correspondence data place the PxxG constrained island at median sequence positions 44-51, substantially downstream of its reference coordinates (positions 20-27). The ∼24-residue offset reflects cumulative N-terminal indels: the reference sequence StonChr6AG199490.1 has a relatively compact N-terminus, whereas most query sequences carry insertions upstream of the hinge. This places the island at or immediately beyond position 50 in more than half of *S. tonkinensis* P450s.

A practical consequence follows: the region upstream of the PxxG hinge— approximately the first 40-50 amino acids of the native sequence—is devoid of any detectable sequence-level constraint. PEACE dispersion in this region is uniformly high (mean 0.622), no secondary constrained islands were detected, and directionality scores are negligible. The PxxG hinge itself, while near-absolutely conserved at Pro^21^ and Gly^24^, derives its constraint from backbone conformational requirements rather than from amino acid coding capacity (see Section 4.3).

This constraint topography provides an empirically grounded rationale for the analytical window adopted in the analyses that follow. The 50-amino-acid cutoff— which might otherwise appear arbitrary—corresponds approximately to the constraint-free zone upstream of the PxxG hinge in the majority of sequences. Positions beyond ∼50 increasingly overlap with the hinge itself and the catalytic domain, where structural constraints would dominate and obscure the subtler informational signals that are the focus of the next three sections. In what follows, we decompose this 50-amino-acid constraint-free zone into two separable information channels—one topological (DIVA, Section 3.6) and one semantic (PIVOT, Section 3.7-3.8)—and show that the absence of sequence-level constraint is not a void, but a substrate.

#### 3.4.7 Directionality and phylogenetic structure analyses yield no N-terminal signal

As an orthogonal layer of constraint interpretation, PEACE quantifies two additional properties at each position:

- **Directionality score**: whether the high-dimensional embedding variation partitions into discrete clusters (Gaussian Mixture Models, with the number of clusters selected by silhouette score). A high directionality score (>0.6) with a low effective rank indicates that variation at that position is *structured*—for example, into residue classes that correspond to CYP family membership—rather than random drift. Low directionality indicates that variation, however extensive, is unstructured with respect to any categorical partition.
- Phylo-structure z-score: whether the GMM cluster assignments at a position are non-randomly distributed across the phylogenetic tree. A high phylo-z (>2) would indicate that clusters map to specific clades, consistent with lineage-specific (episodic) selection; low phylo-z indicates that cluster membership is phylogenetically scattered, consistent with family-wide pervasive variation.

In both REFERENCE and DTW modes, the top ten directional positions mapped exclusively to the catalytic core (REFERENCE: positions 347-489, approximate positions 346-488; DTW: positions 344-442, approximate positions 304-455). None fell within the N-terminal 50 amino acids (both reports: “Of top 10 directional, 0/10 in N-term region”). This indicates that while the N-terminus is highly variable (high dispersion), its variation is *unstructured*—the embedding signatures at N-terminal positions do not partition into discrete residue clusters. This is consistent with neutral relaxation rather than diversifying selection driving N-terminal divergence (Figure 10 & Figure 11).

**Figure 10.**
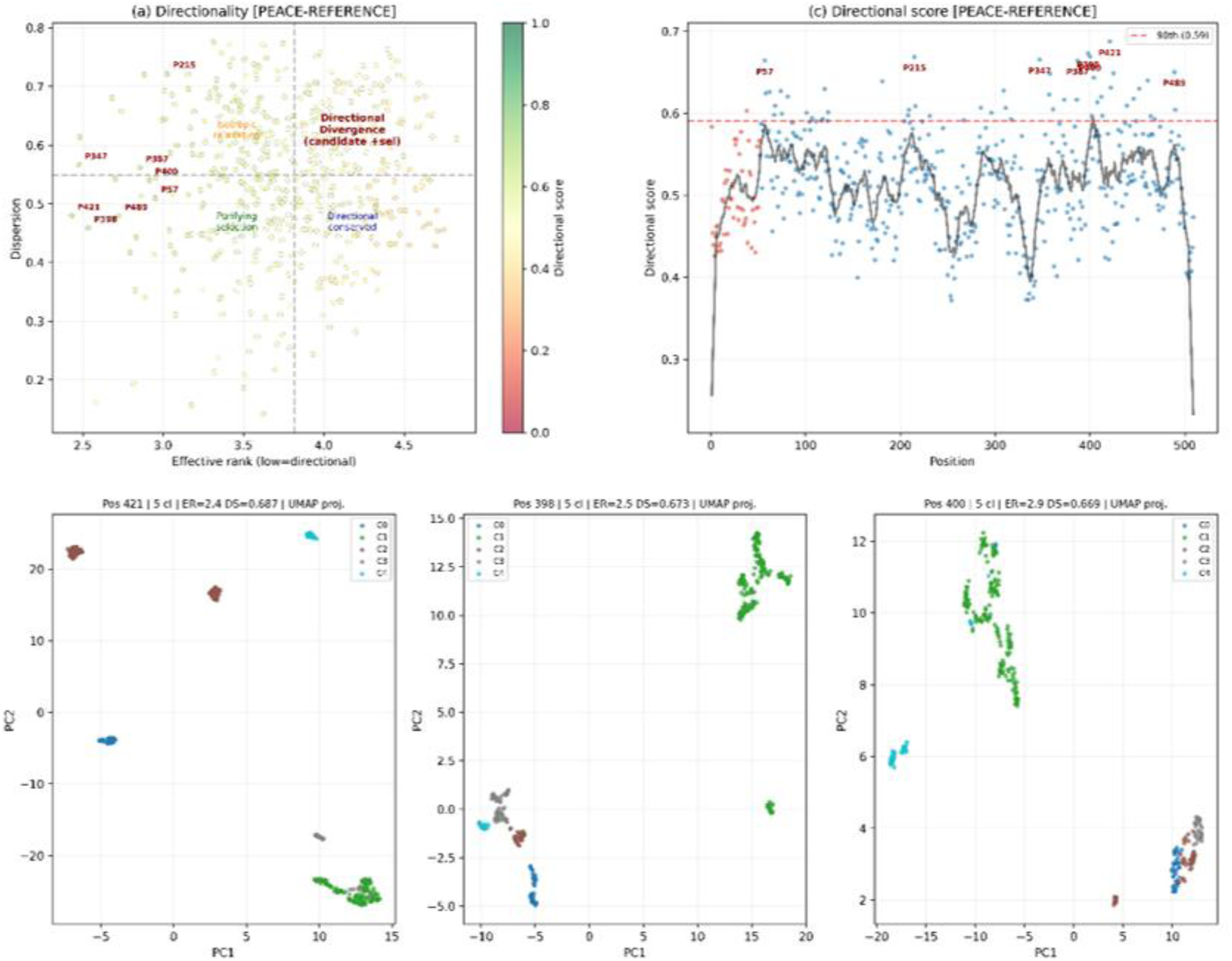
(reference mode) Directionality analysis of the PEACE - REFERENCE mode. (a) Scatter plot of effective rank versus dispersion, with individual points colored by the Directional score (ranging from low in red to high in green). Low effective rank indicates a highly directional embedding distribution, while high dispersion indicates greater embedding divergence. The dashed lines divide the plot into four quadrants representing distinct evolutionary regimes: Directional Divergence (candidate +sel), Isotropic relaxation, Purifying selection, and Directional conserved. Highly directional N-terminal and core positions (e.g., P215, P421, P490) are highlighted in red. (c) Distribution of directional scores across the protein sequence. Blue dots represent raw directional scores, and the black line indicates the smoothed trend. The red dashed line marks the 90th percentile threshold (0.59), with positions exceeding this threshold highlighted as red dots (candidates for episodic divergence or positive selection). Bottom panels show the UMAP (Uniform Manifold Approximation and Projection) embeddings of the three highest-scoring representative positions (Pos 421, 398, and 400). These positions exhibit strong directional scores (DS ≈ 0.67-0.69), 4-5 Gaussian Mixture Model (GMM) clusters (5 cl), and low effective ranks (ER ≈ 2.4—2.9), indicating distinct and structured directional divergence within the sequence family.

**Figure 11.**
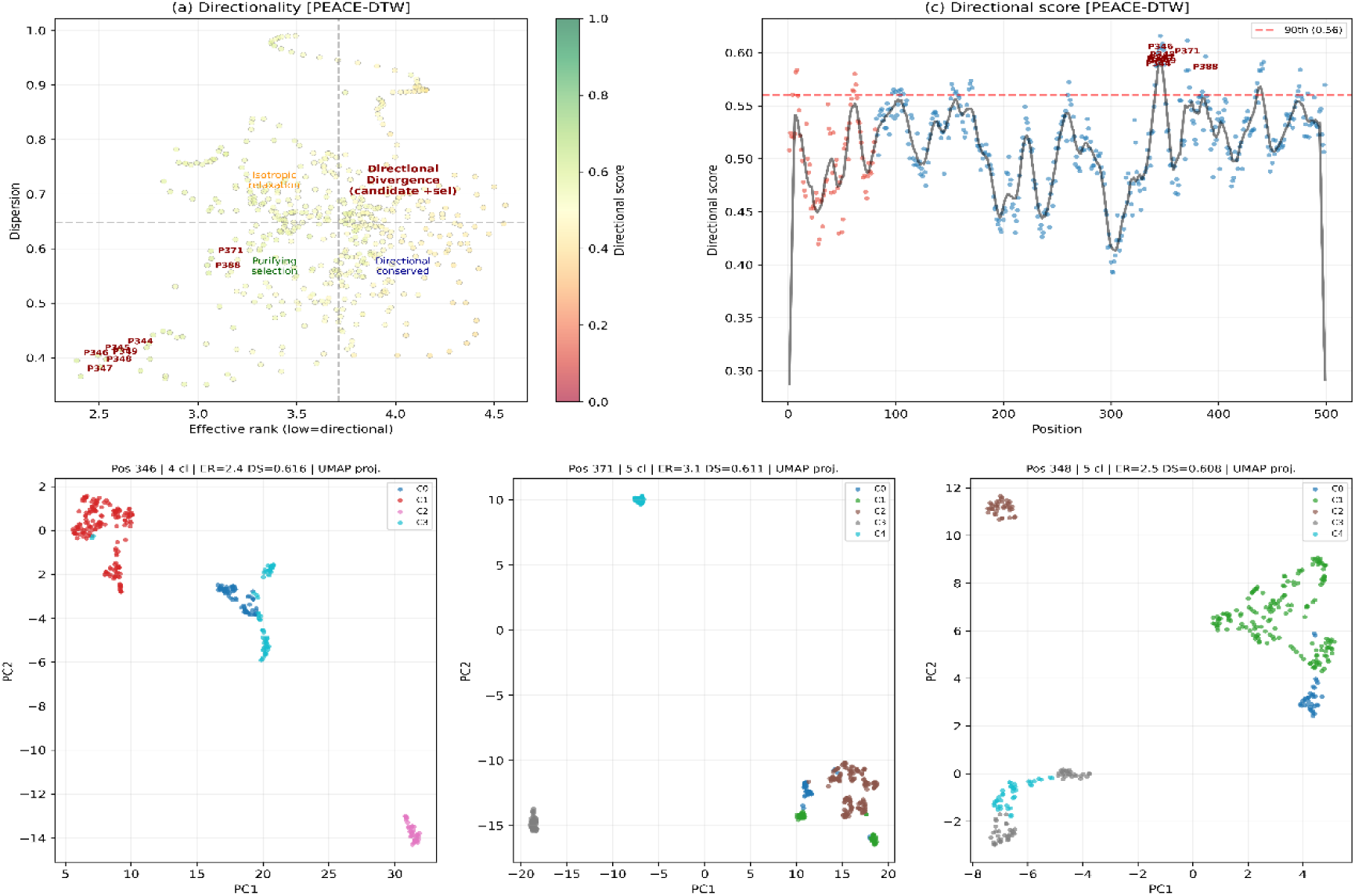
Directionality analysis of the PEACE-DTW mode. (a) Scatter plot of effective rank versus dispersion for the DTW consensus sequence. Points are colored by their Directional score to assess residue-level structural constraint and sequence divergence. The DTW mode generally captures higher dispersion values compared to the reference mode. The four quadrants define distinct evolutionary features: Directional Divergence (candidate +sel), Isotropic relaxation, Purifying selection, and Directional conserved. Highly scored positions (e.g., P371, P388) are labeled in red. (c) Directional score profile along the sequence position. The black smooth line shows the overall trend, and the red dashed line marks the 90th percentile threshold (0.56). Highly directional peaks are predominantly located in the latter half of the sequence (positions 300-500). Bottom panels display the UMAP projections of the three highest-scoring representative positions (Pos 346, 371, and 348). These positions reveal highly structured GMM clusters (4-5 clusters). Despite having moderate effective ranks (ER = 2.4-3.1), their high directional scores (DS ≈ 0.61) indicate strong directional divergence signals captured within the windowed DTW consensus.

Phylogenetic structure analysis revealed no significant signal anywhere in the protein. In REFERENCE mode, the top ten phylo-structure positions were all located in the catalytic core (positions 375-480), but none survived FDR correction (all PhyloQ ≥ 0.628). In DTW mode, two of the top ten phylo-structure positions mapped to the N-terminal region (positions 63-64, approximate position 34), alongside eight core positions; again, none approached FDR significance (all PhyloQ ≥ 0.586). The highest phylo-z in either mode was 1.81 (Reference, position 480), well below the conventional threshold of 2 for candidate episodic selection (Figure 12).

**Figure 12.**
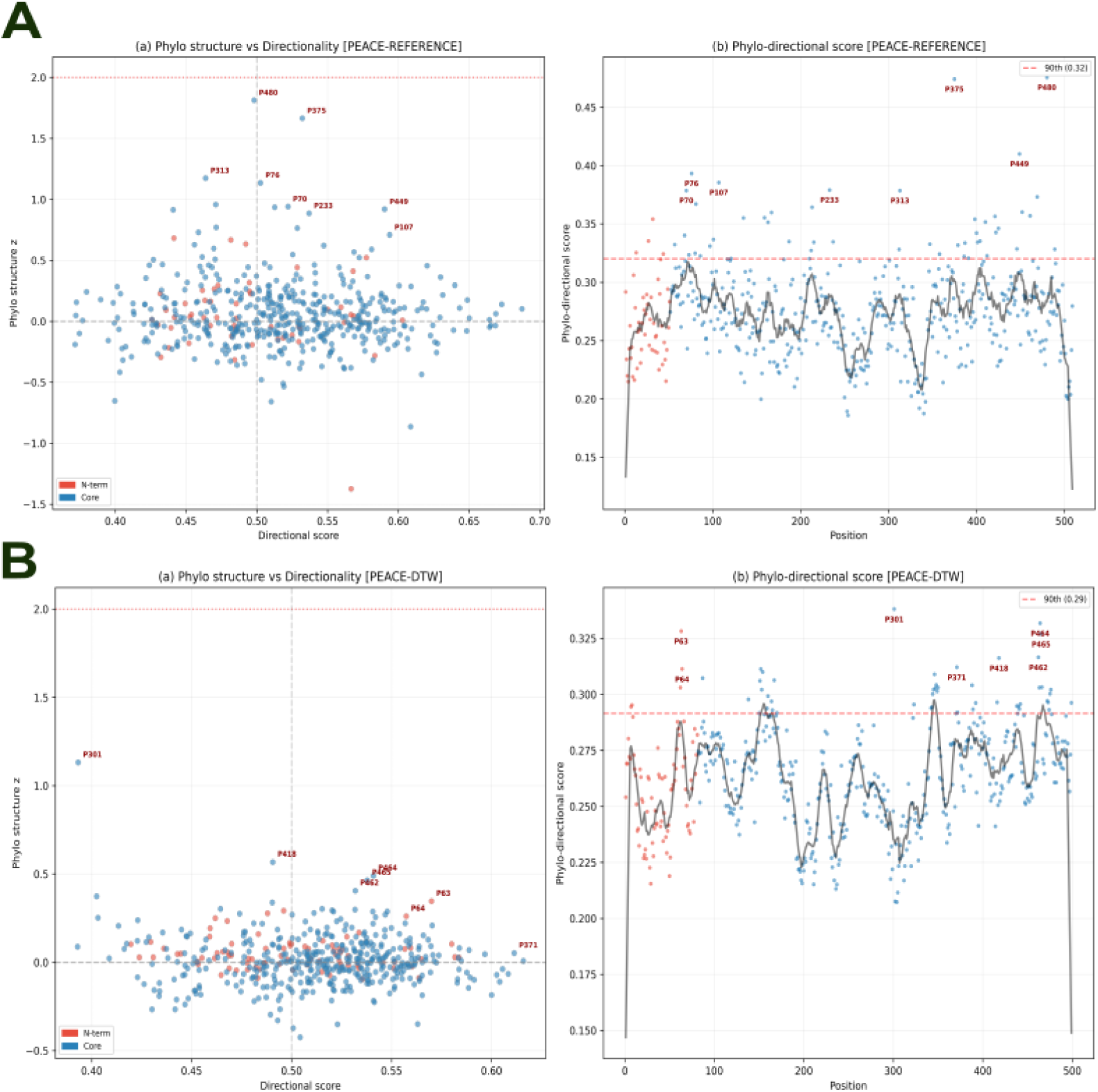
Integration of phylogenetic structure and directional divergence signal. (A) Phylo-structure and directional divergence analysis under the PEACE-REFERENCE mode. (a) Scatter plot of Phylo structure z-score versus Directional score. The z-score quantifies the concordance between GMM-derived directional clusters and the provided phylogenetic tree topology (z > 2, marked by the red dotted line, indicates strong clade-specific clustering). Points are color-coded by region (N-term in red, Core in blue), and the dashed gray line indicates the median Directional score threshold (0.5). The most prominent positions are annotated in red. (b) Phylo-directional score profile along the reference sequence. The black solid line represents the smoothed trend, while the red dashed line indicates the 90th percentile threshold (0.32). Positions exceeding this threshold are considered candidates for lineage-specific adaptation and are highlighted with sequence labels. (B) Phylo-structure and directional divergence analysis under the PEACE-DTW mode. (a) Scatter plot of Phylo structure z-score versus Directional score for the DTW consensus sequence. The z = 2 significance line is marked by a red dotted line. (b) Phylo-directional score profile across the window-aligned sequence positions. The 90th percentile threshold is shown as a red dashed line (0.29). Peaks that exceed the threshold (e.g., P301, P418, P371, P464) are highlighted in red, revealing strong combined signals of local embedding directionality and phylogenetic topology within the DTW framework

The absence of significant phylo-structure in the N-terminus—combined with the complete absence of directional signal—is consistent with the neutral-relaxation model. The N-terminal 50 amino acids are not merely variable; their variation lacks both categorical structure (directionality ≈ 0) and phylogenetic coherence (phylo-z ≈ 0). In contrast, variation in the catalytic core, while constrained in magnitude (low dispersion), shows structured directionality that is phylogenetically pervasive rather than lineage-specific—a pattern consistent with long-term functional constraint punctuated by family-level divergence, not episodic positive selection (Figure 12).

### 3.5 Population-Level Importance Landscapes Converge on the N-Terminus

To identify which regions of the P450 protein sequence carry functional information at the population level, per-residue importance profiles from DIVA (unsupervised) and PIVOT (supervised) were computed across the full protein length for all 345 sequences, and aggregated into normalized population-level density profiles (n = 200 bins spanning the normalized sequence position from 0 [N-terminus] to 1 [C-terminus]; threshold = 80th percentile of non-empty bin means).

Despite their fundamentally different designs—PIVOT identifies residues important for SVM discrimination of predefined functional clusters, while DIVA identifies residues whose embeddings align with each protein’s intrinsic functional specificity vector—both algorithms converged on the same dominant signal: the N-terminal region (Figure 13).

**Figure 13.**
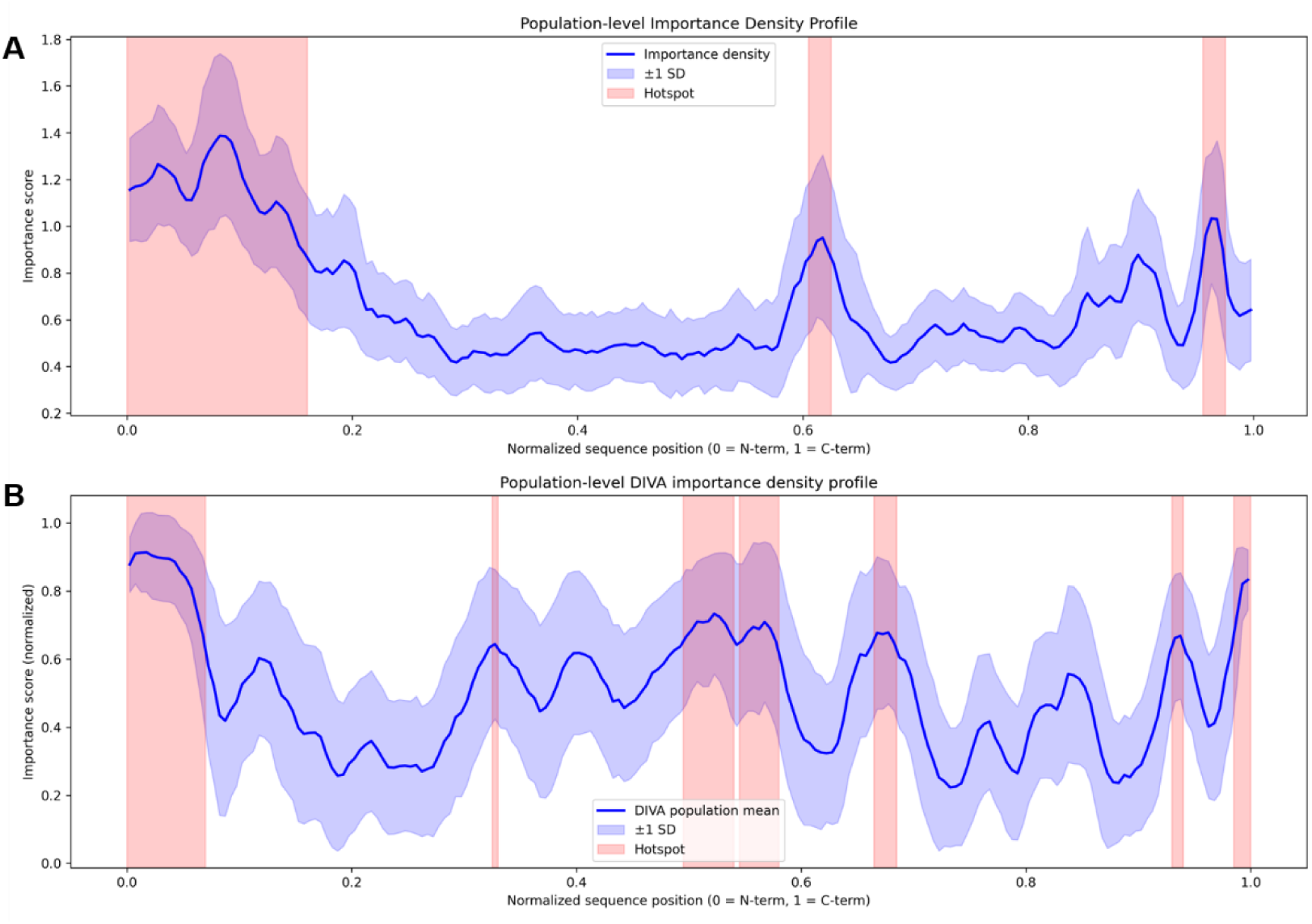
Population-level importance density profiles generated by two complementary residue-level attribution methods (PIVOT and DIVA) **A.** Population importance density profile based on the PIVOT method (AA_Position_Importance_V3(PIVOT).py). PIVOT is a supervised feature attribution algorithm. The method first establishes a multi-class classifier via ESM2 embeddings, PCA dimensionality reduction, and a linear SVM. It then employs a sliding window for local perturbations across full-length sequences, assessing the importance of the residues within each window by calculating the deviation in the SVM decision function before and after perturbation (Δ decision). The blue solid line represents the importance density, the light blue shaded area indicates ±1 standard deviation (SD), and the pink shaded regions denote automatically identified functional hotspots determined by the percentile threshold and coefficient of variation (CV) threshold. **B.** Population importance density profile based on the DIVA method. DIVA is an unsupervised attribution method. The algorithm constructs a directional difference vector using the difference between the ESM2 CLS token embedding and the average residue embedding of the sequence. The importance score for each individual residue is derived from the absolute cosine projection of its embedding onto this directional vector, followed by normalization. The blue solid line represents the DIVA population mean, the light blue shaded area indicates ±1 standard deviation (SD), and the pink shaded regions represent hotspots identified based on a percentile threshold.

Note: Both panels normalize original sequence positions to the [0, 1] interval (0 = N-terminus, 1 = C-terminus) and employ a unified importance score scale. This enables a direct comparison of key functional regions uncovered by fundamentally different strategies (supervised classifier-driven perturbation vs. unsupervised embedding-space directional projection). PIVOT detected three population-level hotspots (Figure 13A). The N-terminal hotspot (bins 0-31, relative position 0.0-0.16, mean importance = 1.171) carried the highest absolute importance, exceeding the C-terminal hotspot (bins 191-194, relative position 0.955-0.975, mean = 0.983) by 19% and the mid-sequence hotspot (bins 121-124, relative position 0.605-0.625, mean = 0.915) by 28%. The prominence of the N-terminal signal is notable: a 32-bin contiguous block at the extreme N-terminus exceeded in mean importance all other detected hotspots, including the heme-binding region and the C-terminal catalytic residues that define the P450 fold.

DIVA independently detected seven hotspots (Figure 13B). The N-terminal region (bins 0-13, relative position 0.0-0.07, mean importance = 0.858) exhibited the highest hotspot-level mean across the entire profile, exceeding the global threshold (0.642). Additional DIVA hotspots were distributed across the mid-sequence (0.325-0.33, mean = 0.644; 0.49 5-0.54, mean = 0.701; 0.545-0.58, mean =0.681) and C-terminal regions (0.665-0.685, mean = 0.669; 0.93-0.94, mean = 0.665; 0.985-1.0, mean = 0.792). However, no region outside the N-terminus matched its mean importance—the strongest non-N-terminal DIVA hotspot (0.495-0.54, mean = 0.701) fell 18% below the N-terminal value, and was only marginally above the global threshold.

The convergence of an unsupervised metric (DIVA) and a supervised metric (PIVOT) on the same N-terminal region—computed from entirely different mathematical principles and without shared parameters—constitutes strong evidence that this region carries functionally significant information. That both algorithms place the N-terminus as the highest-intensity region in their respective profiles confirms that the N-terminal signal is not an artifact of classifier training or embedding normalization, but a genuine feature of the P450 sequence-function landscape. Per-protein importance profiles for all 345 sequences are provided in Supplementary Materials (Dataset S1).

### 3.6 Physicochemical Validation of DIVA Reveals a Topological Template

To determine what biological signal DIVA scores capture—beyond their computational definition—per-residue DIVA scores across all 17,250 residue-position observations (345 proteins × 50 N-terminal positions) were correlated with four classical amino acid properties (hydropathy, helical propensity, side-chain volume, aromaticity) and tested for charge class effects, partitioned into three segments matching the ER signal-anchor topology phases: 1-8 (pre-insertion), 9--40 (transmembrane helix core), and 41-50 (cytosolic exit bend).

Two findings establish that DIVA captures a distinct biological signal rather than a simple correlate of known sequence properties. First, in the 1-8 aa segment—where DIVA scores are highest (Section 3.5; see also Section 3.7)—the correlation between DIVA and position-level Shannon entropy was negligible (p = +0.046), ruling out the interpretation that DIVA merely measures amino acid conservation. Second, the physicochemical correlates of DIVA differed fundamentally across the three segments, revealing a tripartite pattern that precisely matches the biophysical steps of ER signal-anchor topology determination (Fig. 14): **Positions 1-8 (pre-insertion)**: weak positive correlation with hydropathy (p = +0.086, p < 0.001) but negative correlation with helical propensity (p = -0.045, p = 0.019)—a preference for moderately hydrophobic but helix-avoiding residues. Charged residues showed significantly lower DIVA than uncharged residues (mean: 0.868 vs. 0.903, Mann-Whitney U p = 7.3 × 10^-18^). **Positions 9-40 (transmembrane helix core)**: classical transmembrane helix signature—strongest hydrophobicity correlation across all segments (ρ = +0.261, p < 0.001), strong helical propensity (ρ = +0.191, p < 0.001), and significant contributions from side-chain volume (ρ = +0.101, p < 0.001) and aromaticity (ρ = +0.125, p < 0.001). **Positions 41-50 (cytosolic exit bend)**: complete reversal—hydrophobicity correlation inverted to strongly negative (ρ = -0.240, p < 0.001), helical propensity and other properties became non-significant, and charged residues carried substantially higher DIVA than uncharged residues (mean: 0.639 vs. 0.425), with positively charged residues (R, K) showing the highest scores (mean = 0.670)—consistent with the “positive-inside rule” that governs membrane protein orientation (Figure 14).

**Figure 14.**
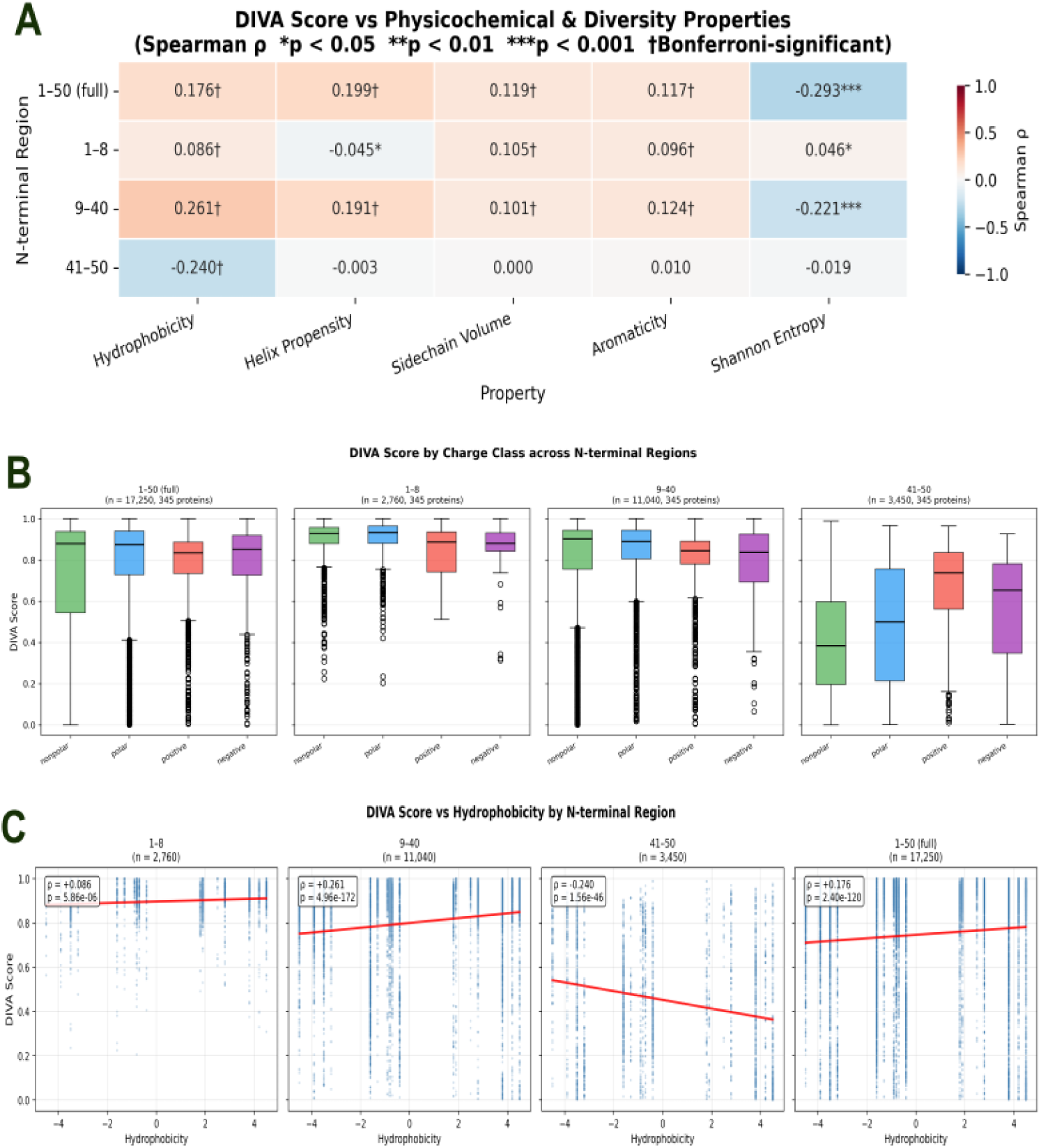
Physicochemical validation of DIVA scores across N-terminal regions reveals a tripartite topological template. **A.** Heatmap of Spearman rank correlations between DIVA scores and five residue-level properties (hydropathy, helix propensity, side-chain volume, aromaticity, and Shannon entropy) across distinct N-terminal segments (1-8, 9-40, 41-50, and 1-50 full). The color gradient represents the Spearman ρ (red for positive, blue for negative). Asterisks and daggers denote statistical significance (*p < 0.05, **p < 0.01, ***p < 0.001, ^†^Bonferroni-adjusted significant). **B.** Distribution of DIVA scores across four charge classes (nonpolar, polar, positive, negative) in the defined N-terminal regions. Boxplots show medians and interquartile ranges. Charged residues are significantly disfavored in the 1-8 segment but strongly favored in the 41-50 segment, consistent with the “positive-inside rule” governing ER membrane protein topology. **C.** Scatter plots and linear regression trends (red lines) of DIVA score versus hydrophobicity for each N-terminal region. Spearman correlation coefficients (ρ) and p-values are annotated in the inset boxes. The correlations exhibit a sharp inversion from a strongly positive slope in the transmembrane helix core (9-40, ρ = +0.261) to a strongly negative slope in the cytosolic exit bend (41-50, ρ = -0.240), validating the dynamic topological shift

This tripartite pattern—weakly hydrophobic pre-insertion → strongly hydrophobic transmembrane helix → hydrophilic, charge-enriched cytosolic exit—recapitulates the three-step biophysical program of ER signal-anchor topology determination (von Heijne, 1989). We therefore conclude that DIVA encodes a “topological template”: a position-dependent physicochemical profile specifying membrane-anchoring architecture, operating as a directional constraint on residue properties rather than a site-specific requirement for residue identity. The complete statistical results are provided in Table T6.

### 3.7 N-Terminal Importance Profiles Reveal Separable Encoding Architectures

To characterize how individual proteins differ in their N-terminal importance architecture, DIVA and PIVOT profiles were independently subjected to hierarchical clustering based on profile shape and peak position (NIPC pipeline; Methods 2.8.4). Proteins were first classified as “N-terminal-driven” (meeting both concentration and peak criteria within the 1-50 aa window) or “Weak-N” (failing either criterion); the N-terminal-driven subset was then clustered and labeled Early-N, Mid-N, or Late-N according to mean peak position.

#### DIVA clustering

identified four profile types from 337 N-terminal-driven proteins, with 8 additional Weak-N genes (Supplementary Table T7). A single cluster dominated: **Early-N** (n = 324, 96.1% of N-terminal-driven), exhibiting a sharp importance peak at positions 1-13 (mean importance > 0.90 at positions 3-13) that decayed monotonically C-terminally to ∼0.24 at position 50. This stereotyped profile is consistent with the topological template identified in Section 3.6: the vast majority of P450s concentrate their DIVA signal in the membrane pre-insertion and early transmembrane segments. Minor clusters included Late-N (n = 9), Mid-N_1 (n = 3), and Mid-N_2 (n = 1), together accounting for only 3.9% of N-terminal- driven proteins (Figure 15 A C).

**Figure 15.**
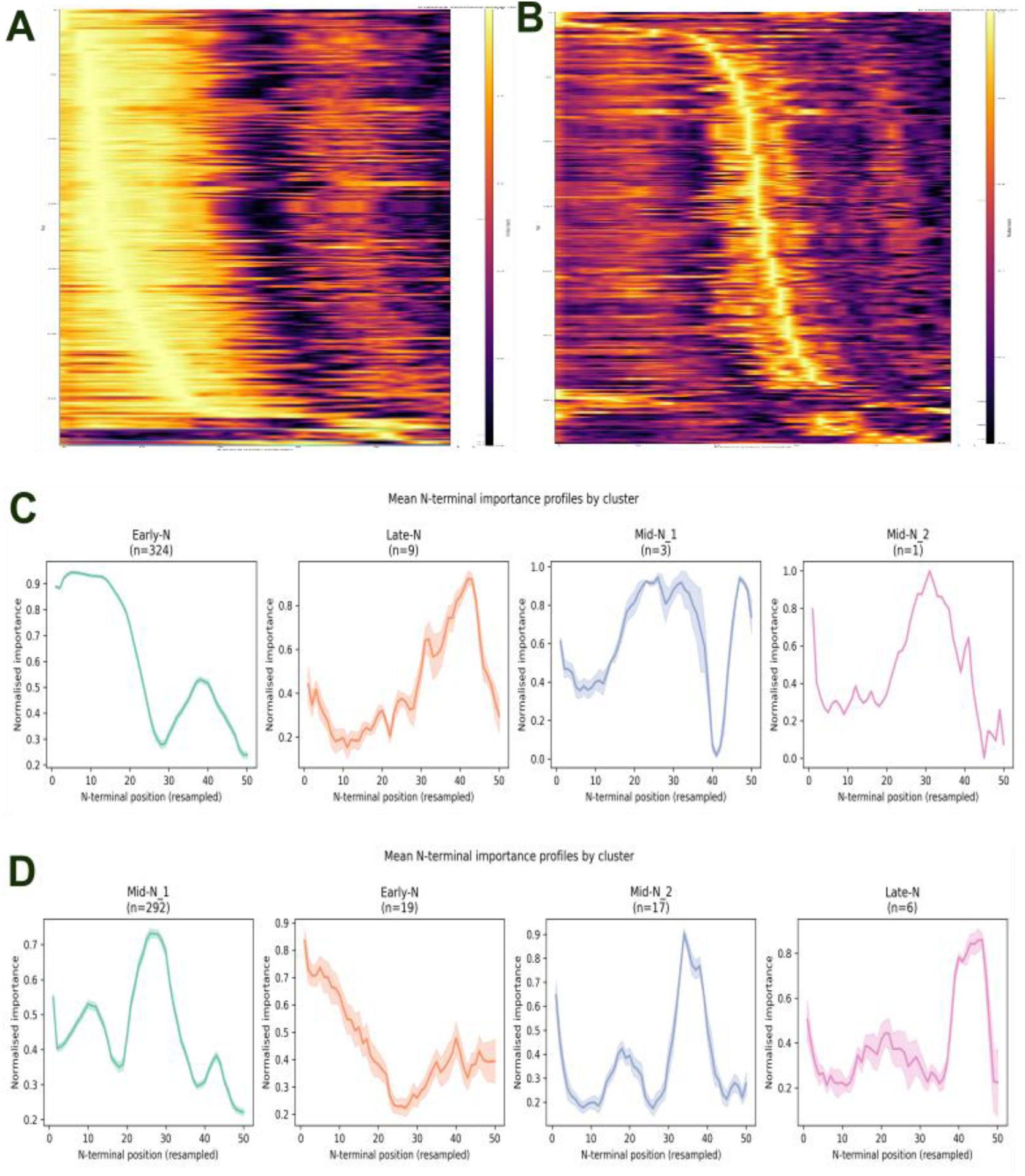
Distinct visual architectures and clustering patterns of DIVA and PIVOT N-terminal importance profiles. (A, B) Per-protein heatmaps of normalized importance across the first 50 N-terminal positions for (A) DIVA and (B) PIVOT. Each row represents an individual sequence, with color intensity from dark purple (low) to bright yellow/white (high). The DIVA heatmap shows a concentrated, stereotypic “Early-N” bright spot that remains tightly anchored within the first 10-15 residues across the majority of proteins. In sharp contrast, the PIVOT heatmap exhibits a progressively migrating diagonal bright band, indicating that high importance regions dynamically shift from the extreme N-terminus toward more C-terminal positions (residues ∼25-30) in a protein-dependent manner. (C, D) Mean N-terminal importance profiles of the four clusters defined by (C) DIVA and (D) PIVOT. Shaded regions denote ±1 standard deviation. The DIVA dominant cluster, Early-N (n=324, 96.1%), features a sharp peak at positions 1-13 that decays monotonically, fully consistent with the visual concentration in panel A. The PIVOT dominant cluster, Mid-N_1 (n=292, 87.4%), features a broad, plateau-shaped profile peaking at positions 25-28, visually corroborating the diagonal migration observed in panel B. These separable architectures indicate that DIVA captures a stereotypic topological template while PIVOT identifies more protein-specific, dynamically shifting functional determinants.

#### PIVOT clustering

produced a fundamentally different distribution from 334 N-terminal-driven proteins (11 Weak-N, Supplementary Table T8). One cluster dominated, but less overwhelmingly than DIVA: **Mid-N_1** (n = 292, 87.4%), showing a broad importance profile that rose gradually from ∼0.55 at position 1 to a plateau at positions 25-28 (mean importance ∼0.71-0.73) and decayed gently toward both termini. Three smaller clusters were identified: Mid-N_2 (n = 17, 5.1%), Early-N (n = 19, 5.7%), and Late-N (n = 6, 1.8%). Early-N proteins exhibited elevated importance in the first ∼10 positions but with substantially lower magnitude than DIVA Early-N (mean at position 1: 0.83 vs. 0.89 for DIVA). Late-N proteins, though few (n = 6), displayed a distinct sharp peak at positions 42-46 (mean importance ∼0.84-0.86). This heterogeneous distribution—with the dominant cluster capturing only 87.4% of proteins compared to 96.1% for DIVA— indicates that residues critical for SVM cluster discrimination (PIVOT) are distributed across the entire N-terminal segment in a more protein-specific manner than the stereotypic DIVA membrane topology template (Figure 15 B D).

#### Cross-metric comparison

revealed three lines of evidence for separable encoding. First, the Weak-N gene sets were completely mutually exclusive: zero overlap between the 8 DIVA Weak-N genes and 11 PIVOT Weak-N genes. Genes lacking N-terminal importance in one metric invariably retain it in the other—DIVA and PIVOT identify fundamentally different subsets of proteins as N-terminally “important.”

Second, cluster assignments showed minimal correspondence. DIVA placed 96.1% of N-terminal-driven proteins in a single Early-N cluster, whereas PIVOT distributed proteins across four clusters with its dominant cluster (Mid-N_1) capturing 87.4%. A gene’s DIVA profile type did not predict its PIVOT profile type.

Third, per-protein peak positions within the 1-50 aa window were compared across the 326 shared genes. DIVA peaks were overwhelmingly concentrated at the extreme N-terminus (mean relative position = 0.08 ± 0.10, median = 0.08), while PIVOT peaks spanned the entire window with a clear downstream shift (mean = 0.35 ± 0.26). The Spearman correlation between DIVA and PIVOT peak positions was weak but statistically significant (ρ = 0.38, p = 4.4 × io^-13^), with only 6% of shared variance (r^2^≈ o.o6)—consistent with the minimal coupling expected from their shared 50-residue backbone. In 84.4% of cases, the PIVOT peak fell downstream of the DIVA peak (mean Δ = +0.27, corresponding to ∼13.5 residues), with only 10.2% of genes showing close alignment (|Δ| ≤ 0.15)(Figure 16 A).

**Figure 16.**
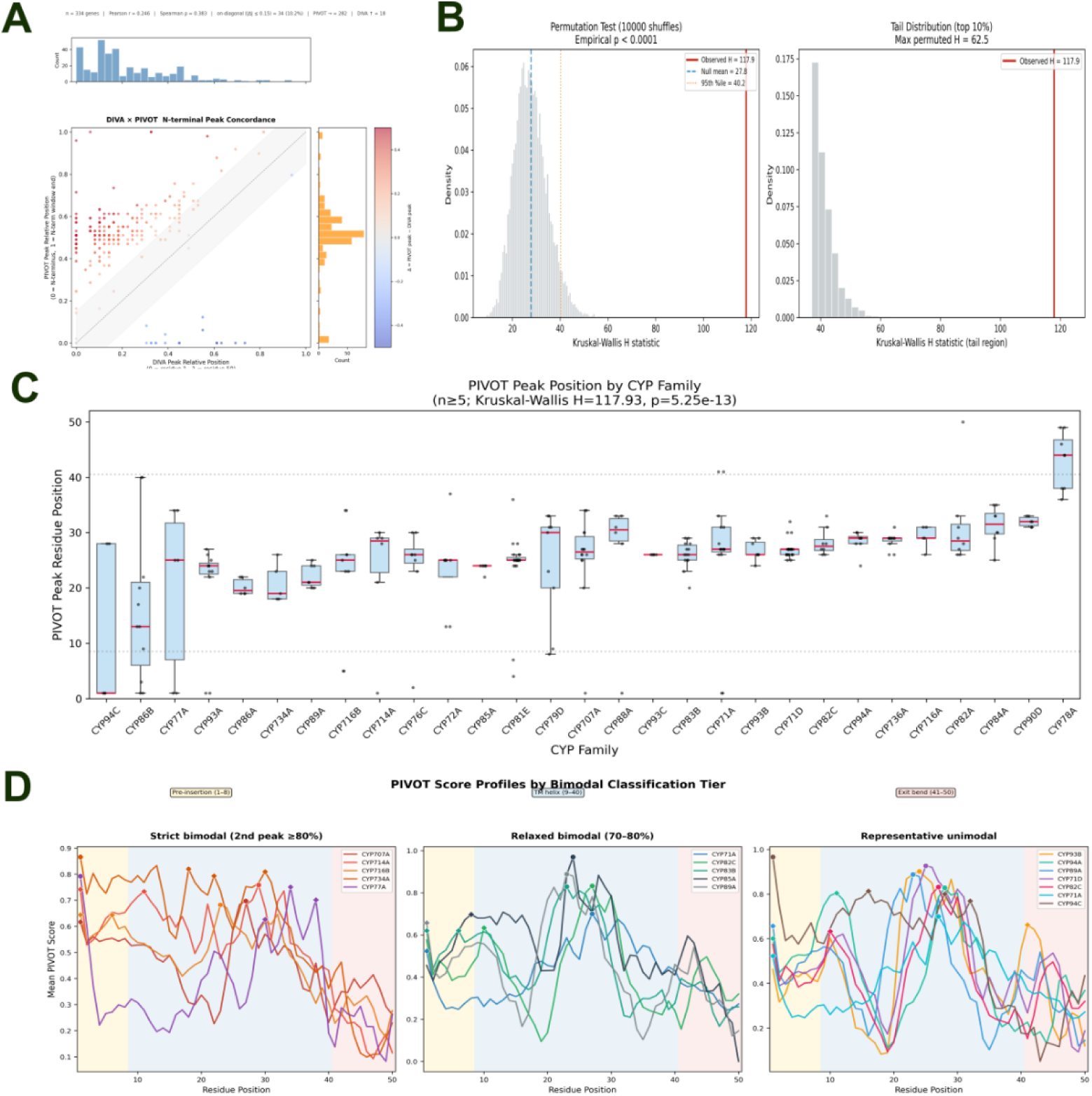
Cross-metric comparison and statistical validation of PIVOT peak positions as a family-discriminatory signal. **A.** Scatter plot of normalized PIVOT peak positions versus DIVA peak positions within the 1-50 N-terminal window across 334 shared proteins (n=334 genes). The diagonal dashed line indicates perfect agreement; the gray shaded band marks close alignment (| Δ |≤ 0.15). Points are color-coded by the shift distance Δ (PIVOT peak - DIVA peak). Only 10.2% of genes show close alignment, with PIVOT peaks falling downstream of DIVA peaks in 84.4% of cases, indicating weak coupling (Spearman *ρ* = 0.383, *r***_2_** ≈ 0.06). **B.** Permutation test demonstrating that this association is not an artifact of family size imbalance. The observed Kruskal-Wallis H statistic (117.93, red solid line) drastically exceeds the null distribution generated from 10,000 random shuffles of family labels (maximum permuted H = 62.5). The resulting empirical p-value is p< 0.0001. **C.** Boxplot showing the distribution of PIVOT peak residue positions across CYP families with at least five members (η≥5). A Kruskal-Wallis test reveals a highly significant association between PIVOT peak position and family identity (*H* = 117.93,*p* = 5.25 × 10**^-13^**,*̅***^2^** = 0.44), indicating a large effect size where 44% of the variance is explained by family membership. **D.** Mean PIVOT score profiles categorized by bimodal architecture tiers. Three representative profile types are shown: **Strict bimodal** (secondary peak ≥80% of the global peak, e.g., CYP707A, CYP734A), **Relaxed bimodal** (secondary peak 70-80%), and **Representative unimodal** (single sharp peak, e.g., CYP93B, CYP71D). Families with strict bimodal profiles possess two distinct semantic identity hotspots, reflecting complex evolutionary constraints. The background colors denote the three functional zones defined by DIVA: pre-insertion (yellow), TM helix (blue), and exit bend (pink).

Taken together, these results establish that DIVA and PIVOT measure two largely separable importance architectures on the same N-terminal segment. DIVA importance is overwhelmingly concentrated at the extreme N-terminus (positions 1-13), with a stereotypic Early-N profile shared by >96% of proteins—consistent with a conserved topological template (Section 3.6). PIVOT importance is distributed more broadly across the 50-amino-acid segment, with protein-specific peak positions that show weak but non-zero coupling with DIVA peaks (ρ = 0.38). The weak residual correlation is expected: both signals operate on the same linear peptide and are thus subject to shared physical constraints. The mutual exclusivity of Weak-N gene sets, the divergence in cluster distributions, and the systematic downstream offset of PIVOT peaks collectively support the interpretation of separable information channels within a shared peptide scaffold, developed further in the Discussion (Section 4.2).

To determine whether the PIVOT signal carries family-discriminatory information, we tested for an association between PIVOT peak residue positions and CYP family identities. A Kruskal-Wallis test across 29 families containing at least five members revealed a highly significant association (*H* = 117.93,*p* = 5.25 × 10^-13^; **Figure 16C**). The effect size was large (*ε^2^* = 0.44), indicating that approximately 44% of the variance in peak position is explained by family membership. To confirm that this association was genuine rather than a consequence of imbalanced family sizes, we performed a permutation test. Among 10,000 random shuffles of family labels, no permutation exceeded the observed H statistic (maximum permuted *H* = 62.5), yielding an empirical p-value of *p* < 0.0001 (**Figure 16B**). The observed H statistic was 4.24-fold higher than the mean of the null distribution (27.8), confirming a robust, non-random association.

Despite this strong overall signal, pairwise differences between CYP families were remarkably sparse. Dunn’s post-hoc test with Holm correction identified only 2 out of 406 possible pairwise comparisons (0.5%) as statistically significant (adjusted *p* < 0.05). This pattern is the statistical hallmark of a continuous gradient rather than a categorical partition. Thus, PIVOT peak positions vary systematically across CYP families along a spectrum without discrete boundaries, consistent with functional diversification through gradual substrate specialization.

We further examined the architectural diversity of family-level PIVOT profiles using a local-maximum-based bimodality detection algorithm. Based on the ratio of secondary peaks to the global peak, families were classified into three tiers: *Strict bimodal* (secondary peak ≥ 80%; e.g., CYP707A, CYP77A, CYP78A), *Relaxed bimodal* (70-80%; e.g., CYP71A, CYP82C, CYP85A), and *Representative unimodal* (e.g., CYP71D, CYP72A, CYP93C) (**Figure 16 D**). Strict bimodal families displayed two distinct semantic hotspots—typically one at the extreme N-terminus and a second in the C-terminal portion of the N-terminal window (positions 25-40 or 41-50), often reflecting the dual demands of membrane insertion and catalytic geometry. In contrast, representative unimodal families exhibited a single sharp peak centered near position 25. The presence of clear bimodal architectures in 11 out of 29 families (38%) further indicates that PIVOT captures a complex, multi-faceted identity signal that is not reducible to a single positional determinant.

### 3.8 DeepLoc-2.1 subcellular localization predictions support the separable encoding model

As an independent test of whether the N-terminal sequence features defined by DIVA and PIVOT are recognized by modern localization-prediction algorithms, the full-length protein sequences of all 345 *S. tonkinensis* P450s were submitted to DeepLoc 2.1 (Nielsen, 2025). Of these, 328 (95.1%) were predicted to localize to the endoplasmic reticulum (ER), consistent with the expected membrane-anchored P450 localization (Werck-Reichhart & Feyereisen, 2000). The remaining 17 proteins (4.9%) received non-ER predictions distributed across five compartments: plastid (n = 5), cytoplasm (n = 5), nucleus (n = 3), mitochondrion (n = 2), and cell membrane (n = 2) (Figure 17).

**Figure 17.**
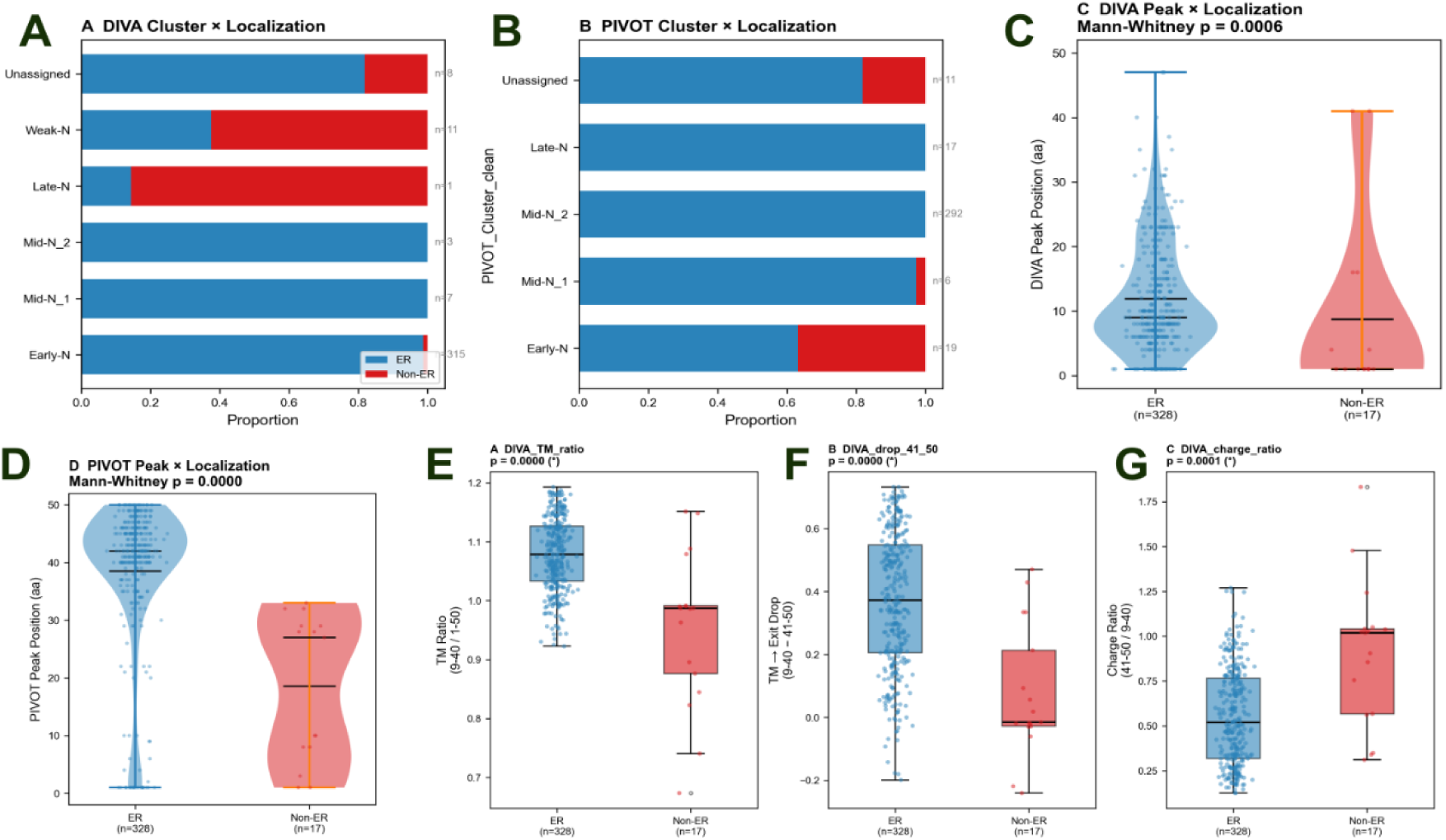
DeepLoc-2.1 validation of DIVA and PIVOT reveals a topological template and separable localization signals. **(A-B)** Stacked bar plots showing the proportion of ER-localized (blue) versus non-ER (red) proteins across **A.** DIVA clusters and **B.** PIVOT clusters. The DIVA Weak-N cluster is strongly enriched for non-ER predictions, whereas PIVOT Weak-N proteins are exclusively ER- localized. **(C-D)** Violin plots of **C.** DIVA peak positions and **D.** PIVOT peak positions for ER vs. non-ER proteins. DIVA peaks are significantly concentrated at the extreme N-terminus in ER proteins (Mann-Whitney p=0.0006), while PIVOT peaks show a complementary spatial shift. **(E-G)** Compliance with the DIVA topological template across ER and non-ER proteins. ER-predicted proteins display significantly higher TM-ratio (DIVA 9-40/1-50, p=0.0000), a pronounced TM→Exit drop (p=0.0000), and a reduced Charge ratio (41-50/9-40, p=0.0001), consistent with the canonical N-terminal ER-signal-anchor architecture.

#### DIVA detects topology-relevant features; PIVOT does not

To assess whether the topological template captured by DIVA is recognized by DeepLoc 2, we correlated DIVA and PIVOT segment scores with the predicted ER probability. Over the 9-40 window—which encompasses the hydrophobic transmembrane region in the topological template model—the mean DIVA score showed a significant positive correlation with ER probability (Spearman *ρ* = +0.272, p < 0.001; Figure 18A). By contrast, the mean PIVOT score over the same 9-40 interval showed no correlation with ER probability (ρ = -0.030, p = 0.573; Figure 17B). This asymmetry directly supports the separable encoding model: the topological template captured by DIVA constitutes a recognized input for subcellular localization prediction, whereas the identity-encoding features tracked by PIVOT in the transmembrane region are irrelevant to the ER targeting problem.

**Figure 18.**
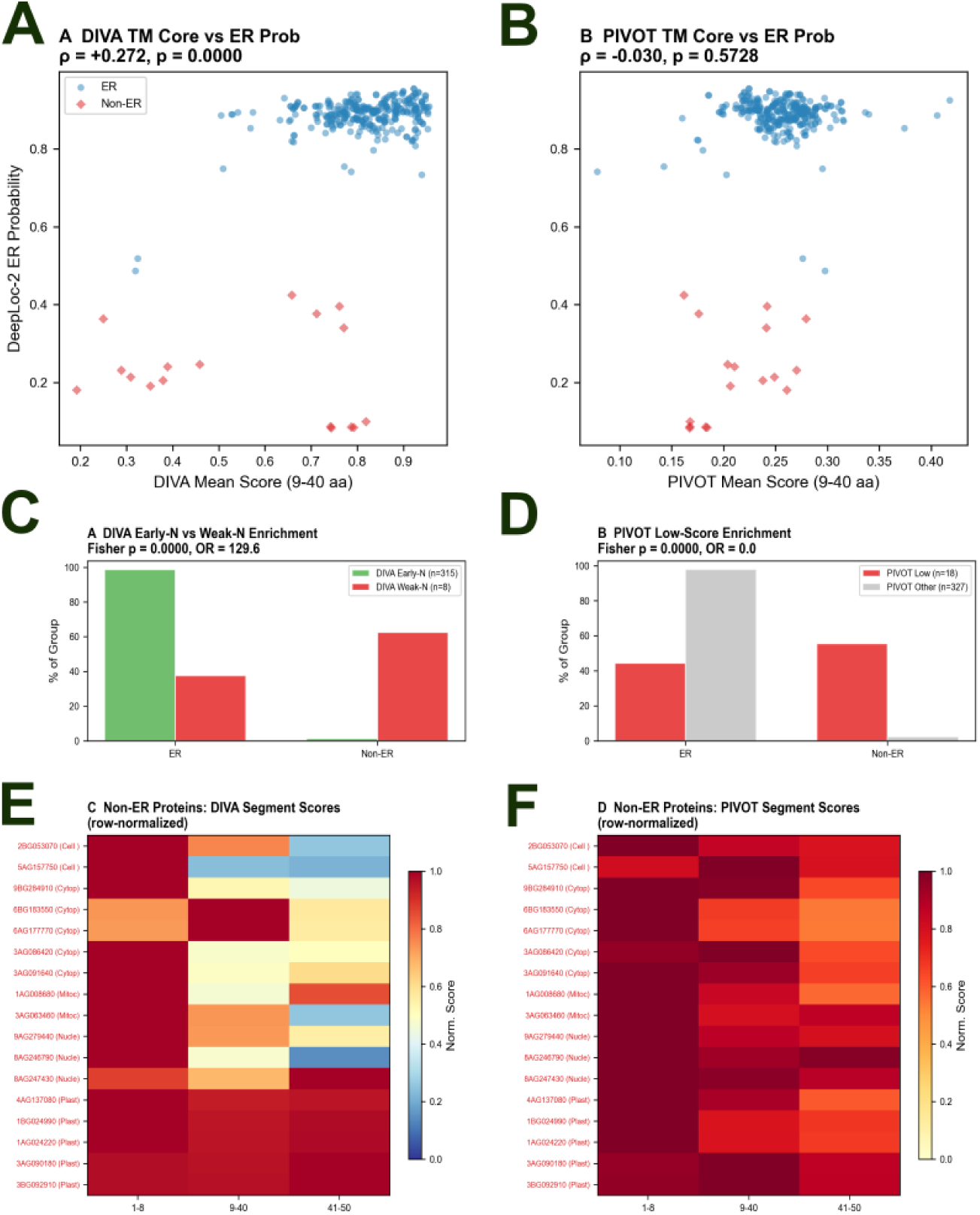
Continuous DeepLoc-2.1 probabilities and non-ER protein profiles confirm the independence of DIVA and PIVOT channels. **A.** Scatter plot of DIVA mean score in the transmembrane region (9-40) versus DeepLoc-2.1 ER probability, showing a significant positive correlation (*ρ* =+0.272,p = 0.0000). **B.** Scatter plot of PIVOT mean score in the same region (9-40) versus ER probability, showing no correlation (*ρ* = -0.030,p = 0.5728). **C.** Enrichment analysis of DIVA Early-N vs. DIVA Weak-N proteins. DIVA Weak-N proteins are highly enriched in the non-ER category (Fisher’s exact p=0.0000, OR=129.6). **D.** Enrichment analysis of PIVOT low-score vs. other PIVOT proteins. In sharp contrast, PIVOT low-score proteins are exclusively localized to the ER. **(E-F)** Row-normalized heatmaps showing DIVA and PIVOT segment scores (1-8, 9-40, 41-50) for individual non-ER proteins (n=17). The target subcellular compartments (Plastid, Mitochondrion, Cytoplasm, Nucleus, Cell membrane) are indicated. The divergence in segmental profiles between the two metrics underscores the separable, largely orthogonal nature of the topological and identity signals.

The dissociation extends beyond ER probability alone. DIVA mean 9-40 showed an even stronger positive correlation with predicted transmembrane probability (ρ = +0.562, p < 0.001), whereas PIVOT mean 9-40 showed a significant *negative* correlation with the same metric (ρ = -0.300, p < 0.001). That is, sequences conforming more strongly to the DIVA topological template are assigned higher transmembrane confidence by DeepLoc 2, while sequences with stronger PIVOT identity signatures in this region are assigned lower transmembrane confidence —consistent with the interpretation that PIVOT captures evolutionary features separable from—and in this region potentially antagonistic to—the structural demands of membrane anchorage.

#### DIVA and PIVOT both discriminate ER from non-ER, but through different regions

Although the PIVOT 9-40 segment is uncorrelated with ER probability, PIVOT is not globally uninformative for localization. Both DIVA and PIVOT scores differ significantly between ER- and non-ER-predicted proteins, but their discriminative signal is carried by distinct N-terminal sub-regions (Supplementary Table T9). DIVA mean scores over the 1-8 anchoring segment (0.908 vs. 0.714, Mann-Whitney p < 0.001) and the 9-40 transmembrane segment (0.819 vs. 0.553, p < 0.001) were both substantially lower in non-ER proteins, while the 41-50 juxtamembrane region showed no difference (p = 0.647). PIVOT showed the inverse regional pattern: no difference in the 1-8 segment (p = 0.180) but significant differences in the 9-40 (p < 0.001) and 41-50 (p < 0.001) windows, as well as in peak position (ER median = 42.0 vs. non-ER median = 27.0, p < 0.001). This complementary regional architecture—DIVA signals concentrated in the N-terminal anchoring and transmembrane regions, PIVOT signals concentrated in the C-terminal portion of the 50-aa window—further reinforces the separable-channel interpretation: the two information layers occupy different, though partially overlapping, spatial domains within the N-terminus.

#### Topological template compliance metrics distinguish ER from non-ER

Three derived metrics from the topological template model—the transmembrane ratio (DIVA 9-40 mean / DIVA 1-50 mean), the juxta membrane drop (DIVA 9-40 mean - DIVA 41-50 mean), and the charge ratio (DIVA 41-50 mean / DIVA 9-40 mean)—all differed significantly between ER- and non-ER-predicted proteins (all p < 0.001; Figure 17 E-G). ER-predicted proteins exhibited a high transmembrane ratio (1.079 vs. 0.954), a pronounced juxta membrane drop (0.369 vs. 0.076), and a low charge ratio (0.549 vs. 0.905), consistent with the canonical N-terminal topology of ER-anchored P450s: a hydrophobic transmembrane segment followed by a positively charged juxta membrane linker. Non-ER proteins deviate from this pattern on all three metrics, indicating that DeepLoc 2 is sensitive to the same physicochemical features encoded in the DIVA decomposition.

#### DIVA Weak-N proteins are strongly enriched among non-ER predictions

The most pronounced signal emerged from the set of DIVA Weak-N proteins (n = 8), defined by the absence of a discernible DIVA peak in the 1-8 membrane-proximal anchoring segment (see Section 3.7). These proteins were highly enriched among non-ER predictions: 5 of 8 (62.5%) DIVA Weak-N proteins fell outside the ER category, compared with only 4 of 324 (1.2%) among DIVA Early-N proteins possessing a strong 1-8 peak (Fisher’s exact test, p < 0.001; odds ratio = 129.6; Figure 18 C). Among the 5 non-ER DIVA Weak-N proteins, predicted localizations included mitochondrion (n = 1), cytoplasm (n = 1), nucleus (n = 2), and cell membrane (n = 1), suggesting a loss of ER-targeting fidelity when the membrane-proximal anchoring signal is absent or severely attenuated.

Intriguingly, the DIVA Late-N cluster (n = 9; peak position shifted beyond residue 40) was also strongly enriched in non-ER predictions: 6 of 7 Late-N proteins were predicted as non-ER, and all five plastid-predicted proteins carried chloroplast transit peptide signals and belonged to this cluster. This pattern suggests that a right-shifted DIVA peak may reflect a genuine alternative topological configuration—consistent with chloroplast outer-membrane targeting rather than ER retention—rather than a mere degradation of the canonical template.

#### DIVA and PIVOT Weak-N sets remain disjoint in localization behavior

Consistent with the mutual exclusivity of DIVA and PIVOT Weak-N assignments reported in Section 3.7, the two Weak-N sets showed non-overlapping localization profiles. All 11 PIVOT Weak-N proteins were predicted as ER-localized, whereas 5 of 8 DIVA Weak-N proteins were non-ER. This reinforces the interpretation that the two information channels are functionally separable: loss of the PIVOT identity signal does not compromise ER targeting, whereas loss of the DIVA topological signal strongly predicts mislocalization.

Together, these results demonstrate that the N-terminal sequence features governing the topological template model—particularly the integrity of the 1-8 anchoring segment and the physicochemical profile of the 9-40 transmembrane region—are specifically detected by a state-of-the-art localization predictor that was agnostic to our DIVA/PIVOT decomposition (Figure 18). The separable behavior of DIVA and PIVOT in the DeepLoc 2 analysis provides independent computational validation of the two-channel encoding framework.

## 4. Discussion

### 4.1 The N-terminus as a neutrally relaxed scaffold

PEACE analysis of the 50-residue N-terminal segment across 345 *S. tonkinensis* P450 paralogs reveals a constraint landscape characterized by near-uniformly elevated dispersion (mean = 0.62 ± 0.09). Only a single constrained island is detectable, spanning positions 20-27 in reference coordinates and corresponding to the WPVVGMLP motif. Within this island, Pro^21^ (dispersion = 0.356) and Gly^24^ (dispersion = 0.289) are each conserved at ≥98% across structurally confirmed sites. The remaining 84% of N-terminal residues show no evidence of purifying selection.

Embedding-based soft correspondence maps this island to median positions 44-51 in target sequences—at or beyond the 50-residue N-terminal boundary— confirming that it corresponds to a structural hinge lying downstream of the region analyzed by DIVA and PIVOT. Within the 50-residue segment proper, site-specific constraint is largely undetectable—the N-terminus approximates a neutrally evolving scaffold in the sense of the neutral theory (Kimura, 1983), with dispersion values (mean 0.62) consistent with near-neutral rather than strongly deleterious or strongly advantageous dynamics (Ohta, 1973).

This finding carries a methodological implication. The plant P450 N-terminus is an indel hotspot, with length variation spanning an order of magnitude (∼15 to >180 residues). In such regions, traditional MSA-based selection inference is structurally vulnerable: alignment algorithms must commit to arbitrary gap placements, and the resulting column-wise mismatches are interpreted by downstream dN/dS methods as substitution events, systematically inflating signals of positive selection (Fletcher & Yang, 2010; Spielman & Wilke, 2015). PEACE circumvents this by replacing column-wise homology with embedding-space soft correspondence. The finding of near-complete neutral relaxation across 84% of N-terminal residues suggests that previous reports of prevalent positive selection in this region should be interpreted with caution—they may predominantly reflect alignment noise rather than adaptive evolution.

### 4.2 Two information channels on a neutral scaffold

If the N-terminus is under virtually no sequence constraint, how does it encode functional information? Decomposition of the ESM2 embedding space reveals two functionally distinct signals—DIVA and PIVOT—that coexist within the same 50- residue segment yet are largely separable (Figure 19 A B).

**Figure 19.**
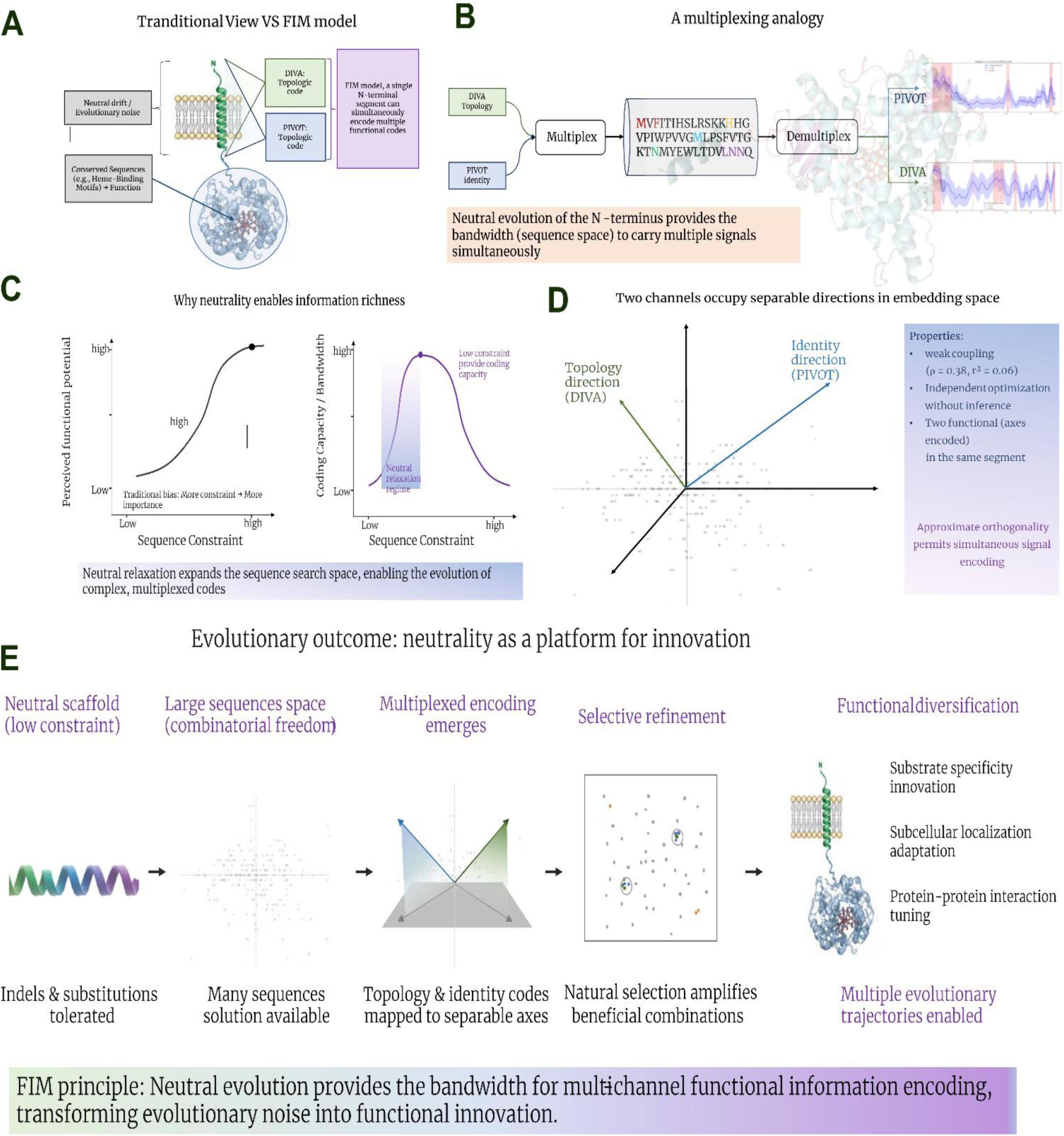
Conceptual framework for Functional Information Multiplexing (FIM) and holographic encoding in the P450 N-terminus. **A.** Comparison of the traditional linear encoding model and the FIM model. In the FIM framework, a single 50-amino-acid N-terminal segment serves as a holographic carrier, compressing global family identity signals (captured by PIVOT) and local topological signals (captured by DIVA) into a single scaffold. **B.** Multiplexing analogy representing the N-terminal carrier (only ∼10% of the full protein) as the physical medium for encoding two independent signal sources. **C.** FIM’s perspective on neutrality: an intermediate relaxation regime maximizes coding capacity by expanding the accessible sequence space. **D.** Functional orthogonality in ESM2 embedding space. The topology (DIVA) and holographic identity (PIVOT) codes map to approximately orthogonal directions with weak coupling *(p* = 0.38,r**_2_** = 0.06), enabling independent optimization. **E.** Evolutionary outcome. Neutral evolution at the N-terminus amplifies sequence diversity and enables the holographic mapping of global functional identity, which is subsequently refined by natural selection to drive substrate specialization, subcellular adaptations, and multi-trait evolutionary innovation.

DIVA, derived from unsupervised decomposition without functional labels, captures membrane-anchor-associated variation concentrated at the extreme N-terminus. This signal likely reflects topological constraints common to all microsomal P450s: the N-terminal transmembrane helix and its orientation relative to the ER membrane. Consistent with this interpretation, DIVA scores correlate systematically with established physicochemical properties such as hydrophobicity and helical propensity (Figure 14 A). Its emergence without explicit labeling is consistent with the observation that protein language models learn structural features from sequence data alone (Rives et al., 2021).

PIVOT, derived from supervised decomposition, captures a family-associated semantic signal that is systematically offset from the DIVA peak. Critically, the classifier underlying PIVOT was trained not on CYP family labels—the conventional sequence-based classification—but on 12 functional clusters defined by ESM2 embedding similarity. This design avoids circularity: the classifier learned to separate clusters defined purely by embedding geometry, never having been taught CYP family labels. Yet the per-residue importance scores produced by this classifier (PIVOT) prove capable of distinguishing CYP families: PIVOT peak positions are significantly associated with family identity (Kruskal-Wallis H = 117.93, p = 5.3 × 10^−13^, ε^2^ = 0.44, large effect). That the resulting importance scores carry family- discriminatory information indicates that the embedding space itself organizes P450 sequences along functional axes that partially align with—but do not reduce to—the traditional family classification.

The nature of this alignment is instructive. A permutation test confirmed that the association was not an artifact of group size imbalance: in 0/10,000 random shuffles of family labels did H exceed the observed value (empirical p < 0.0001; H_obs_/null mean = 4.24×). However, Dunn post-hoc tests with Holm correction identified only 2 of 406 (0.5%) pairwise family comparisons as significant. This pattern—highly significant omnibus association coupled with near-complete absence of significant pairwise differences—is the statistical signature of a continuous gradient, not a categorical partition. PIVOT peak positions vary systematically across CYP families, but they do so along a spectrum without discrete boundaries. This is consistent with functional diversification through gradual substrate specialization (Hamberger & Bak, 2013; Hansen et al., 2021) and suggests that the N-terminal semantic center-of-gravity is a spectral, non-categorical family trait.

Across 334 paralogs, PIVOT peaks fall downstream of DIVA peaks in 84.4% of cases, with only 10.2% showing close alignment (|Δ| ≤ 0.15 in normalized coordinates). The statistical coupling is weak (Spearman ρ = 0.38, r^2^ = 0.06). The PIVOT signal is not a byproduct of differential conservation: PEACE constraint scores show no significant correlation with PIVOT magnitude at any position (|ρ| < 0.24, p > 0.09).

Further evidence for channel independence comes from the Weak-N classification (Section 3.7). Eleven proteins (3.2%) are PIVOT Weak-N: they possess a clear DIVA topological template but lack any prominent PIVOT peak within the N-terminal region. An additional eight proteins are DIVA Weak-N: they lack the topological template while retaining PIVOT identity signal. The two Weak-N sets are completely mutually exclusive (zero overlap). This natural double dissociation— DIVA without PIVOT, and PIVOT without DIVA—confirms the two signals can exist independently.

### 4.3 The N-terminus as an integration point of selective pressures

A central paradox emerges from the PIVOT analysis: the N-terminal 50 residues—comprising merely ∼10% of the average P450 sequence and evolving under near-complete neutral relaxation—encode sufficient information to distinguish CYP families with an accuracy comparable to the full-length protein. CYP family classification is conventionally operationalized through full-length sequence homology, a phylogenetic criterion dominated by the conserved catalytic core. How can the most rapidly evolving, indel-riddled segment of the protein carry the most diagnostic signal for a classification system built on whole-protein similarity?

The resolution requires distinguishing between homology and information. The neutral theory predicts that unconstrained regions accumulate variation at the mutation rate (Kimura, 1968), but Ohta’s nearly neutral theory (Ohta, 1973) further implies that such regions can carry slight functional effects that drift to fixation— providing a theoretical basis for the coexistence of apparent neutrality and functional information. Full-length homology comparisons are dominated by the most constrained regions—the catalytic core, the heme-binding motif, and the substrate recognition sites—precisely because these align most reliably across divergent sequences. However, residues invariant across all P450s carry no family-specific information; they indicate that a sequence *is* a P450, but not *which* P450. The N-terminus, liberated from the structural constraints that immobilize the catalytic core, explores a broader region of sequence space, capturing a signal that full-length homology comparisons systematically obscure.

This accounts for why the core is uninformative, but not why the N-terminus— physically distanced from the catalytic residues—should carry family- discriminatory information at all. The answer lies in the N-terminus functioning as a unique integration point for distinct selective pressures.

Unlike other variable regions, such as flexible linkers which may drift neutrally, the N-terminus is physically positioned to simultaneously respond to constraints from two disparate domains: the cellular environment, which demands correct ER anchoring via hydrophobicity and helical propensity (the DIVA signal), and the protein’s functional core, whose substrate specificity dictates the precise membrane approach geometry required for catalysis (the PIVOT signal). Consequently, the N-terminal sequence is actively sculpted by a multi-dimensional fitness landscape defined by the dual demands of efficient membrane insertion and optimal catalytic orientation. For a CYP family specialized for a specific substrate class, the optimal N-terminal sequence—the one that best balances anchoring stability with catalytic accessibility—will differ systematically from that of another family.

The N-terminus thus does not passively record co-evolutionary history; it actively integrates the selective pressures that define the protein’s function. It is the physical projection of the catalytic domain’s identity onto the membrane interface. Just as a keystone bears the imprint of the forces acting upon an arch, the N-terminus bears the physical signature of the functional integration required for that specific P450 to operate within the cell. This explains why PIVOT—trained on embedding clusters that carry no explicit CYP family labels—nonetheless distinguishes families: it detects the specific sequence configurations that represent the optimal solution to this integration problem for each family. This principle of multi-objective optimization at a physical integration point provides the mechanistic foundation for the multiplexed encoding framework developed below.

The spatial separation of DIVA and PIVOT, together with the PxxG hinge position, reveals how these information channels are physically organized. Three sequentially arranged functional layers can be distinguished. **Layer I — Topology (DIVA, ∼1-20 aa):** The DIVA signal is concentrated at the extreme N-terminus, consistent with the established role of the first ∼20 residues in ER membrane targeting and insertion (Schuler & Werck-Reichhart, 2003; Hansen et al., 2021). Its emergence without functional labels suggests that the embedding space spontaneously organizes around membrane-anchor features. **Layer II — Identity (PIVOT, ∼20—50 aa):** The PIVOT signal occupies the middle portion of the 50- residue segment, with its peak centered at 25-28 (Mid-N1 84.6%) Critically, this layer operates within the neutrally relaxed portion of the scaffold (PEACE dispersion ∼0.62), meaning that family-associated encoding is achieved through multi-objective optimization captured by the embedding space rather than through sequence-level constraint. **Layer III — Structure (PxxG hinge, ∼44-51 aa, soft- corresponded):** The PxxG hinge maps via soft correspondence to positions at or beyond the 50-residue boundary. It is the only constrained island in an otherwise neutrally relaxed scaffold, consistent with a structural transition region demarcating the N-terminal regulatory segment from the catalytic domain.

This layered architecture provides a mechanistic basis for the hypothesis that the N-terminus functions as an evolutionary localization switch (Schuler & Werck-Reichhart, 2003). The near-complete separation of DIVA from PIVOT (84.4% downstream offset, r^2^ = 0.06) implies that the topology template and the family-associated signal can vary quasi-independently—the ability to alter *where* a P450 goes without necessarily altering *what* it does. The neutral relaxation of the scaffold (PEACE dispersion = 0.62) provides the mutational tolerance necessary for the topology template to explore alternative targeting configurations without disrupting the identity signal layered onto the same segment. This dual-channel architecture may explain why the N-terminus, despite being the most variable region of plant P450s, is also the most information-rich: evolutionary permissiveness enables simultaneous encoding of location and function (Figure 19 D).

### 4.4 Functional Information Multiplexing and the holographic organization of protein sequence information

The convergence of these findings motivates a framework termed **Functional Information Multiplexing (FIM)**: the principle that multiple, functionally independent codes can coexist within a single short peptide segment. By expanding the accessible sequence space, the evolutionary permissiveness of the scaffold provides the combinatorial degrees of freedom necessary for these codes to coexist without mutual interference. FIM thus proposes that neutrality is not merely an absence of constraint, but a substrate for encoding complexity (Figure 19 C).

FIM departs from the assumption that functional information requires sequence conservation. In the P450 N-terminus, information resides in the combinatorial organization of the embedding space—the distribution, sharpness, and relative positions of signal peaks—rather than in site-wise amino acid identity. A tightly constrained sequence would force the channels to covary, collapsing the multiplexed architecture.

FIM is distinct from related concepts. Unlike moonlighting proteins, where a single sequence performs different functions in different contexts (Jeffery, 1999), FIM describes simultaneous encoding of multiple signals in the same molecular scaffold. Unlike gene sharing (Piatigorsky, 2007), where a single coding sequence is maintained by pleiotropic constraints, FIM operates on a segment largely unconstrained and free to explore sequence space. FIM also occupies a distinct conceptual space relative to the robustness-evolvability framework (Wagner, 2005), which demonstrates how neutral networks enable the accumulation of cryptic genetic variation as a reservoir of potential phenotypic diversity. FIM extends this logic: neutrality can be more than a reservoir of unrealized potential; it can be actively exploited by natural selection to construct a concurrent multi-channel encoding architecture. The P450 N-terminus is not waiting for a future mutation to unlock a hidden function; it is already transmitting two distinct functional signals simultaneously on the same physical scaffold. The transition is from “neutrality as reservoir” to “neutrality as bandwidth.”

This transition is possible because neutrality creates the combinatorial degrees of freedom necessary for separable encoding—an evolutionary consequence anticipated, in part, by the neutral (Kimura, 1983) and nearly neutral (Ohta, 1973) theories, which established that the majority of molecular variation is selectively neutral or nearly so. When a single peptide segment faces two functionally independent demands—a hard biophysical constraint (membrane insertion) and a soft adaptive constraint (family-specific functional tuning)—natural selection can, over evolutionary time, discover and amplify sequence configurations that map these two demands onto approximately separable directions in the embedding space. Here “separable” refers to high-dimensional geometric near-independence: the two functional demands can be optimized along distinct axes of sequence variation without mutual interference, consistent with their weak statistical coupling (ρ = 0.38, r^2^ ≈ 0.06; Section 3.7). As described in Section 4.3, the N-terminus serves as the physical integration point where these selective pressures are reconciled; separable encoding explains how this reconciliation is achieved without mutual interference.

The P450 N-terminus may represent an extreme but not unique case of FIM. Signal peptides, transmembrane helices, and intrinsically disordered regions—all indel-dense, low-constraint segments—are candidates for similar multiplexed architectures. Ultimately, neutrality provides the bandwidth for evolution to explore adaptive landscapes. As summarized in Figure 19E, this multiplexed scaffold is subjected to selective refinement, leading to functional diversification in substrate specificity, subcellular targeting, and protein-protein interactions.

This framework invites a conceptual reframing of how protein sequences encode functional information. Historically, sequences have been treated as linear assemblies of independent modules—a “beads-on-a-string” model in which function is localized to specific, conserved motifs. However, these findings, together with the recognition of pervasive co-evolutionary couplings across protein structures (Marks et al., 2011), point toward a more integrated architecture. A holographic metaphor is instructive: in a hologram, a fragment of the recording does not contain a smaller version of the full image, but rather encodes the entire image at reduced signal-to-noise ratio, because information is distributed across the recording surface through interference patterns. The N-terminal segment of P450s operates analogously—it does not contain a local, compressed copy of the catalytic domain’s functional signature, but participates in the distributed encoding of global functional identity through high-order epistatic couplings spanning the entire protein sequence. These couplings are invisible to linear alignment but recoverable by the deep learning architectures of protein language models.

The holographic perspective on protein sequence encoding is grounded in a fundamental shift in how non-conserved regions are viewed in protein evolution. Historically, these regions—encompassing linkers, loops, and terminal segments— were dismissed as evolutionary noise or mere structural spacers. However, the recognition of intrinsically disordered regions as functional hubs (Dyson & Wright, 2005; Tompa, 2012), the identification of the protein periphery as the primary site of evolutionary innovation (Tokuriki & Tawfik, 2009), and the discovery of high-order feature convergence in protein evolution (Cao et al., 2025) have collectively overturned this view. Non-conserved regions are not evolutionary dead ends; they are the dynamic interfaces where functional constraints are negotiated and where lineage-specific adaptations are encoded. The N-terminus of P450s exemplifies this principle: its sequence variability is not noise, but the physical substrate for integrating membrane topology and catalytic identity. Protein language models, by capturing the distributed epistatic couplings embedded in these regions (Rives et al., 2021), provide the computational lens through which this holographic information becomes visible.

This holographic perspective provides a unifying framework for the three observations central to this study. It explains why the least constrained region of the protein can be the most information-rich (Section 4.3): in a distributed encoding scheme, information capacity is not tied to local conservation. It explains why multiplexed encoding is possible: separable information channels can coexist as distinct interference patterns in the same physical segment without mutual disruption. And it explains why alignment-free, embedding-based methods succeed where traditional MSA-based approaches fail: the information resides in statistical couplings—in the relationships between residues—not in site-wise identity.

### 4.5 Methodological implications

Beyond the biological findings, this study demonstrates a broader point about the analysis of indel-dense protein regions. Such regions are common in eukaryotic proteomes—signal peptides, linker domains, activation loops, viral envelope proteins—and, far from being evolutionary noise, harbor indels that are themselves subject to selective constraint (de la Chaux et al., 2007)—yet they have remained systematically challenging for MSA-based evolutionary analysis. This challenge reflects a limitation of the dominant paradigm in sequence analysis, which has treated protein sequences primarily as texts—linear strings of letters to be aligned, compared, and searched for conserved motifs. This approach has been enormously productive, but in indel-dense, rapidly evolving regions, the implicit requirement of align ability becomes difficult to satisfy. The sequence becomes difficult to interpret not because it carries no information, but because the tools presuppose a format it no longer obeys (Wong et al., 2008; Fletcher & Yang, 2010).

The present study demonstrates a complementary analytical perspective: sequences as signals. Protein language models serve as the critical bridge, converting discrete amino acid tokens into continuous embedding-space coordinates that preserve contextual information. Once a sequence is represented as a multidimensional signal along the primary structure axis, signal processing tools—peak detection, correlation analysis, offset estimation, sharpness quantification—become available for functional dissection. DIVA and PIVOT function as demodulators, extracting two approximately separable information channels from the same physical carrier. The traditional question “which residues are conserved?” gives way to new questions: “how many functionally independent signals share this segment?” “what are their relative peak positions and sharpness?” “how strongly do they couple?” This analytical reframing—from aligning texts to processing signals—is operational and quantitative, particularly suited to short, variable peptide segments that have long resisted traditional methods. The combination of alignment-free constraint estimation (PEACE) and embedding-space signal decomposition (DIVA/PIVOT) enables the systematic discovery of multiplexed architectures in regions of the proteome that have, until now, been dismissed as uninformative.

### 4.6 Implications for N-terminal engineering

The layered architecture and channel separation documented here have practical implications for P450 heterologous expression, a persistent challenge in plant metabolic engineering. Current N-terminal modification strategies— truncation, signal-anchor replacement, codon optimization—proceed largely by trial and error. The DIVA/PIVOT framework suggests a design principle: engineer the topology template (DIVA) to optimize membrane insertion and expression yield, while preserving or transplanting the identity signal (PIVOT) to maintain catalytic specificity. CYP73A and CYP98A—whose N-terminal modifications are documented to affect expression and activity—may serve as natural validation cases (Renault et al., 2017; Leonard & Koffas, 2007; Schoch et al., 2003; Hotze et al., 1995). Both show clear DIVA-PIVOT separation in the present dataset, suggesting that their engineering sensitivity reflects disruption of one channel while the other remains intact. Systematic experimental testing remains a target for future work.

### 4.7 Limitations

Several limitations warrant mention. First, PEACE quantifies relative constraint within a paralog set and does not provide an absolute scale comparable across protein families. The dispersion values reported here are interpretable within the *S. tonkinensis* P450 superfamily.

Second, soft correspondence depends on embedding-space quality; the 11% failure rate in PxxG mapping illustrates the boundary where deeply divergent embeddings separate onto distinct manifolds, limiting reliable soft correspondence to within-genus or within-family comparisons.

Third, the FIM framework is observational. While the three information layers are statistically and functionally separable, direct experimental demonstration that altering one layer leaves the others intact remains for future work.

Fourth, sequence-level scanning confirms broad fixation across vascular plants (Section 3.4.5), but PEACE embedding-level decomposition has so far been applied only to *S. tonkinensis*.

Fifth, the signal-processing framework developed here depends critically on the quality and evolutionary representativeness of the underlying embedding space. Exploratory analyses with alternative PLM architectures (e.g., the BERT-based VenusPLM, Li M et al., 2025) yielded results that differed substantially from ESM2, suggesting that DIVA-PIVOT separability is not an invariant property of any protein language model. A likely explanation lies in the training data. ESM2 was trained on the UniRef50/90 and BFD metagenomic databases (Lin et al., 2023), which provide broad, unbiased coverage of natural sequence diversity—including environmental microorganisms, archaea, and non-model eukaryotes—without requiring functional annotation. In contrast, many other PLMs were trained primarily on curated databases enriched for model organisms and functionally annotated proteins. Embeddings from ESM2 are therefore more likely to capture the structural and physicochemical constraints upon which evolutionary diversification operates, whereas embeddings from annotation-biased models may be more strongly aligned with functional categories. For evolutionary studies that seek to recover multiplexed architectural principles—rather than to predict functional labels—the unbiased diversity captured by ESM2’s training data may be essential. Whether the FIM framework extends to other PLMs, and under what training regimes, remains an open question.

## 5. Conclusion

This study provides a revised view of the plant P450 N-terminus. Rather than a conserved core surrounded by adaptively evolving surfaces, it is a neutrally relaxed scaffold hosting two separable information channels—one encoding membrane topology, the other encoding family-associated identity—organized in a layered architecture of topology, identity, and structure. The finding that the least constrained region of the protein (84% neutral relaxation) simultaneously serves as the most information-rich—compressing family-level functional identity into merely 50 amino acids—motivates the concept of Functional Information Multiplexing: the principle that evolutionary permissiveness, by expanding the accessible sequence space, provides the combinatorial capacity for multiple functional codes to coexist without mutual interference.

This principle carries implications beyond the P450 superfamily. If neutrality is not the absence of information but the precondition for its multiplexed encoding, then many regions of the proteome previously dismissed as unconstrained noise— signal peptides, intrinsically disordered segments, viral envelope proteins—may warrant re-examination as potential carriers of layered functional codes. The computational framework demonstrated here, combining alignment-free constraint estimation with embedding-space signal decomposition, provides a transferable toolkit for such re-examination. Sequences are not aligned; they are processed as signals—and in doing so, information may be found where traditional methods see only noise.

## Supporting information

Full Alignemnts

