## Supplementary figures and images for "The N-Terminus of *Sophora tonkinensis* Cytochrome P450s Evolves Neutrally yet Encodes Rich Functional Information: A Protein Language Model Analysis"

### Figure S1

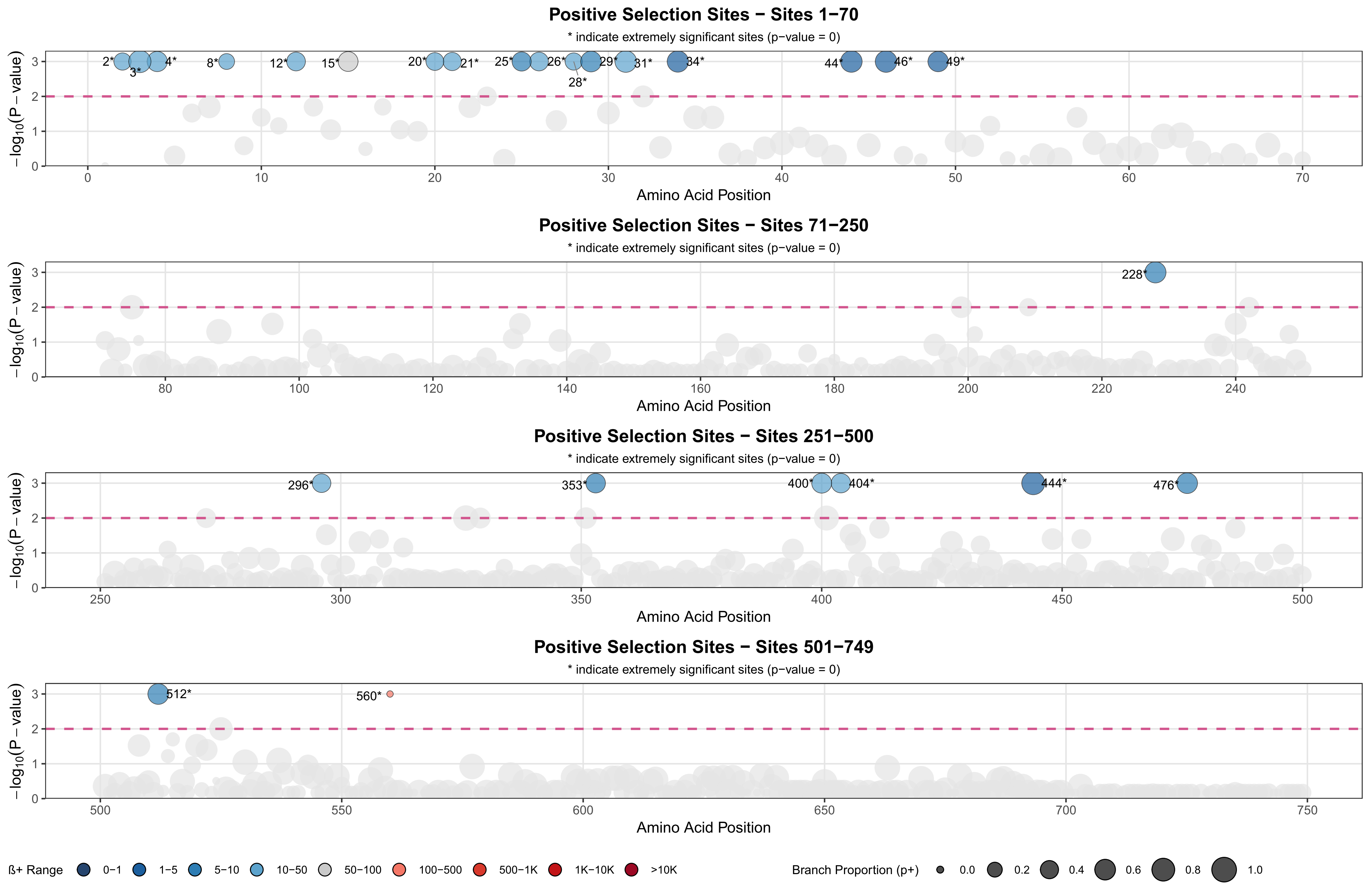

### Figure S2

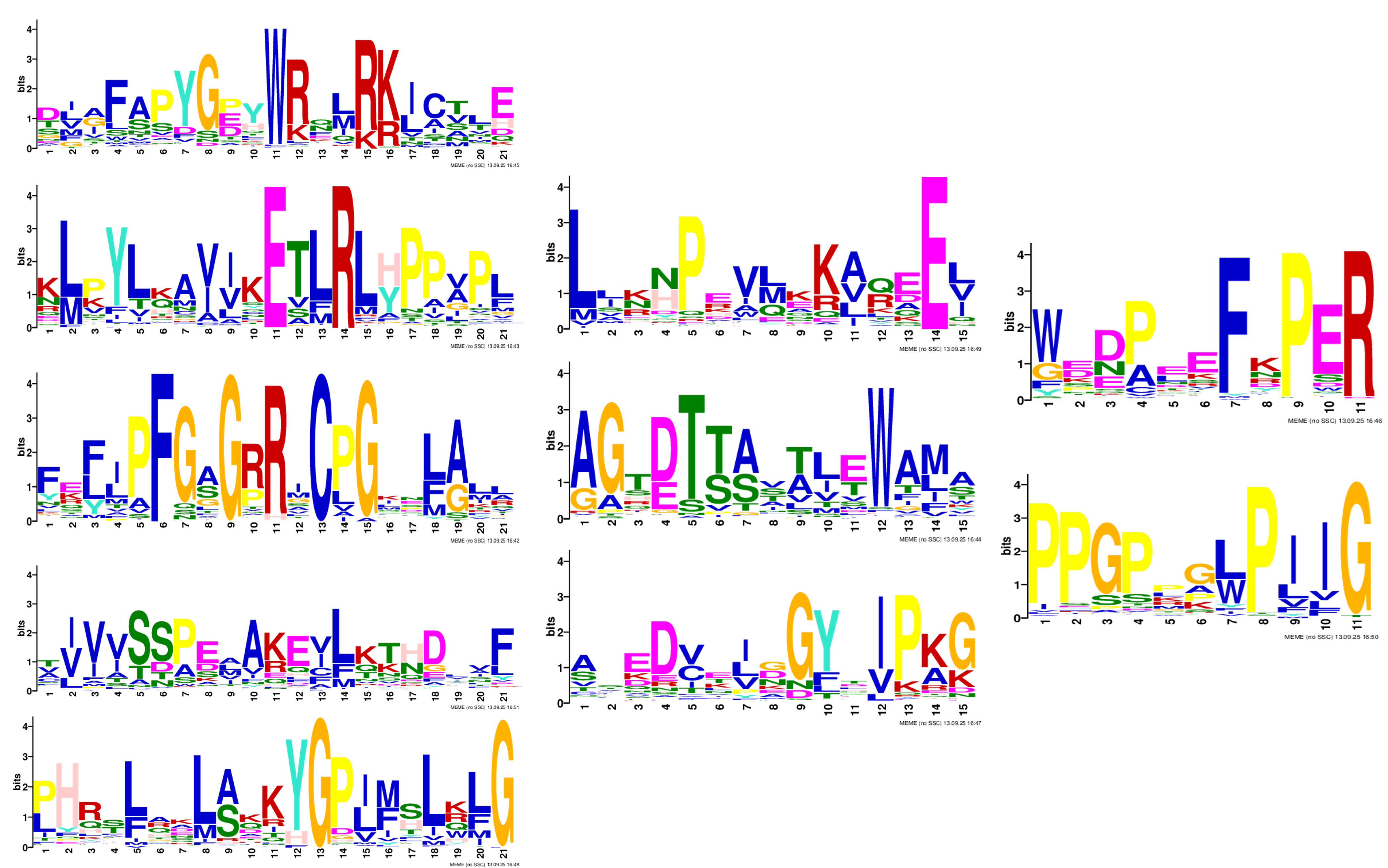

### Figure S3

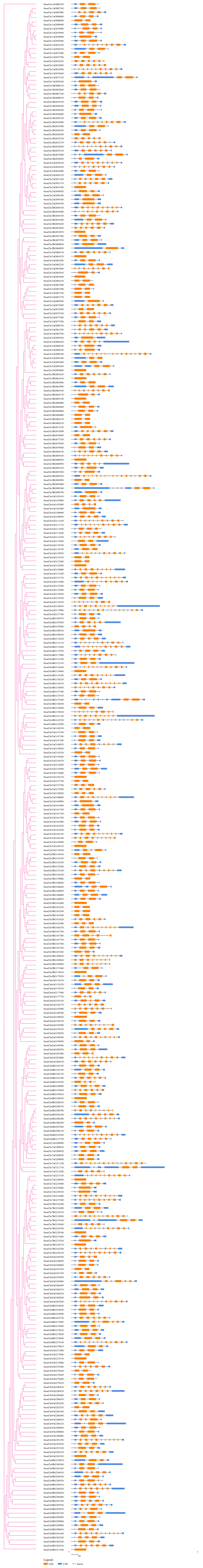

### Figure S4

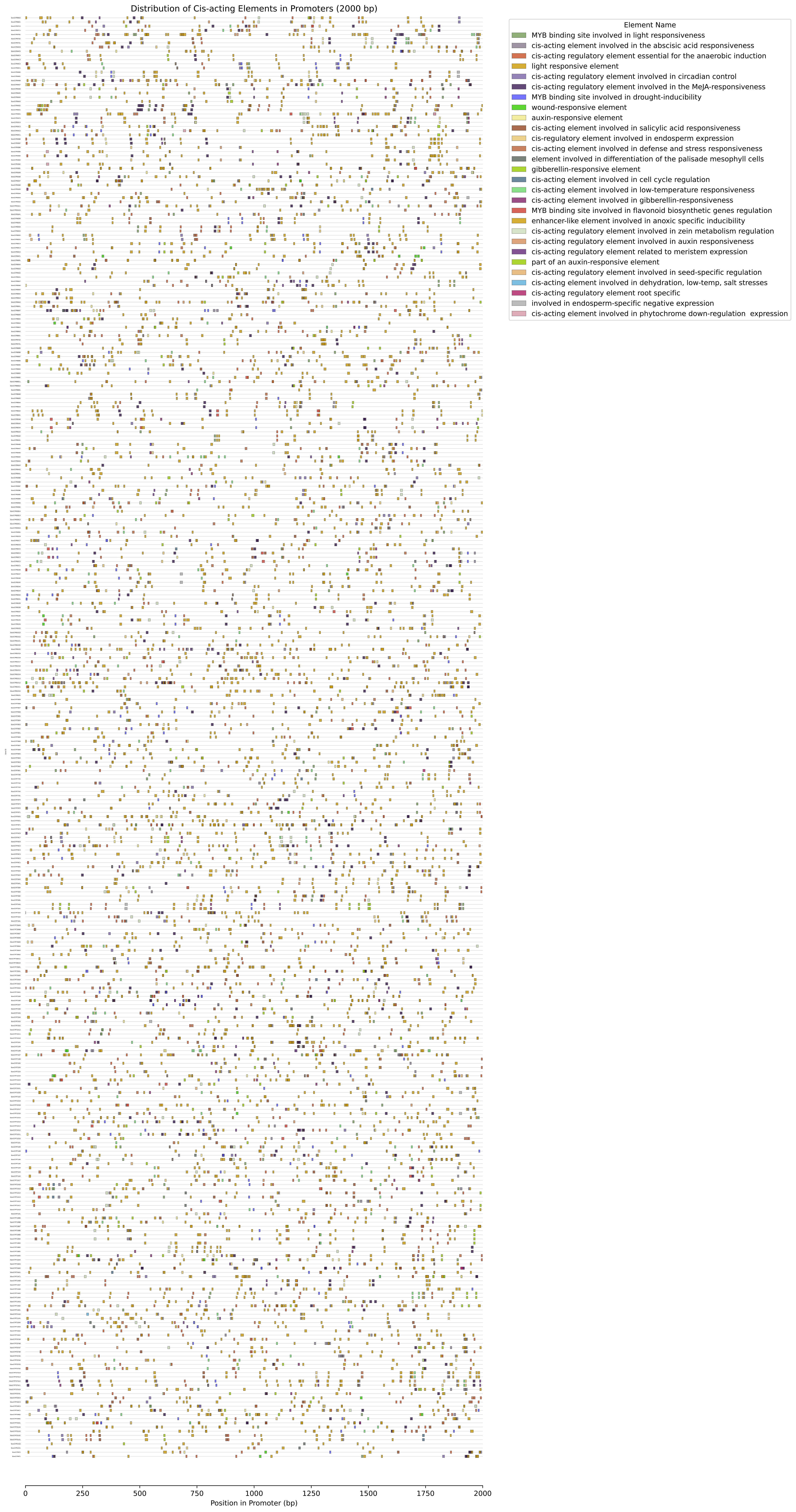

### Figure S5

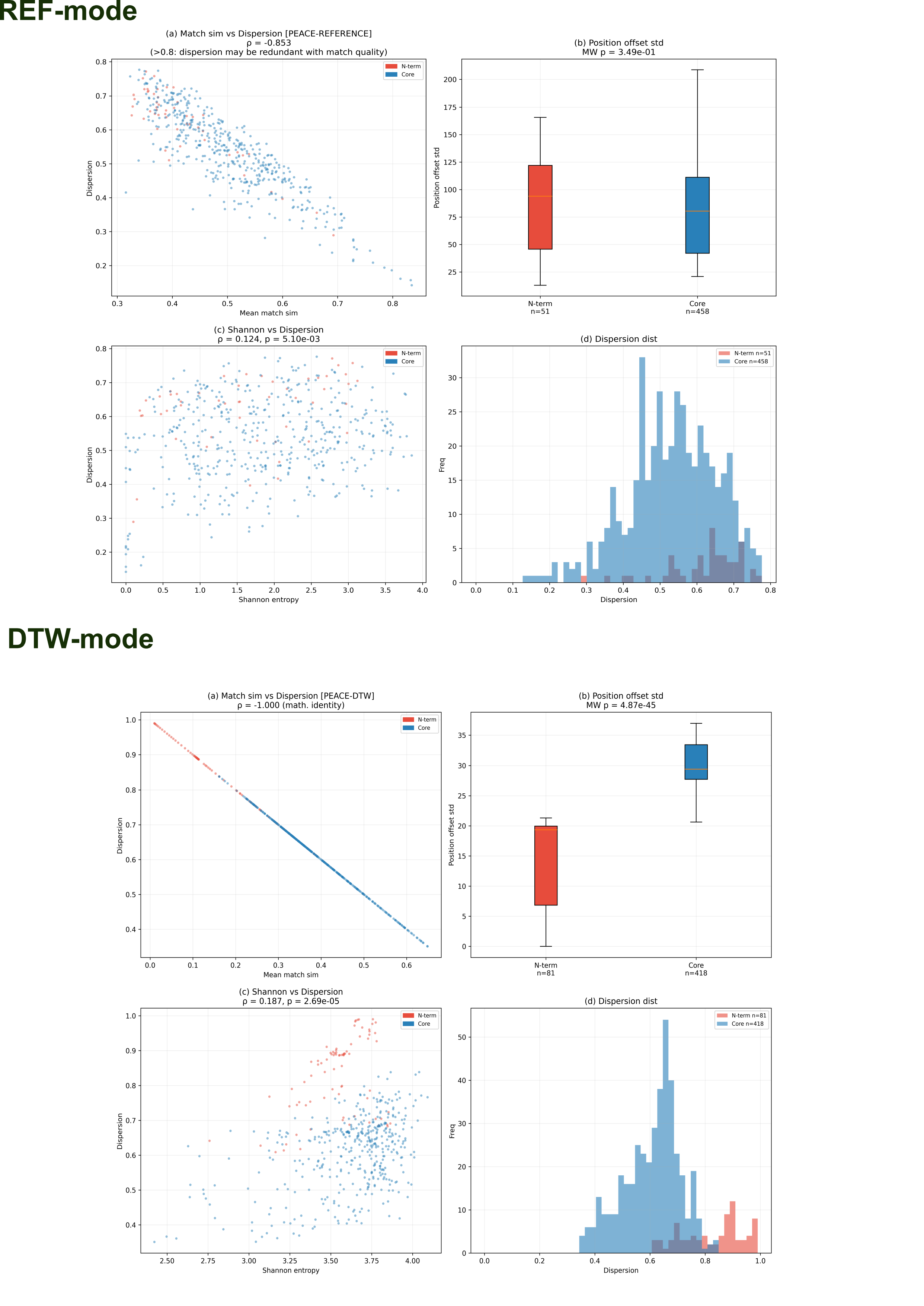

### StonChr2BG045330.1_importance.pdf

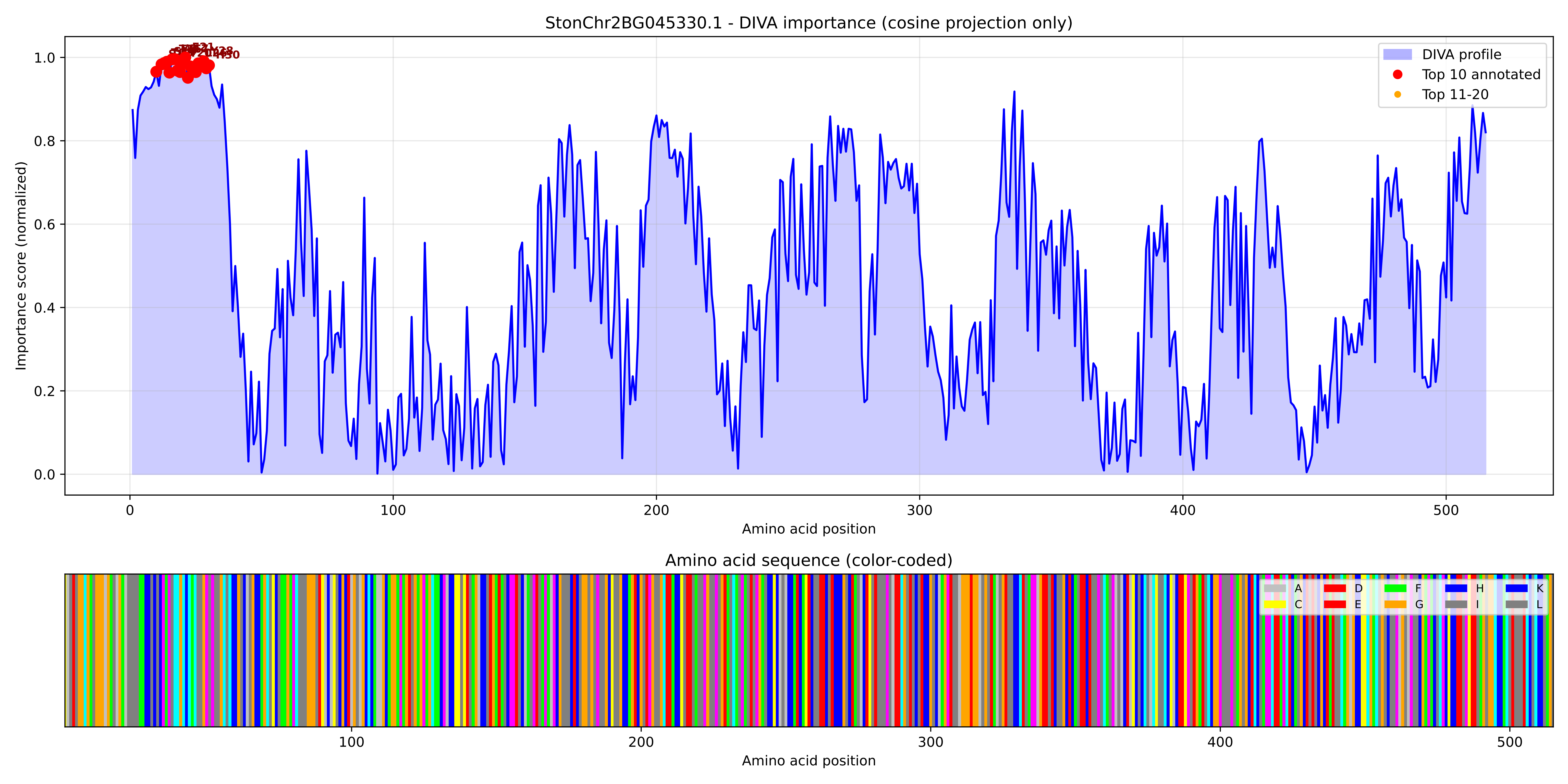

### StonChr2BG045330.1_importance.png

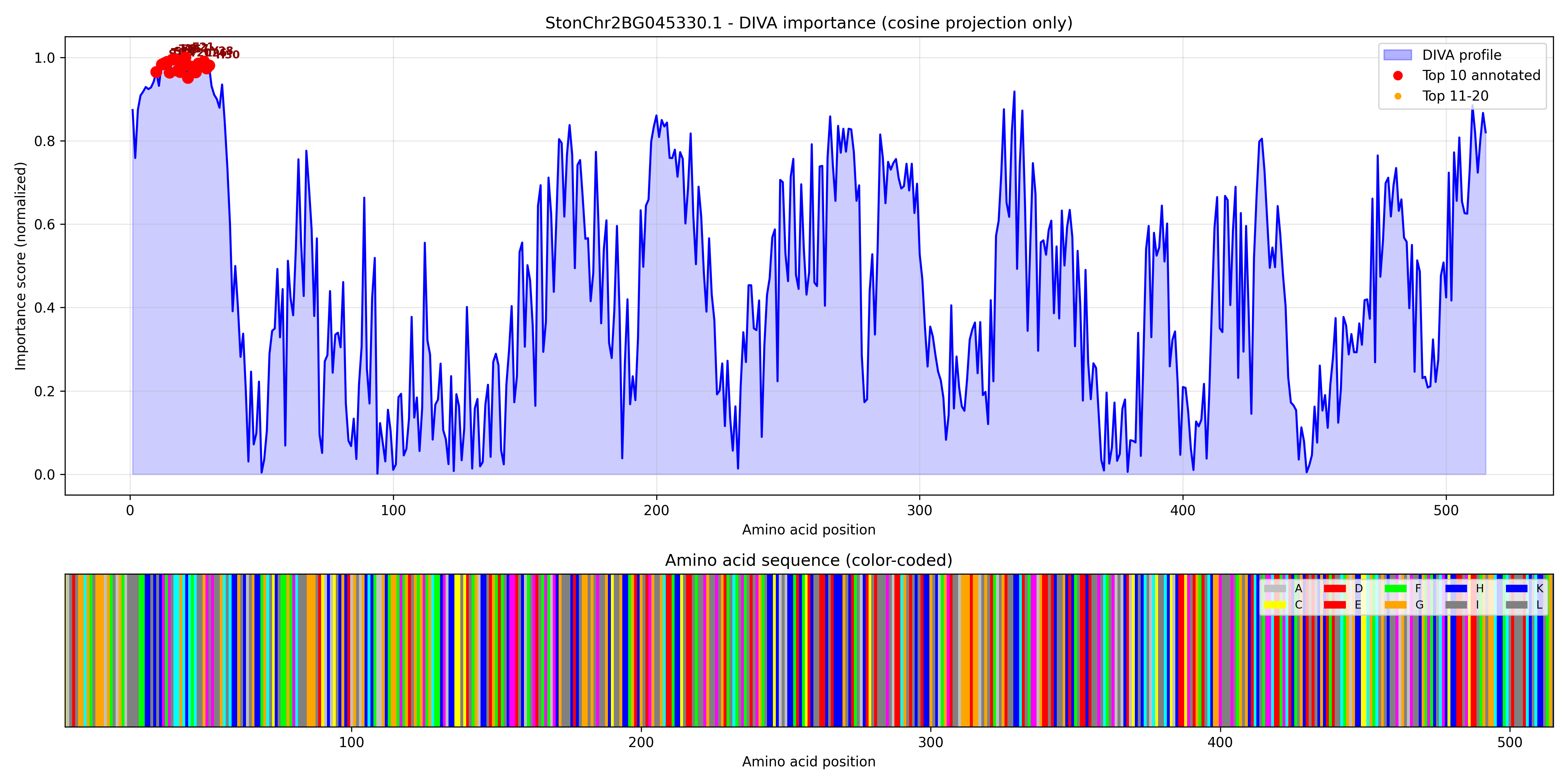

### StonChr2BG045380.1_importance.pdf

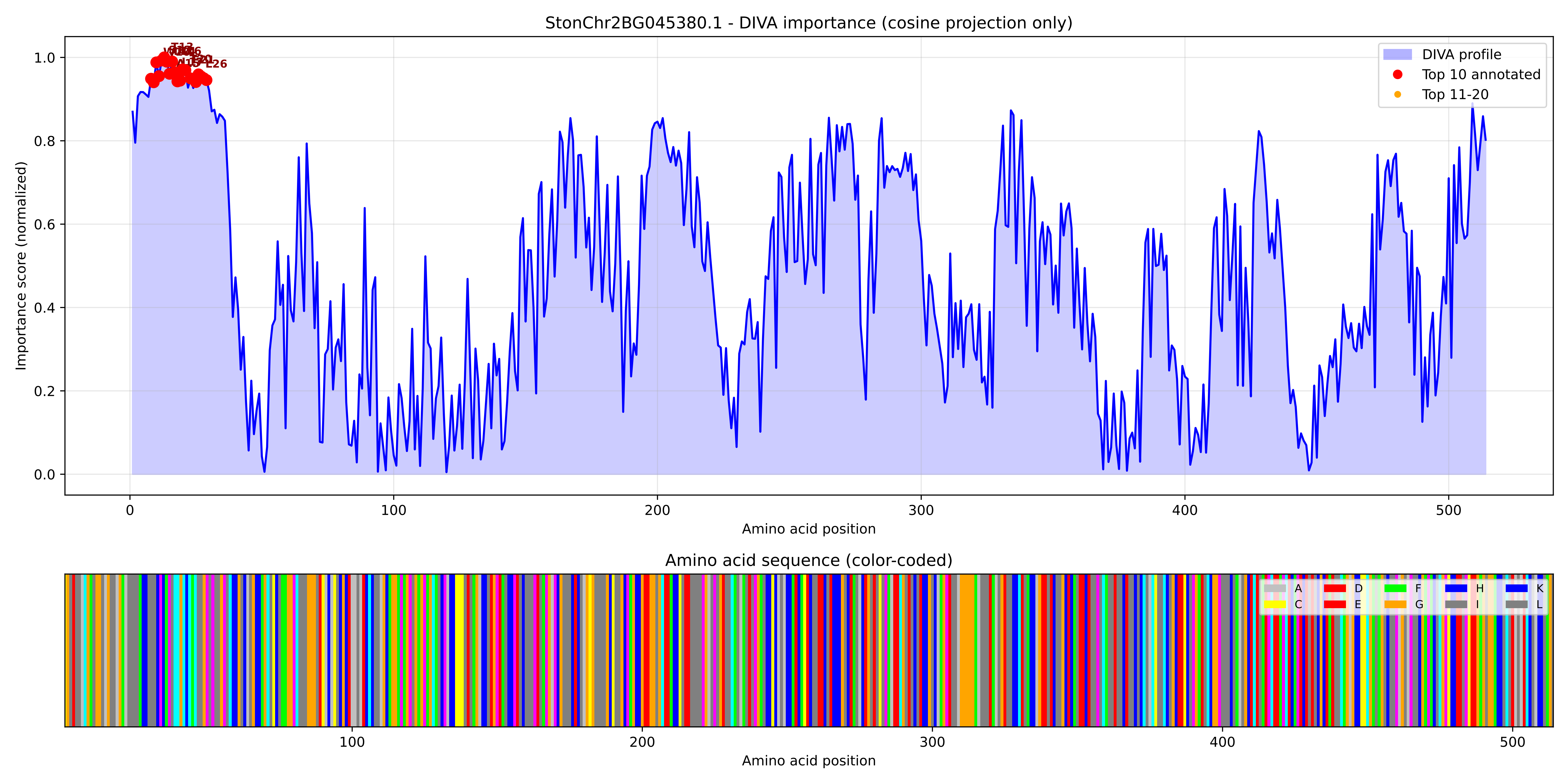

### StonChr2BG045380.1_importance.png

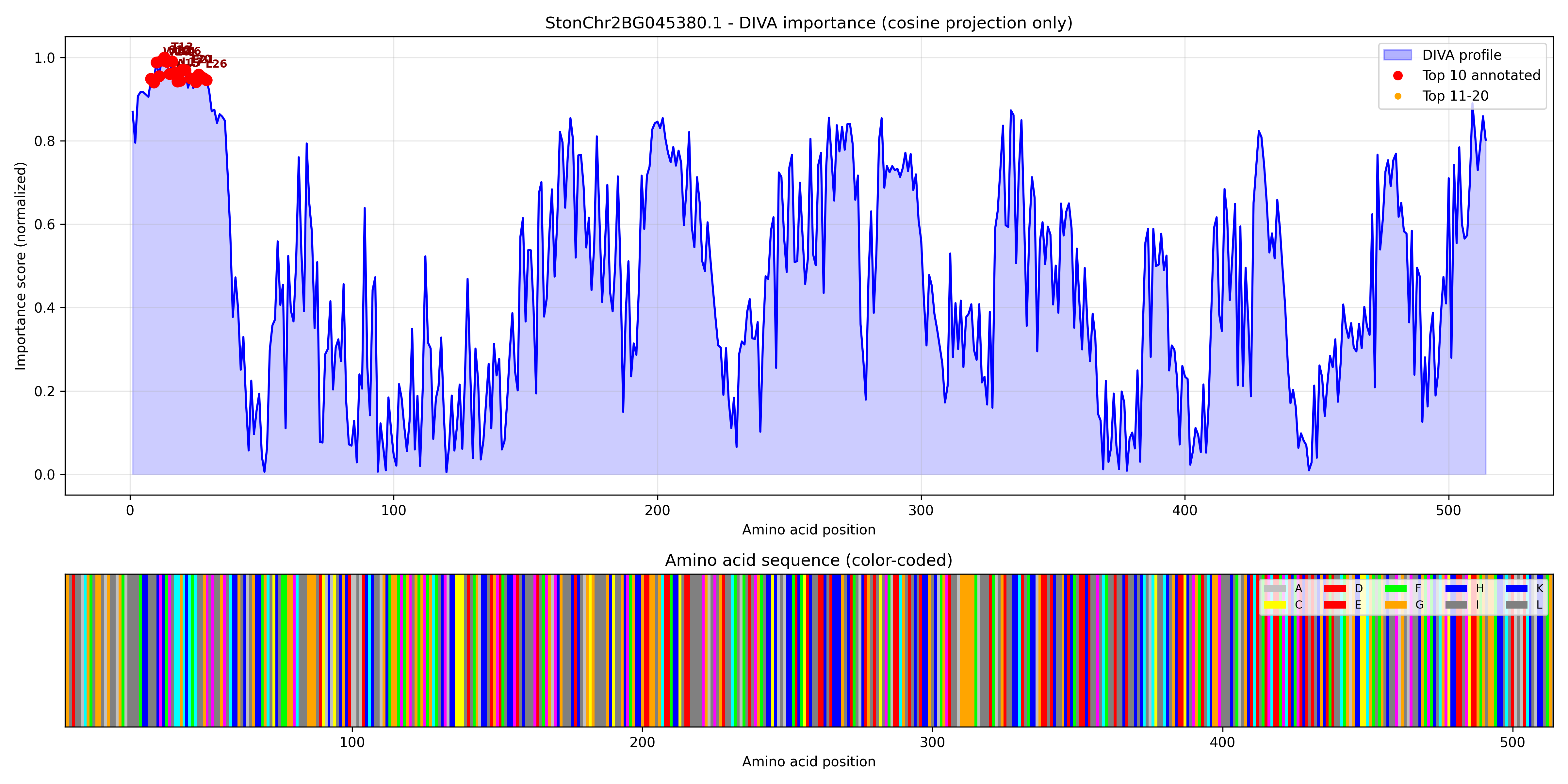

### StonChr2BG052620.1_importance.pdf

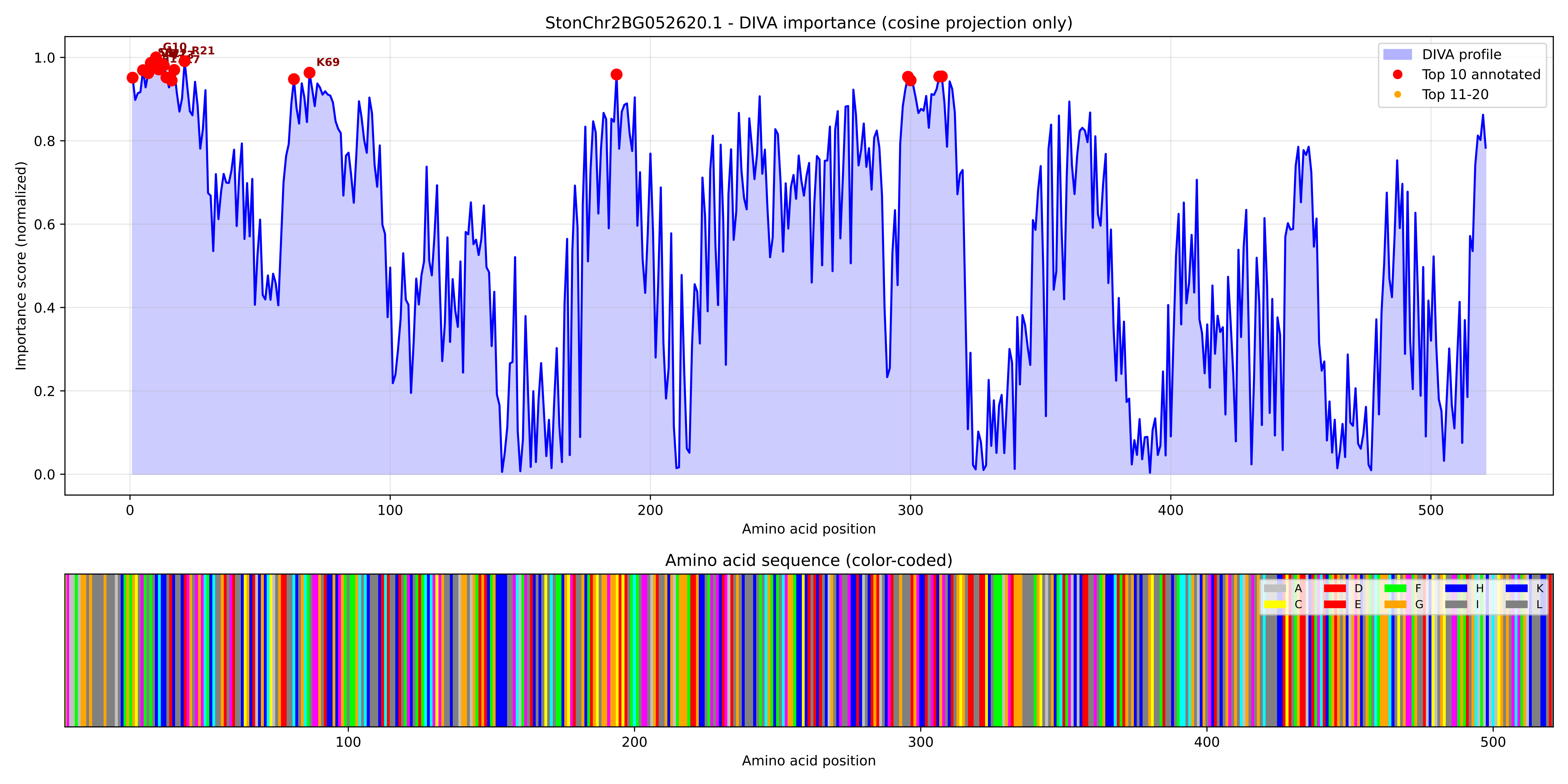

### StonChr2BG052620.1_importance.png

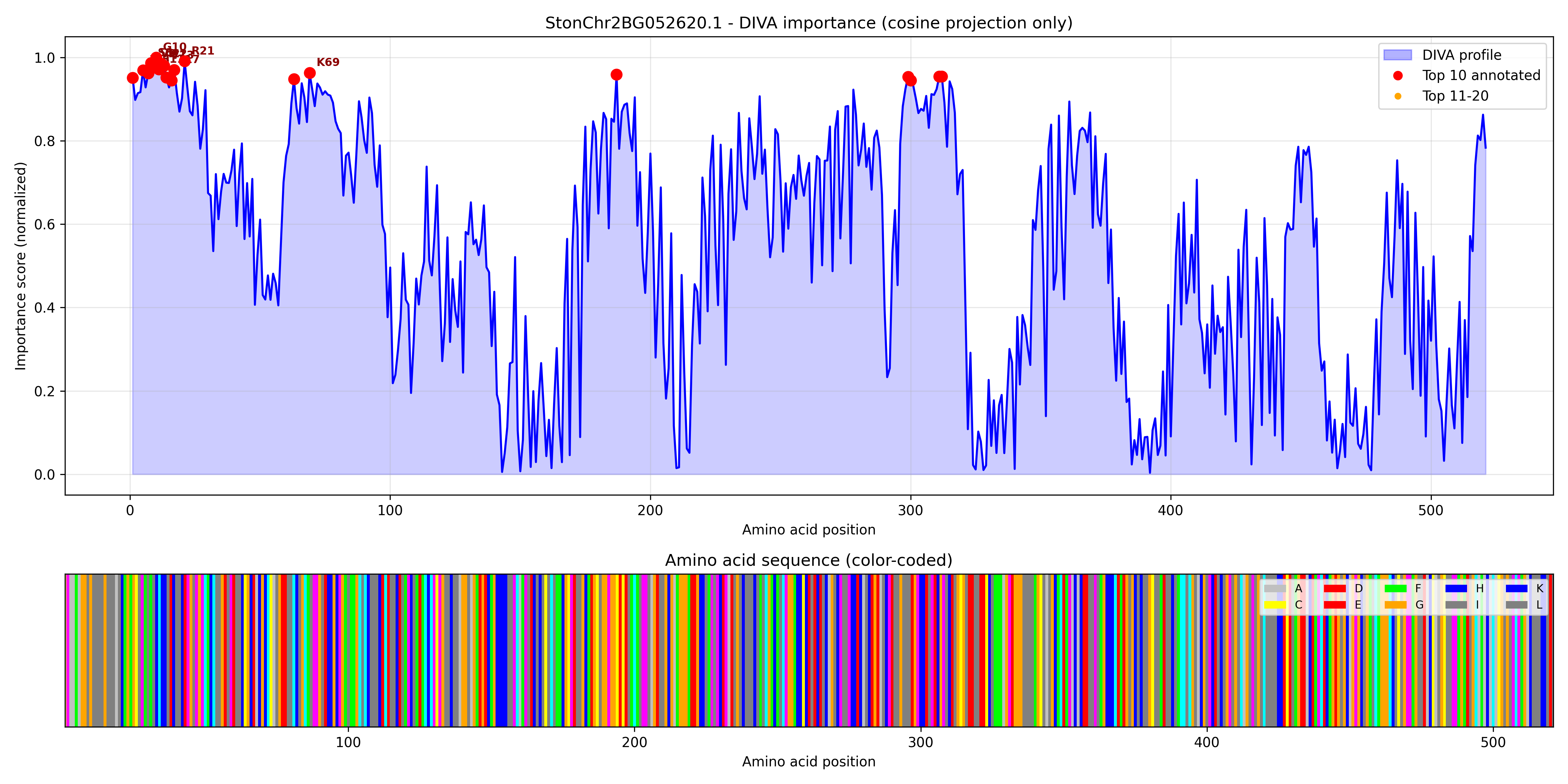

### StonChr2BG052630.1_importance.pdf

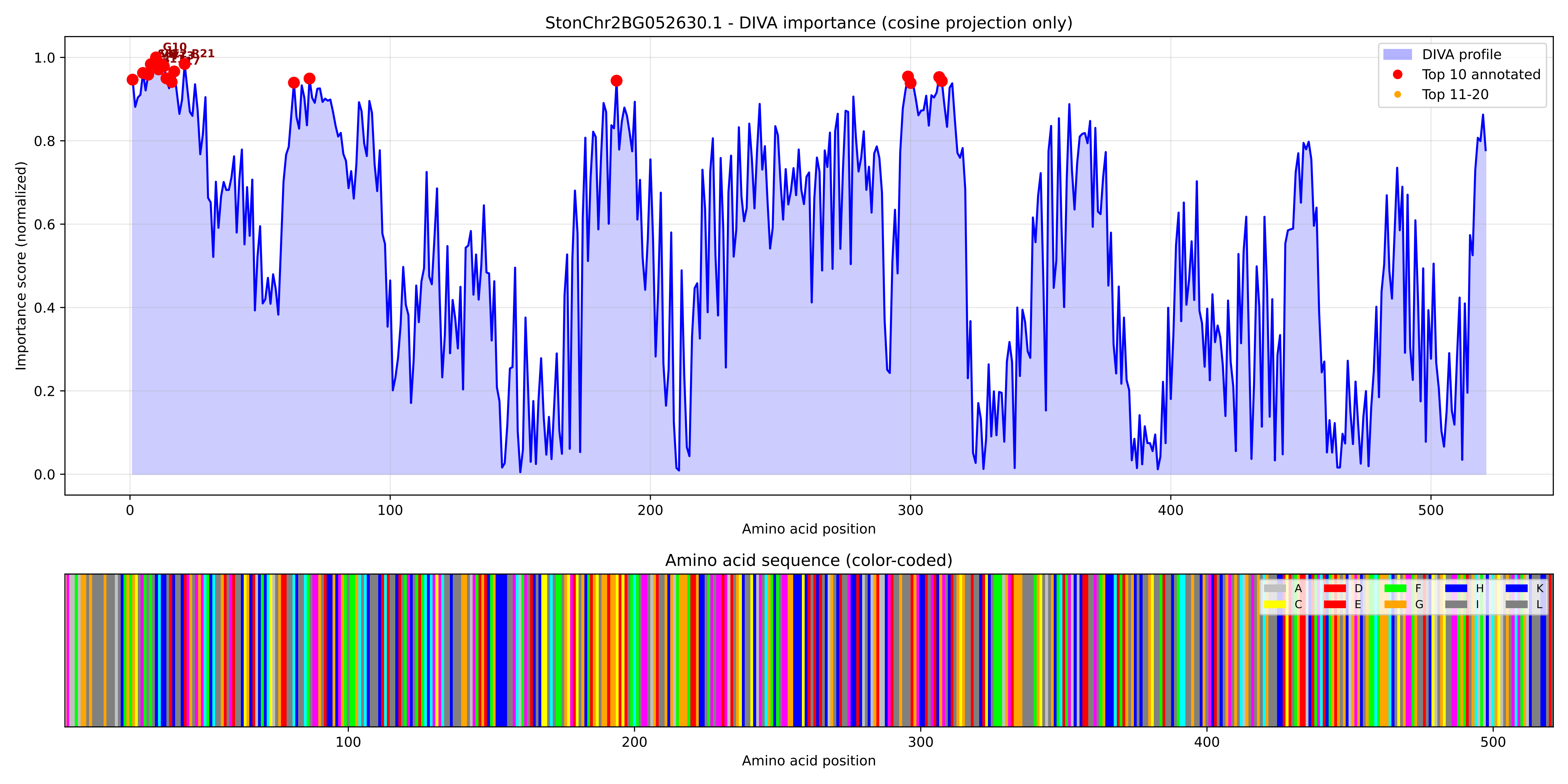

### StonChr2BG052630.1_importance.png

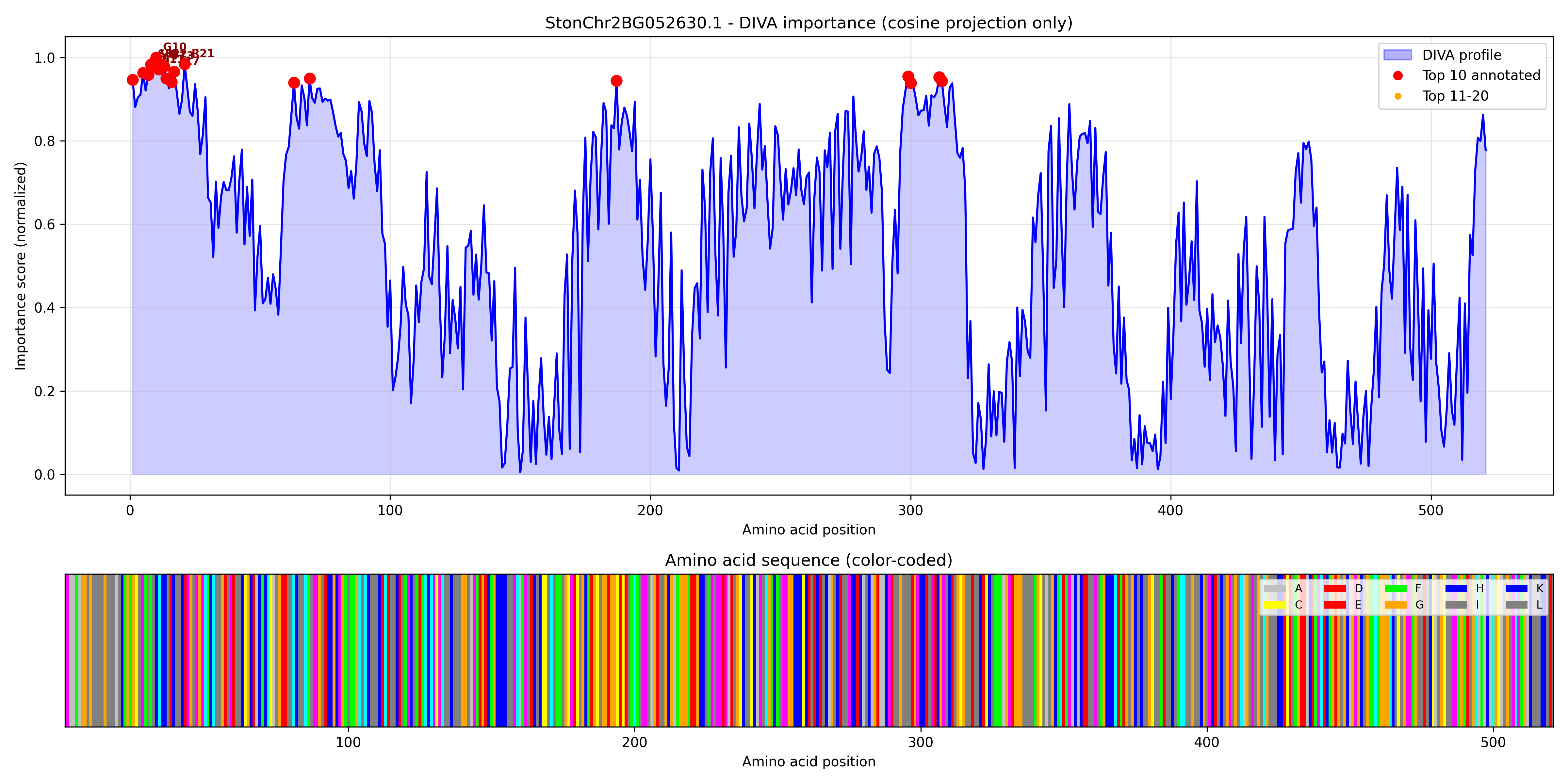

### StonChr2BG053070.1_importance.pdf

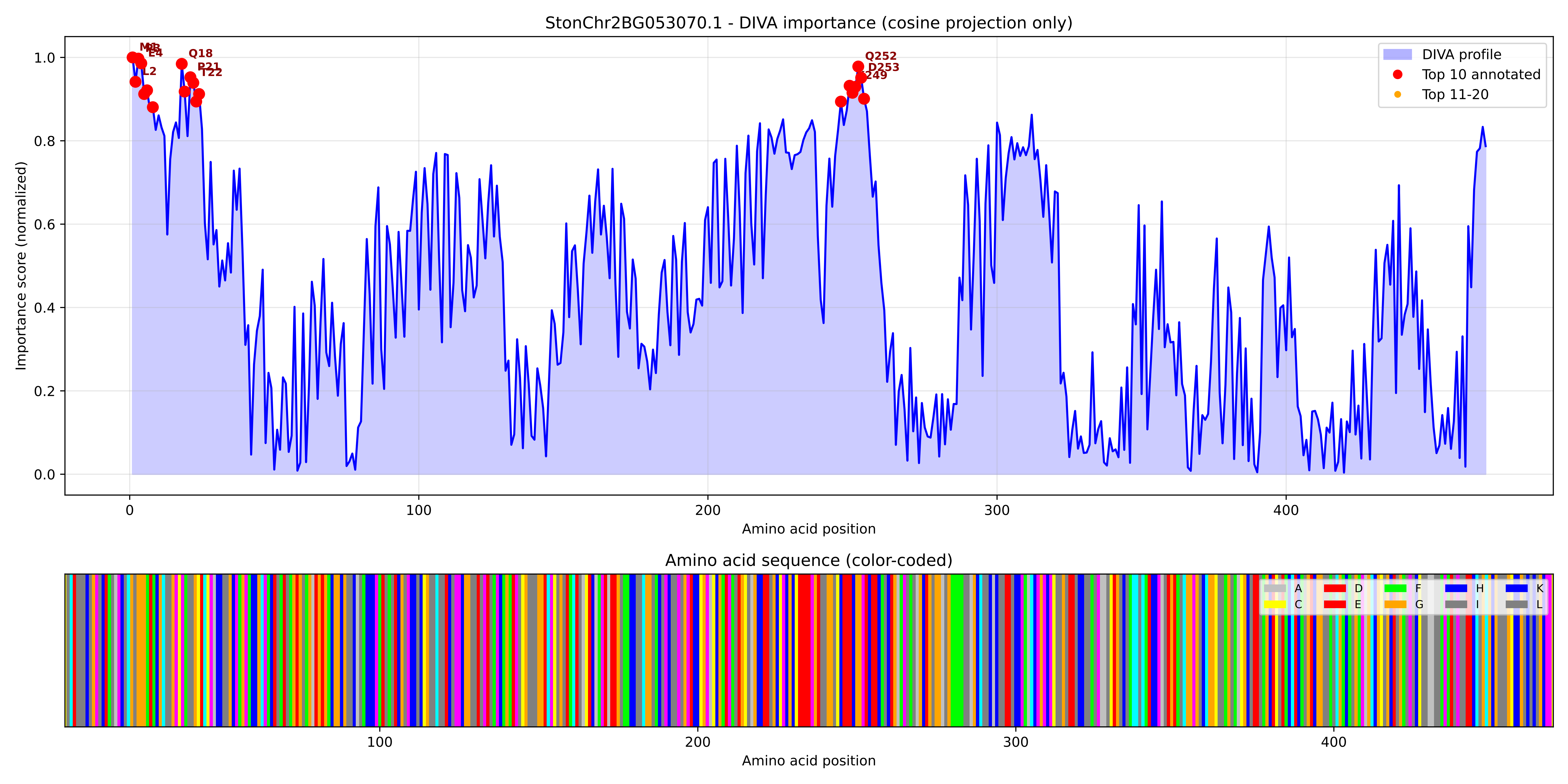

### StonChr2BG053070.1_importance.png

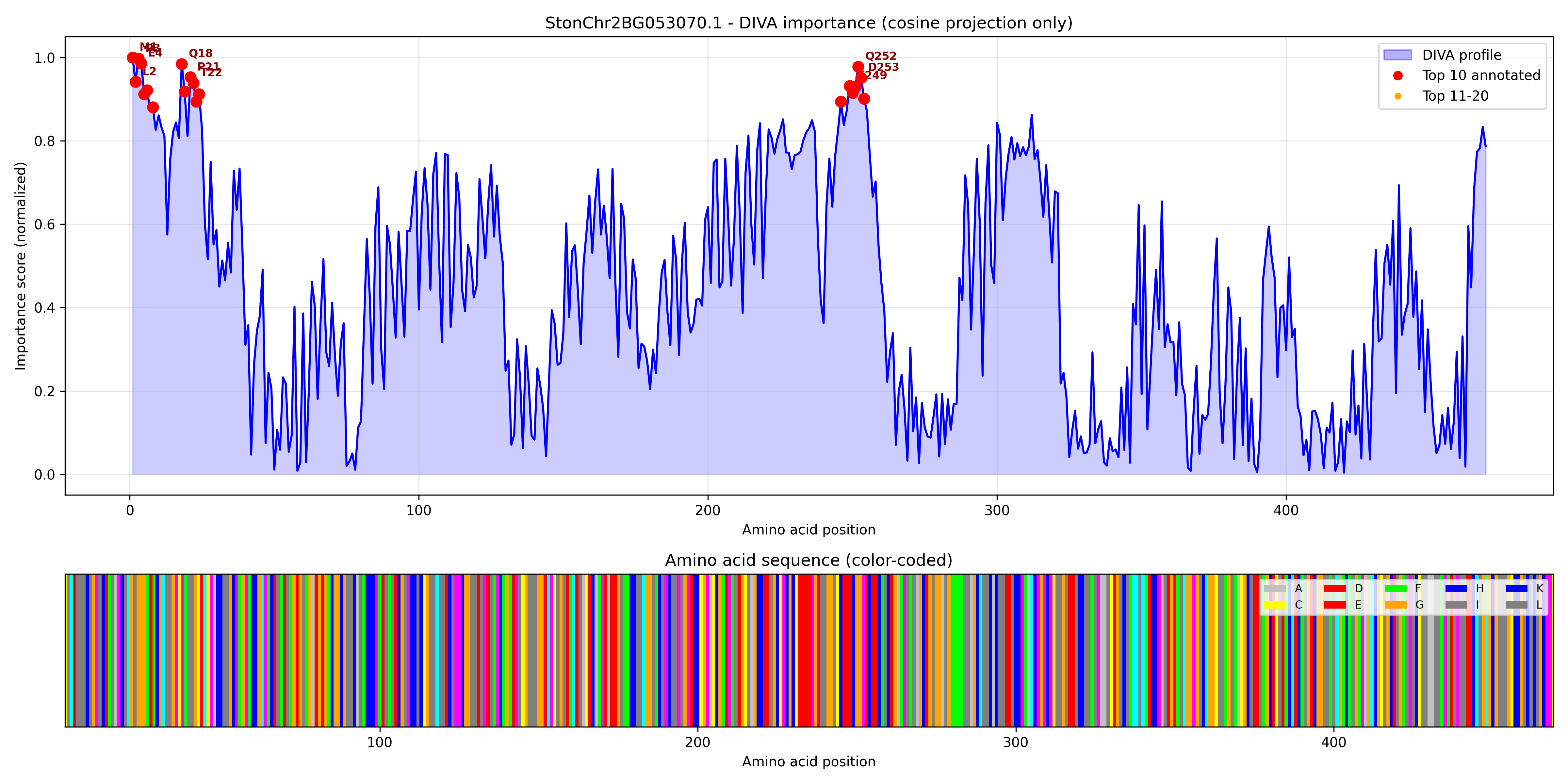

### StonChr2BG057580.1_importance.pdf

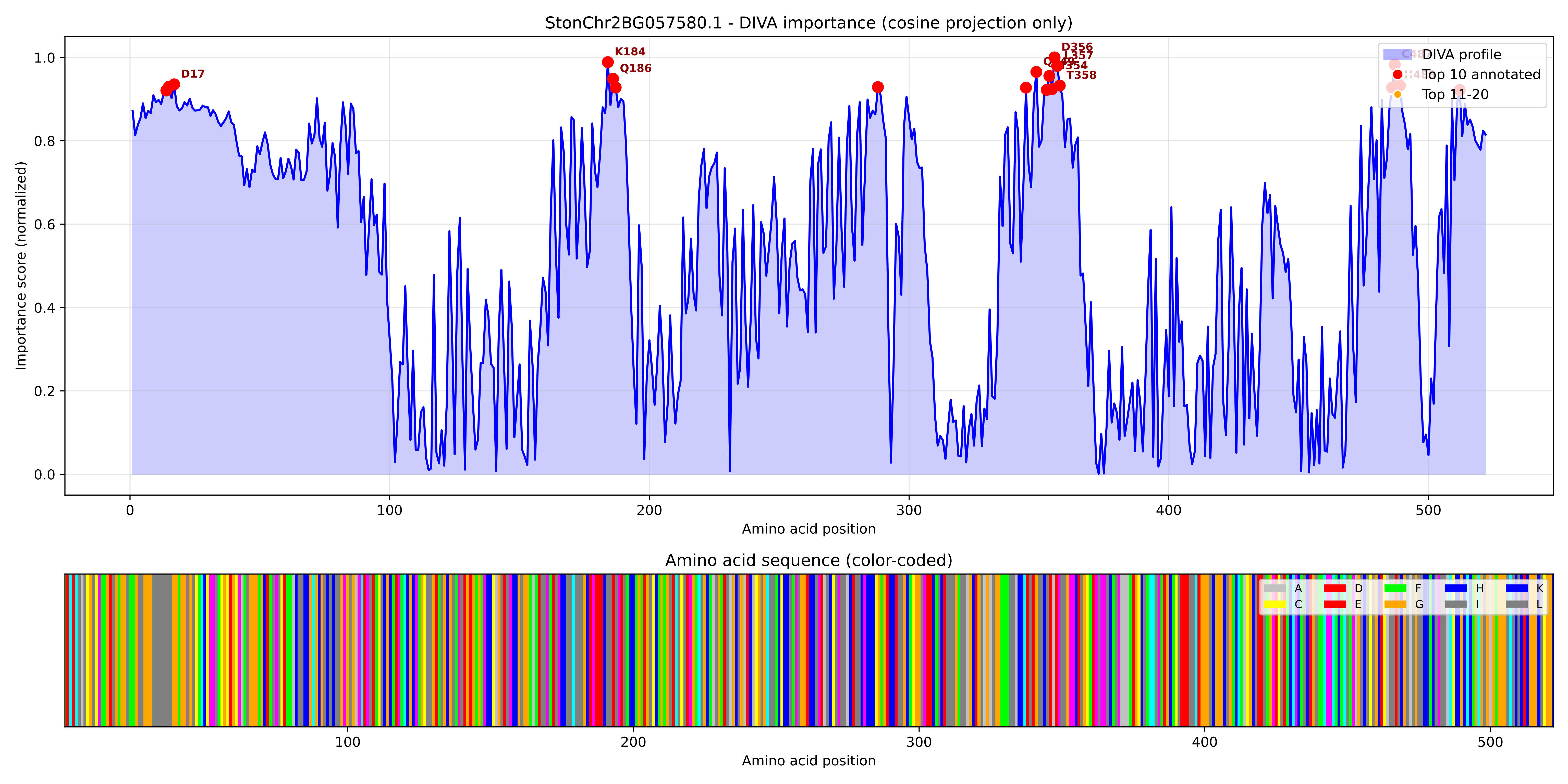

### StonChr2BG057580.1_importance.png

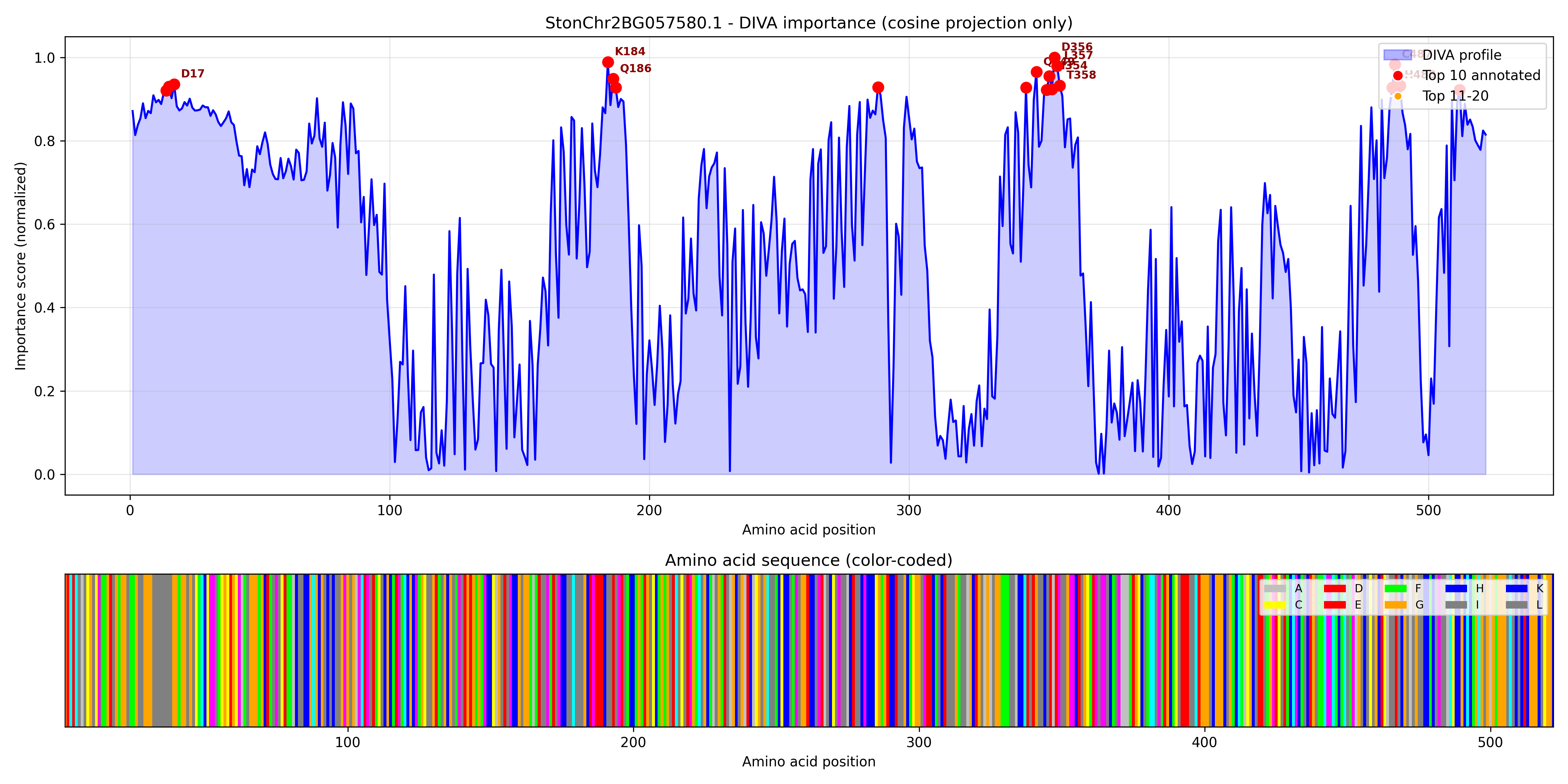

### StonChr2BG057970.1_importance.pdf

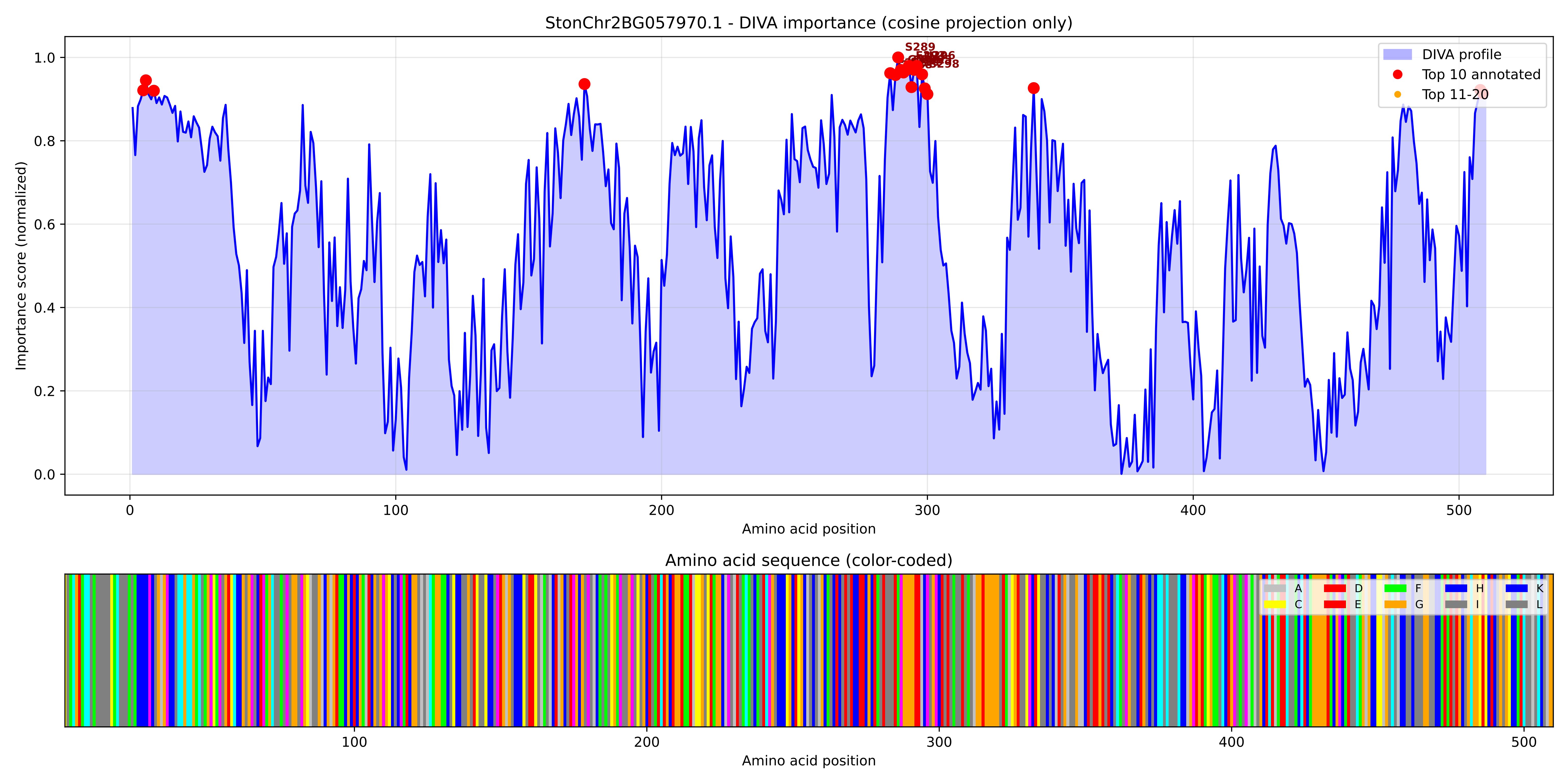

### StonChr2BG057970.1_importance.png

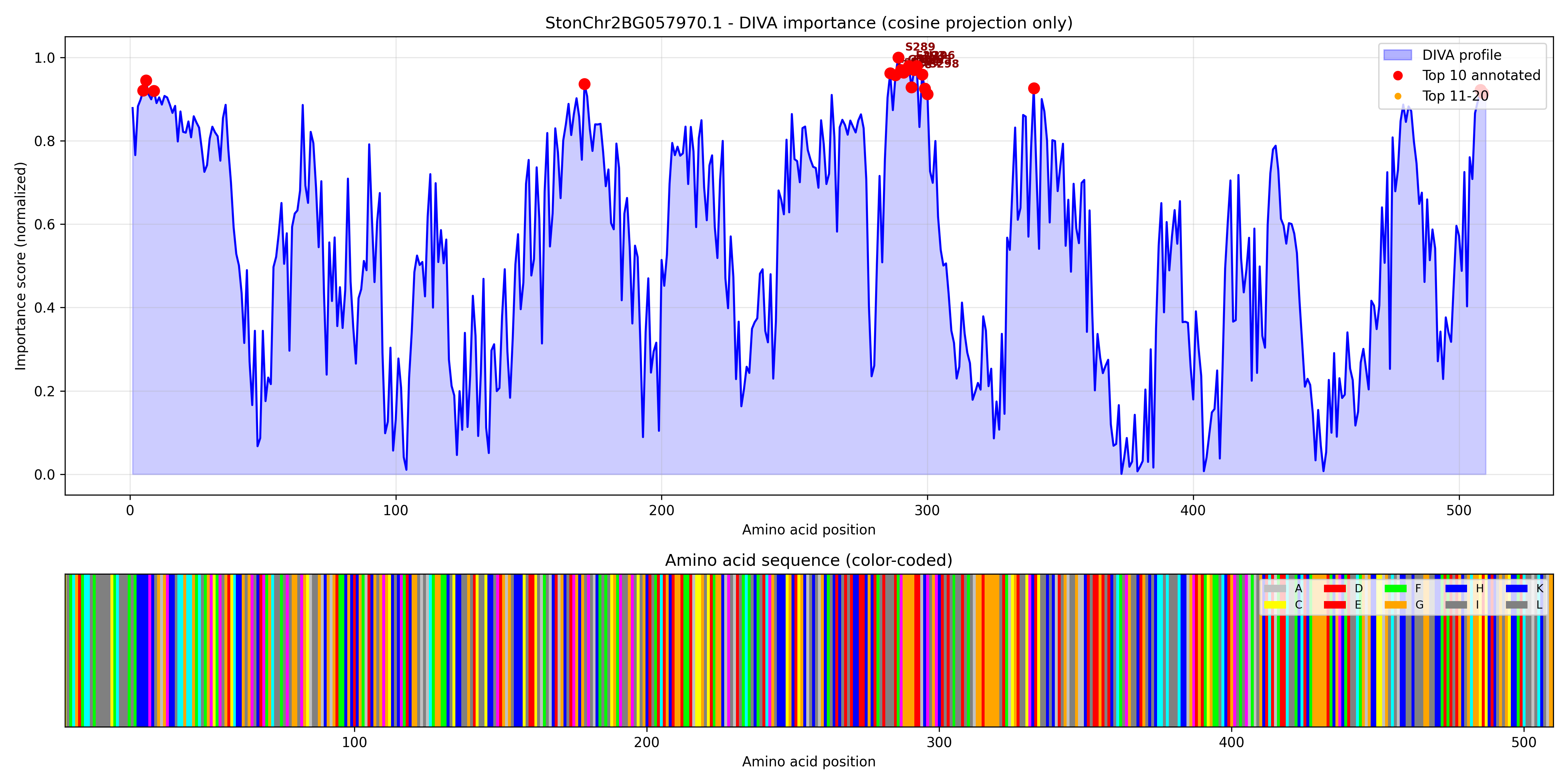

### StonChr2BG060840.1_importance.pdf

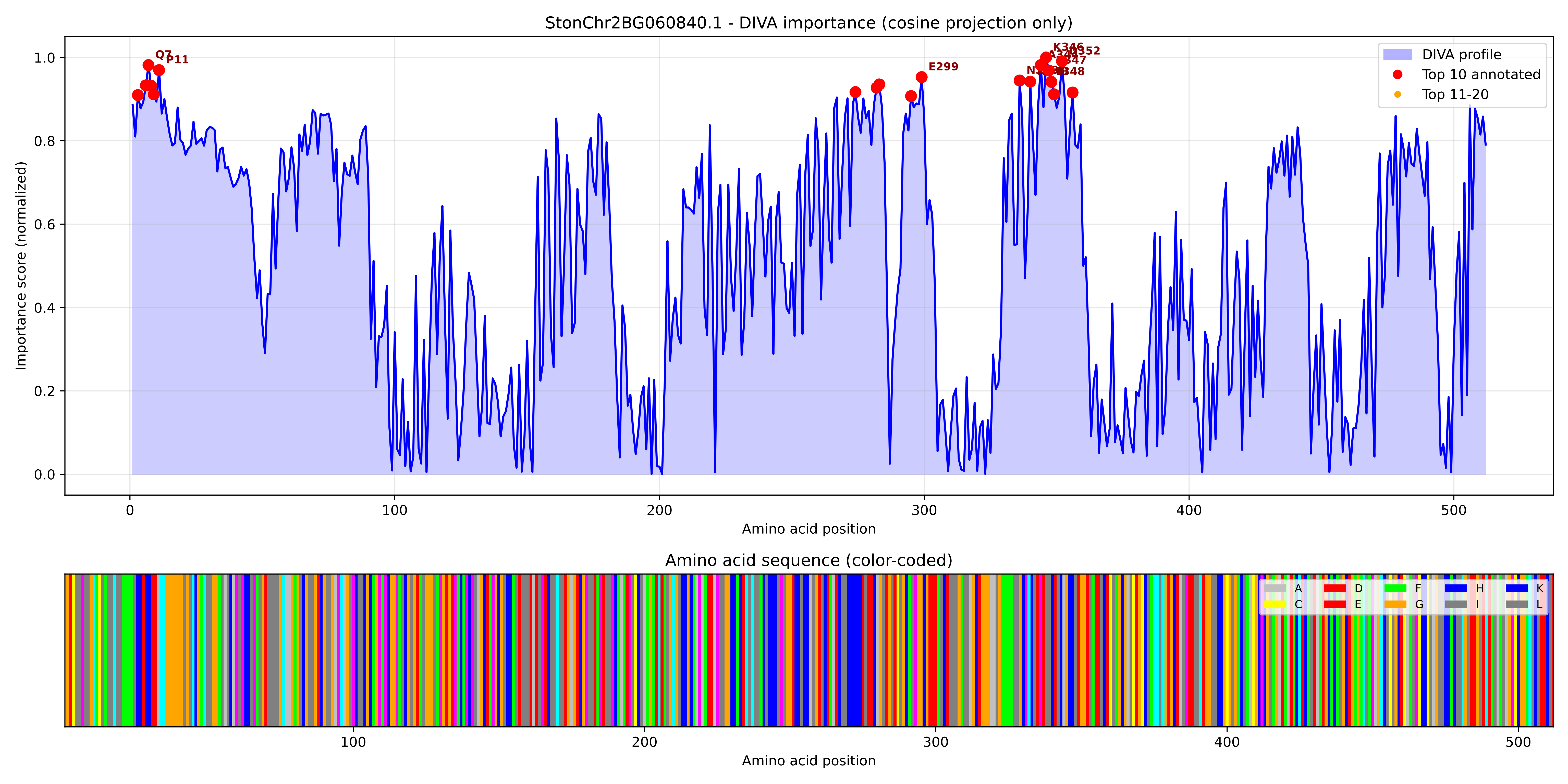

### StonChr2BG060840.1_importance.png

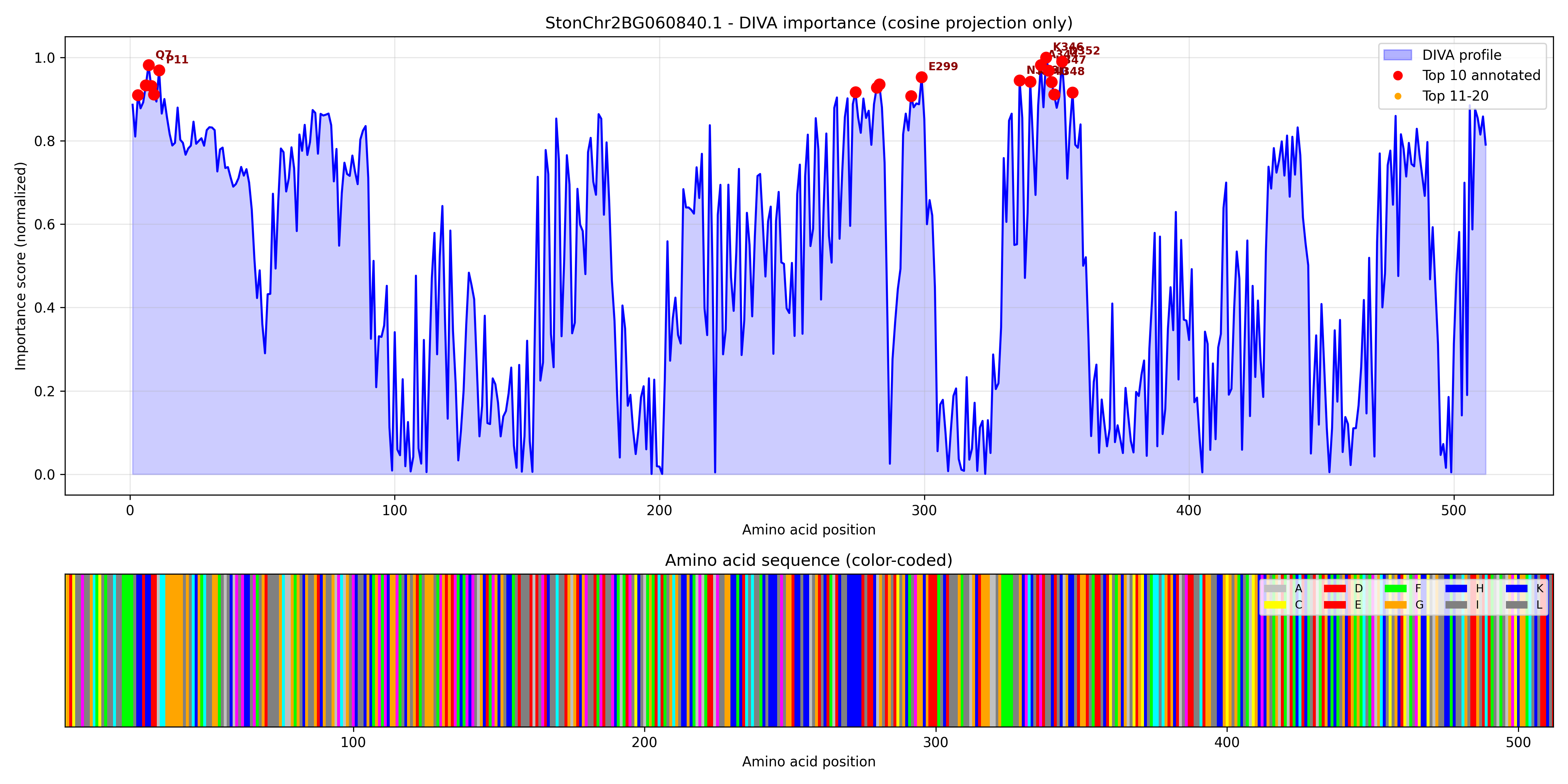

### StonChr2BG060850.1_importance.pdf

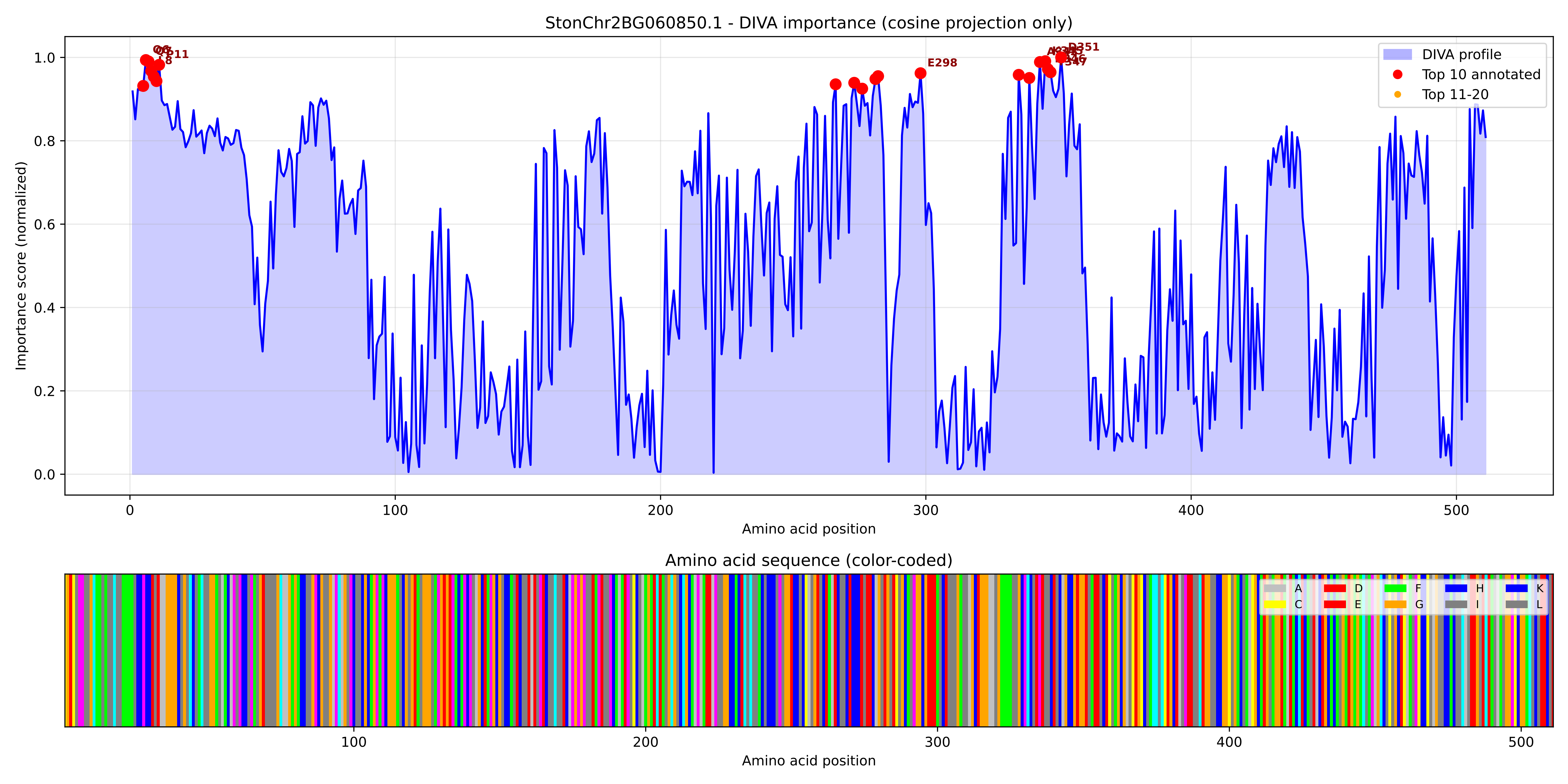

### StonChr2BG060850.1_importance.png

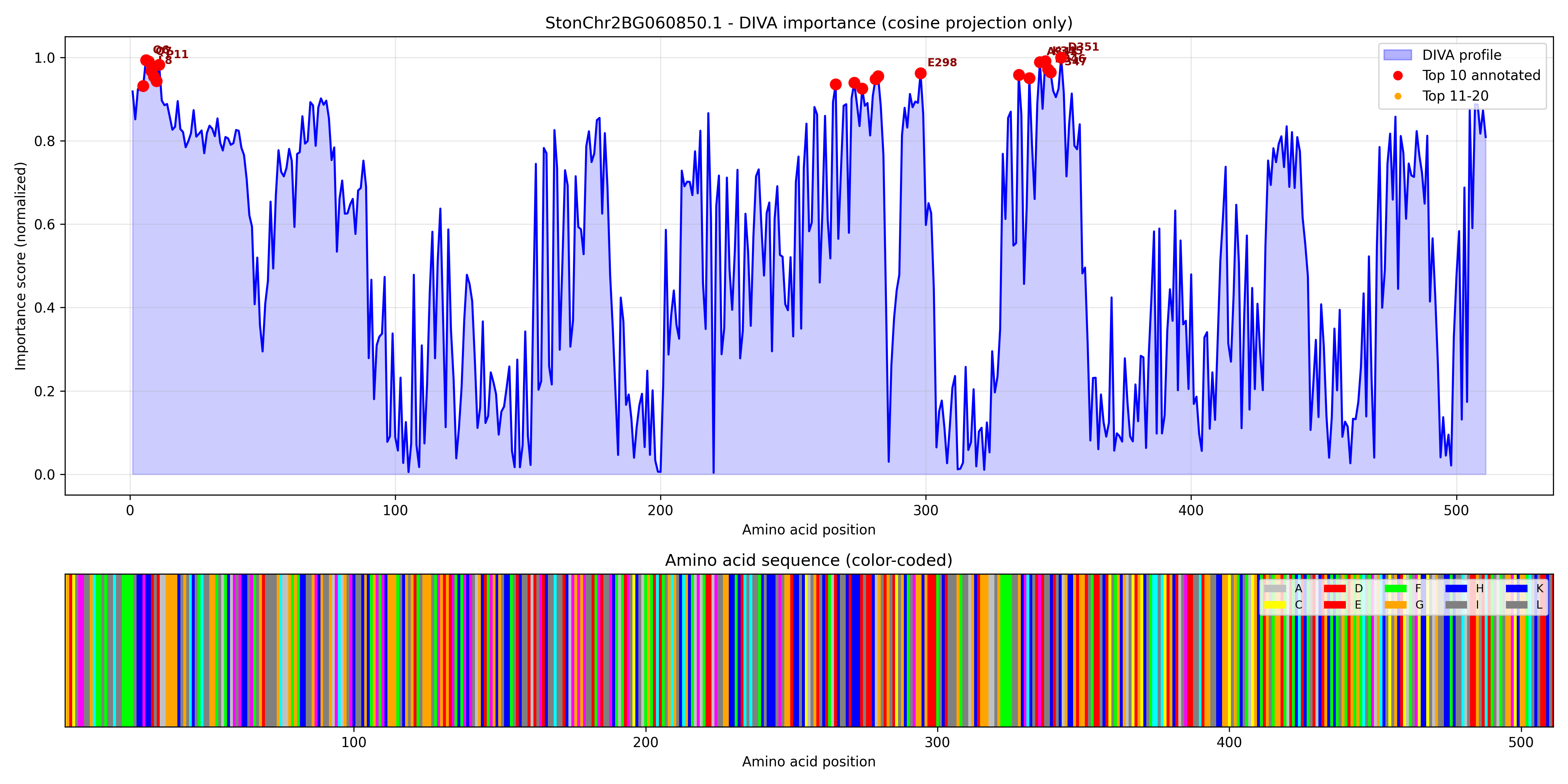

### StonChr3AG060130.1_importance.pdf

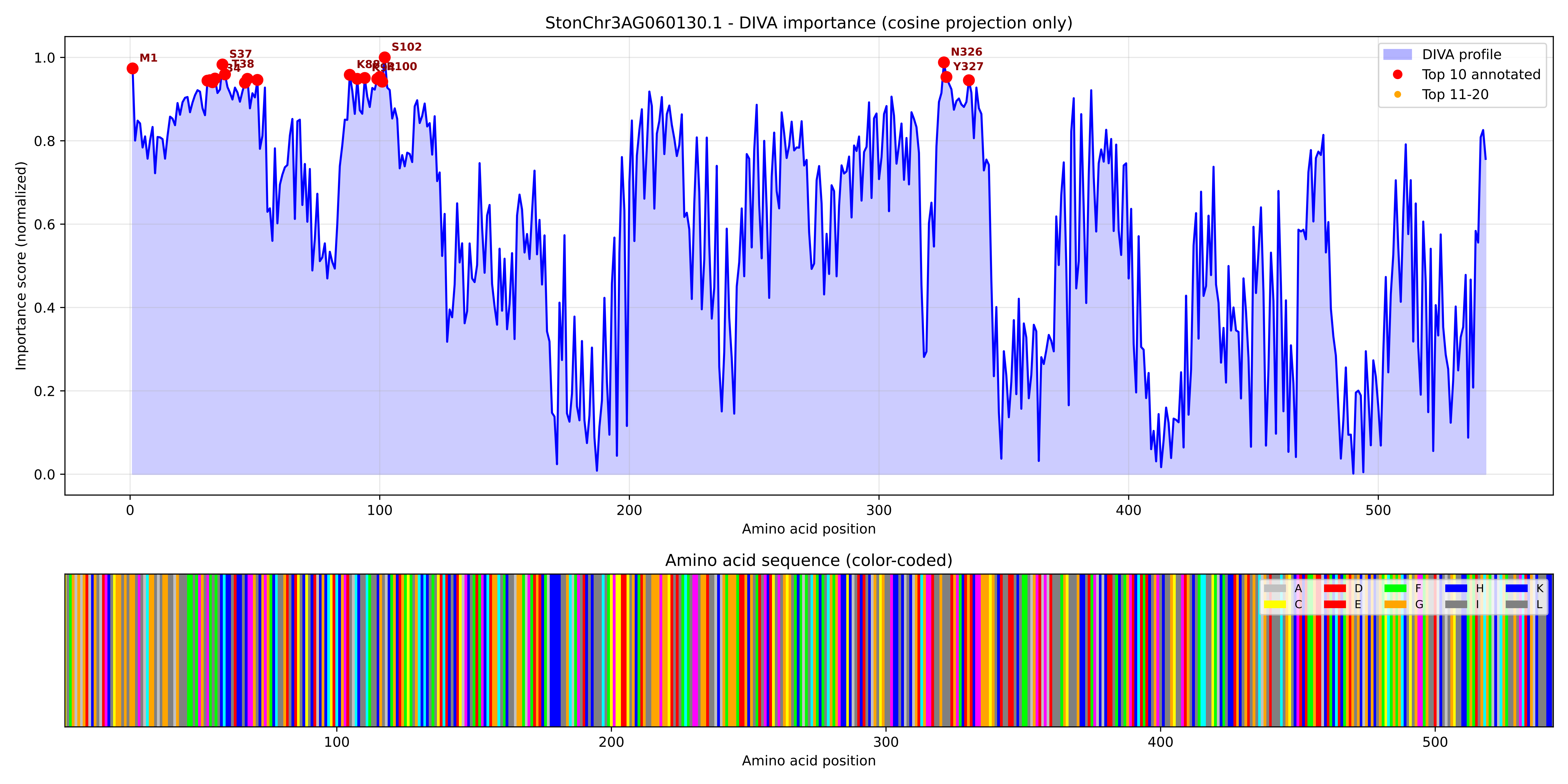

### StonChr3AG060130.1_importance.png

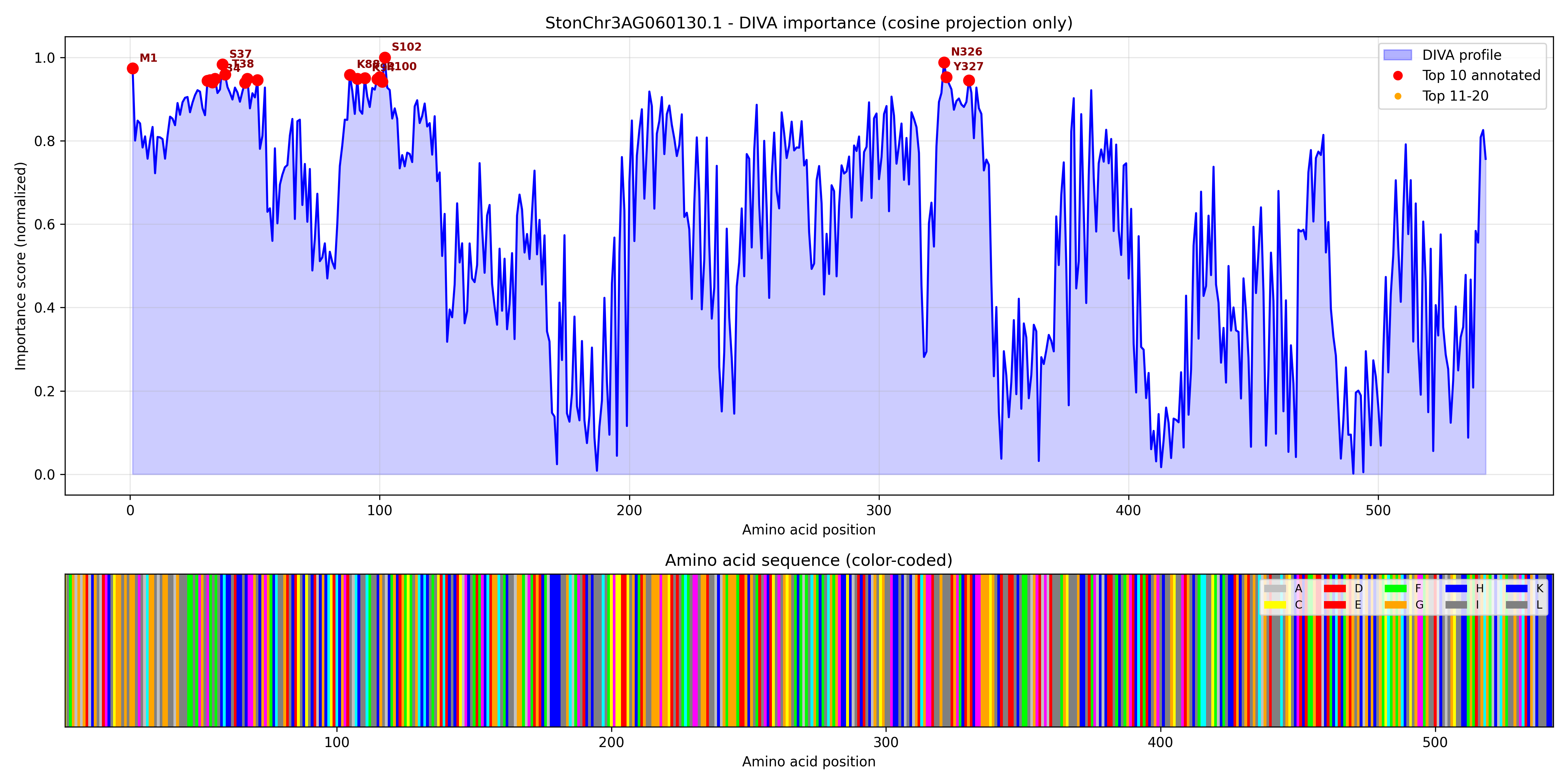

### StonChr3AG061670.1_importance.pdf

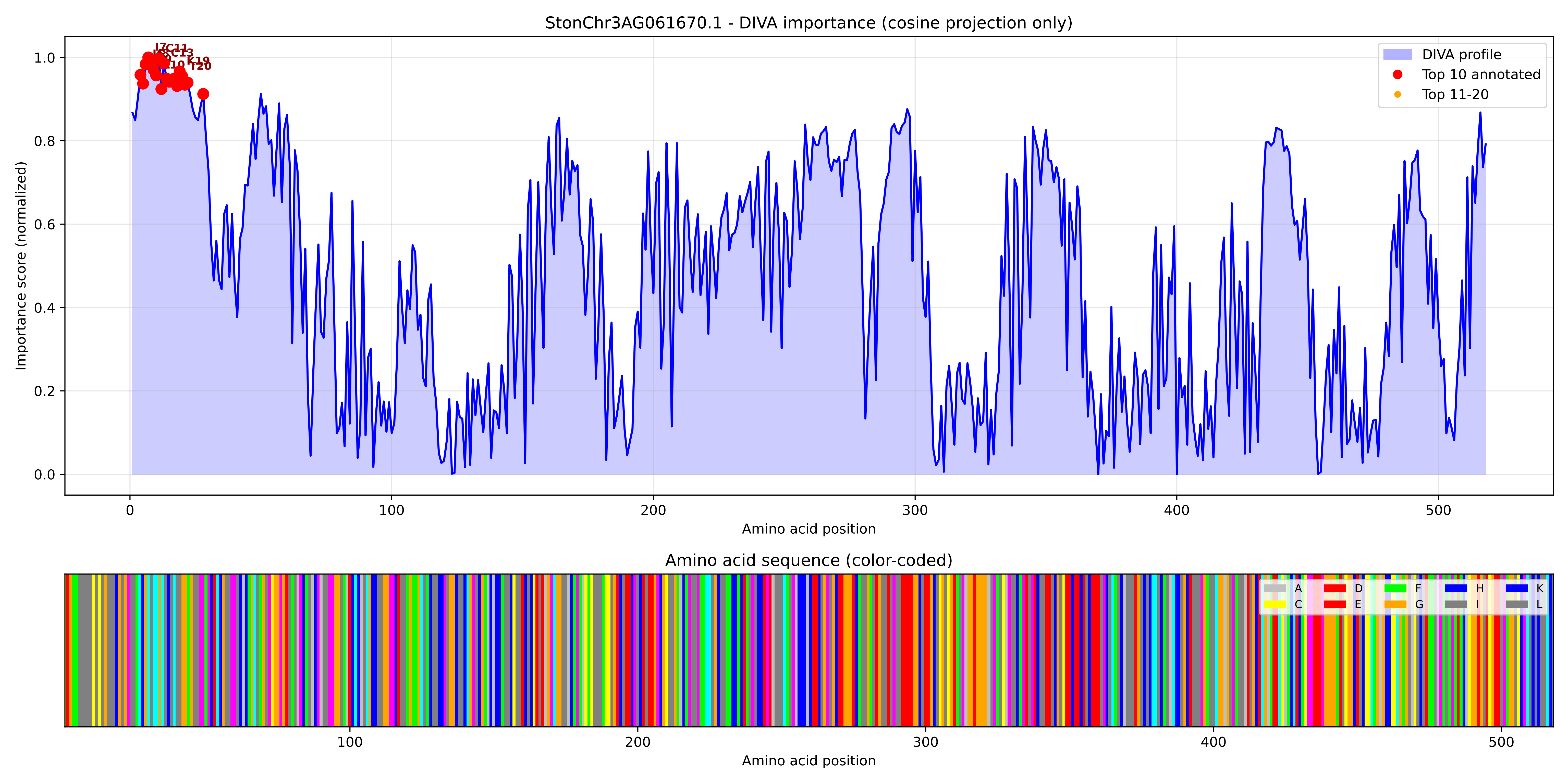

### StonChr3AG061670.1_importance.png

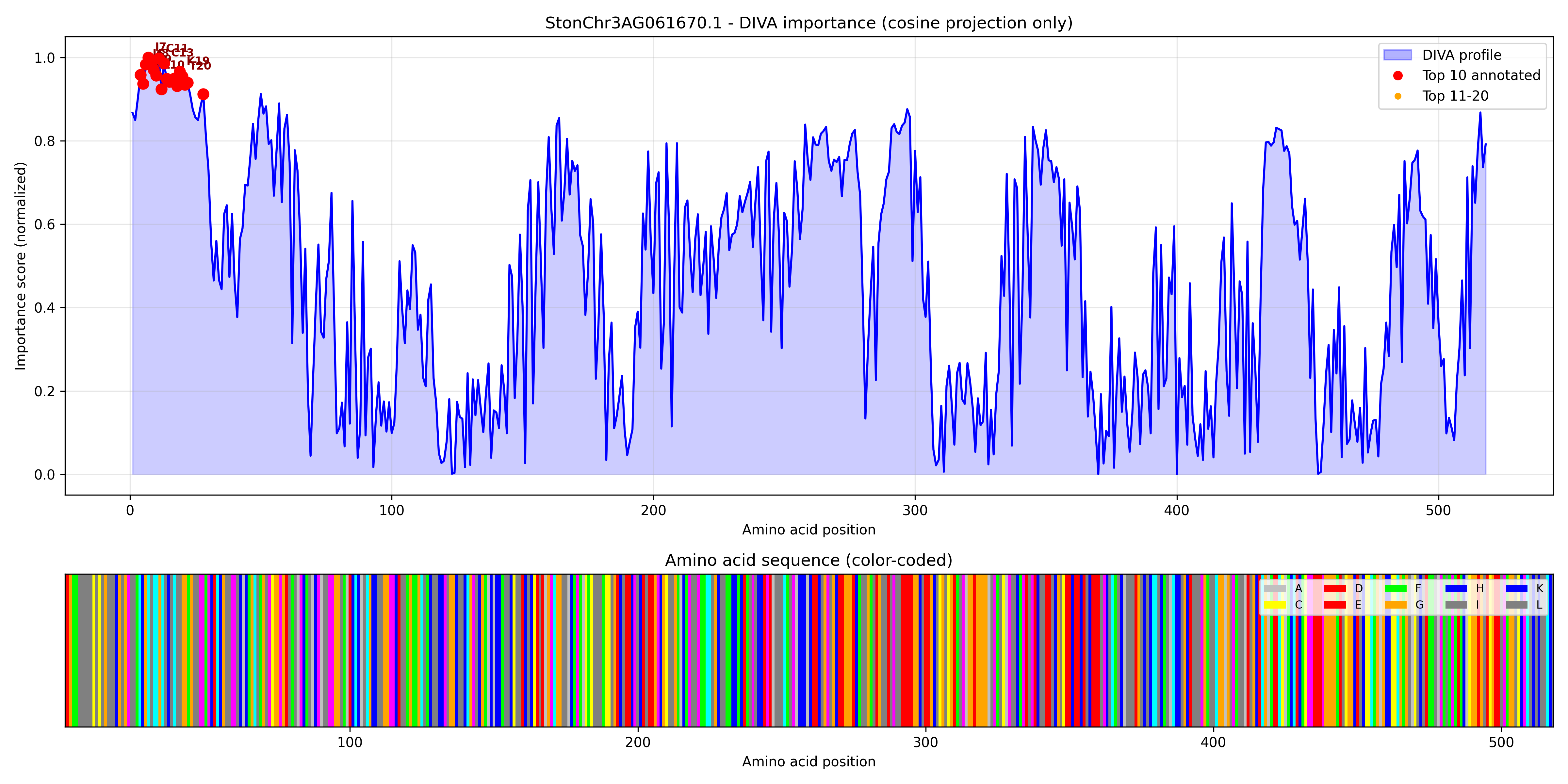

### StonChr3AG062200.1_importance.pdf

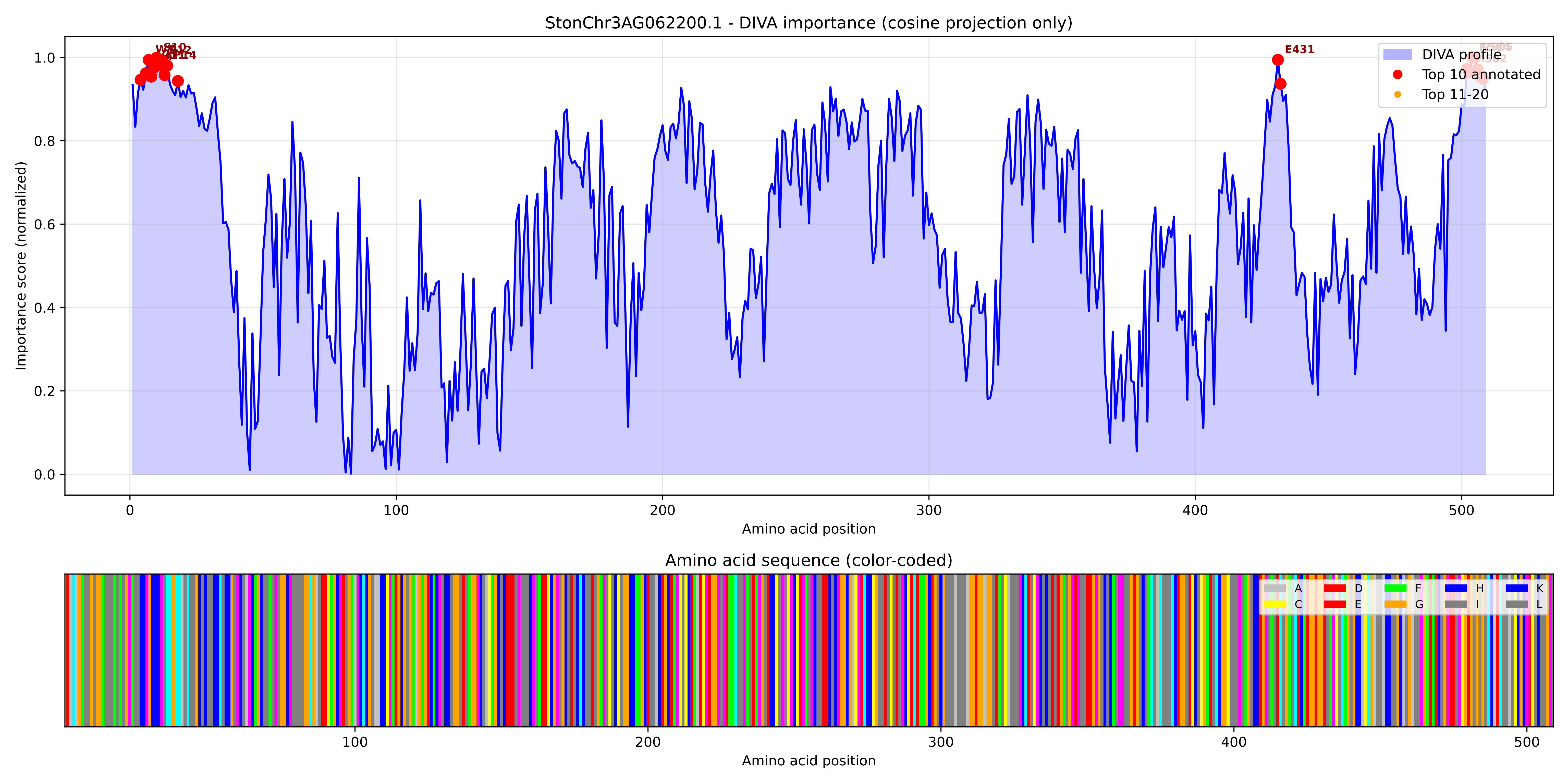

### StonChr3AG062200.1_importance.png

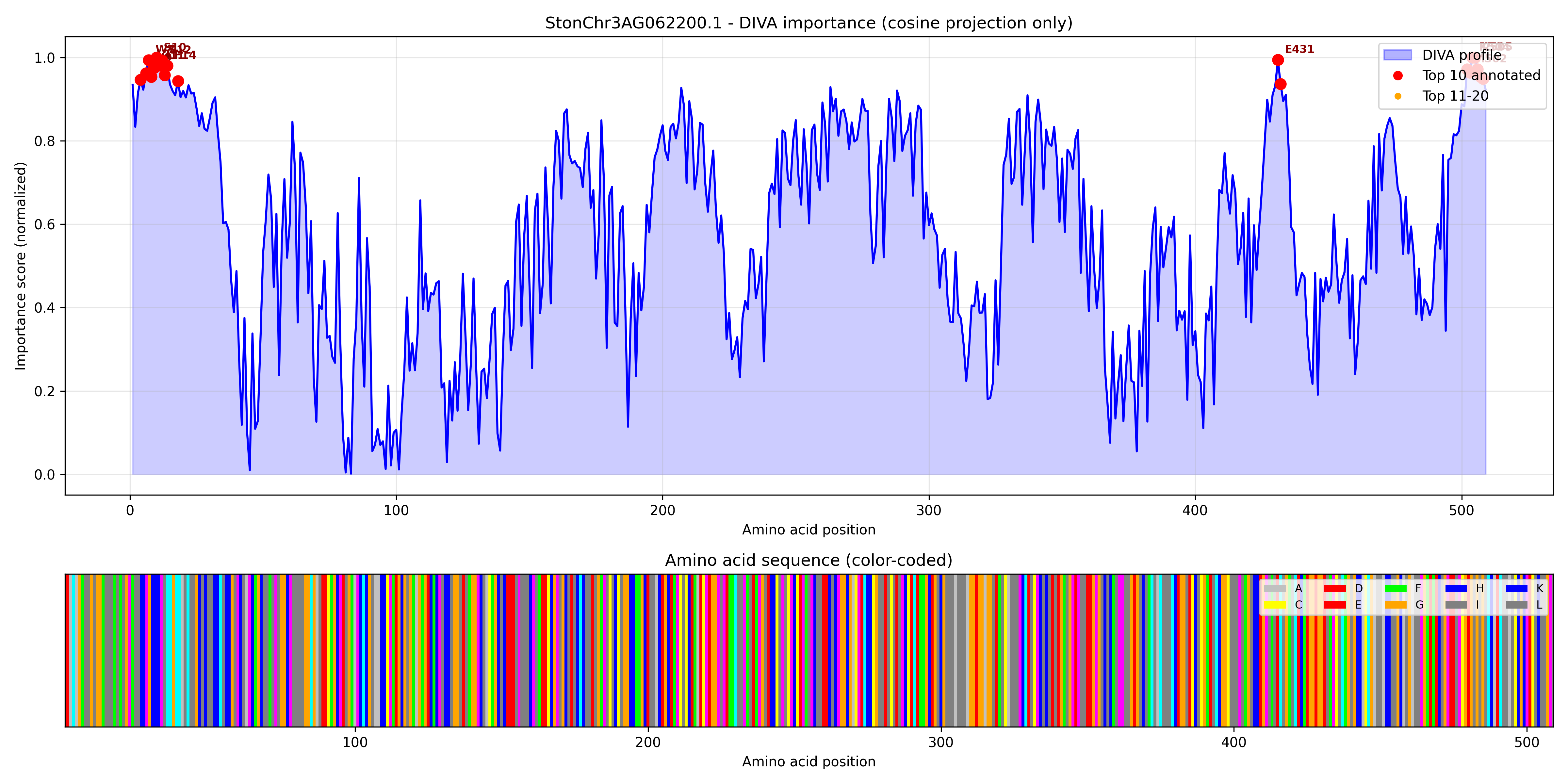

### StonChr3AG062220.1_importance.pdf

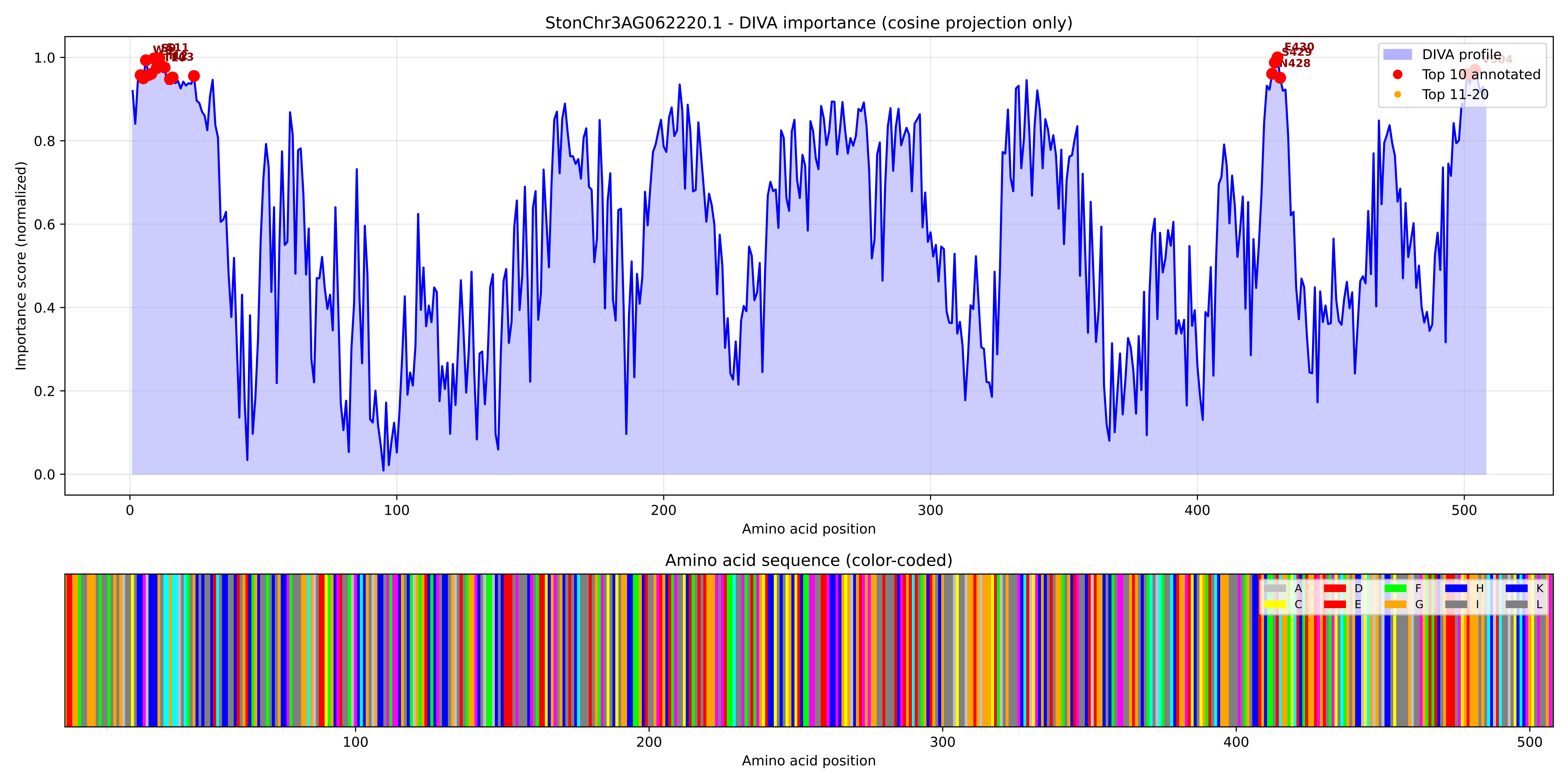

### StonChr3AG062220.1_importance.png

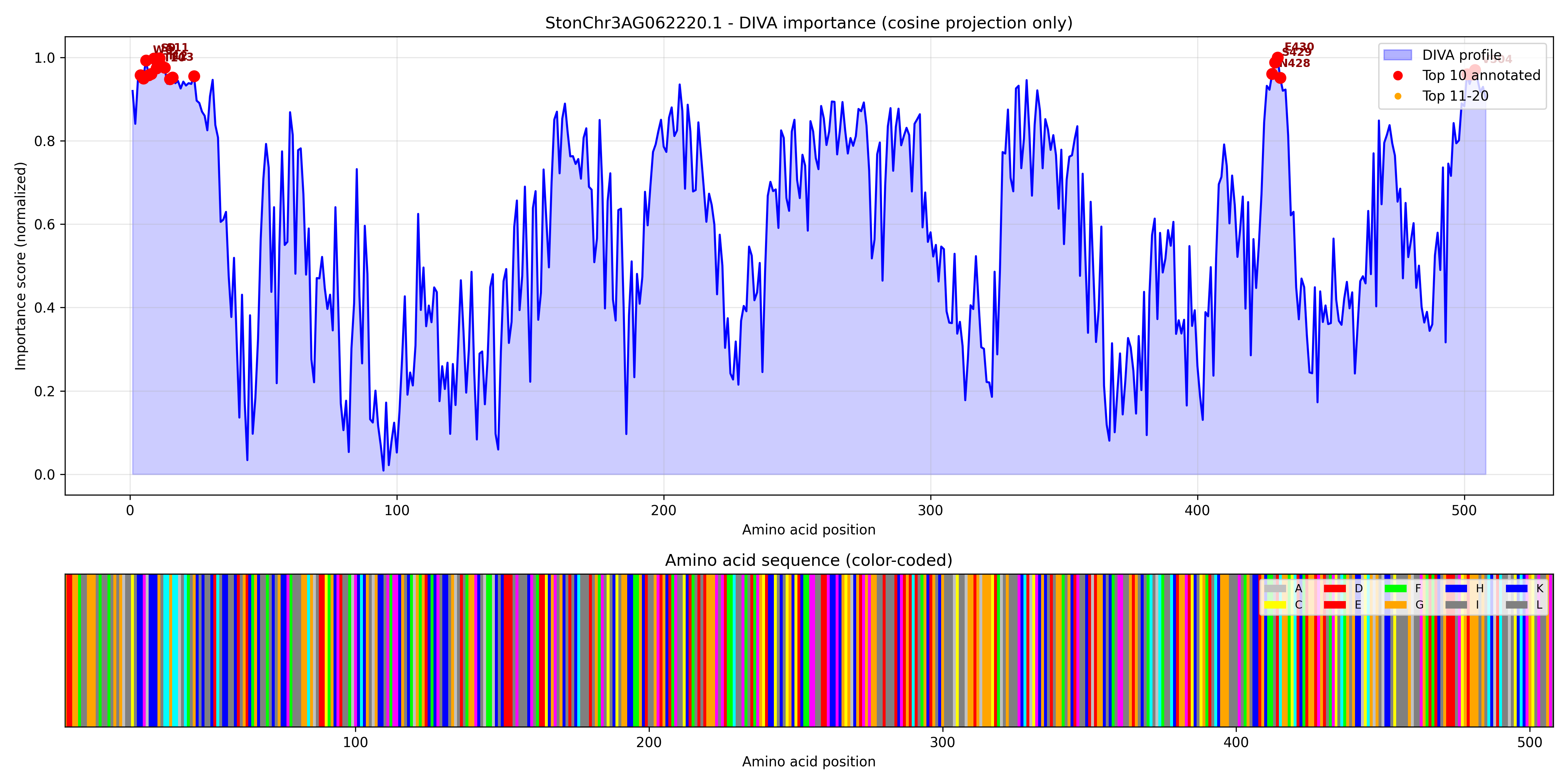

### StonChr3AG062980.1_importance.pdf

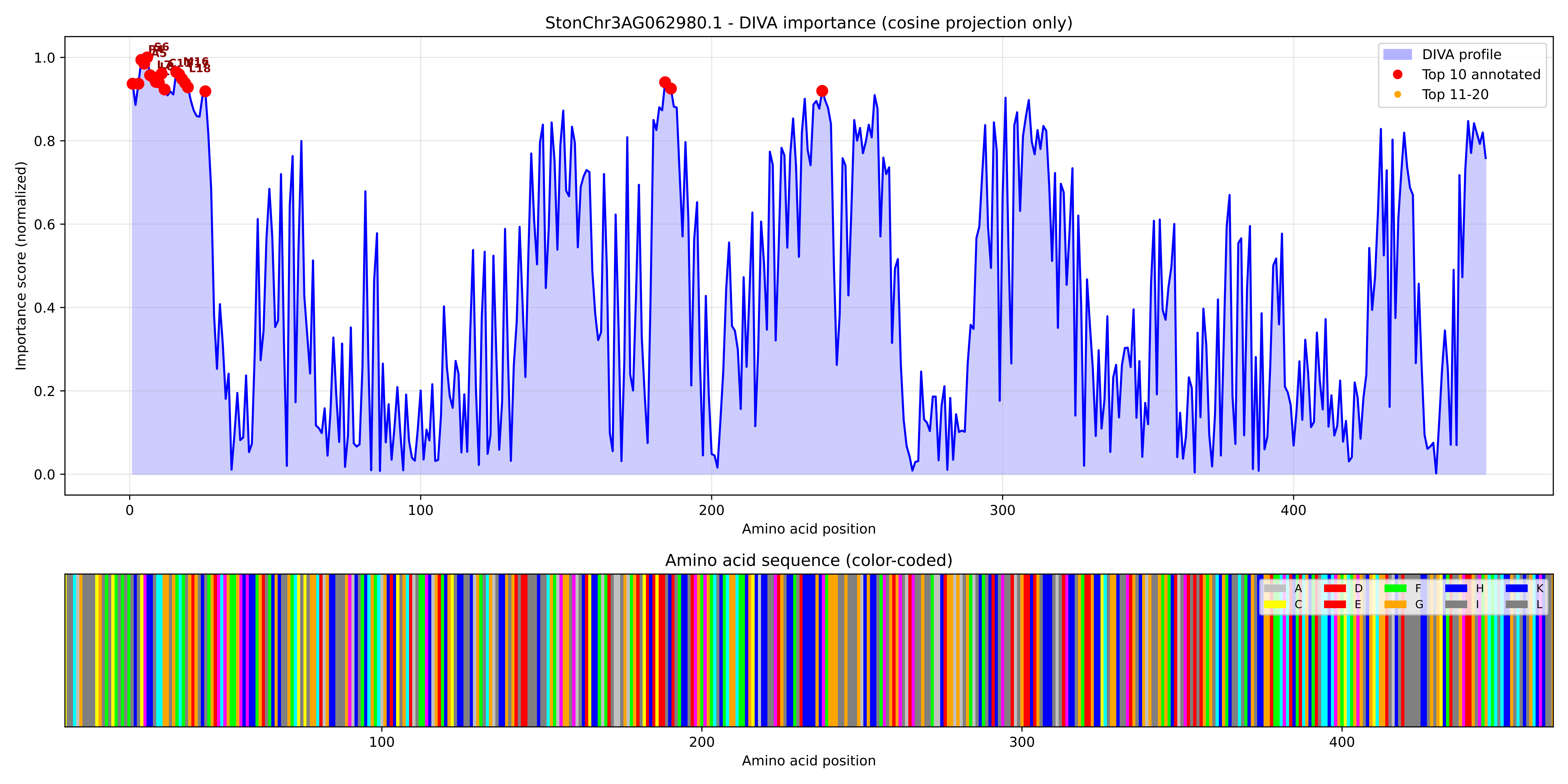

### StonChr3AG062980.1_importance.png

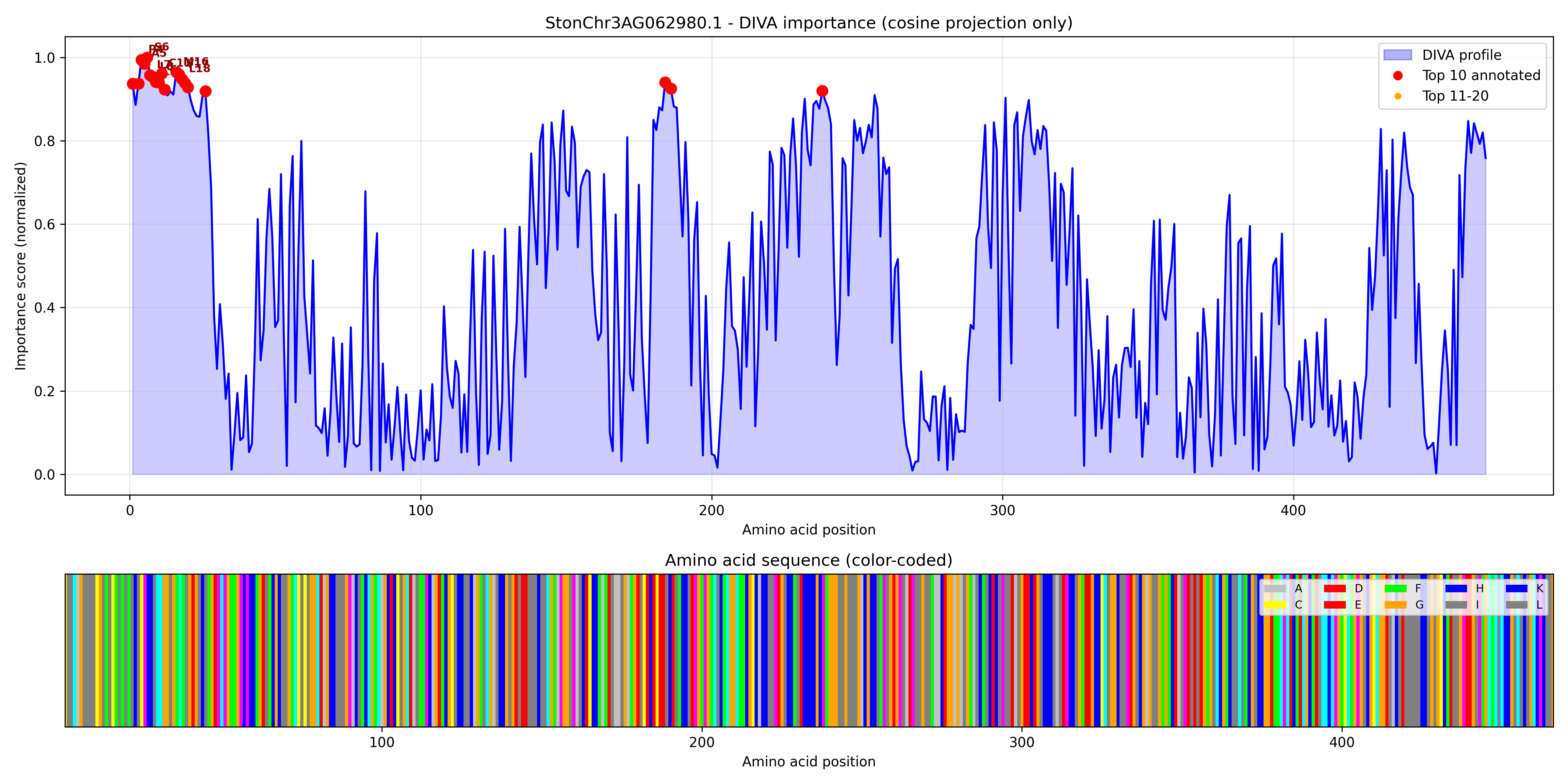

### StonChr3AG063010.1_importance.pdf

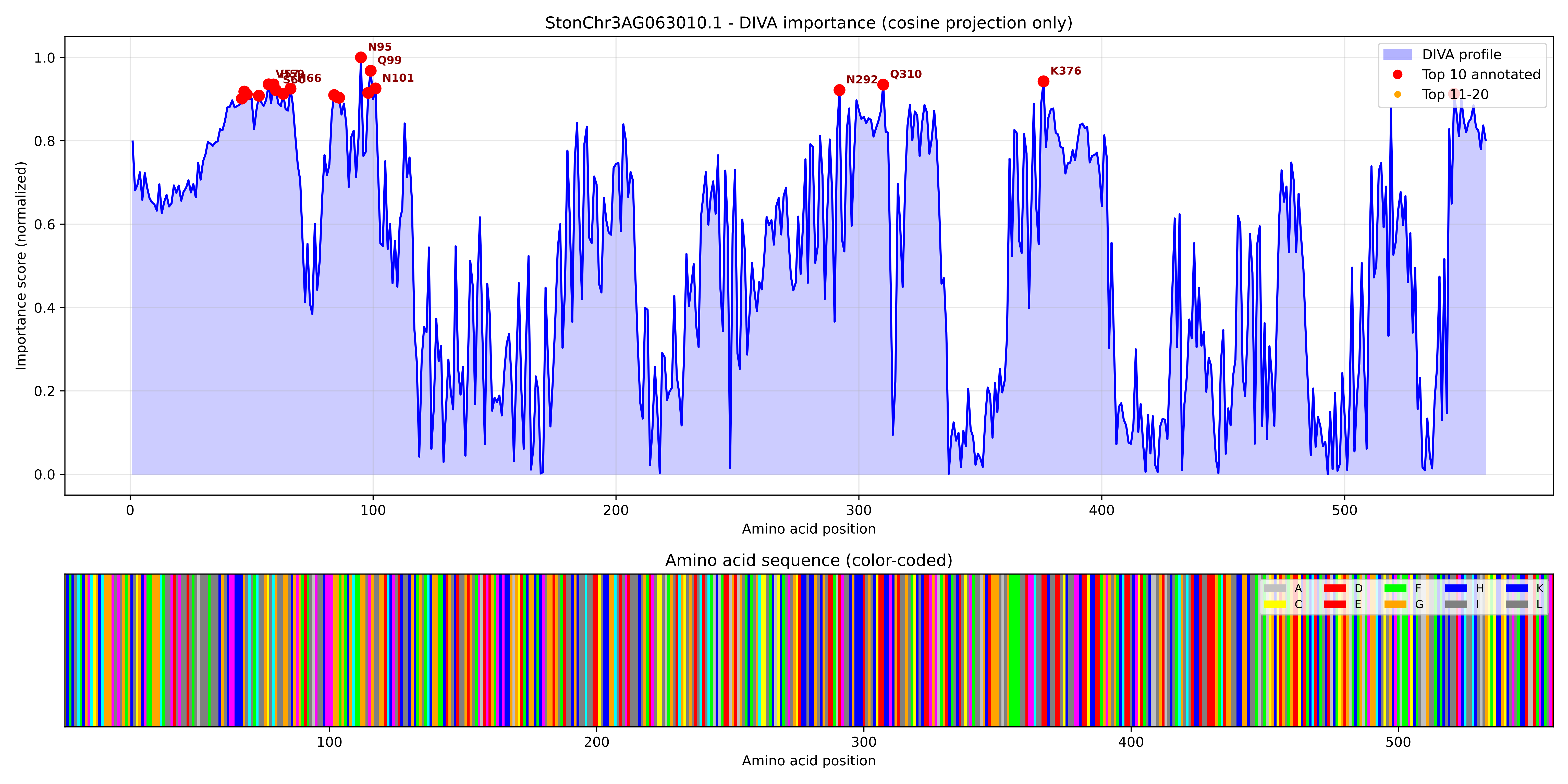

### StonChr3AG063010.1_importance.png

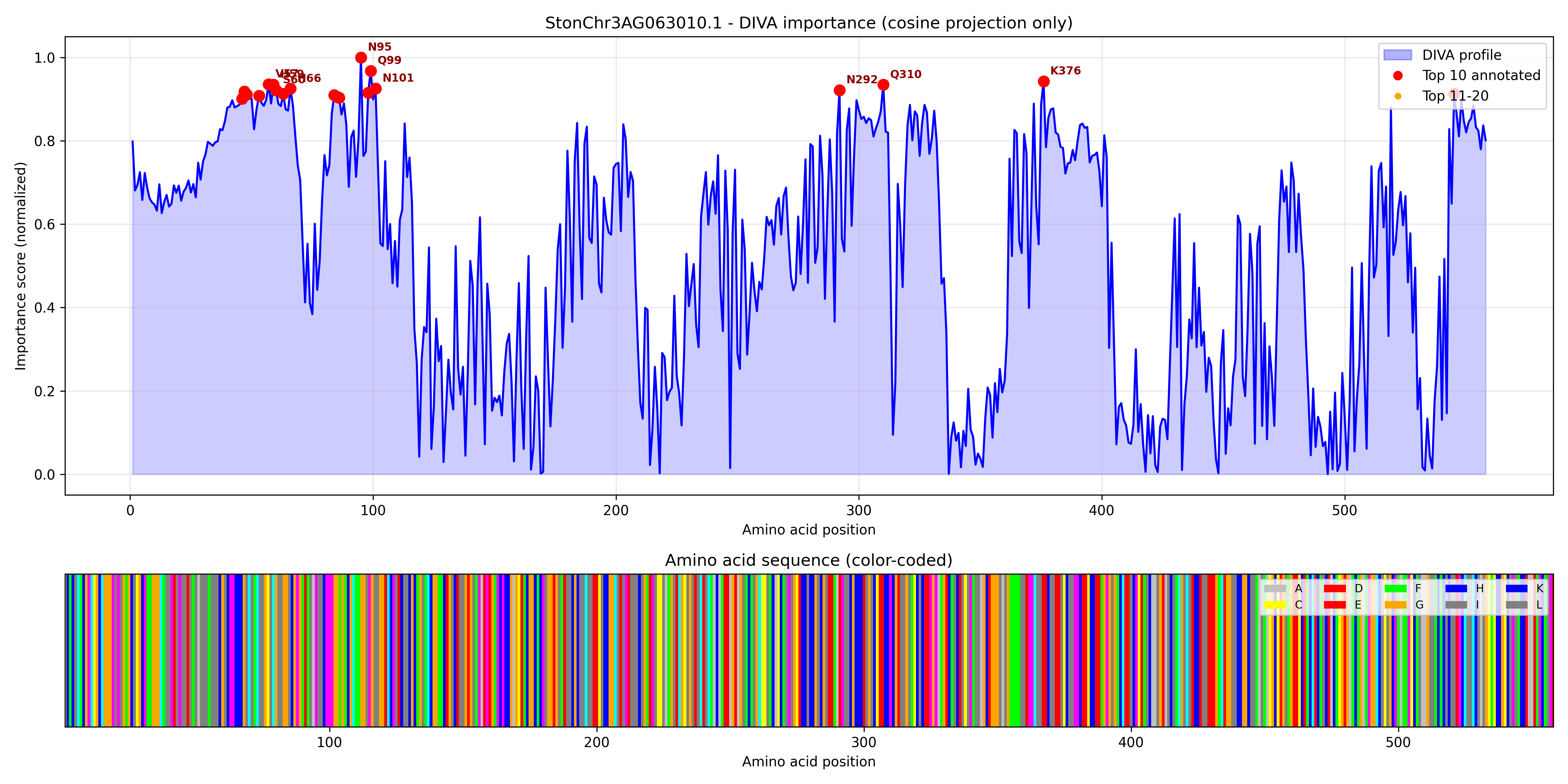

### StonChr3AG063460.1_importance.pdf

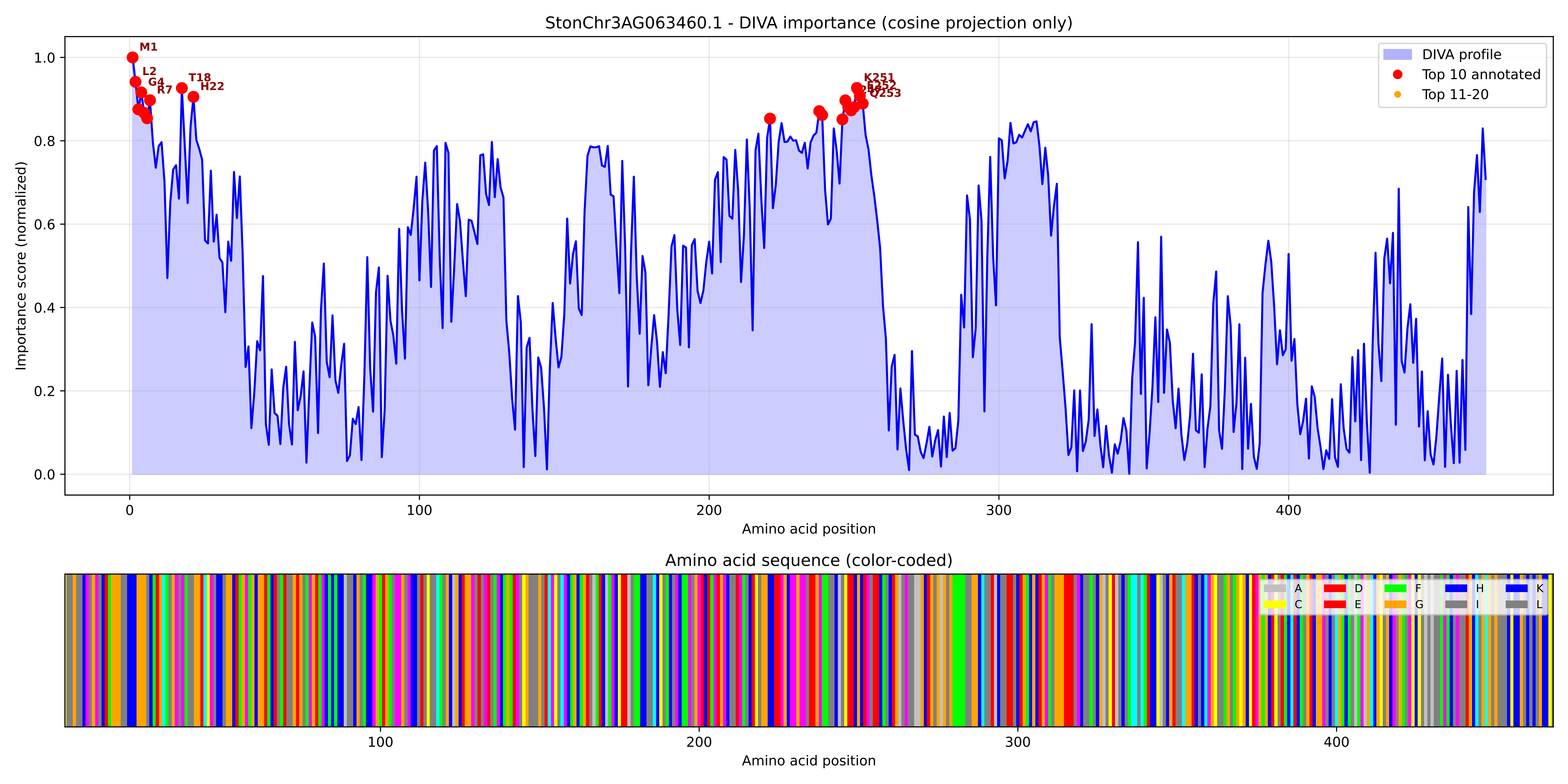

### StonChr3AG063460.1_importance.png

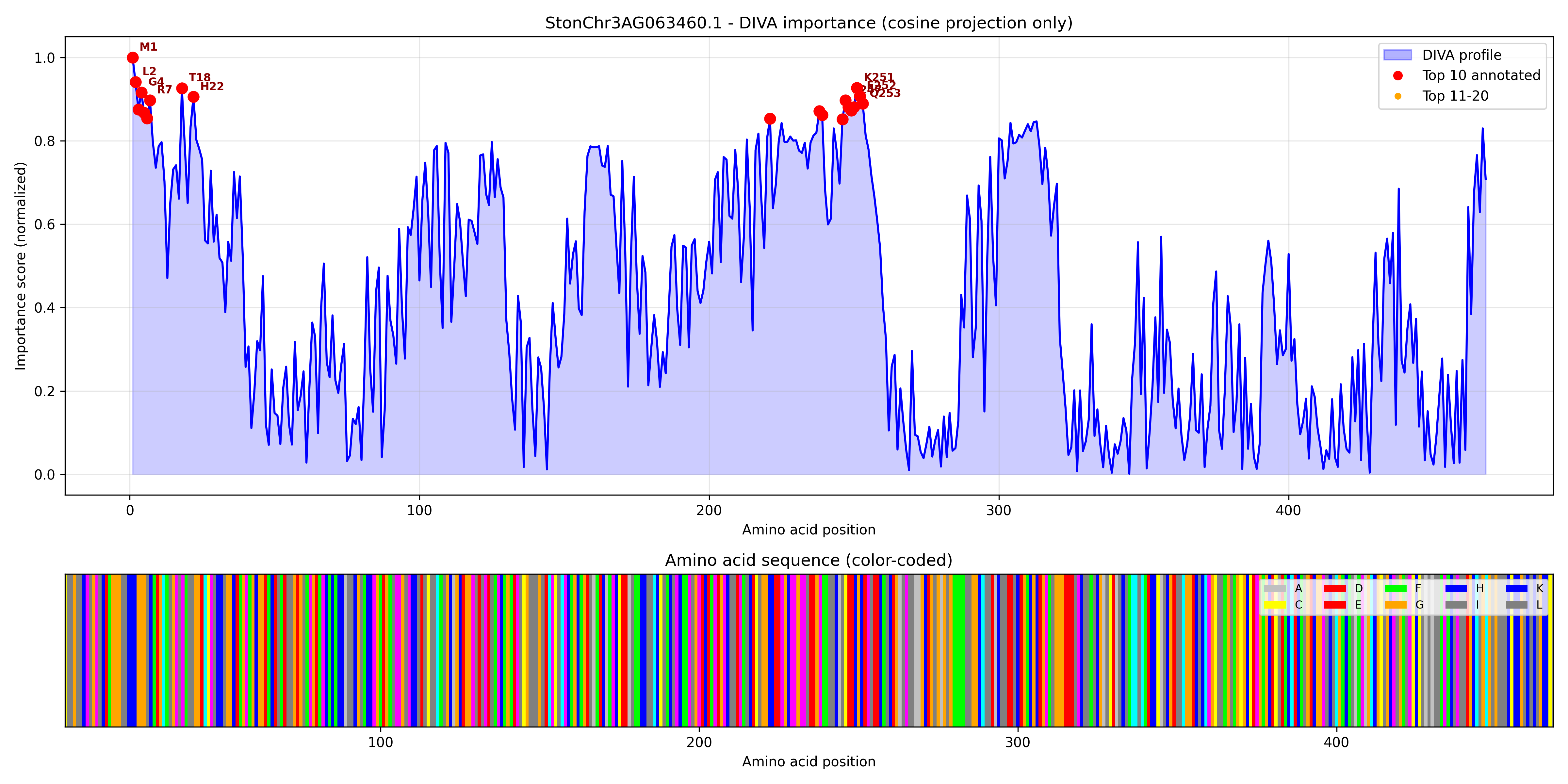

### StonChr3AG064210.1_importance.pdf

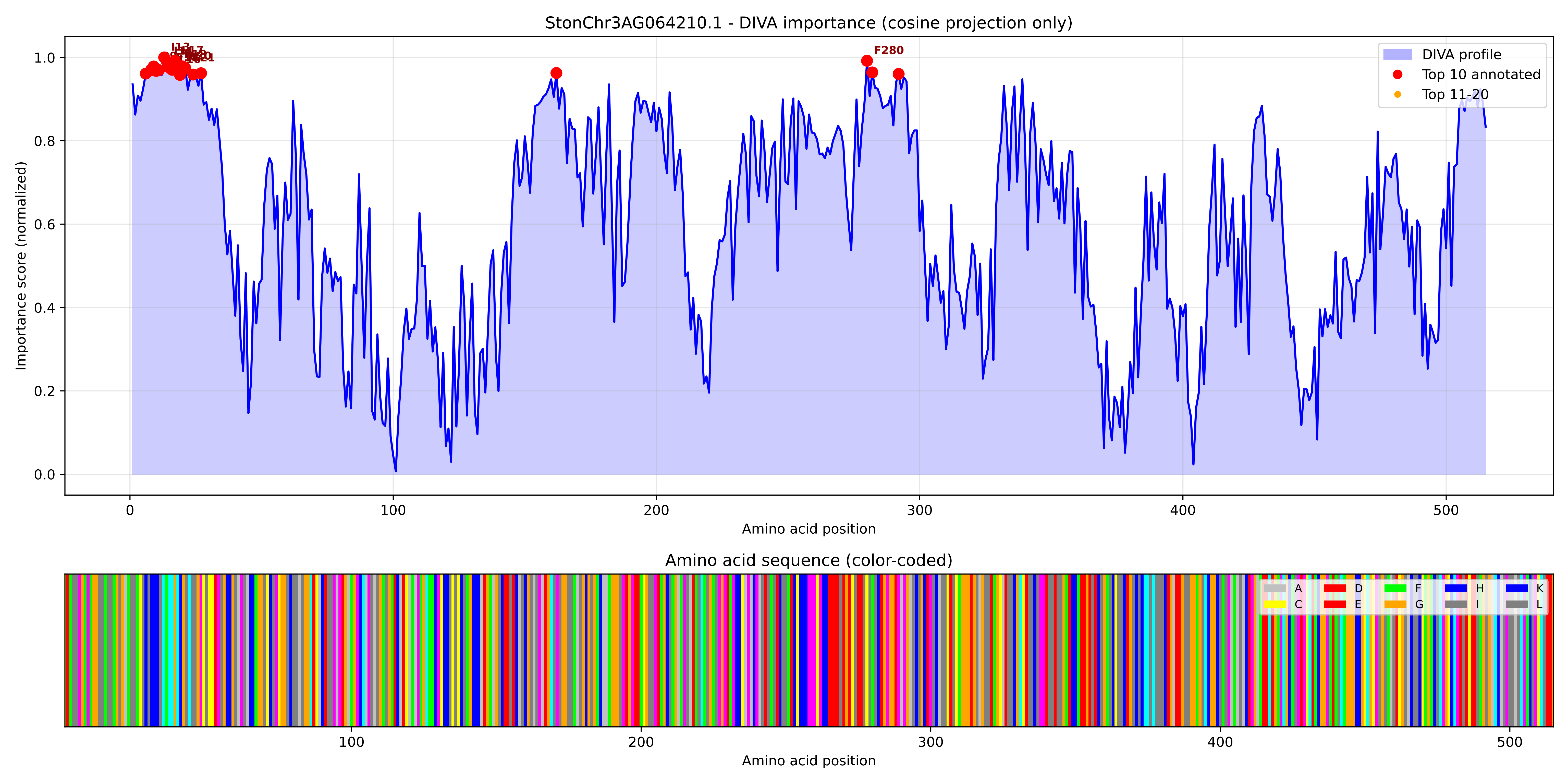

### StonChr3AG064210.1_importance.png

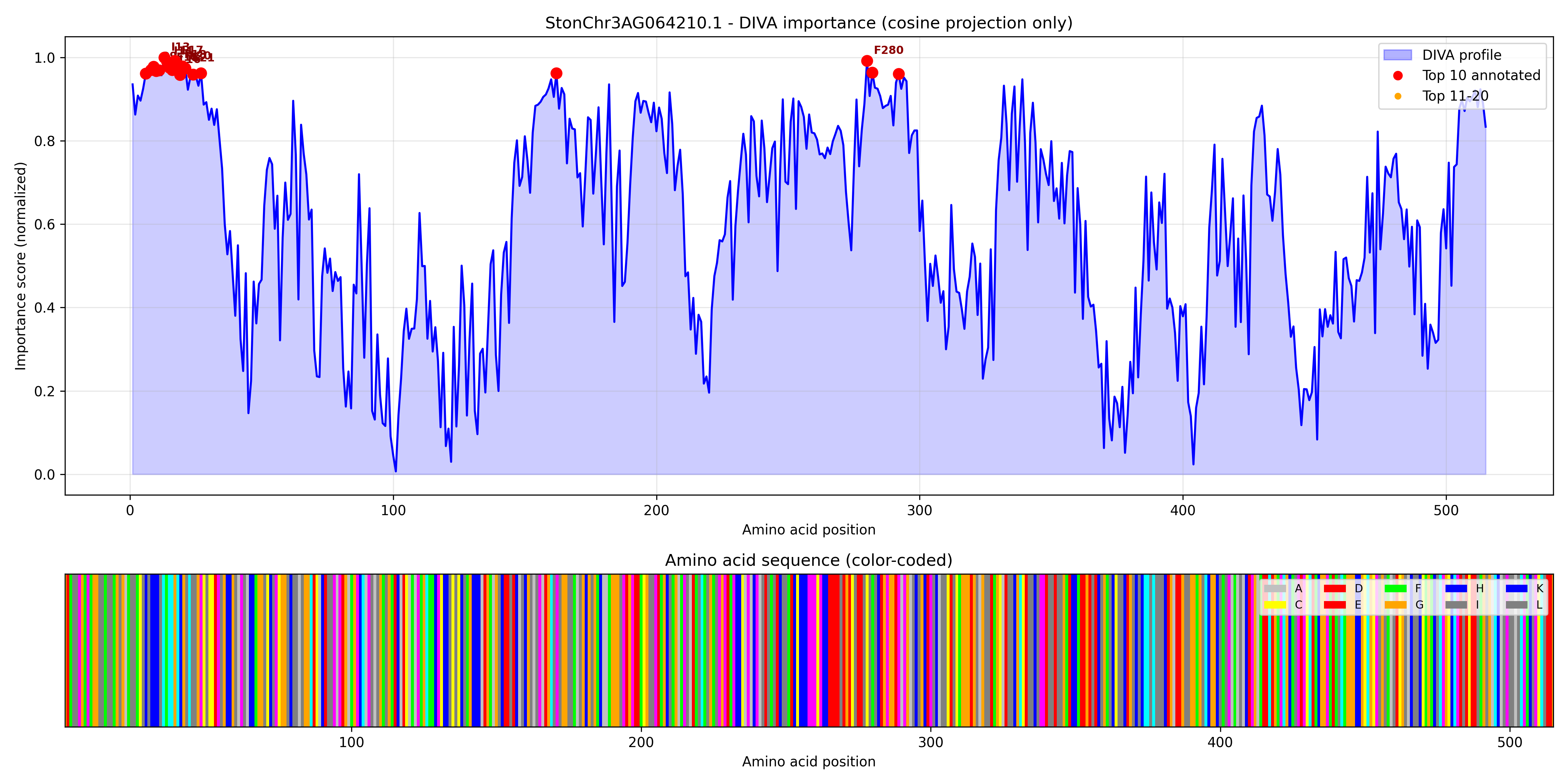
