## Supplementary figures and images for "The N-Terminus of *Sophora tonkinensis* Cytochrome P450s Evolves Neutrally yet Encodes Rich Functional Information: A Protein Language Model Analysis"

### population_profile.pdf

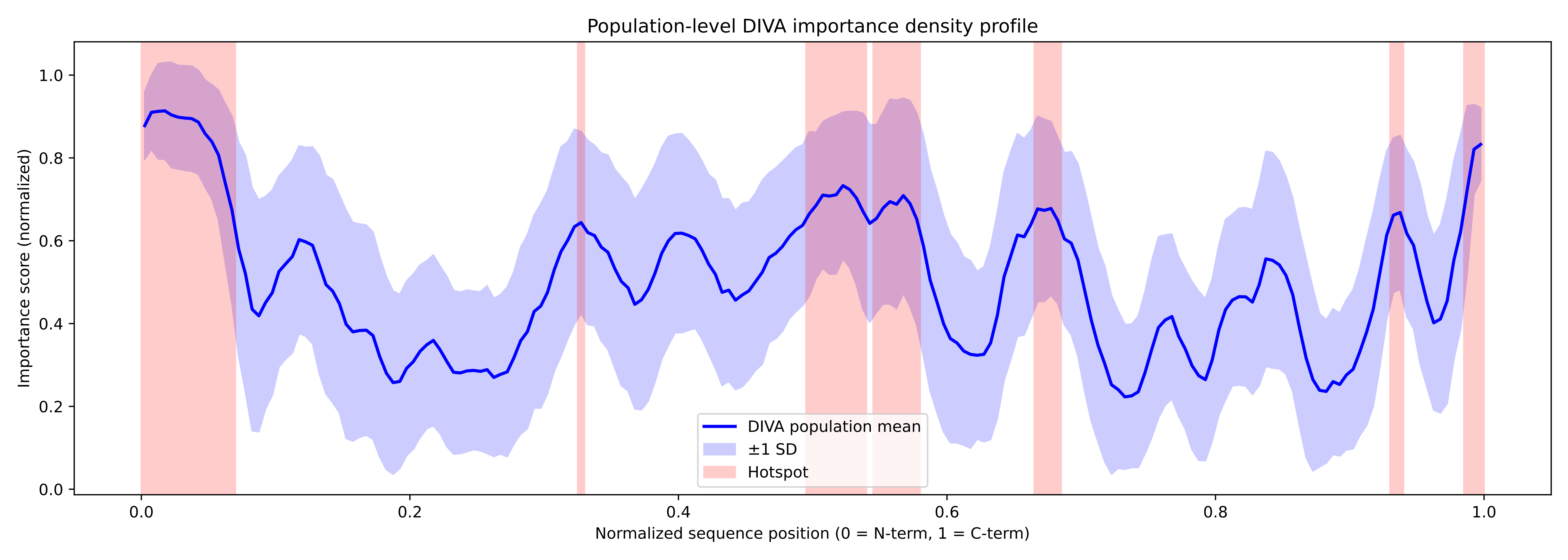

### population_profile.png

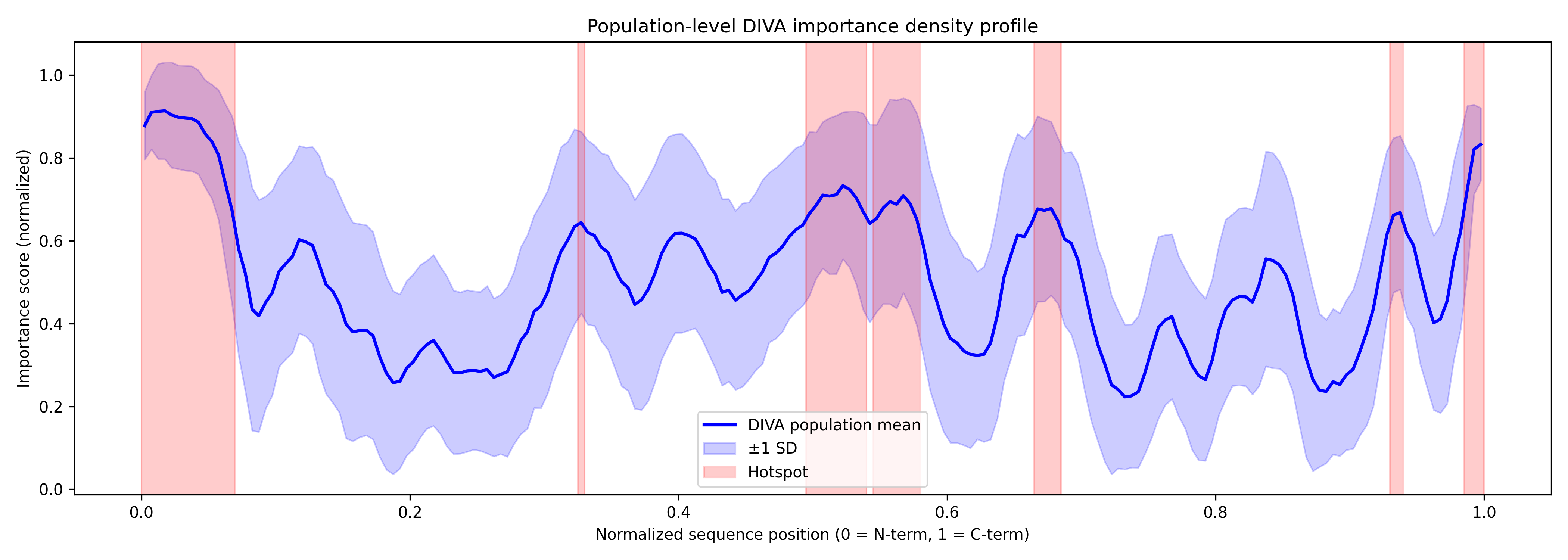

### StonChr1AG000330.1_importance.pdf

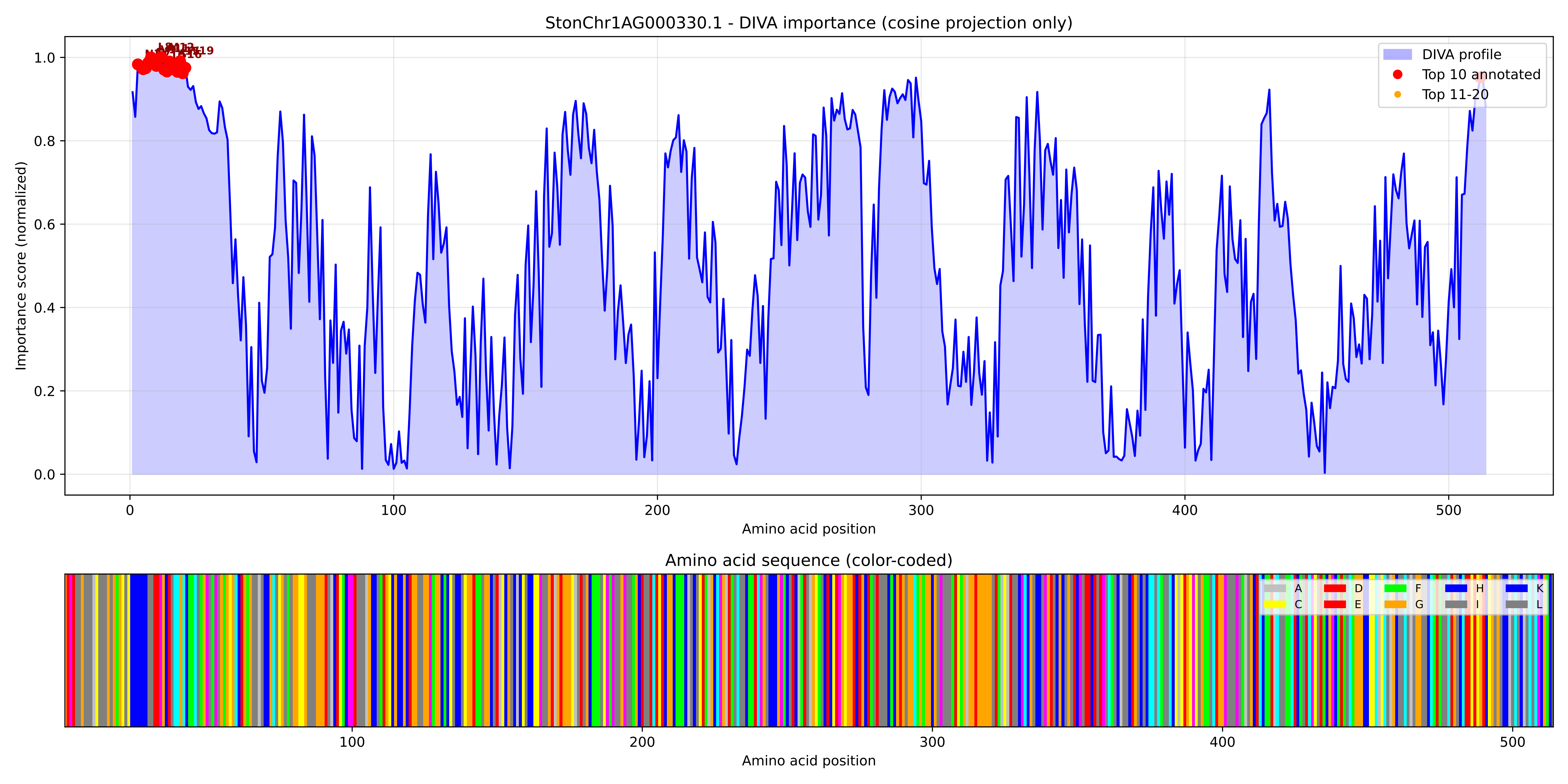

### StonChr1AG000330.1_importance.png

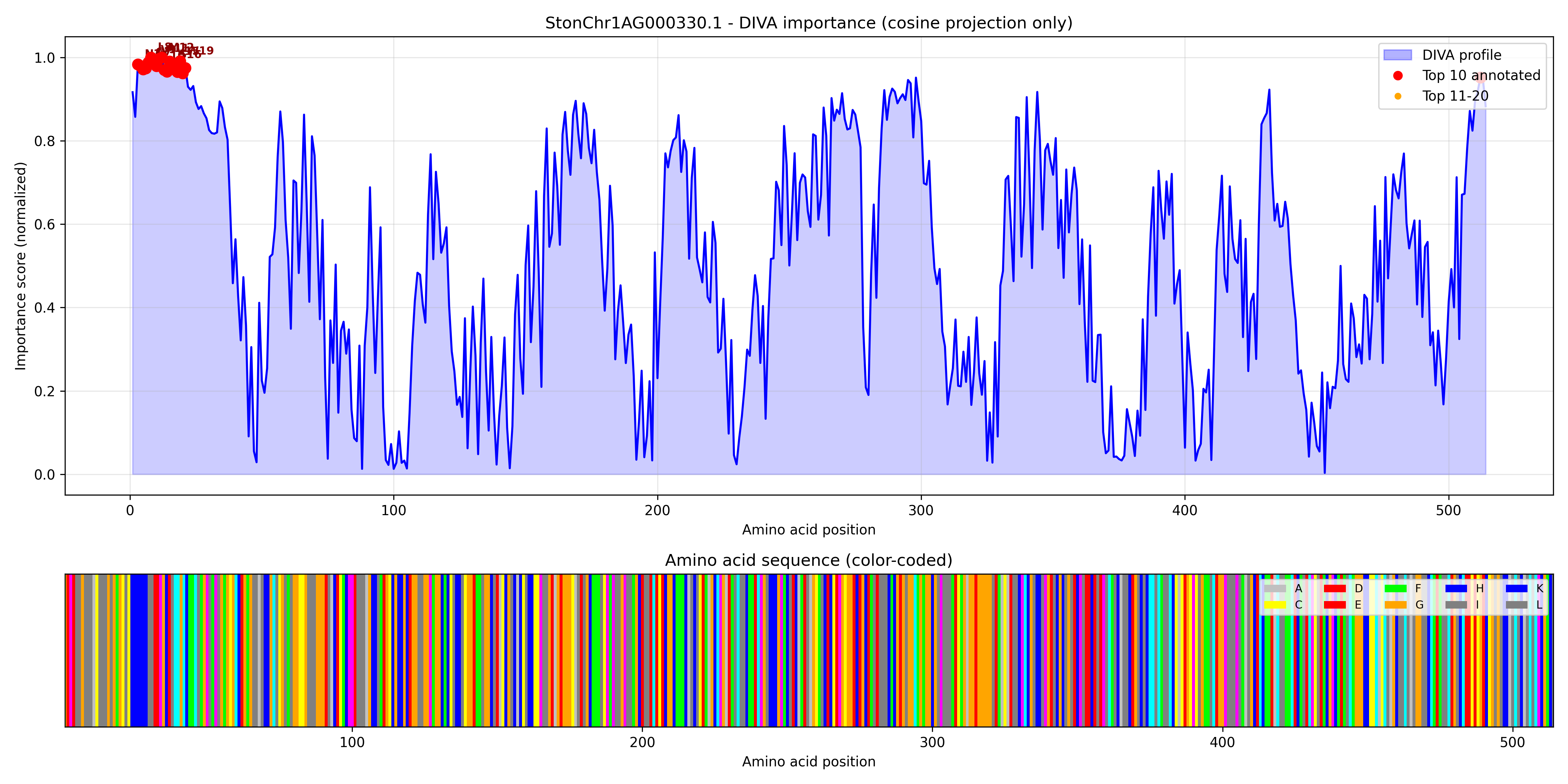

### StonChr1AG002740.1_importance.pdf

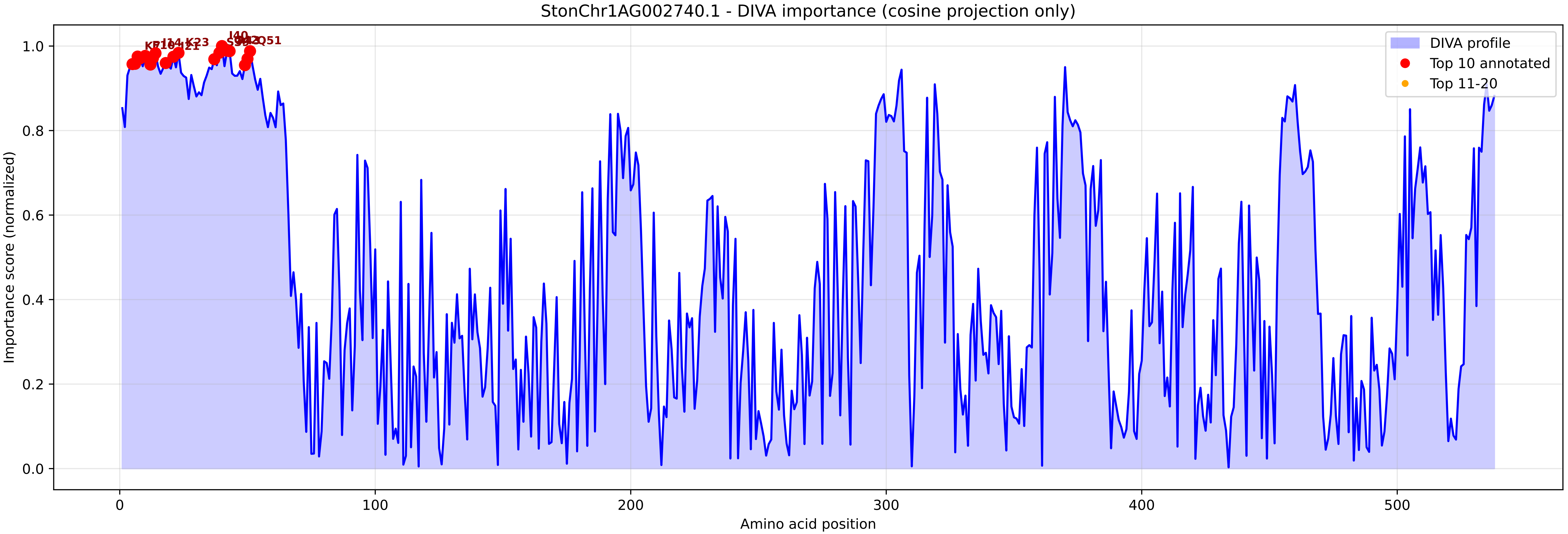

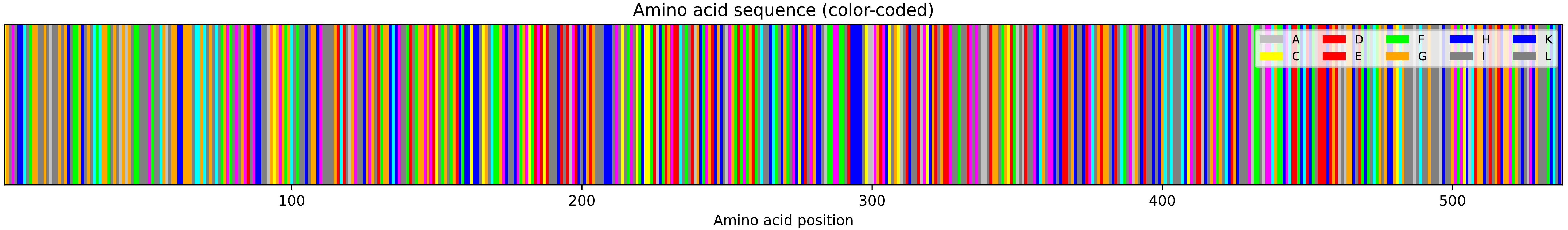

### StonChr1AG002740.1_importance.png

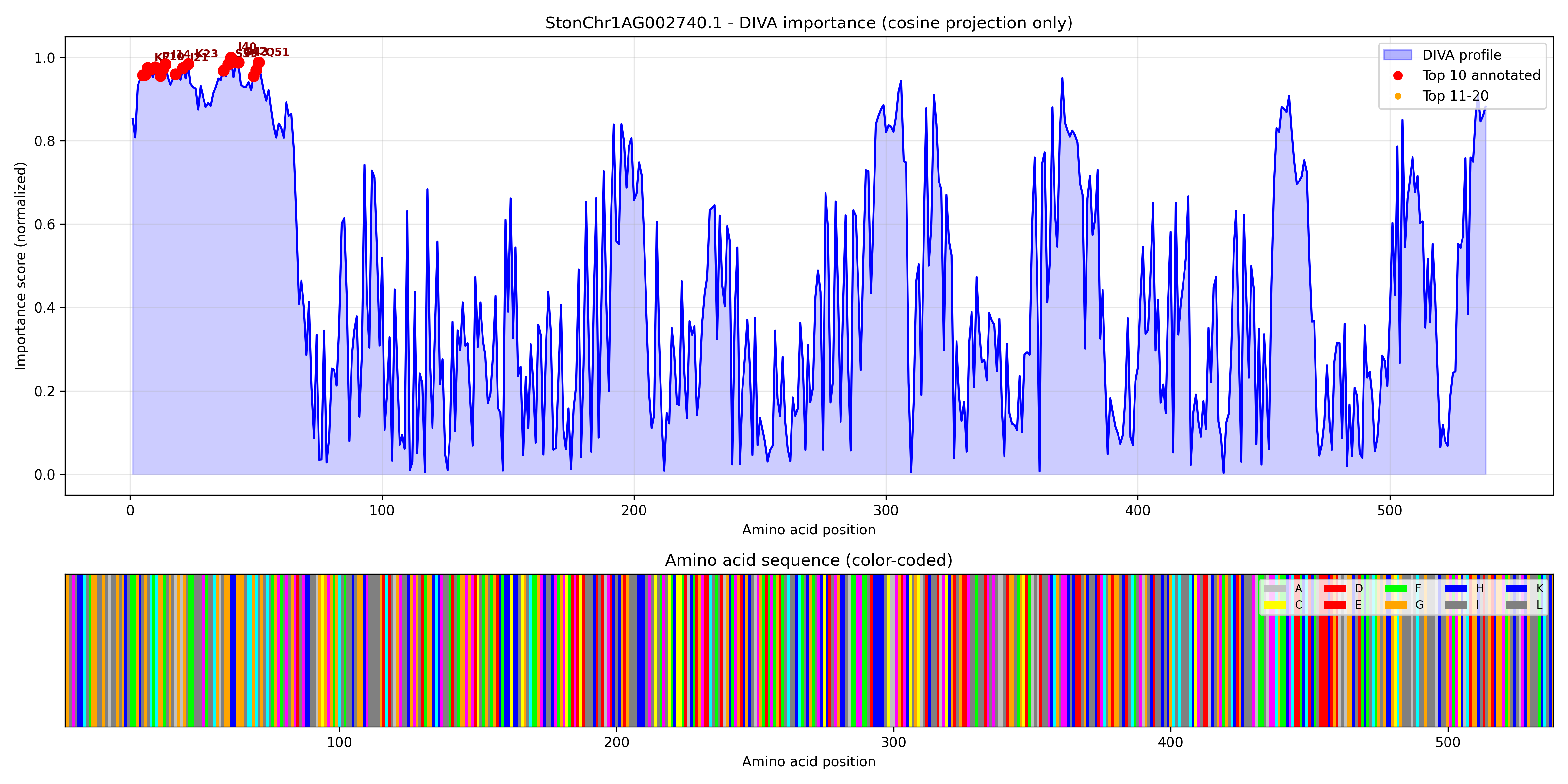

### StonChr1AG007090.1_importance.pdf

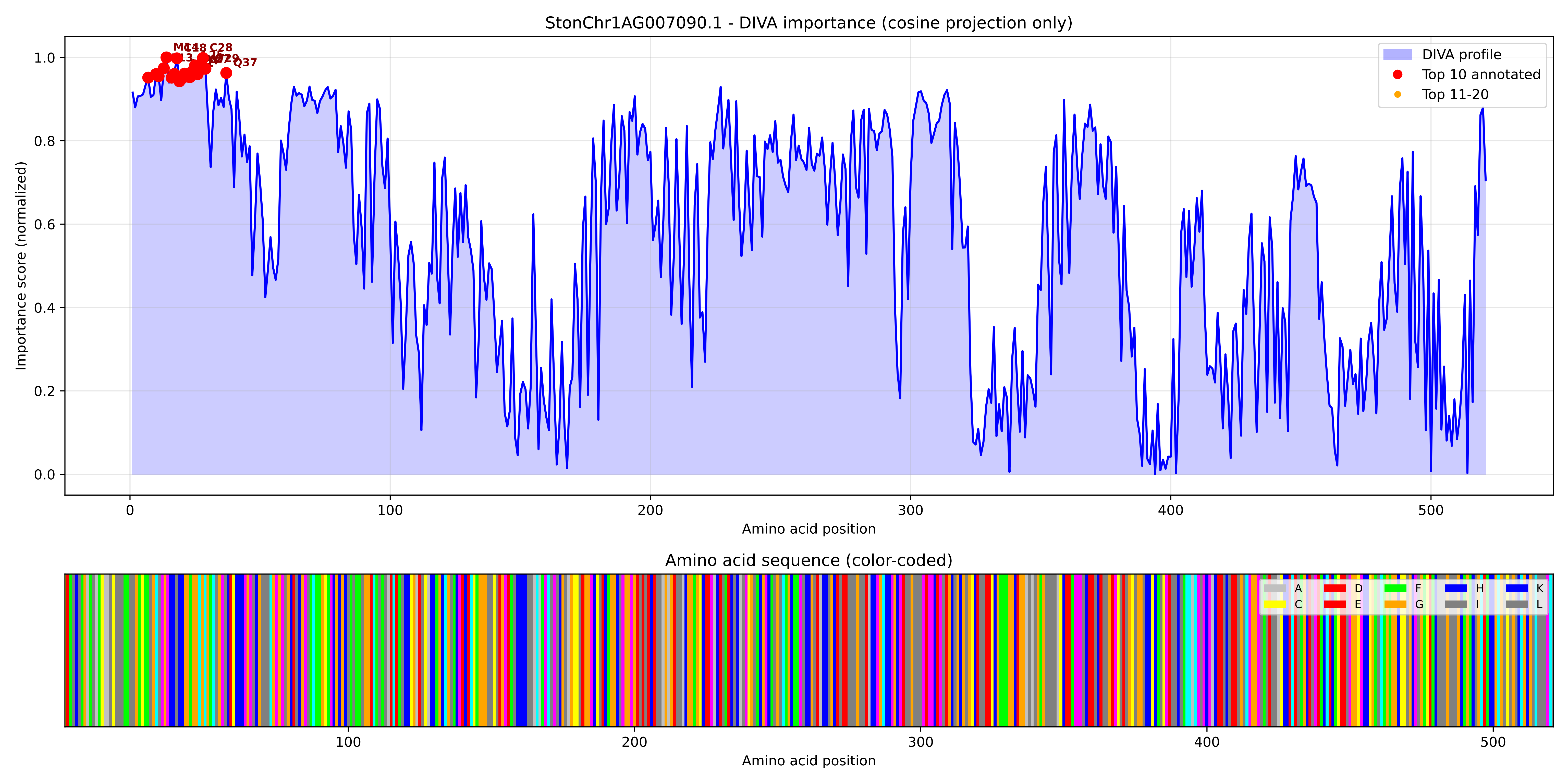

### StonChr1AG007090.1_importance.png

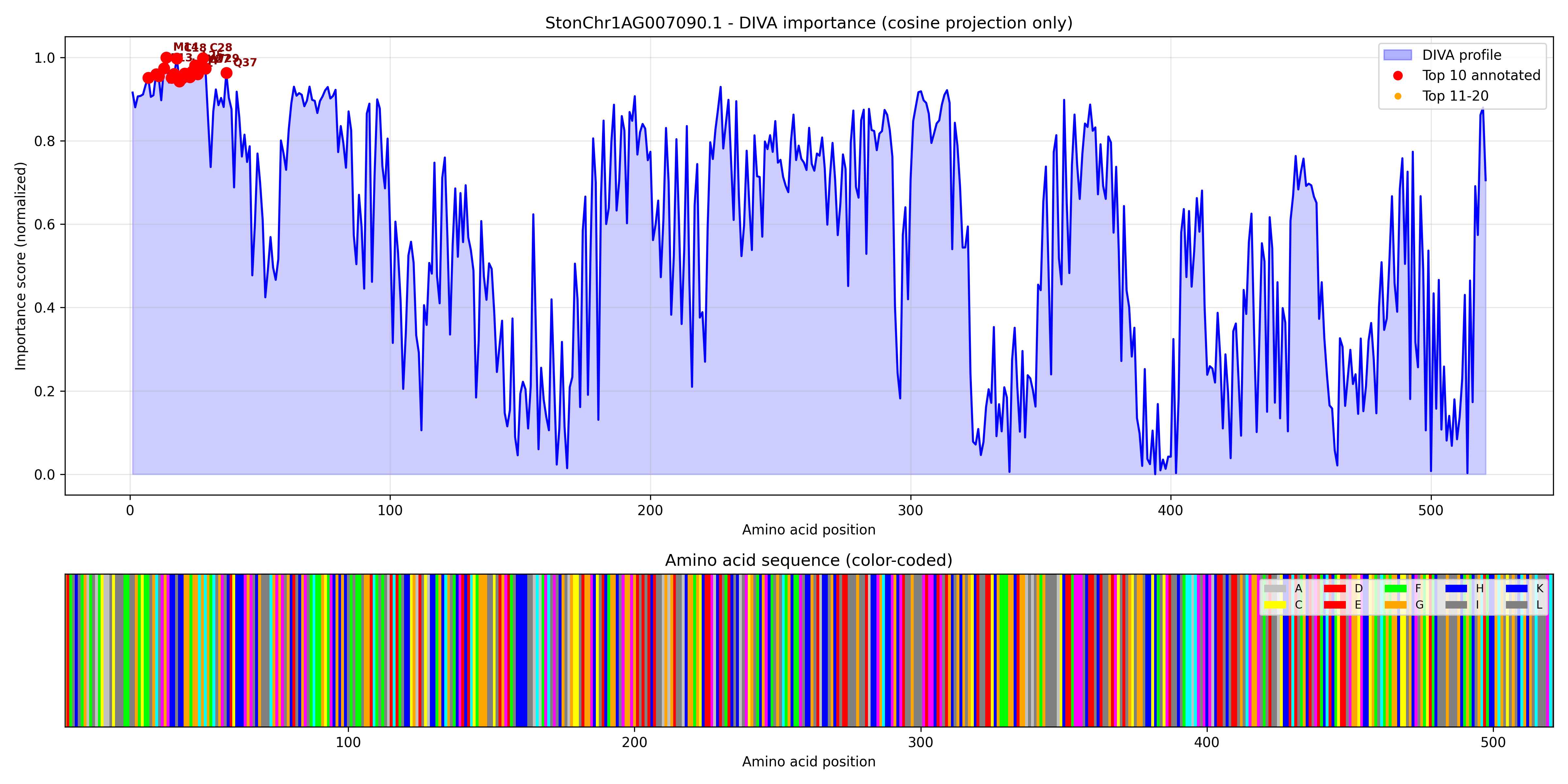

### StonChr1AG008680.1_importance.pdf

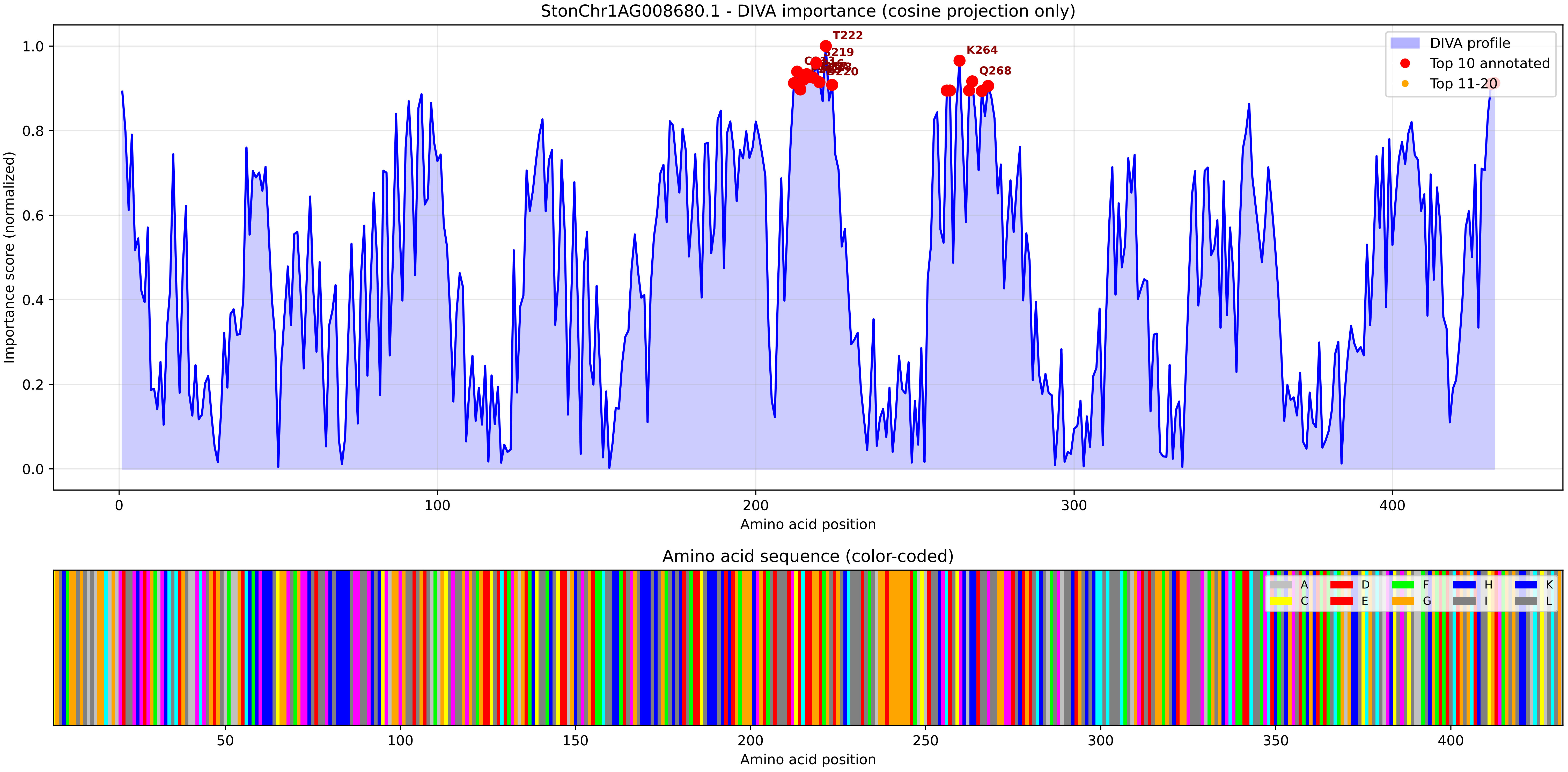

### StonChr1AG008680.1_importance.png

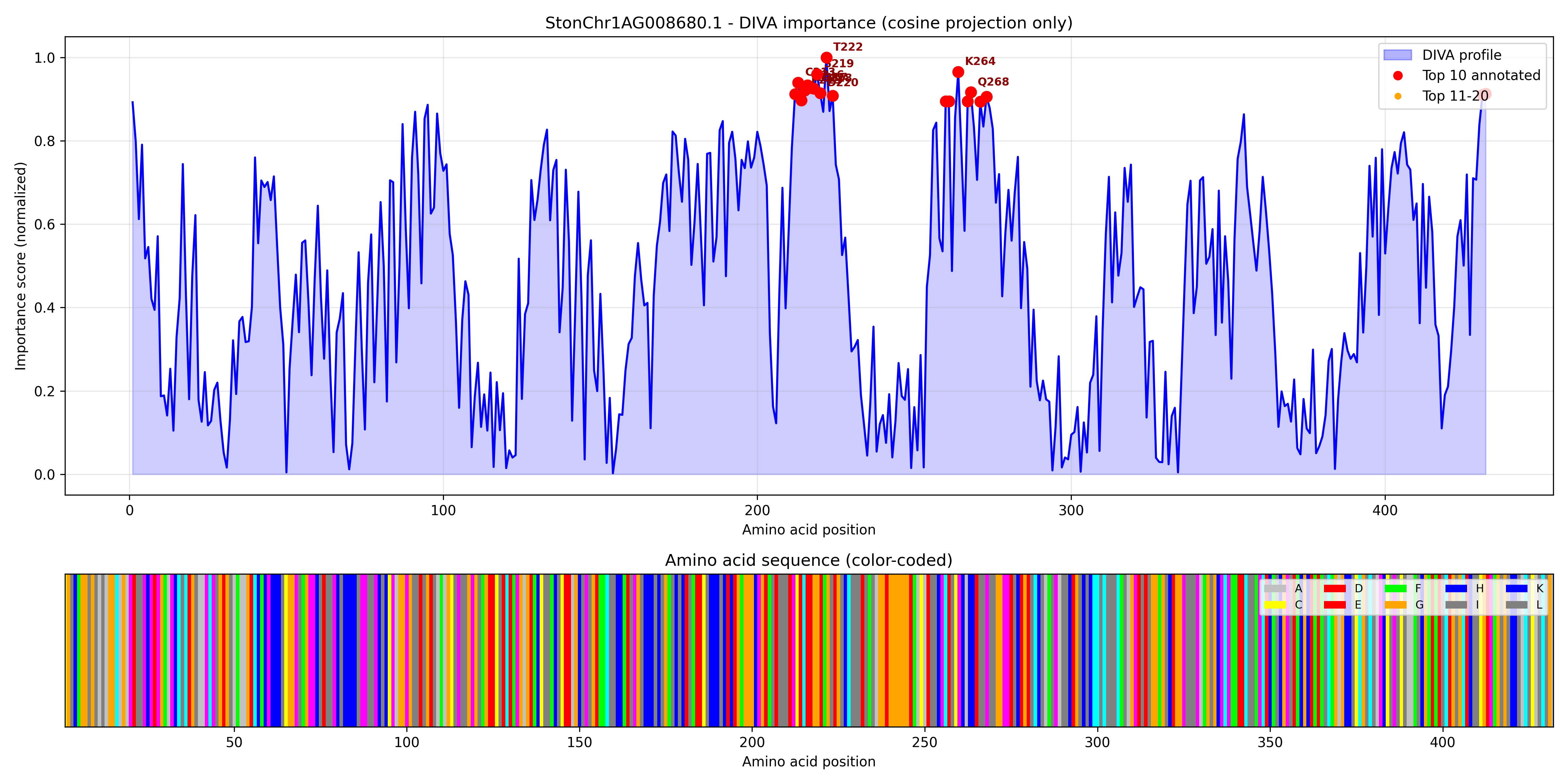

### StonChr1AG008690.1_importance.pdf

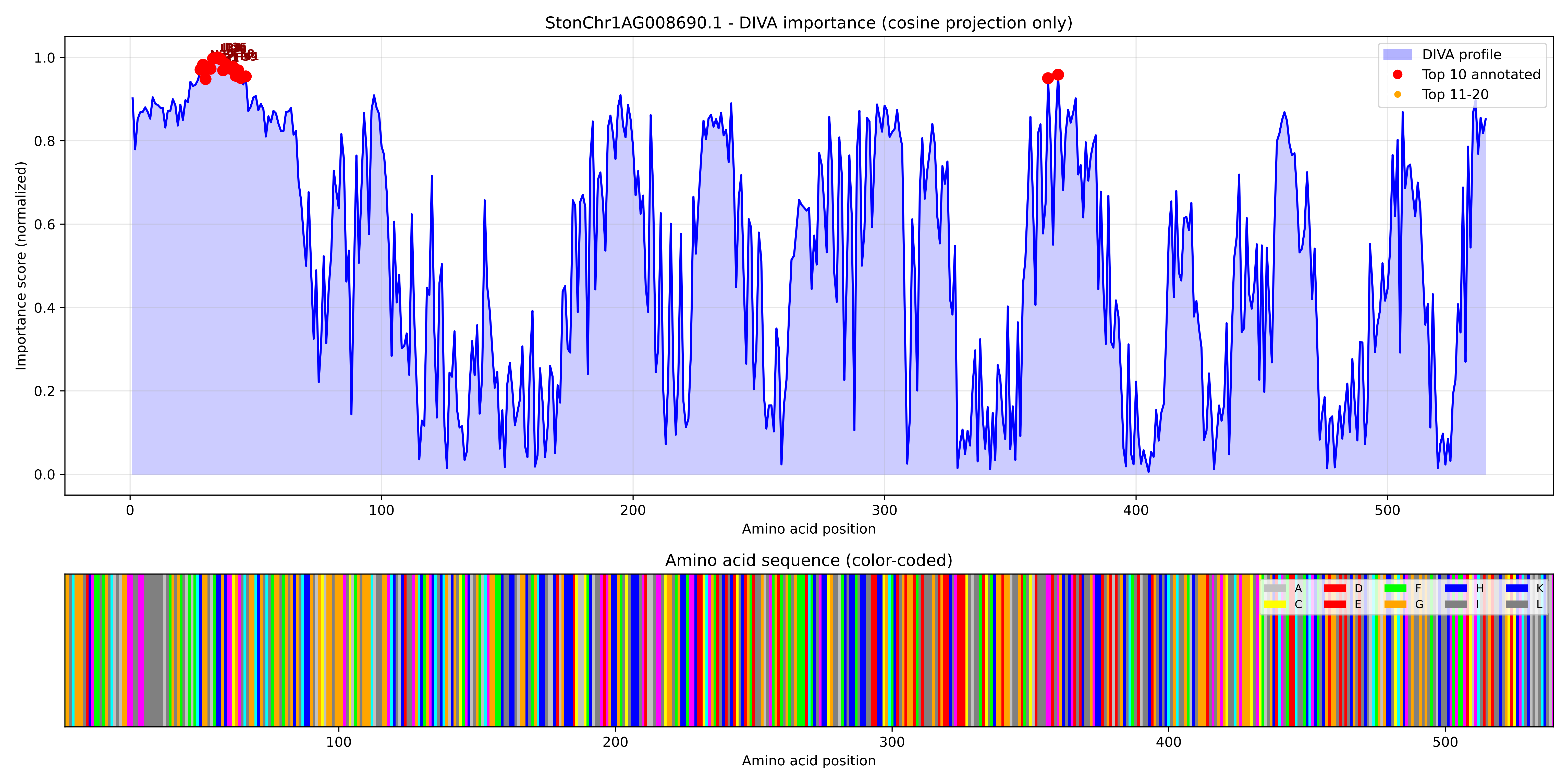

### StonChr1AG008690.1_importance.png

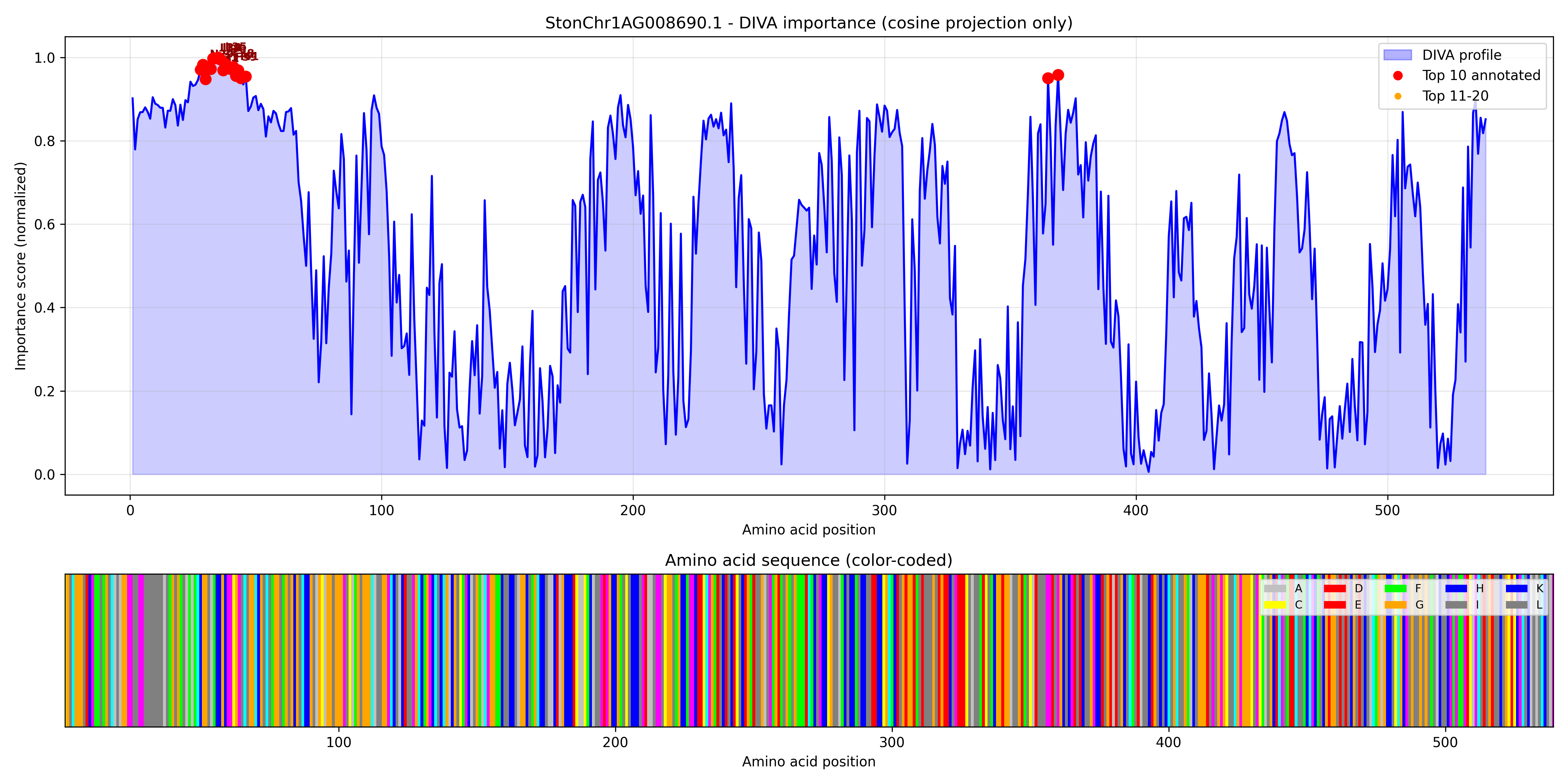

### StonChr1AG009990.1_importance.pdf

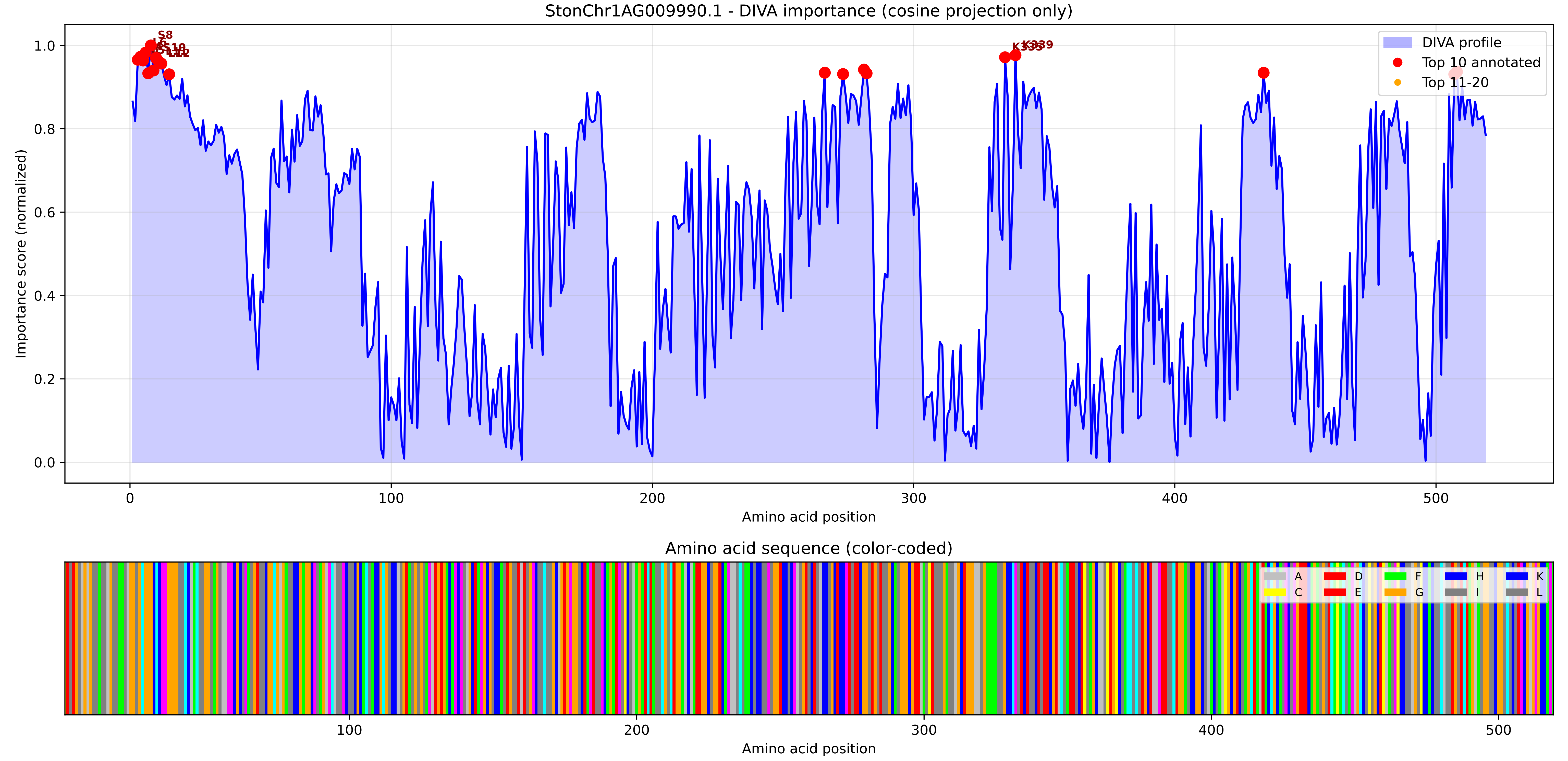

### StonChr1AG009990.1_importance.png

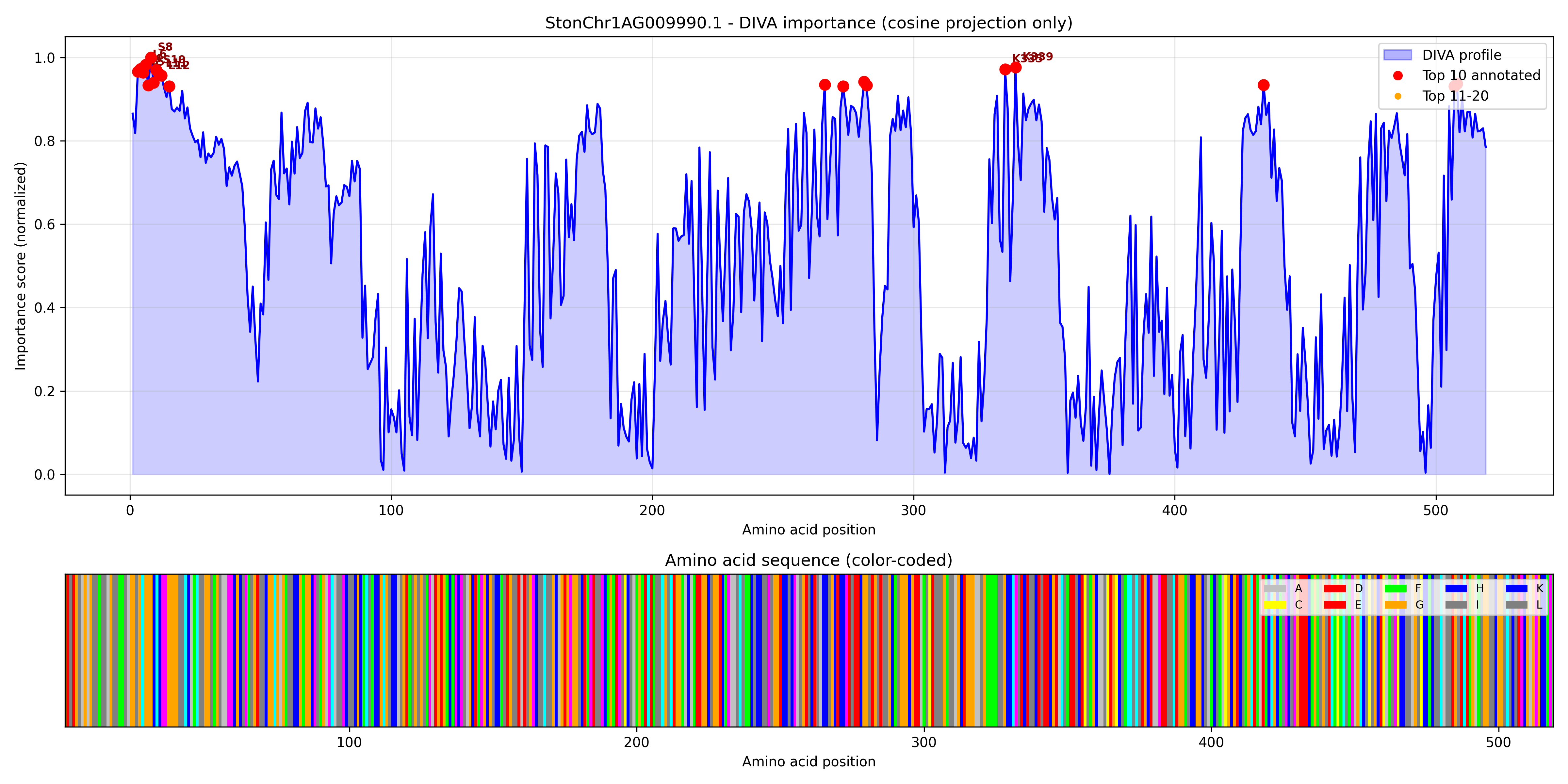

### StonChr1AG019900.1_importance.pdf

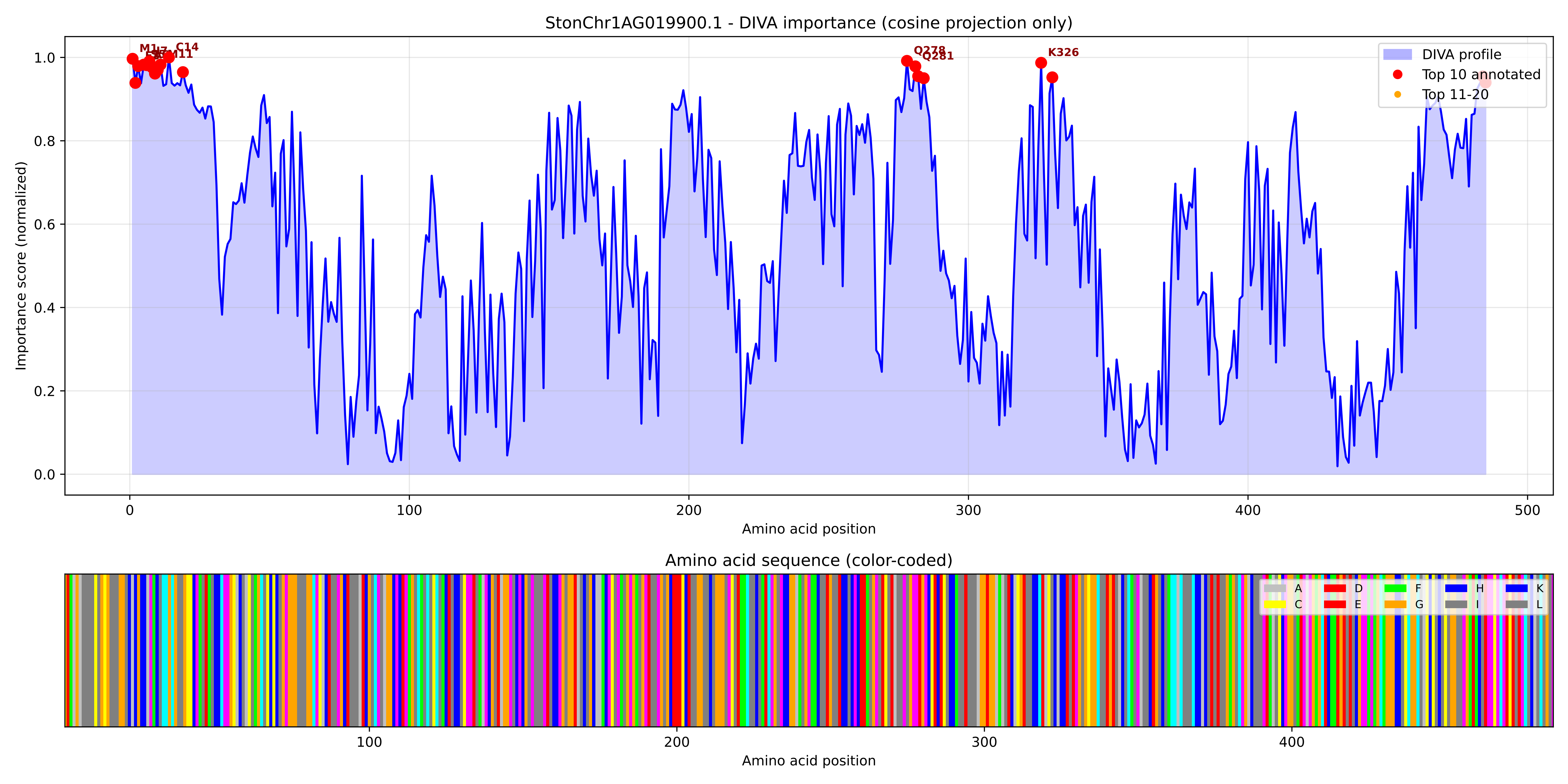

### StonChr1AG019900.1_importance.png

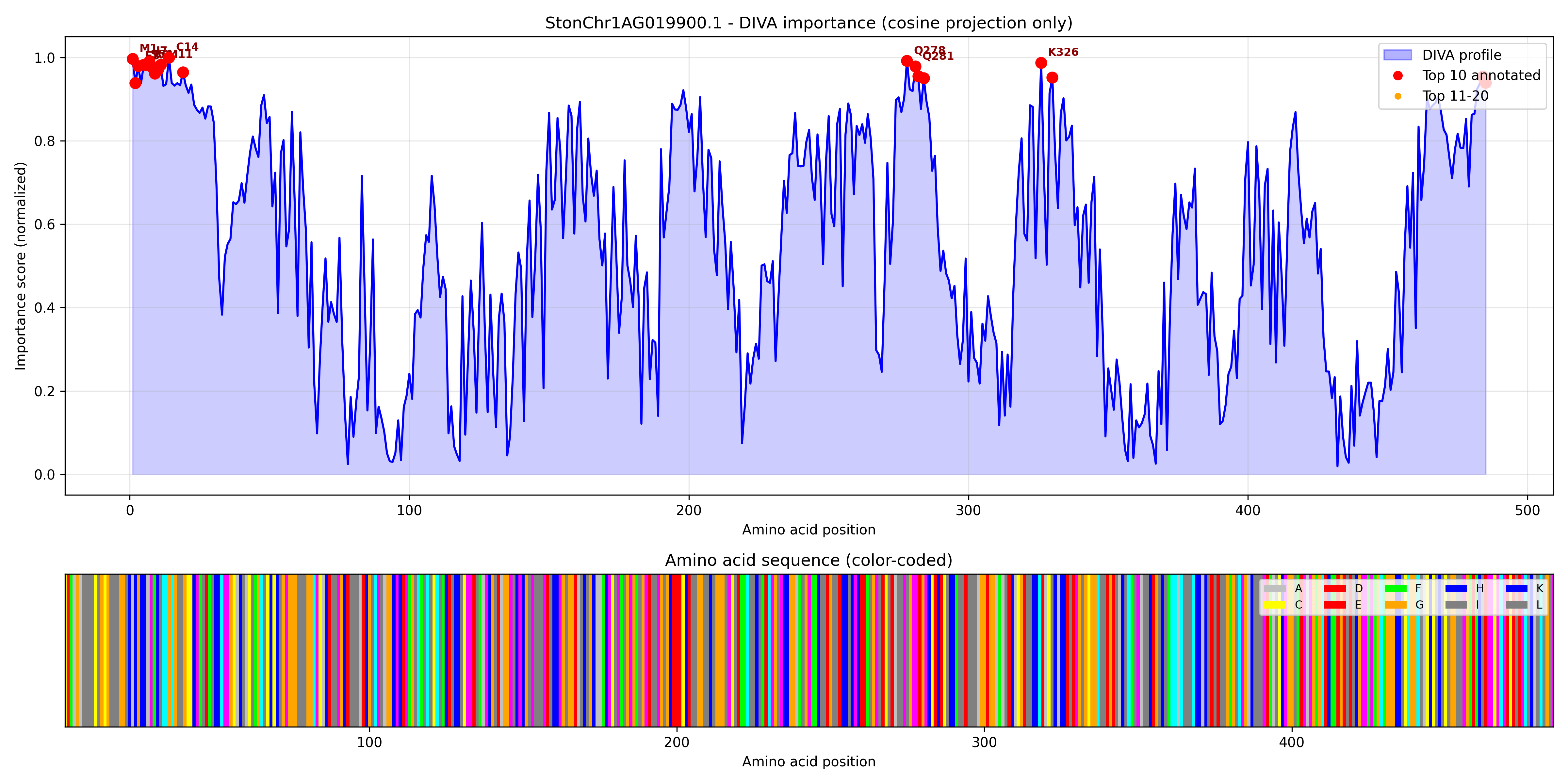

### StonChr1AG019940.1_importance.pdf

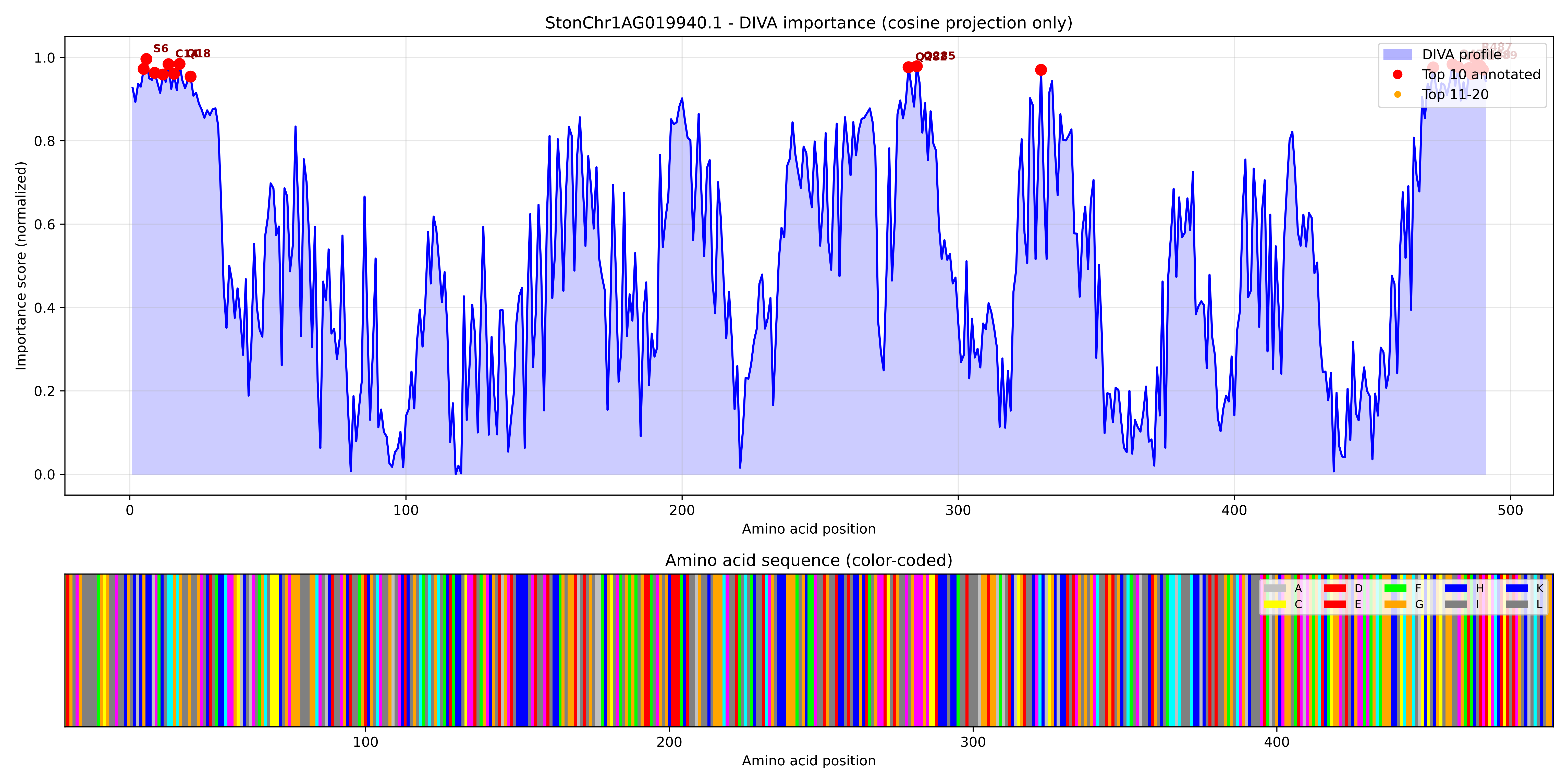

### StonChr1AG019940.1_importance.png

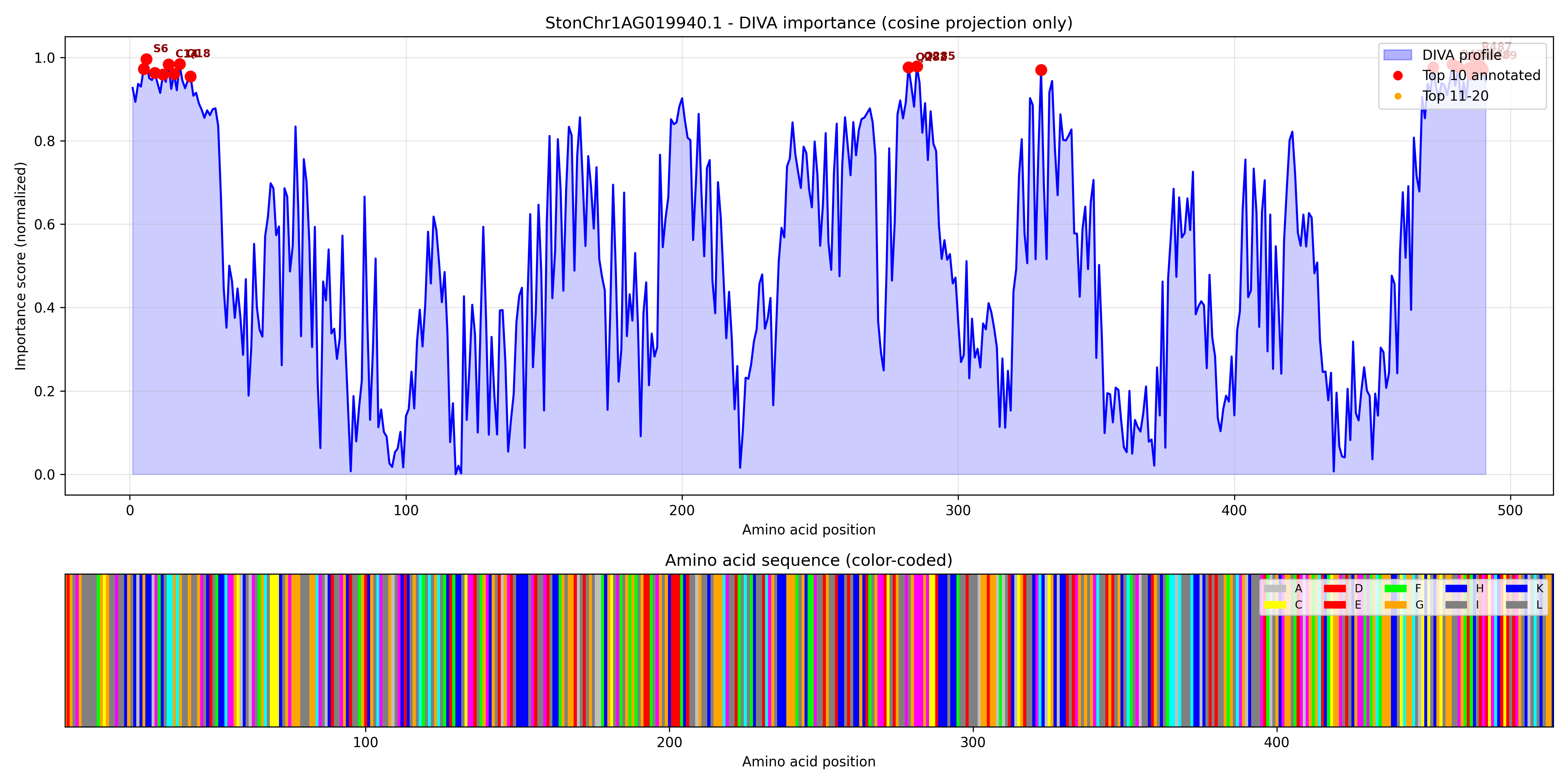

### StonChr1AG020090.1_importance.pdf

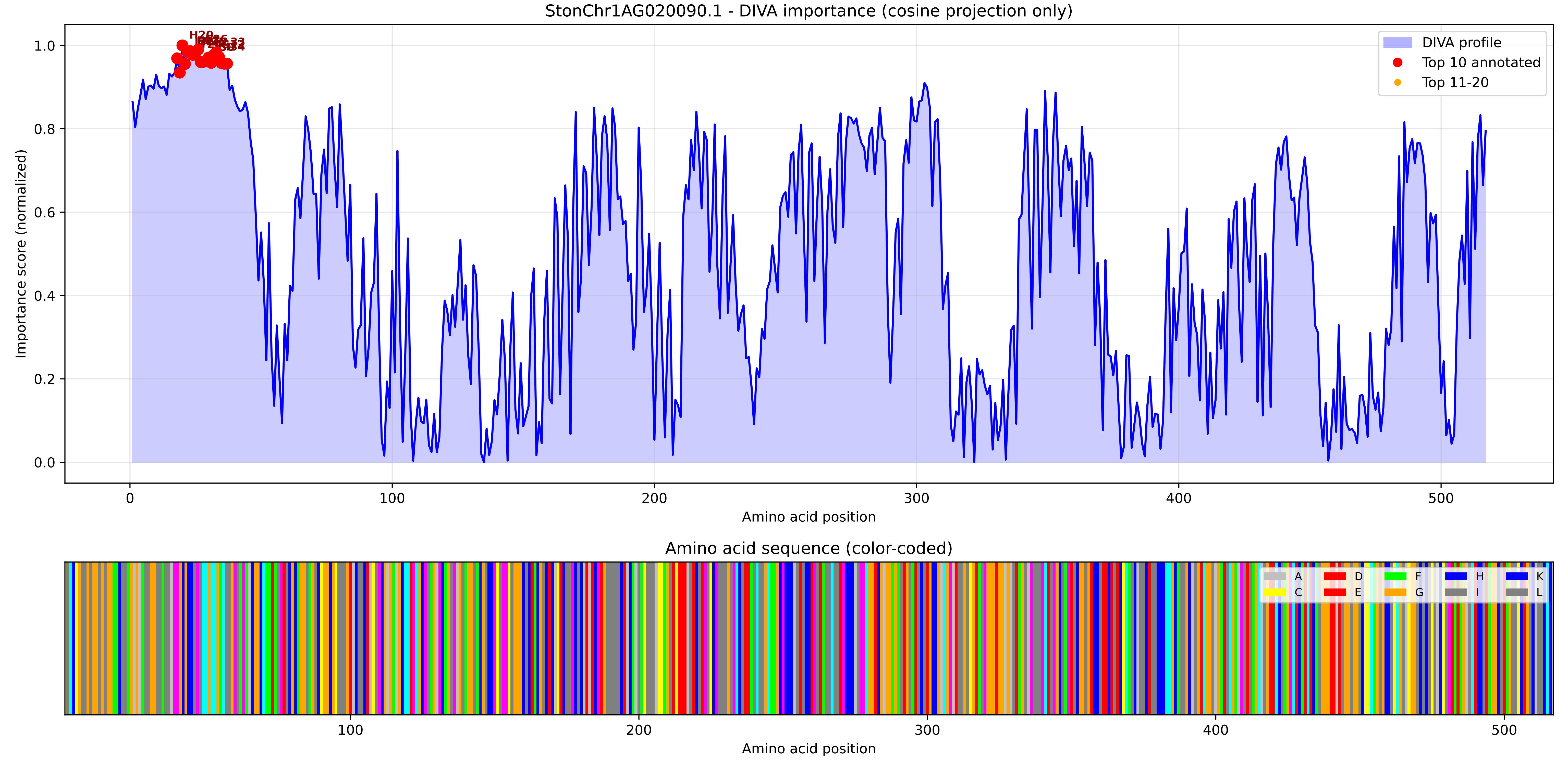

### StonChr1AG020090.1_importance.png

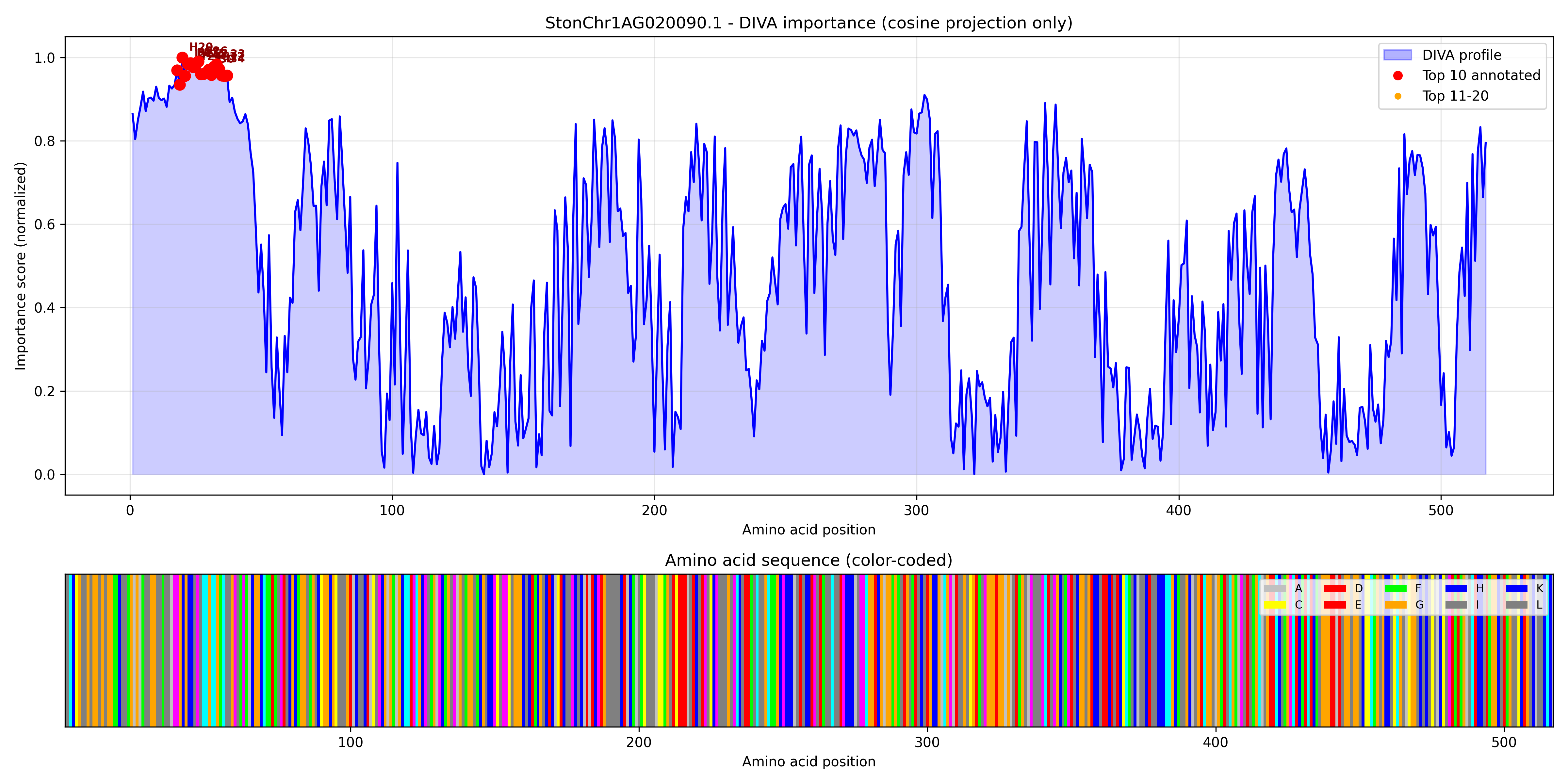

### StonChr1AG020200.1_importance.pdf

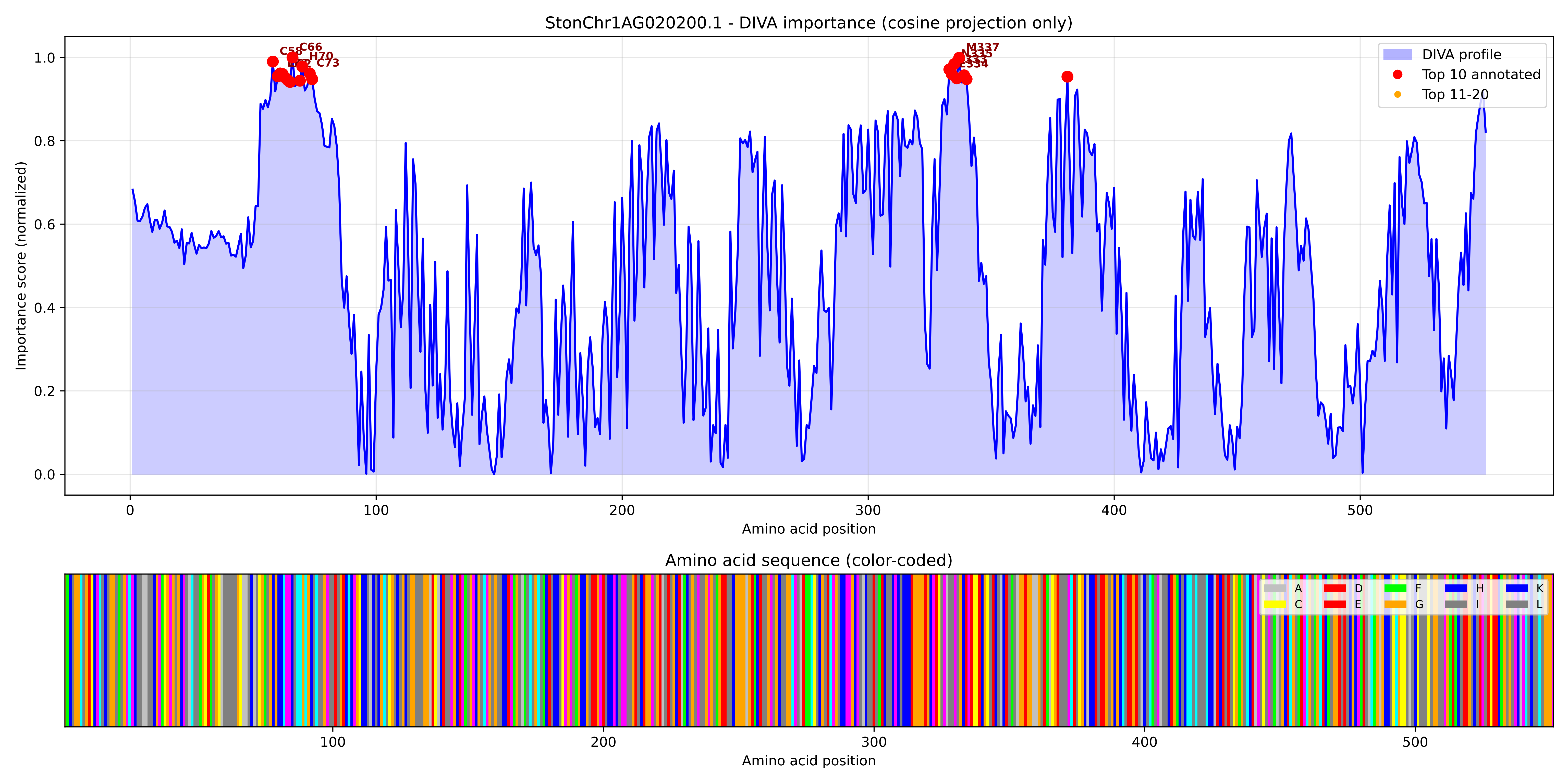

### StonChr1AG020200.1_importance.png

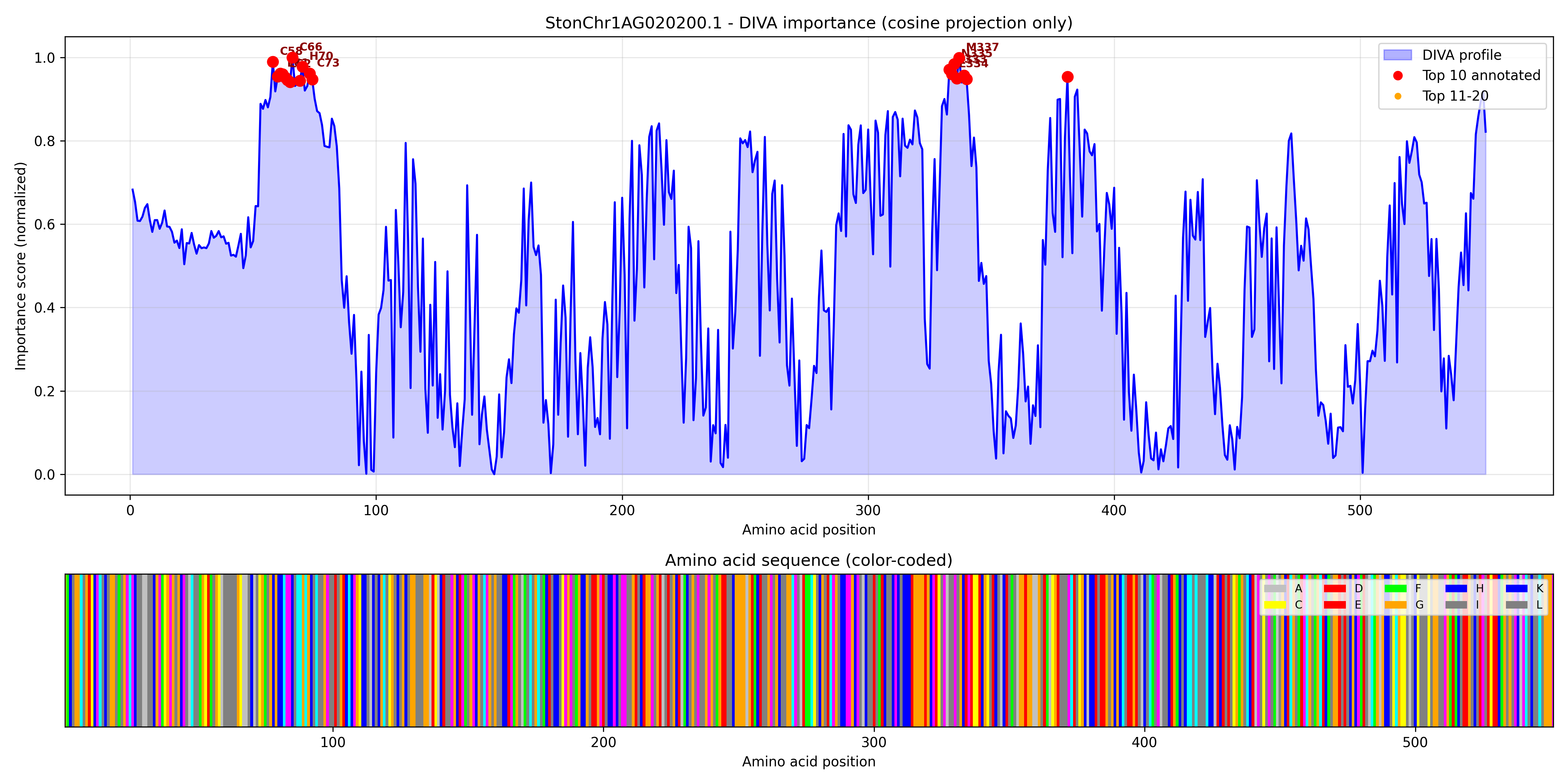

### StonChr1AG024220.1_importance.pdf

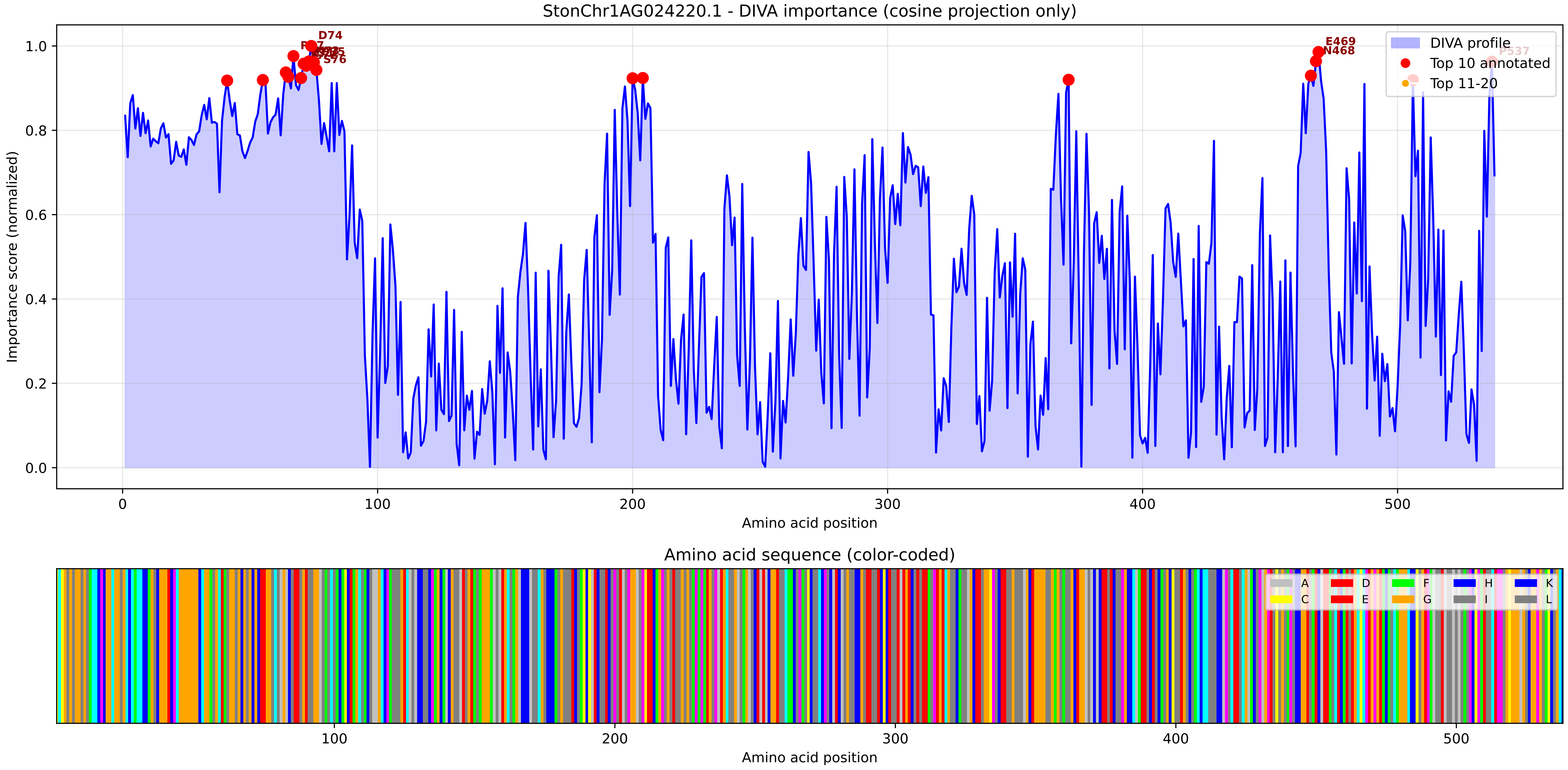

### StonChr1AG024220.1_importance.png

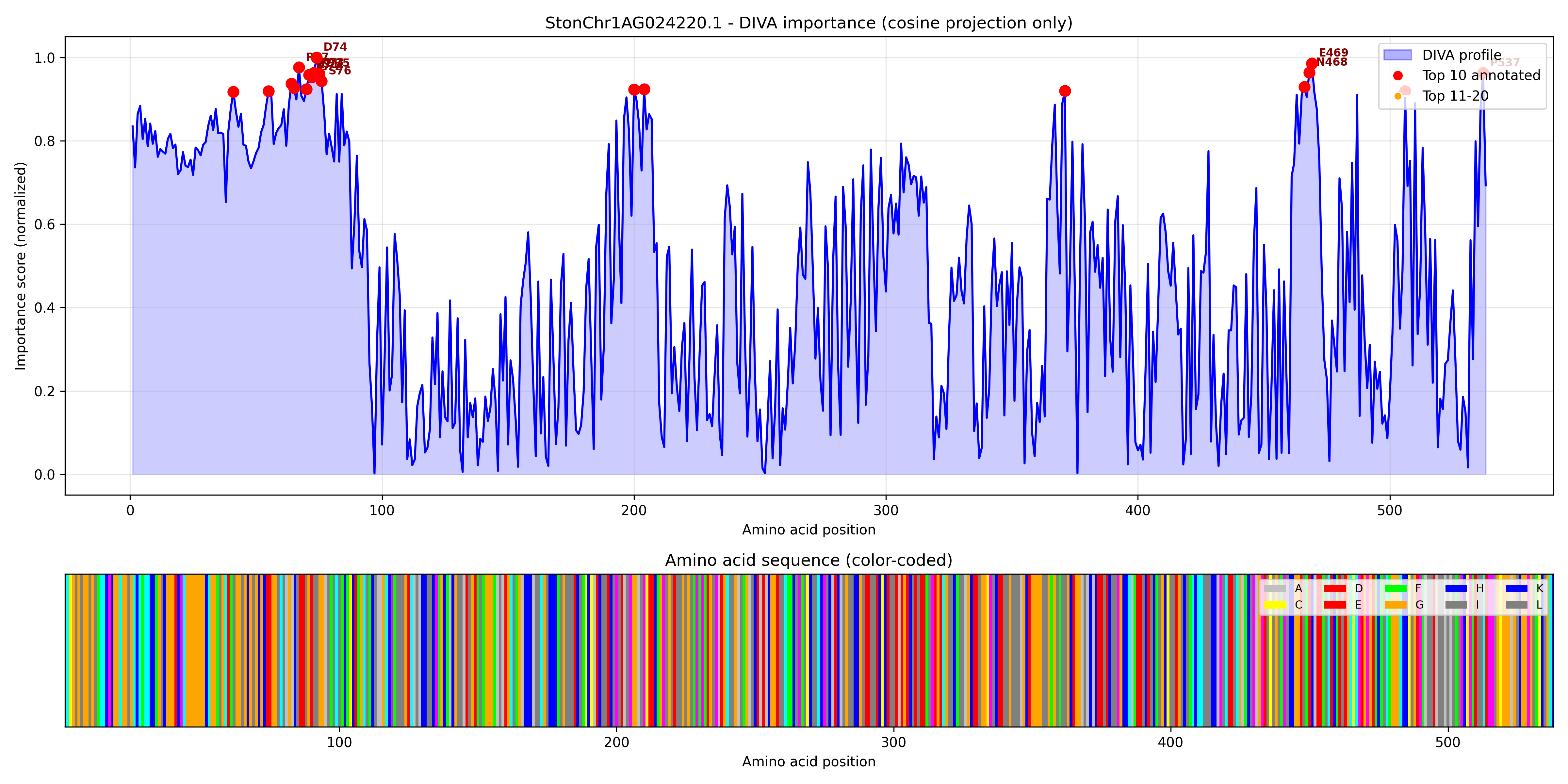

### StonChr1AG026330.1_importance.pdf

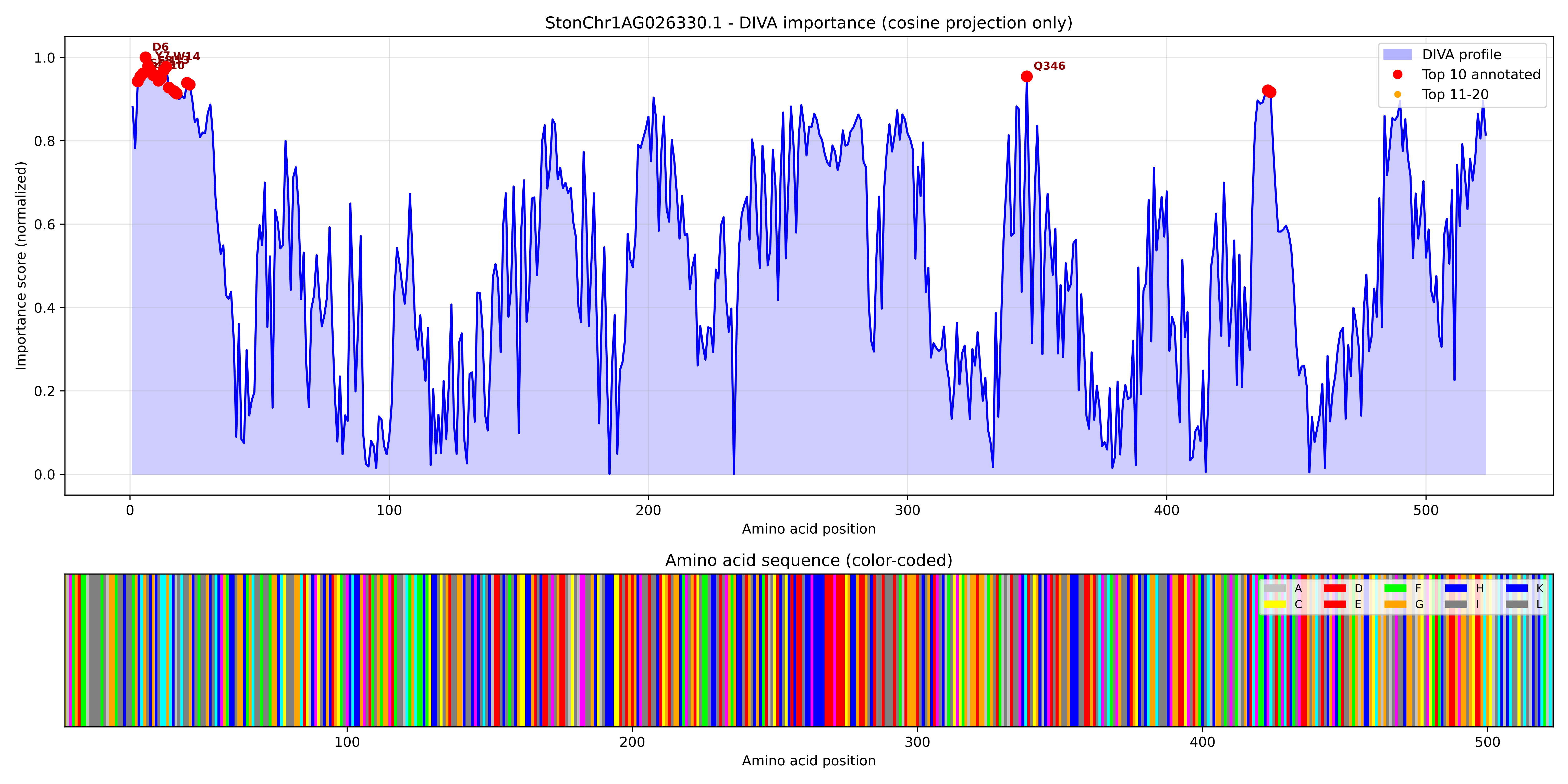

### StonChr1AG026330.1_importance.png

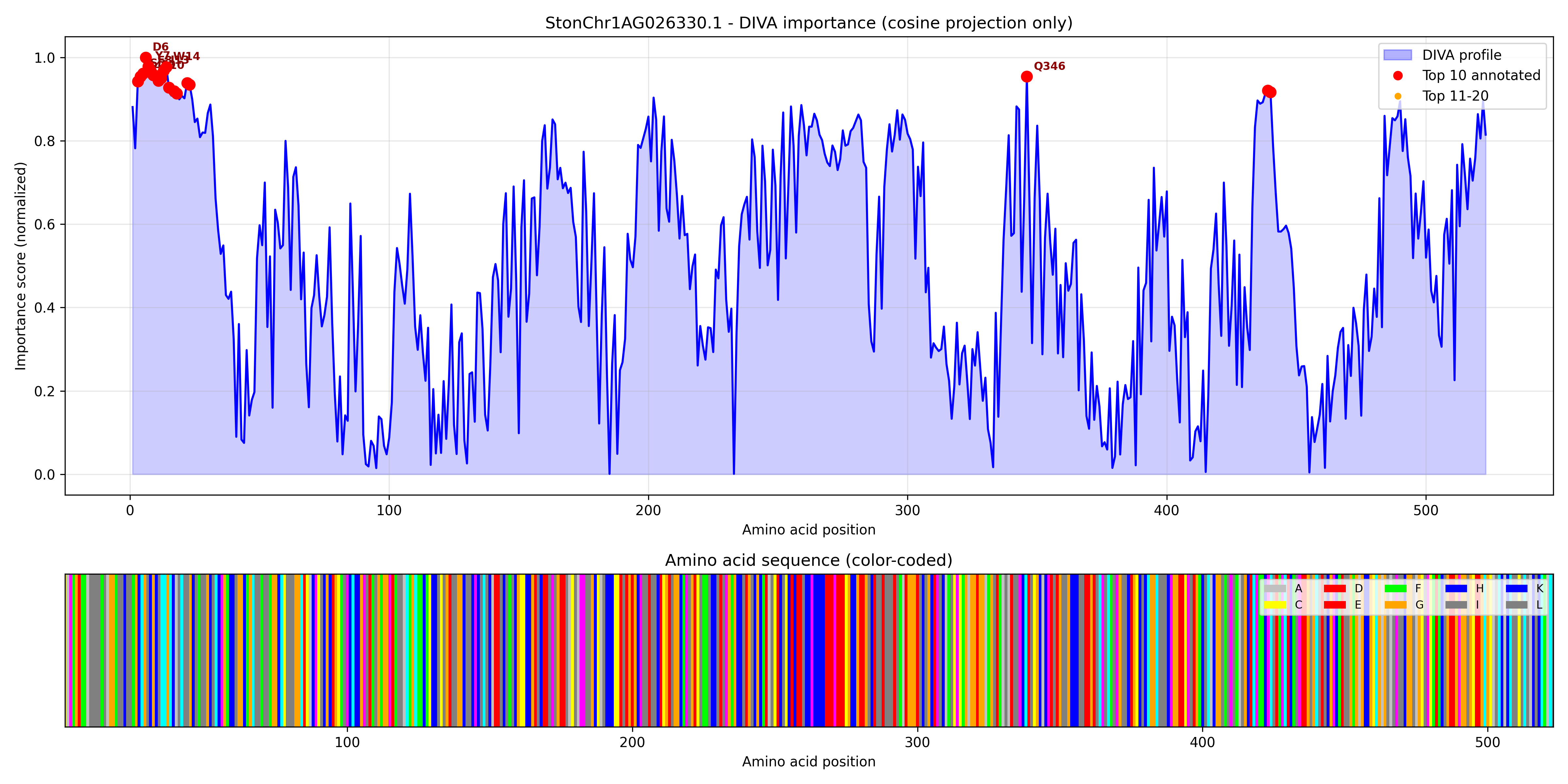

### StonChr1AG027680.1_importance.pdf

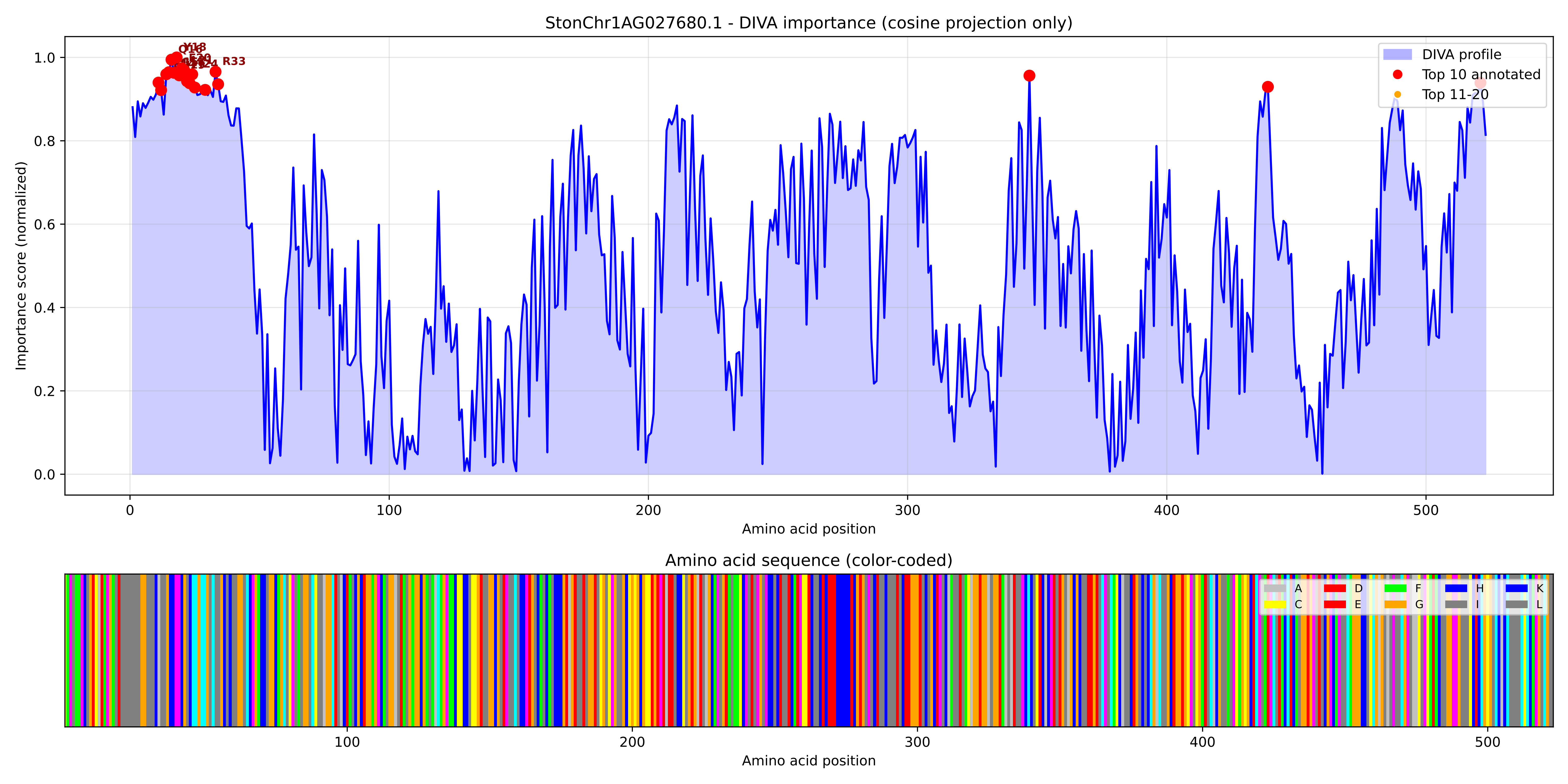

### StonChr1AG027680.1_importance.png

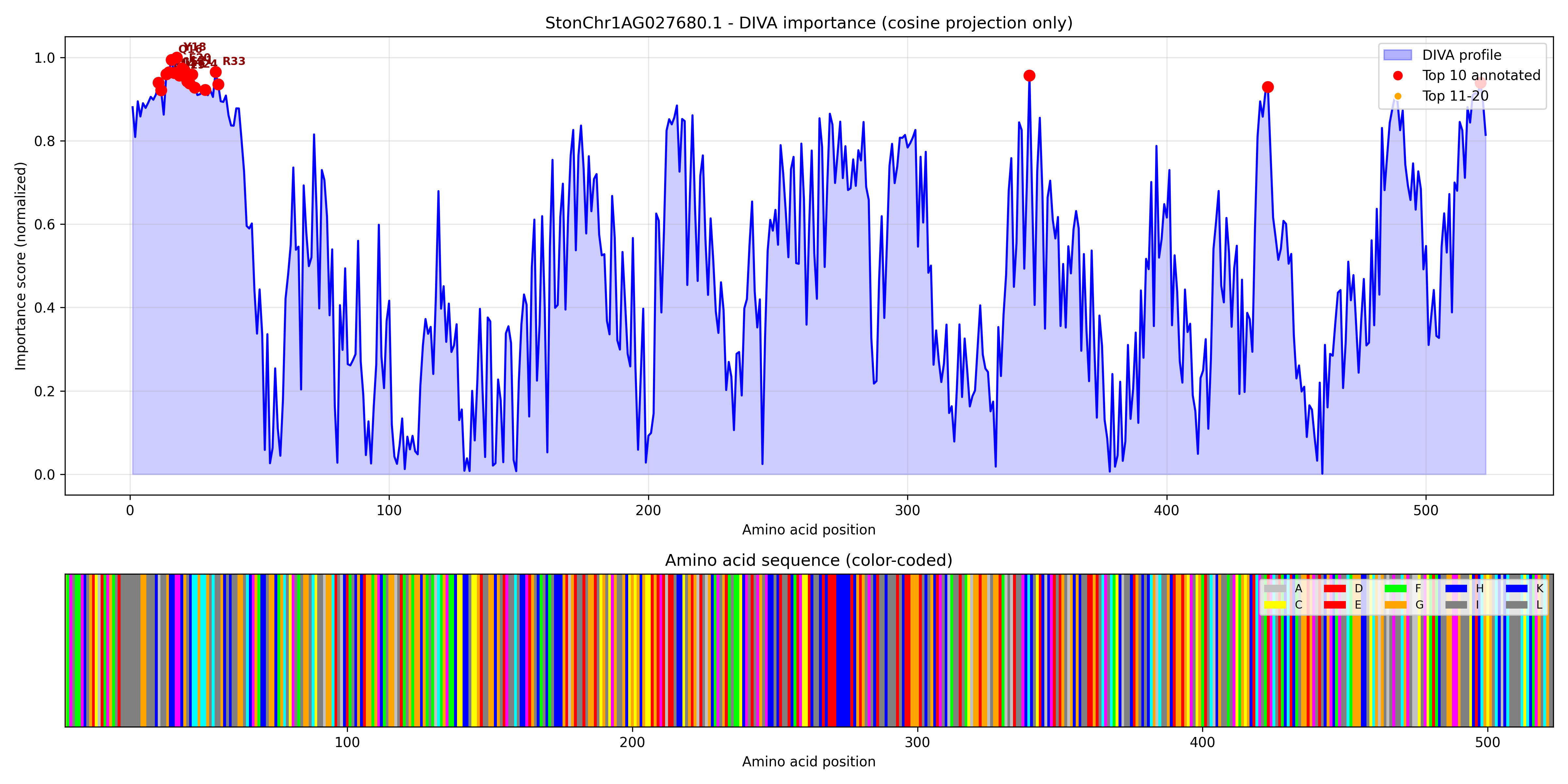

### StonChr1AG027700.1_importance.pdf

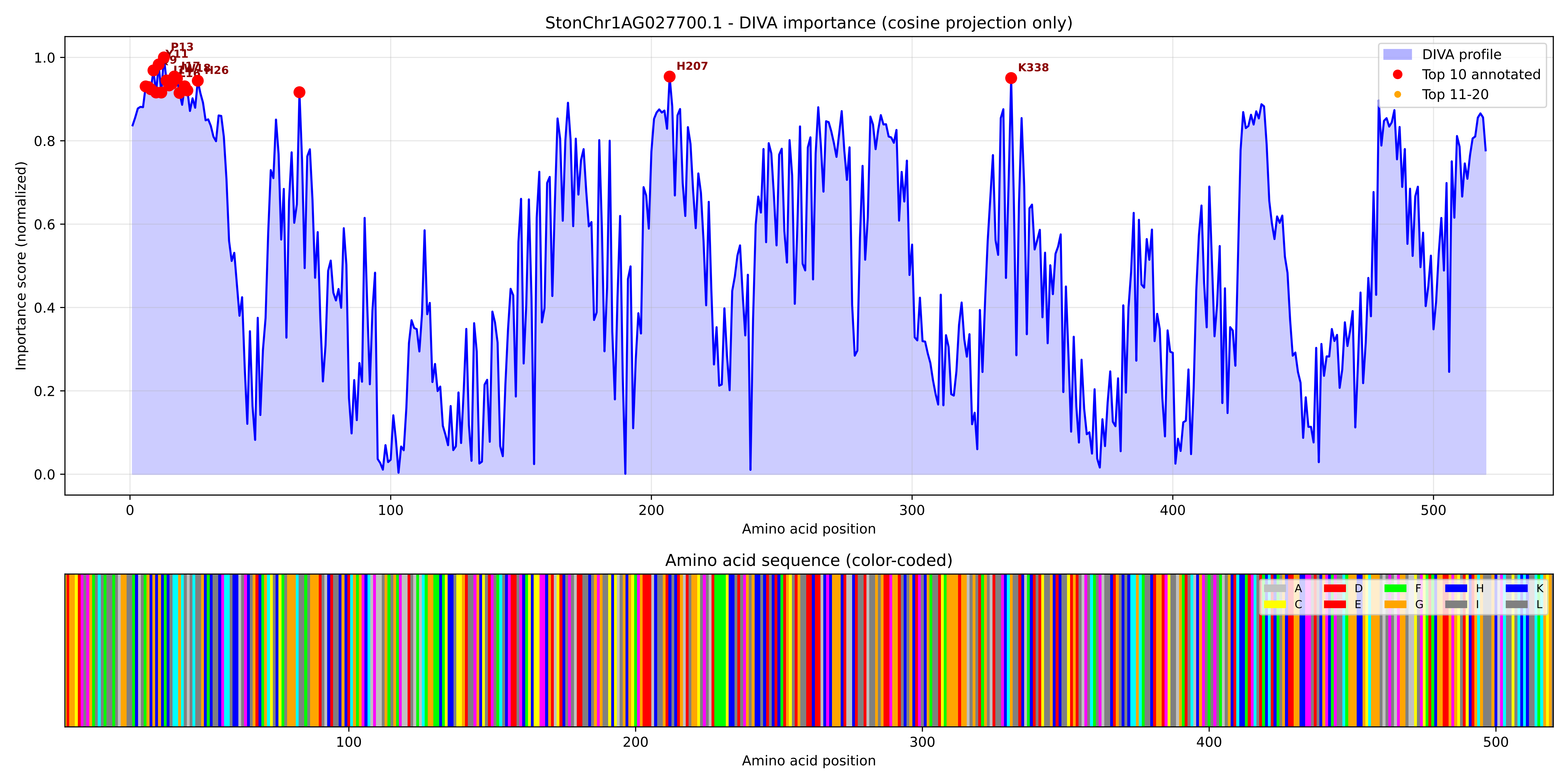

### StonChr1AG027700.1_importance.png

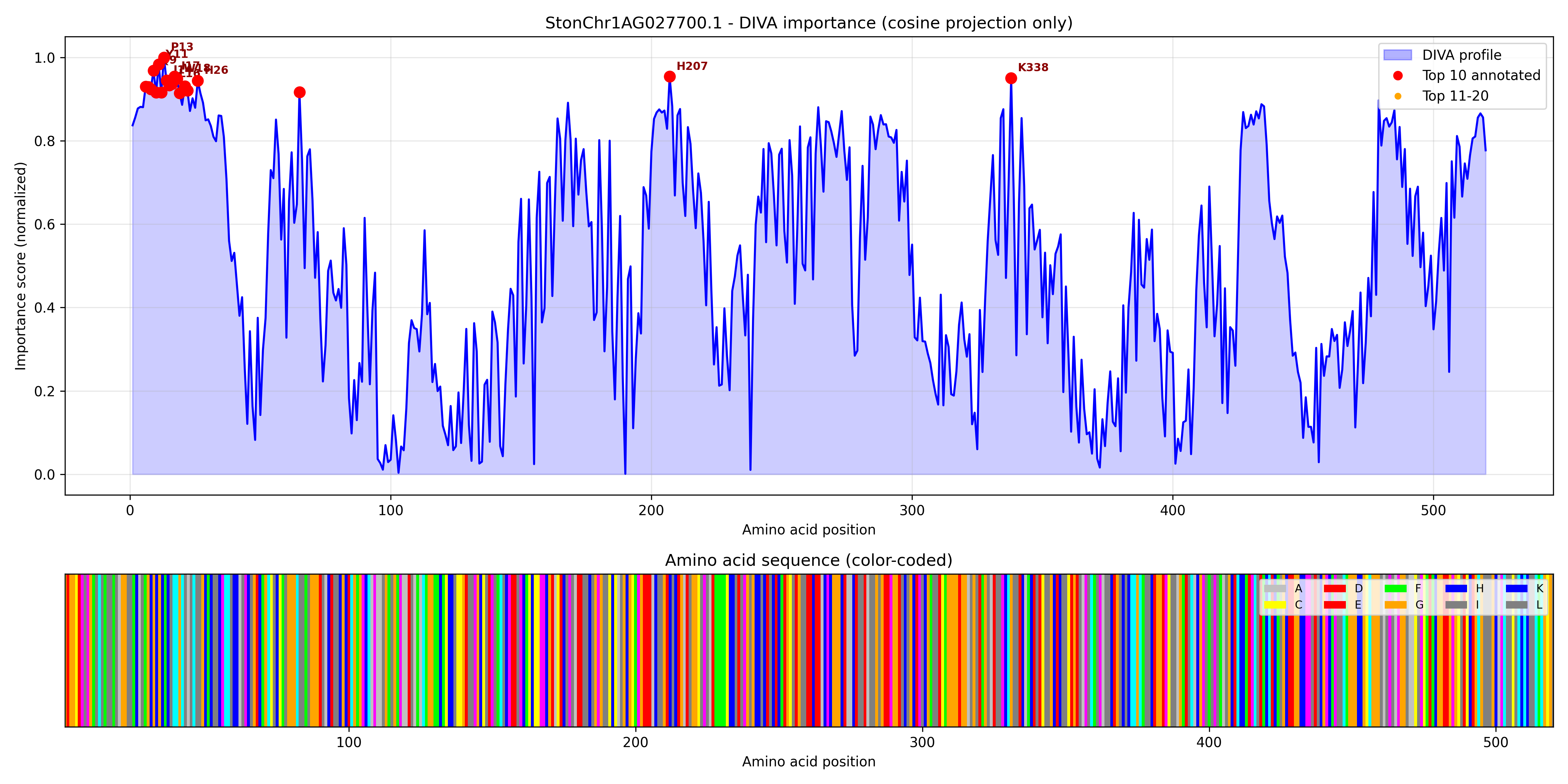

### StonChr1AG032010.1_importance.pdf

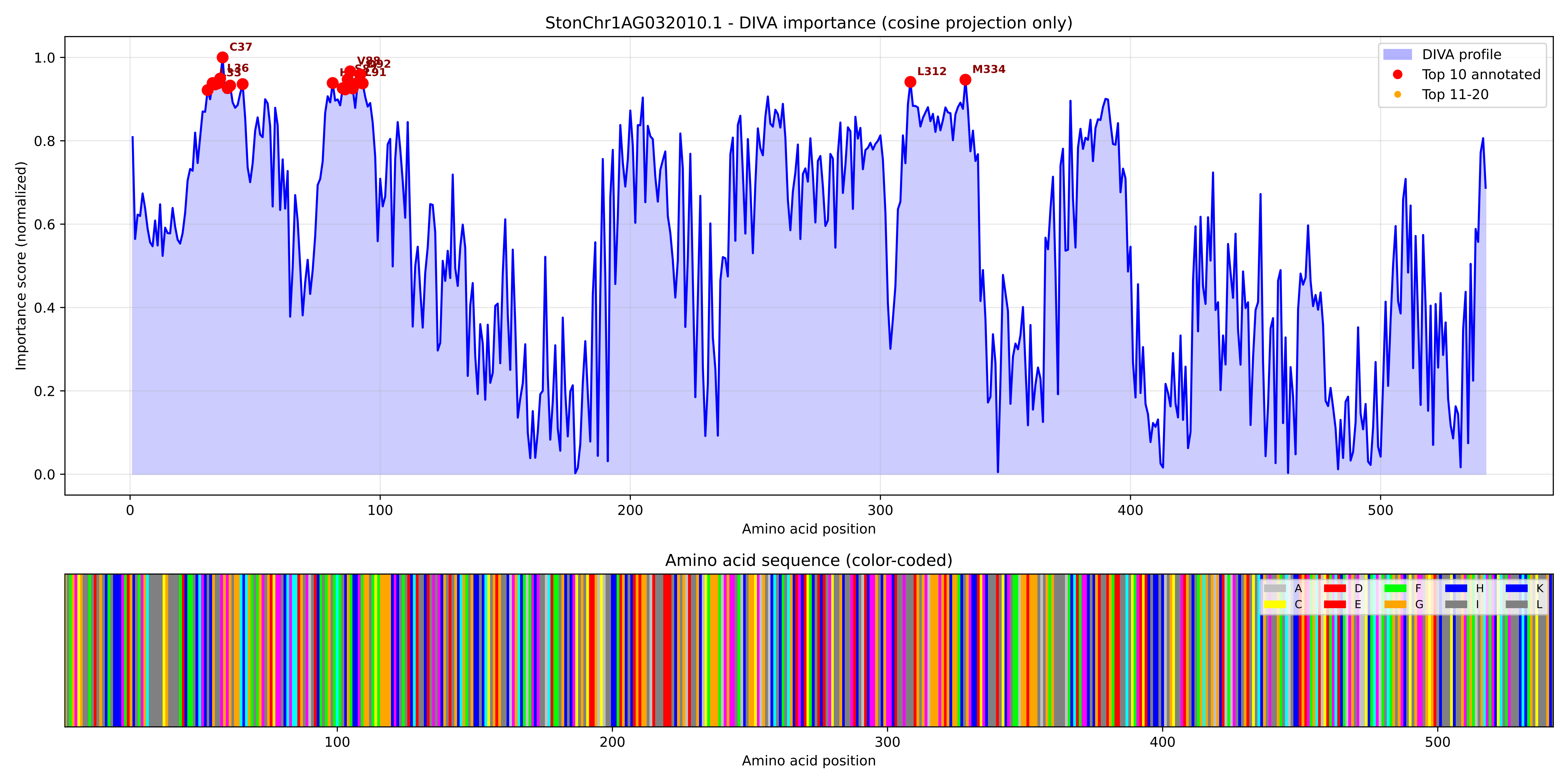

### StonChr1AG032010.1_importance.png

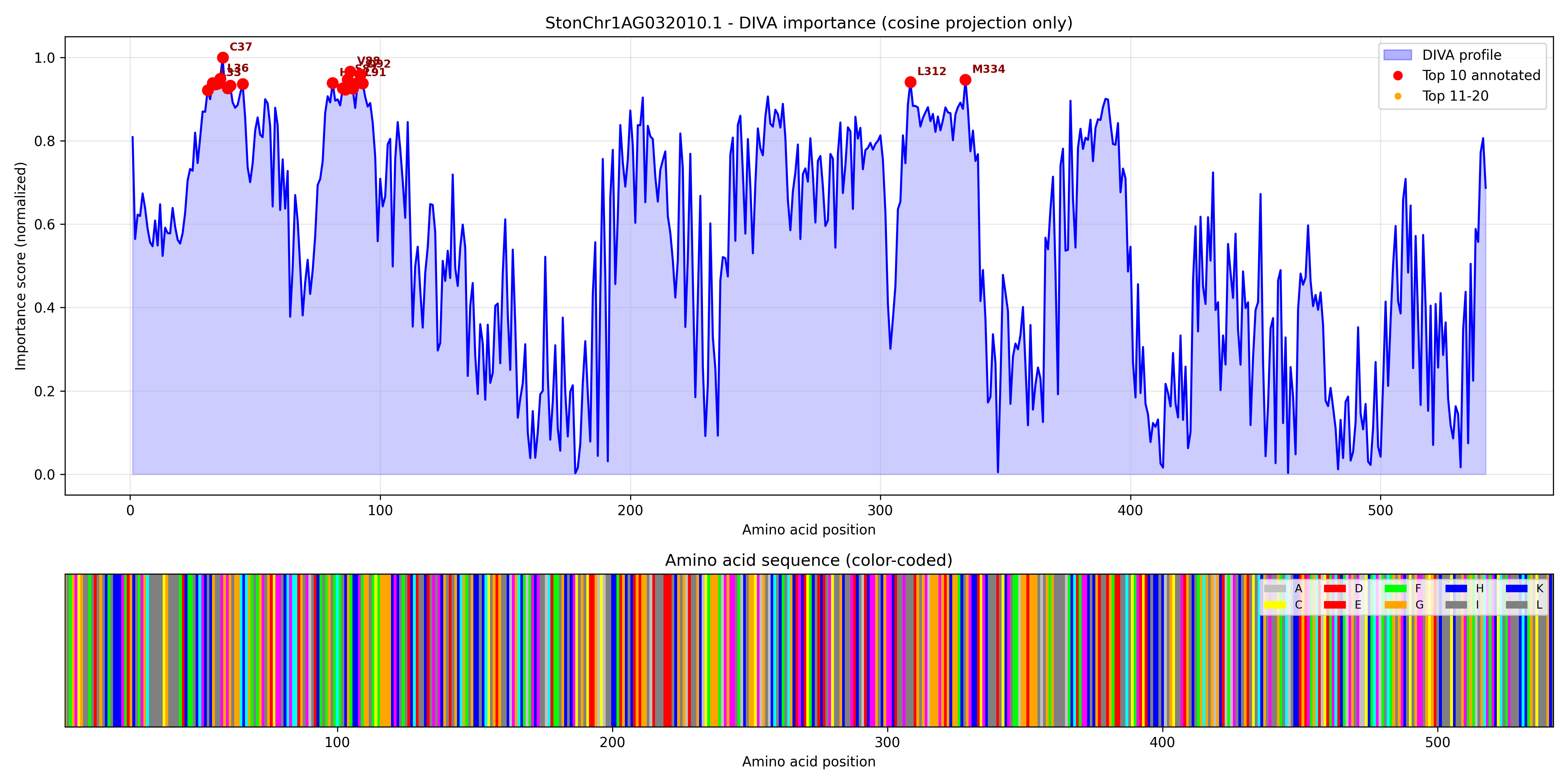

### StonChr1AG032040.1_importance.pdf

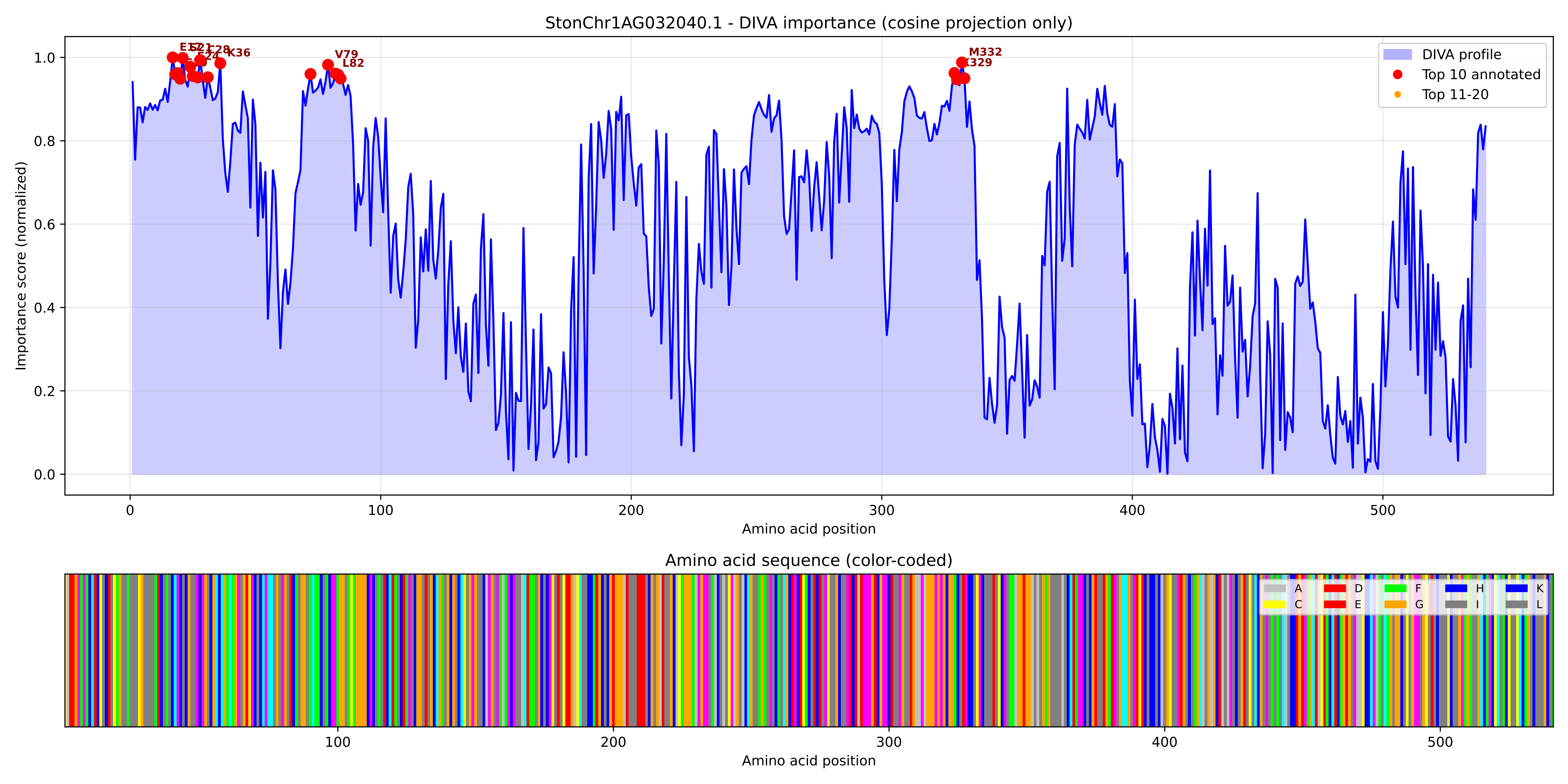

### StonChr1AG032040.1_importance.png

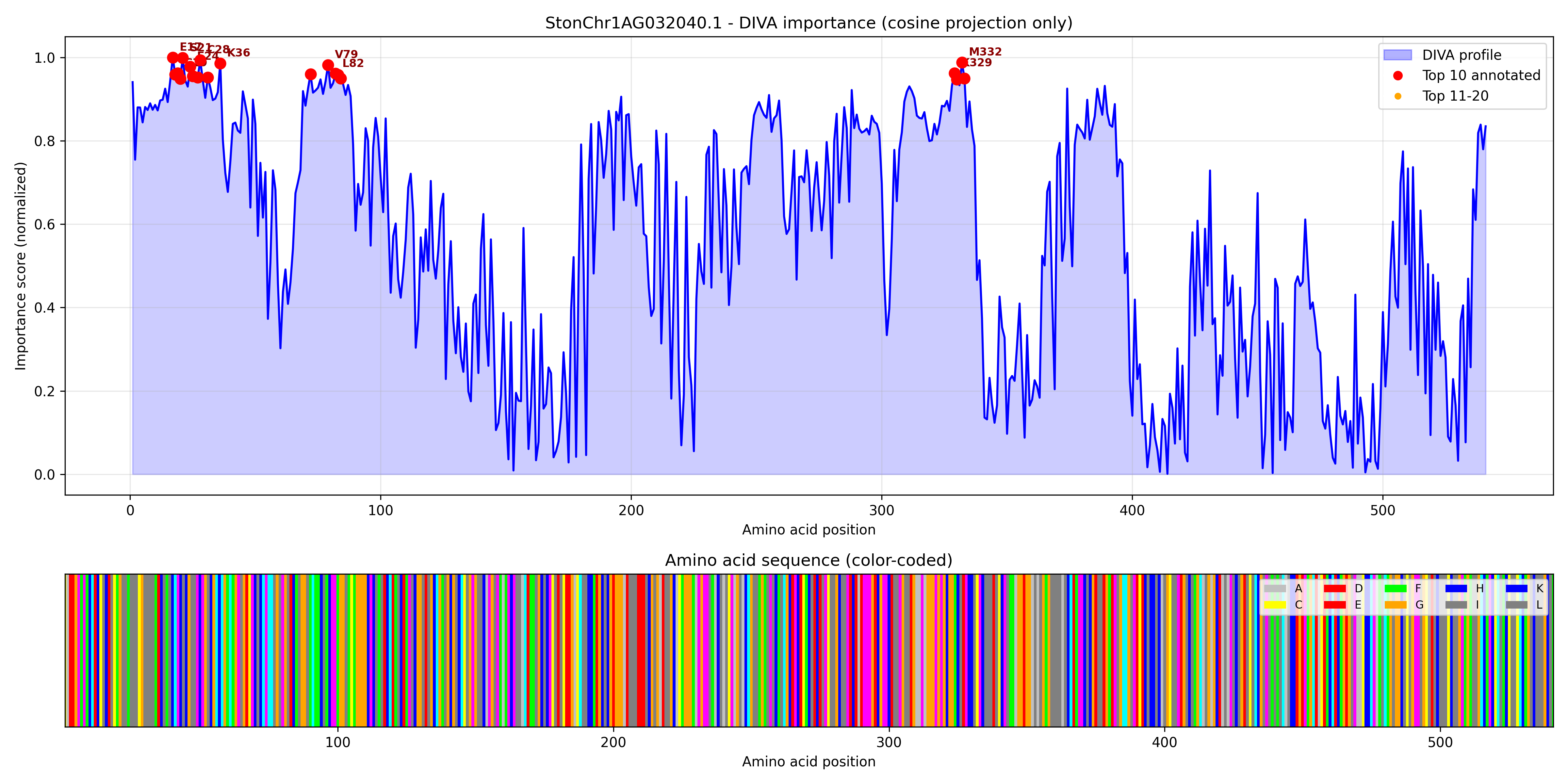

### StonChr1AG034590.1_importance.pdf

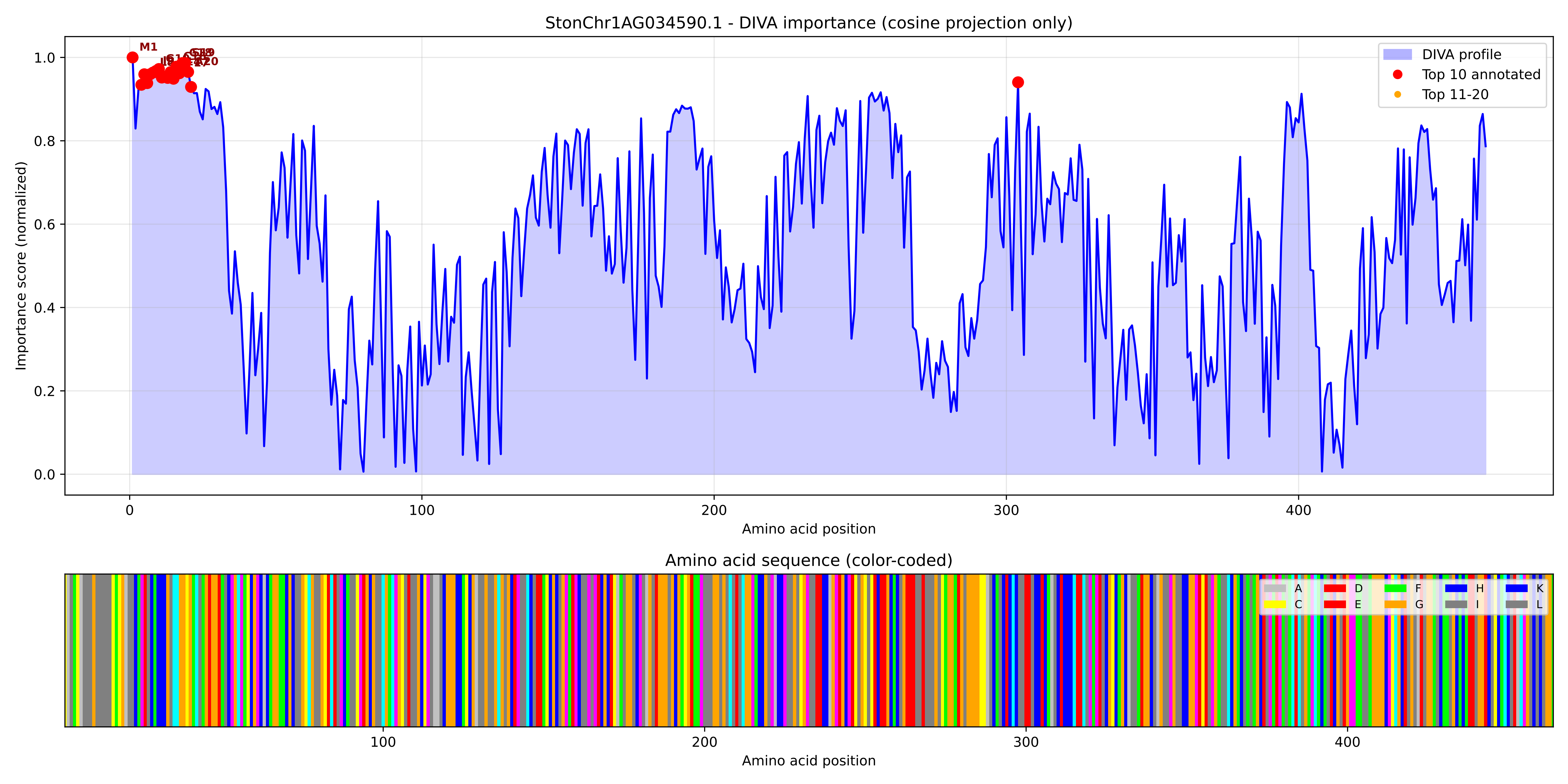

### StonChr1AG034590.1_importance.png

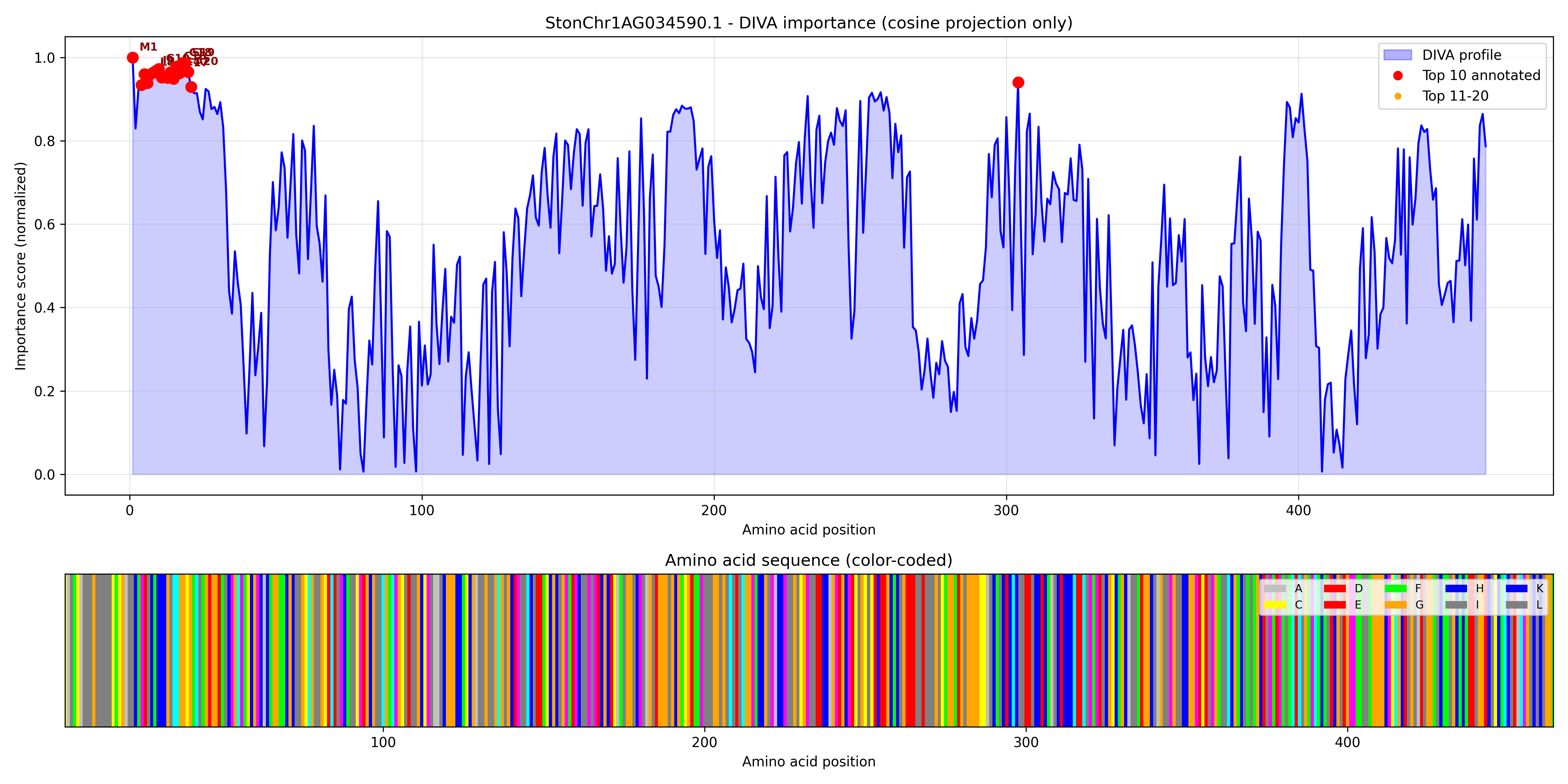

### StonChr1AG034840.1_importance.pdf

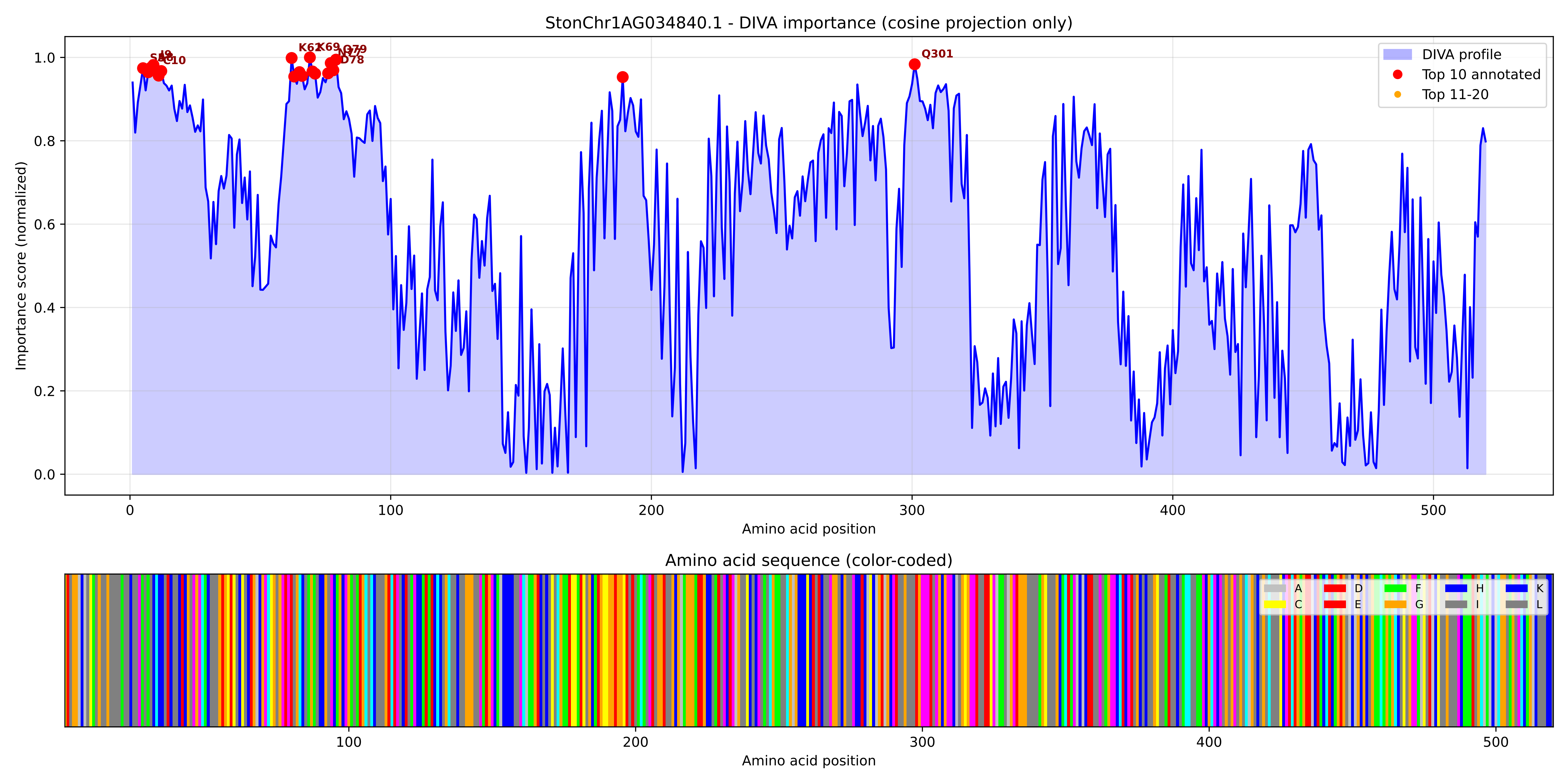

### StonChr1AG034840.1_importance.png

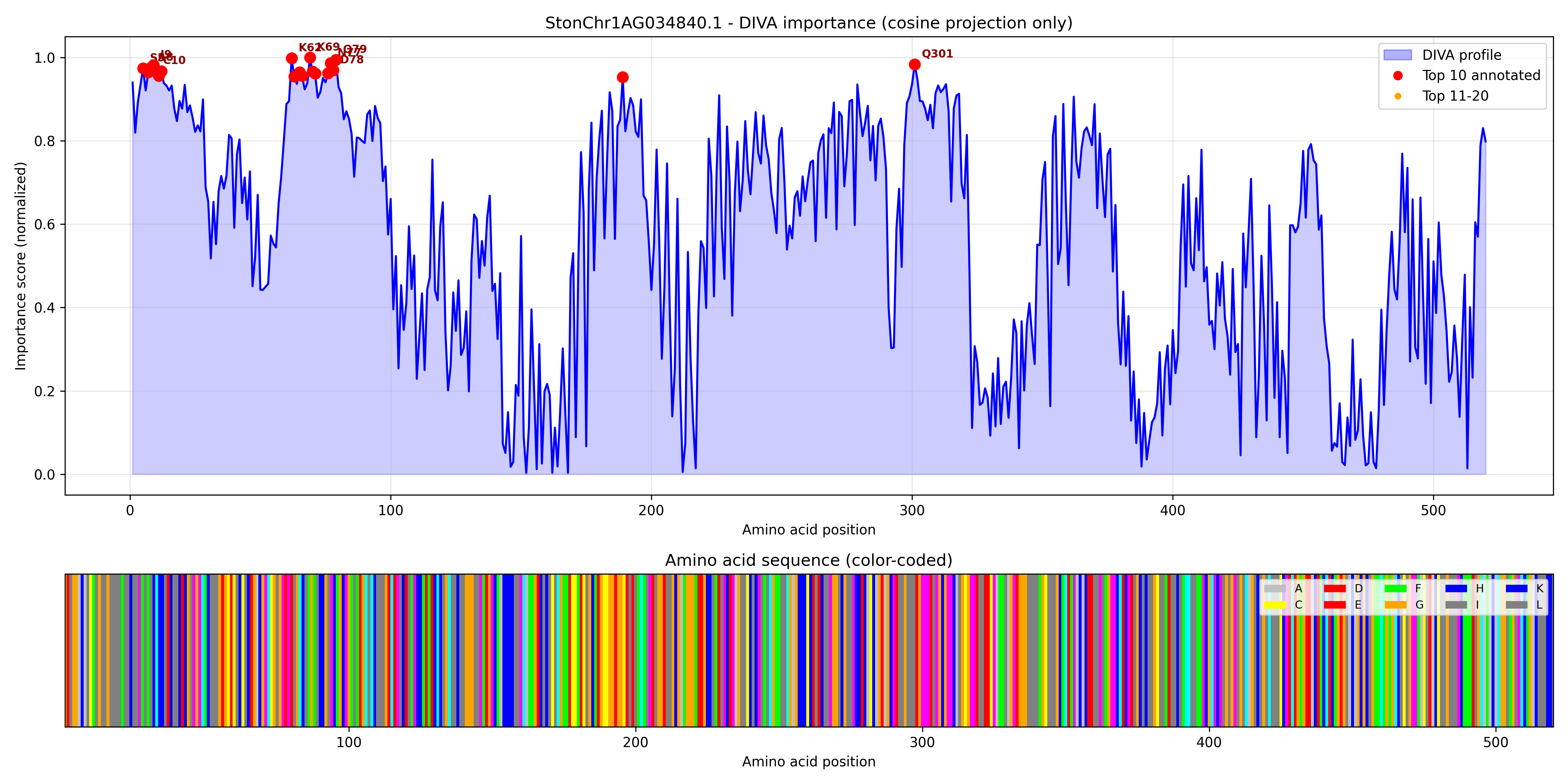

### StonChr1AG037120.1_importance.pdf

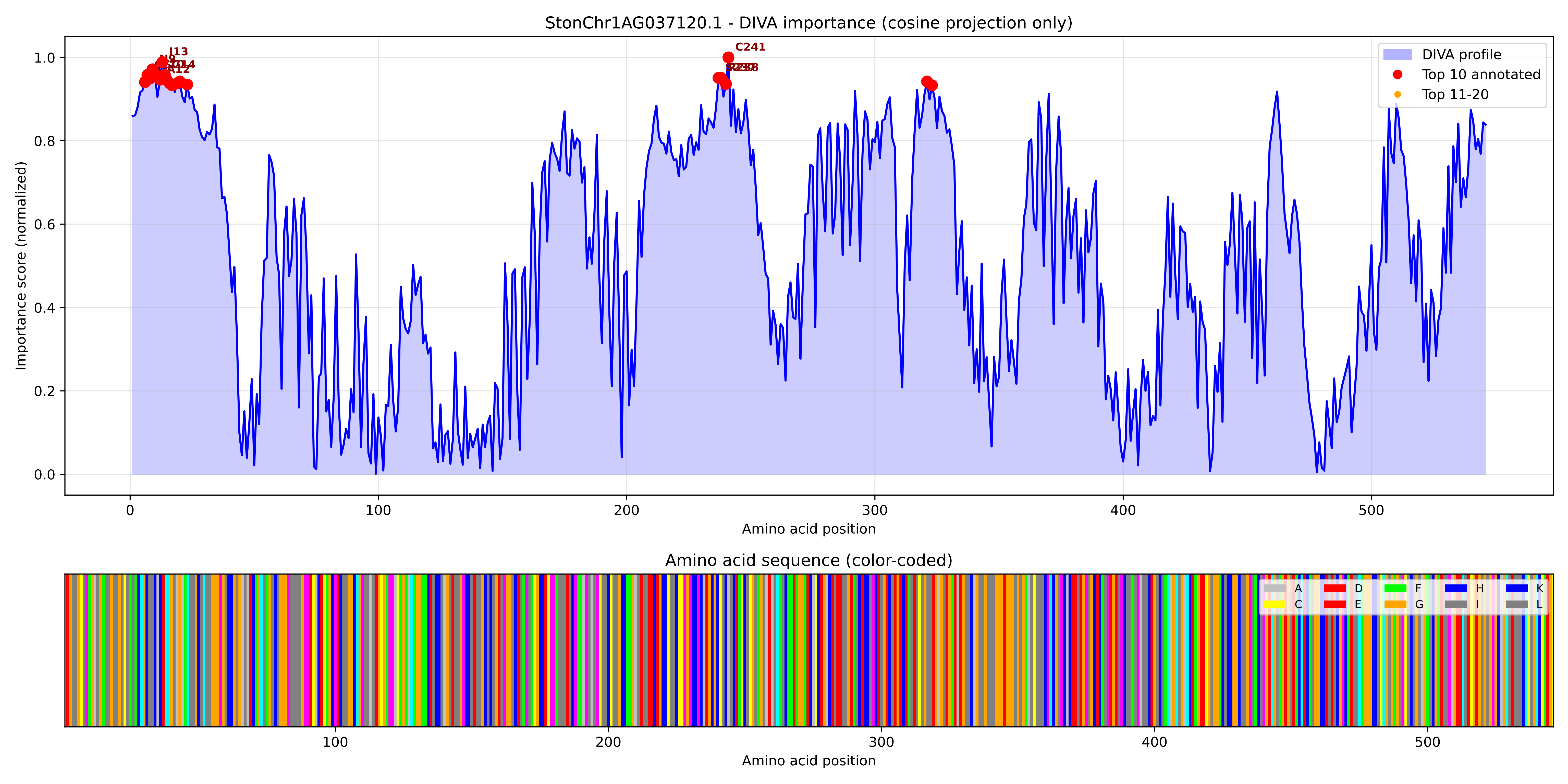

### StonChr1AG037120.1_importance.png

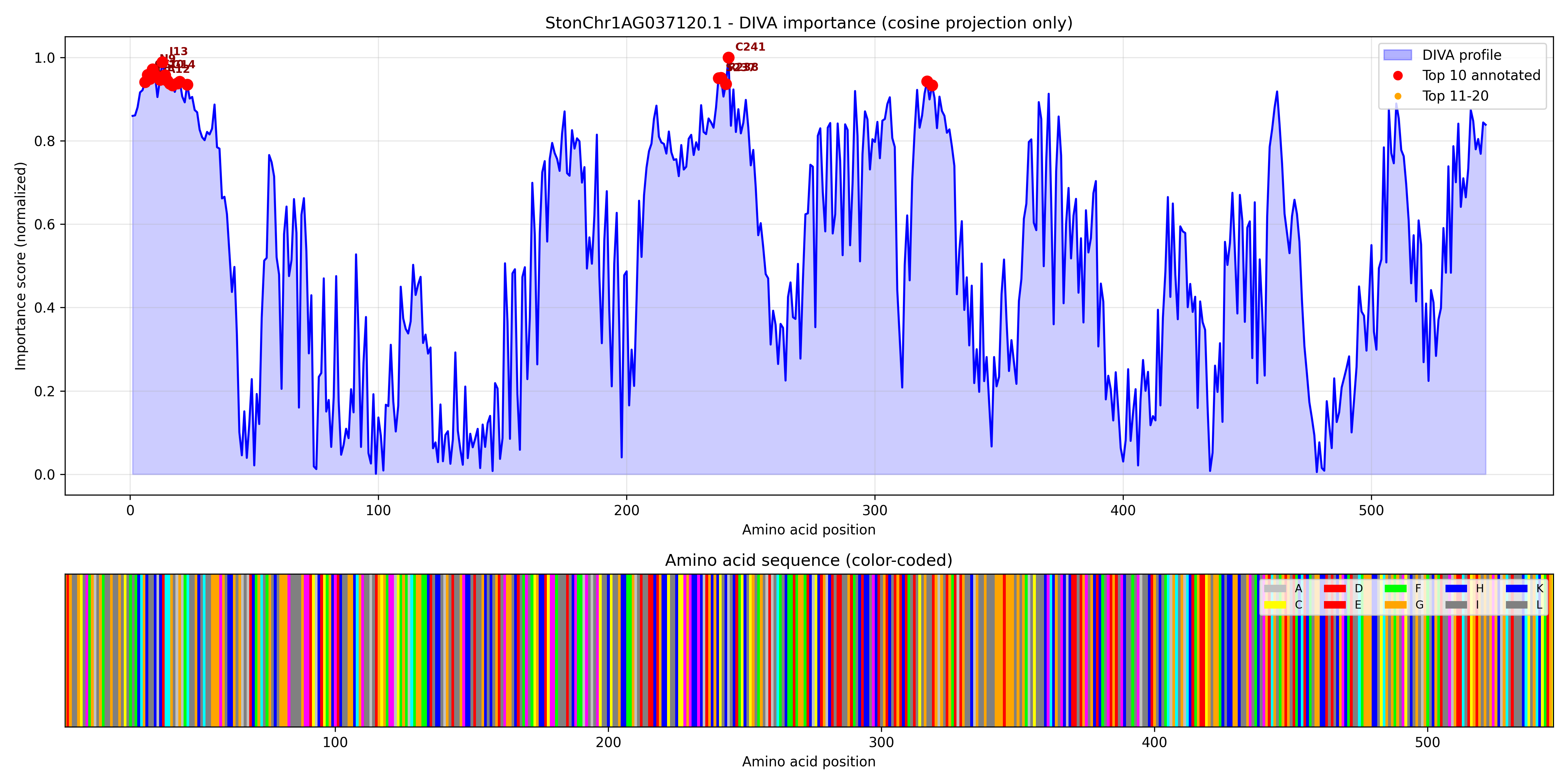

### StonChr1AG038340.1_importance.pdf

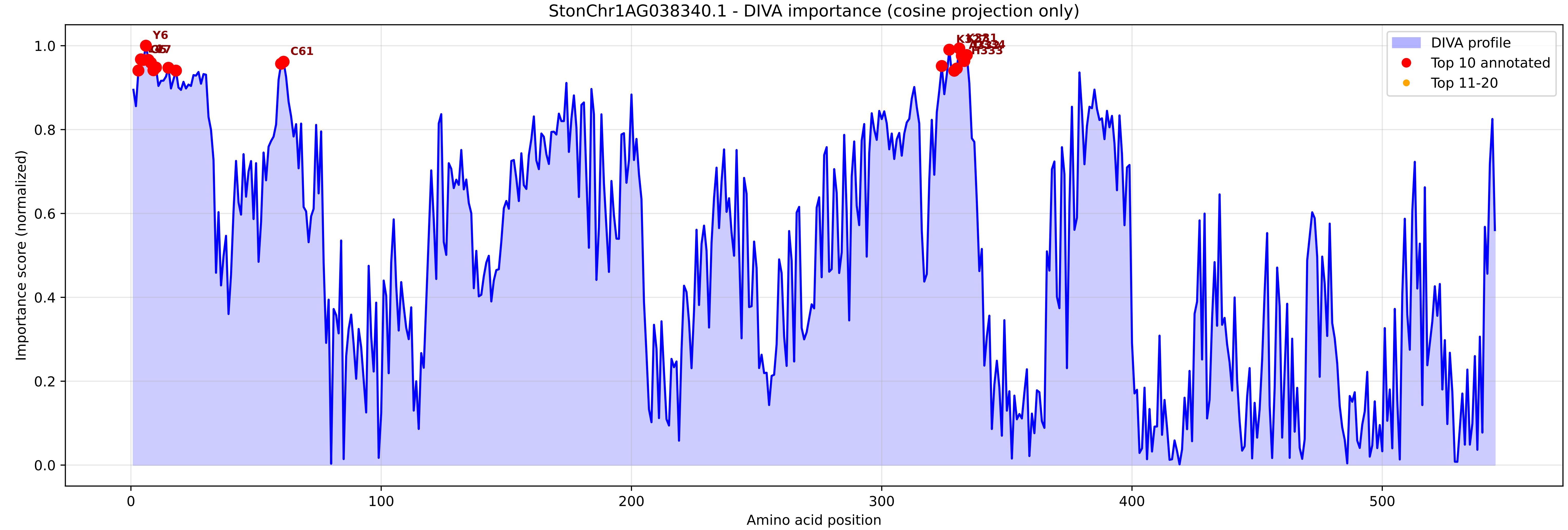

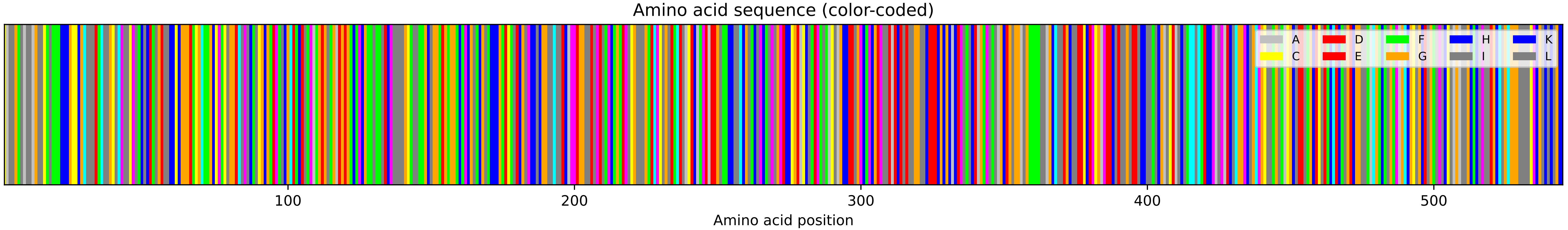

### StonChr1AG038340.1_importance.png

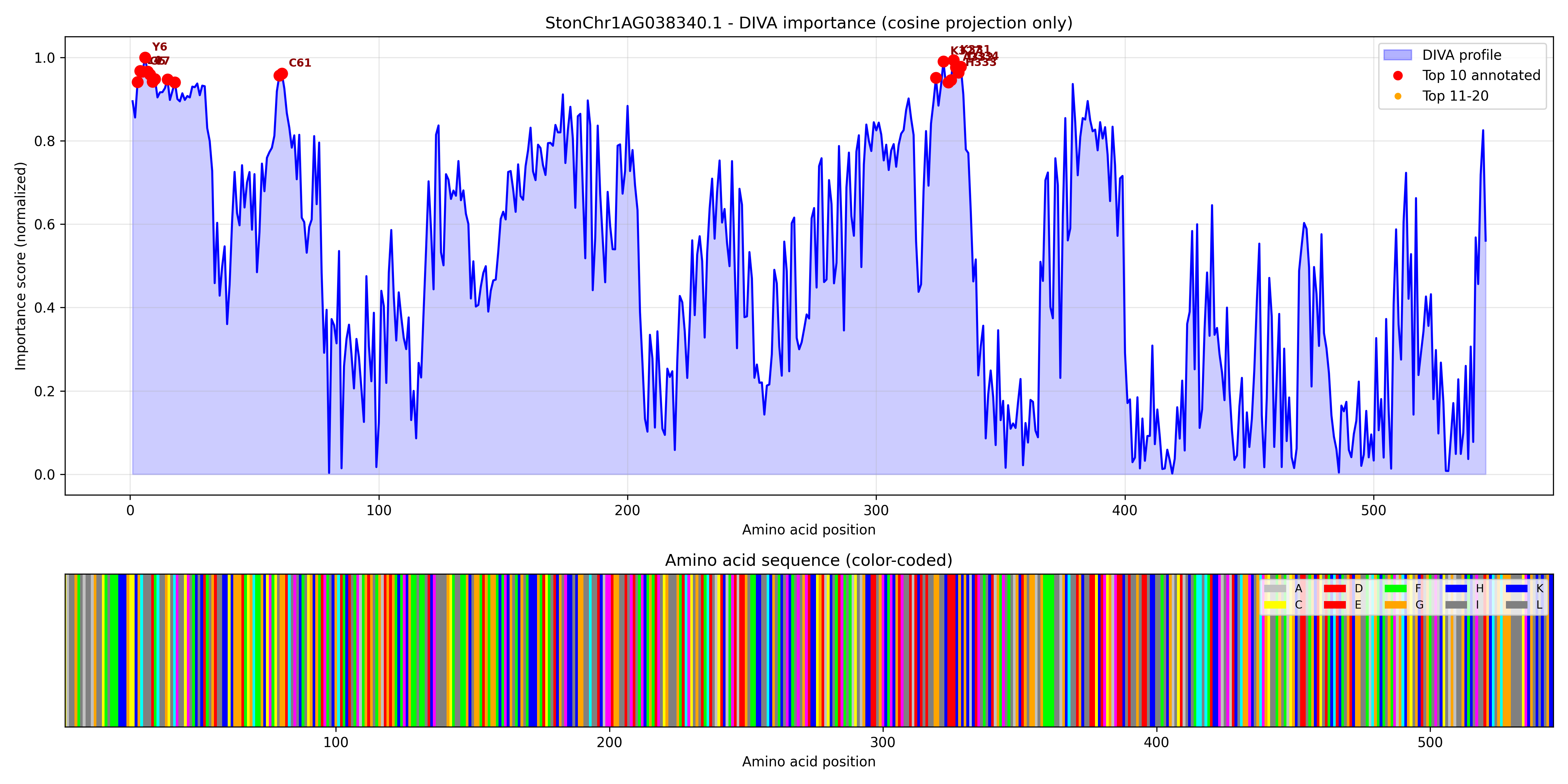

### StonChr1BG000310.1_importance.pdf

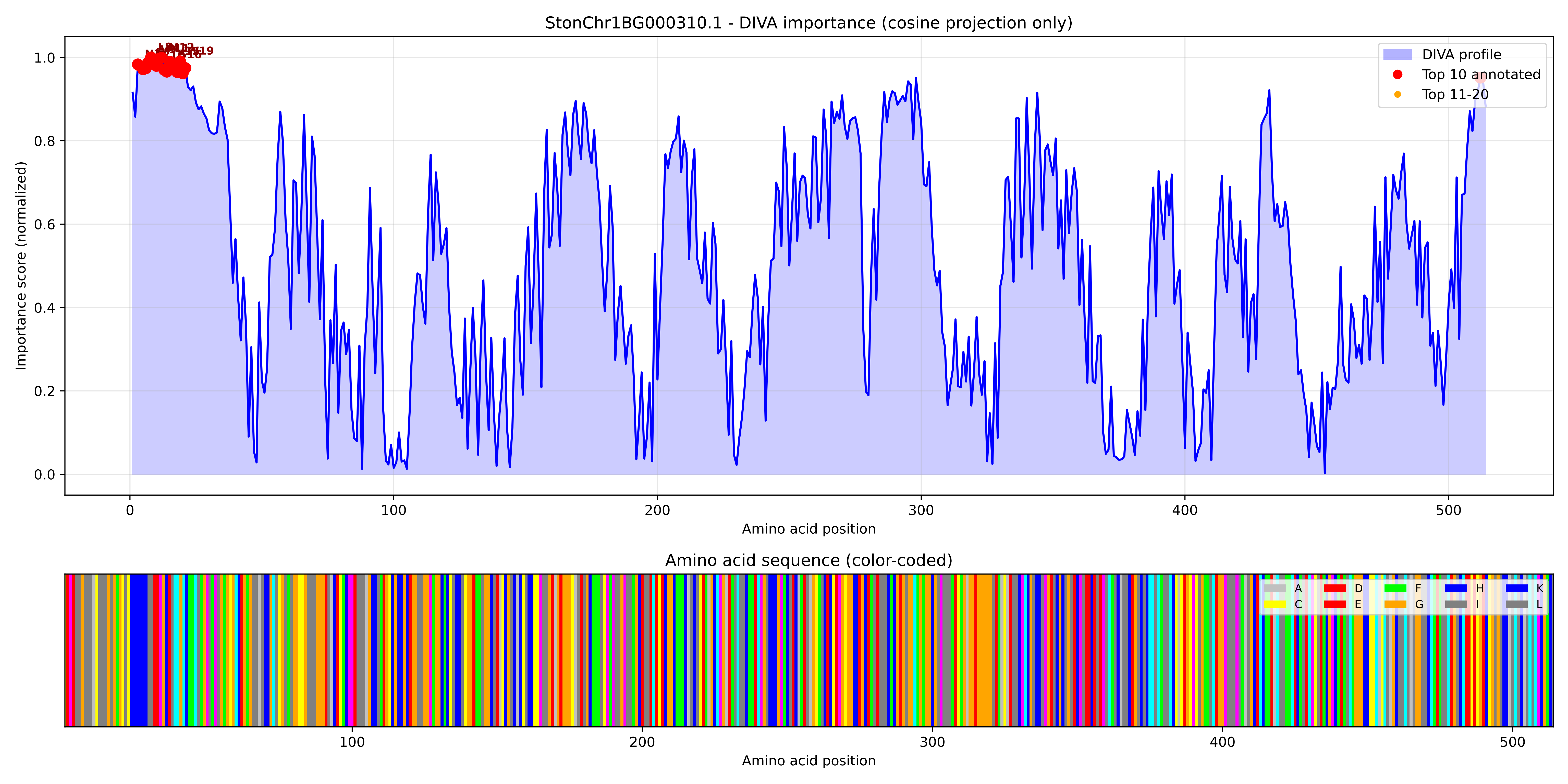

### StonChr1BG000310.1_importance.png

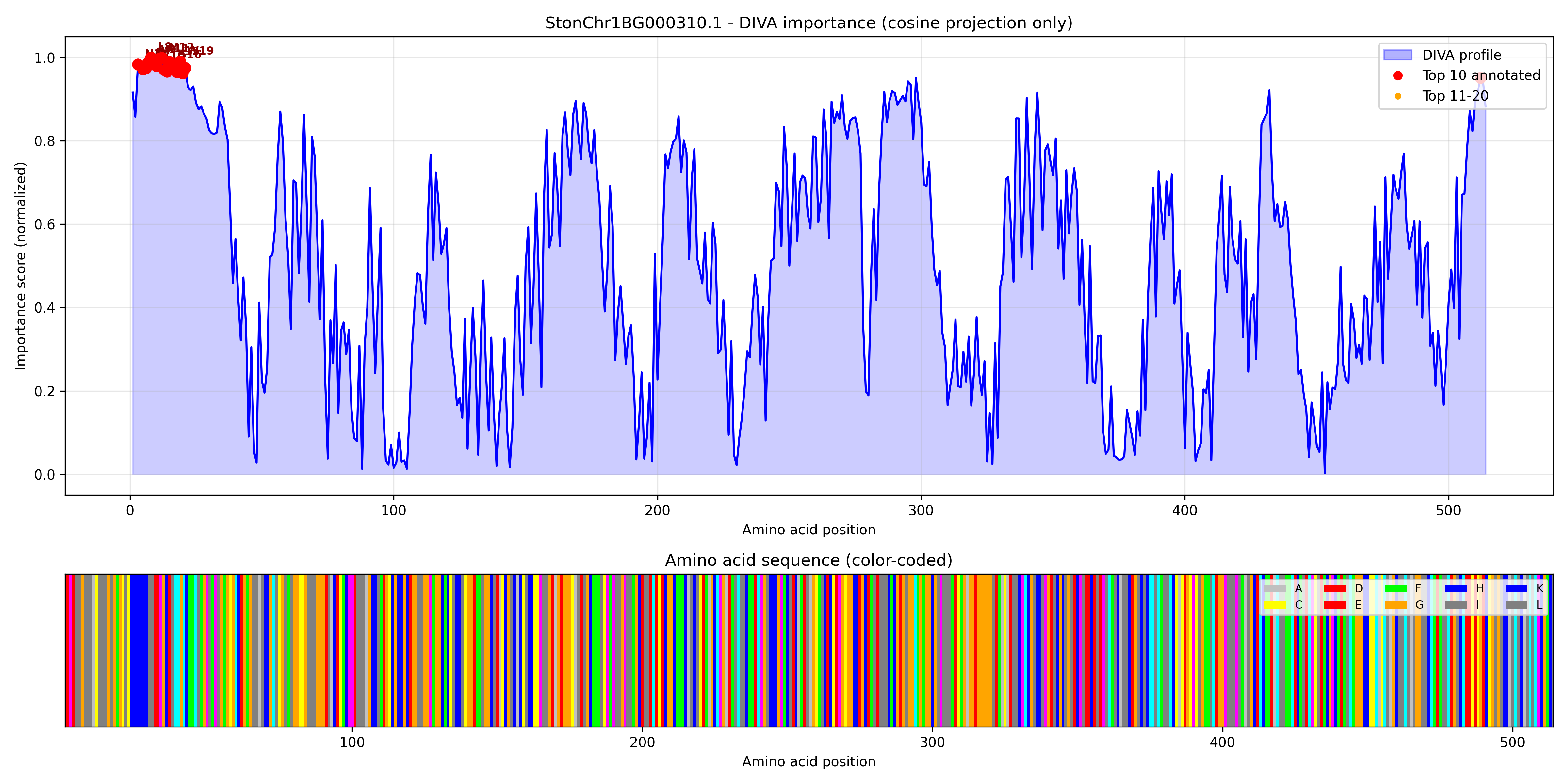

### StonChr1BG002840.1_importance.pdf

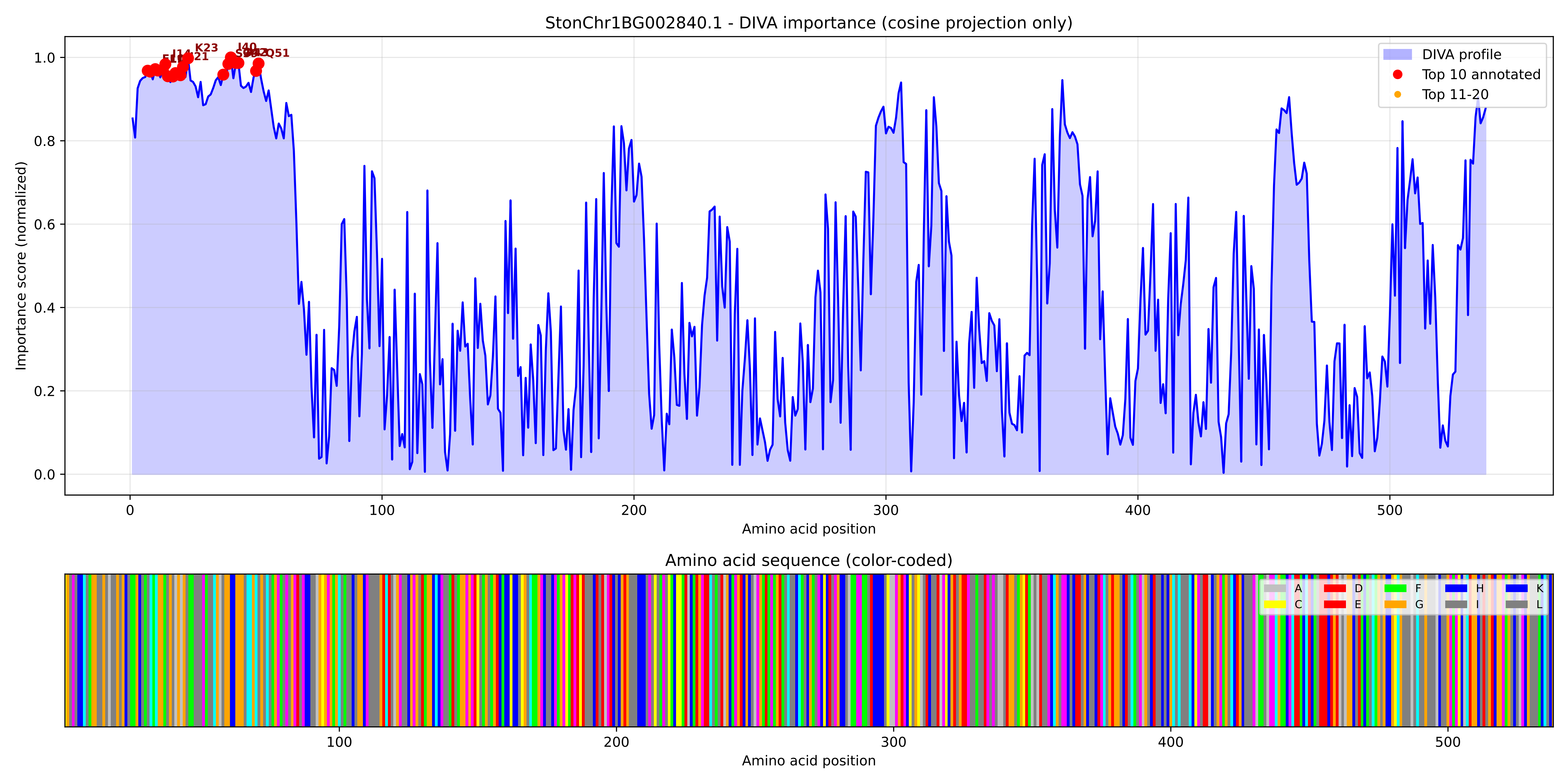

### StonChr1BG002840.1_importance.png

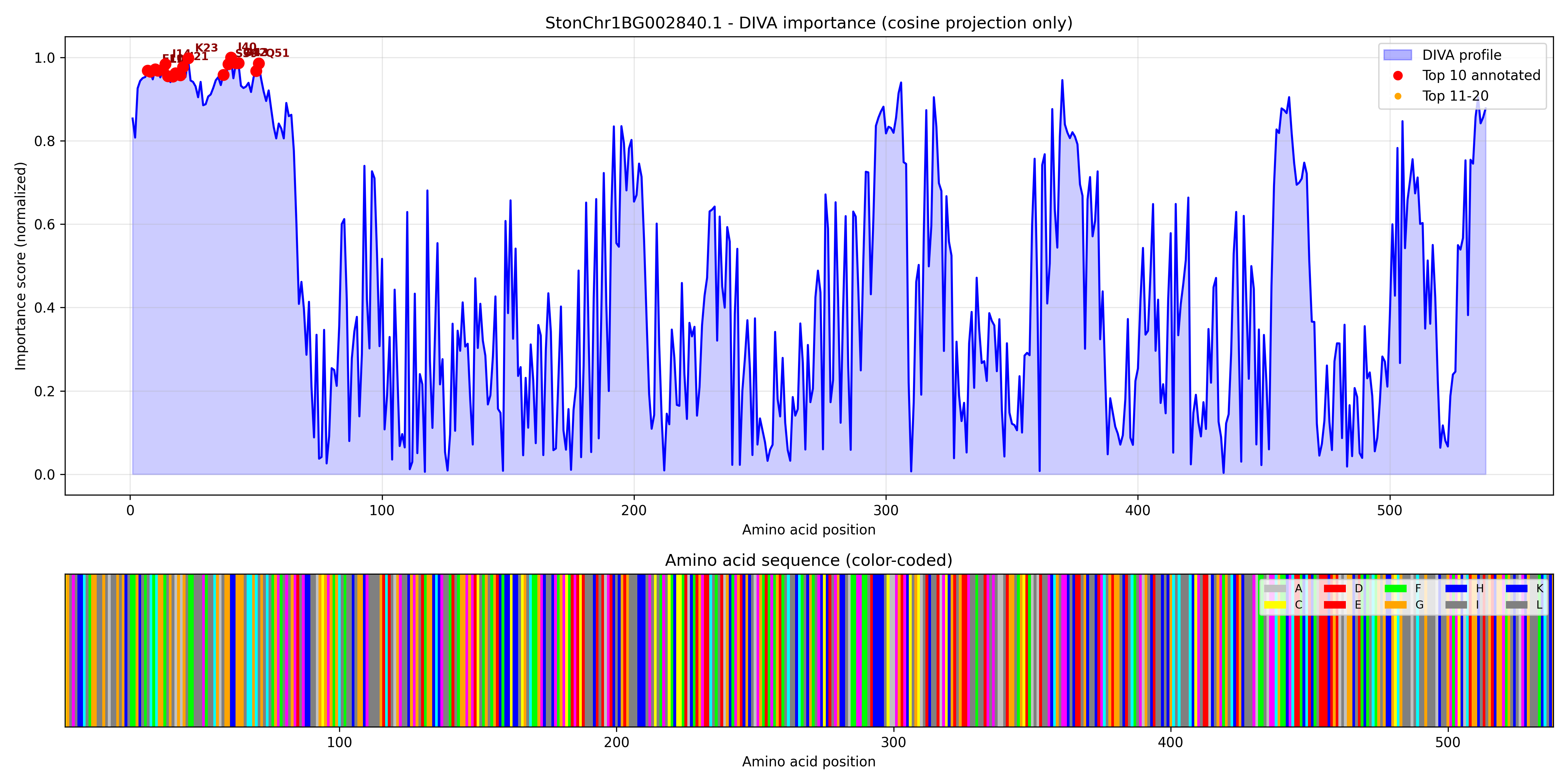

### StonChr1BG008910.1_importance.pdf

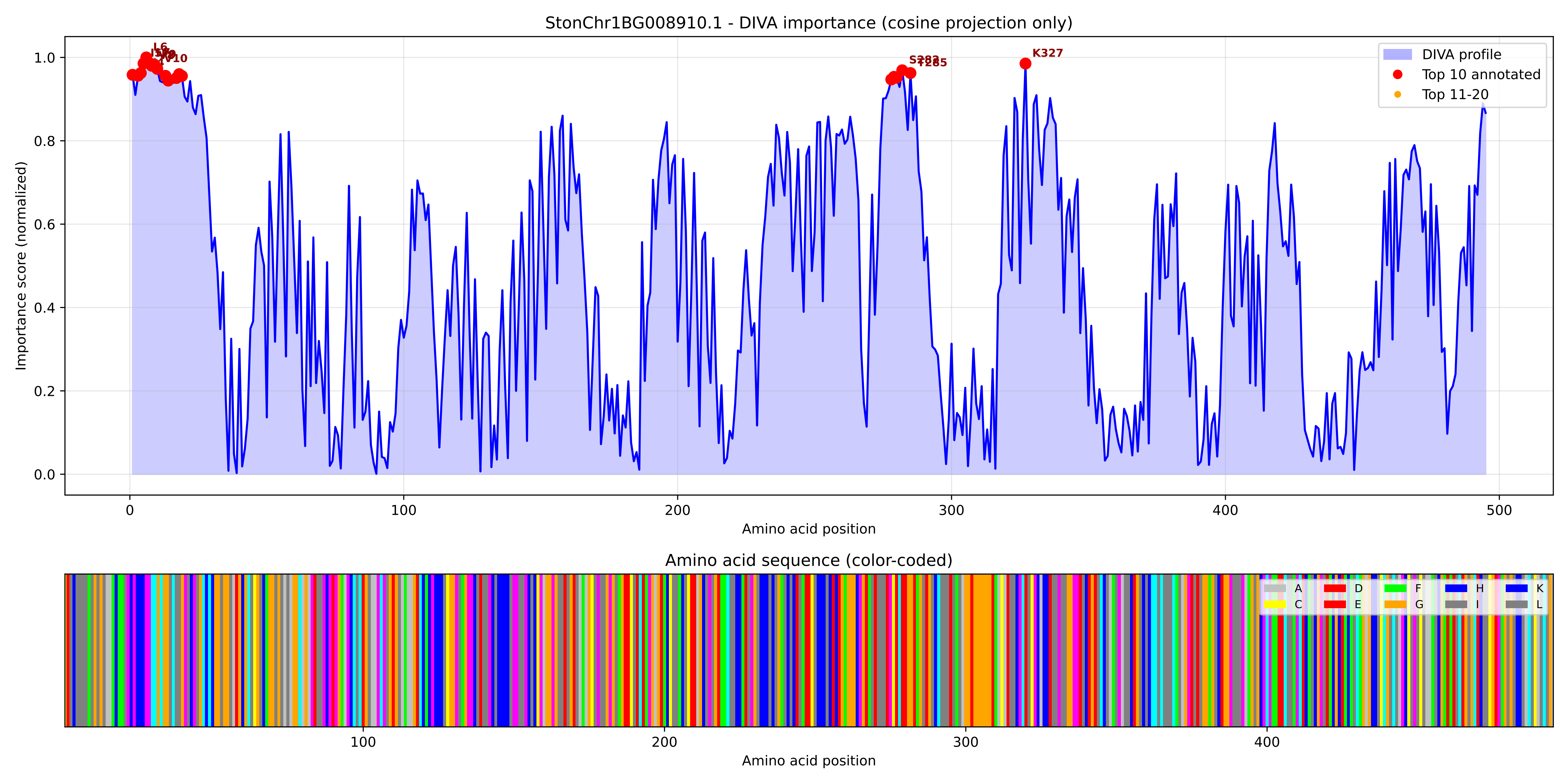

### StonChr1BG008910.1_importance.png

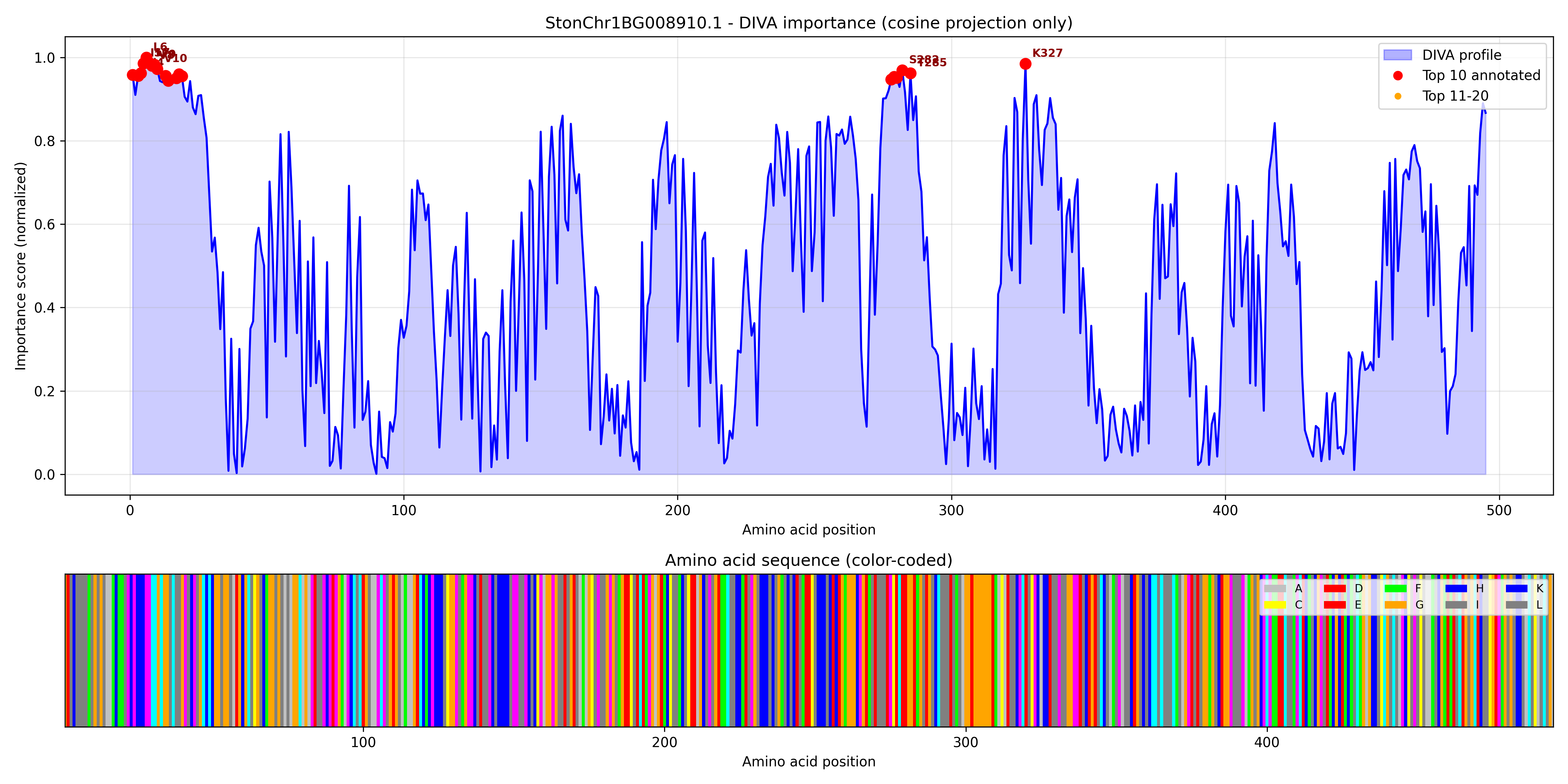

### StonChr1BG010150.1_importance.pdf

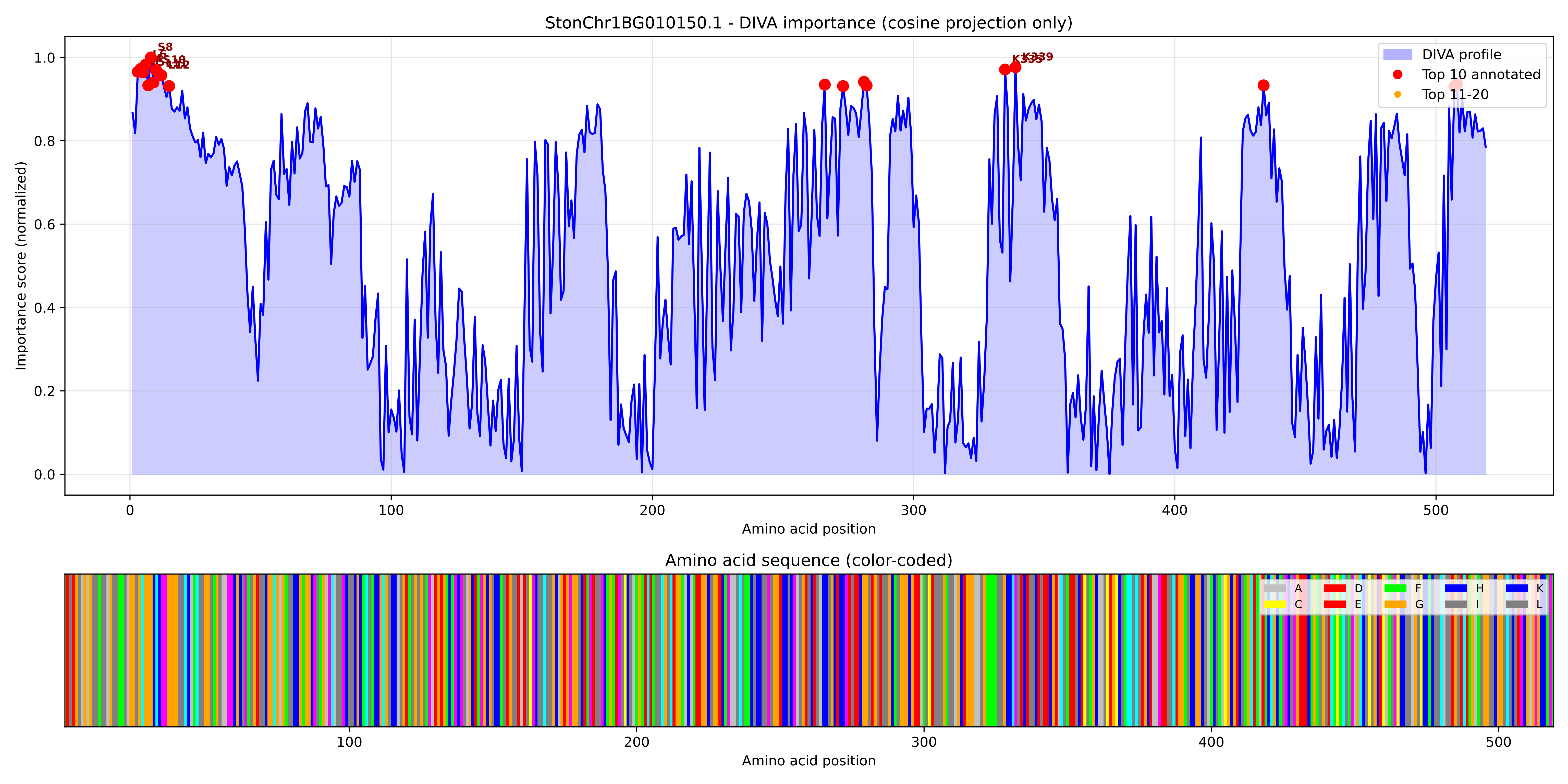

### StonChr1BG010150.1_importance.png

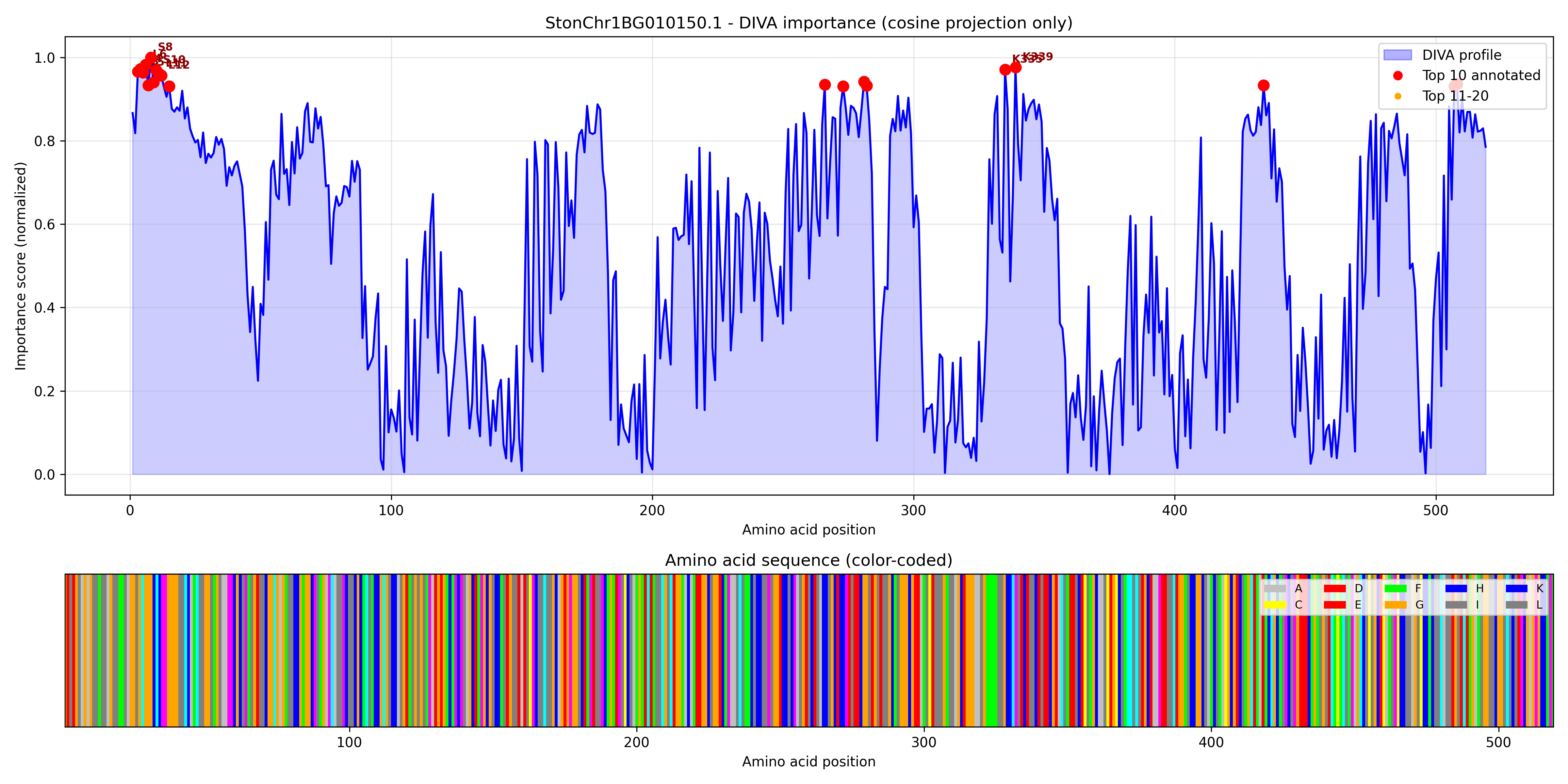

### StonChr1BG020360.1_importance.pdf

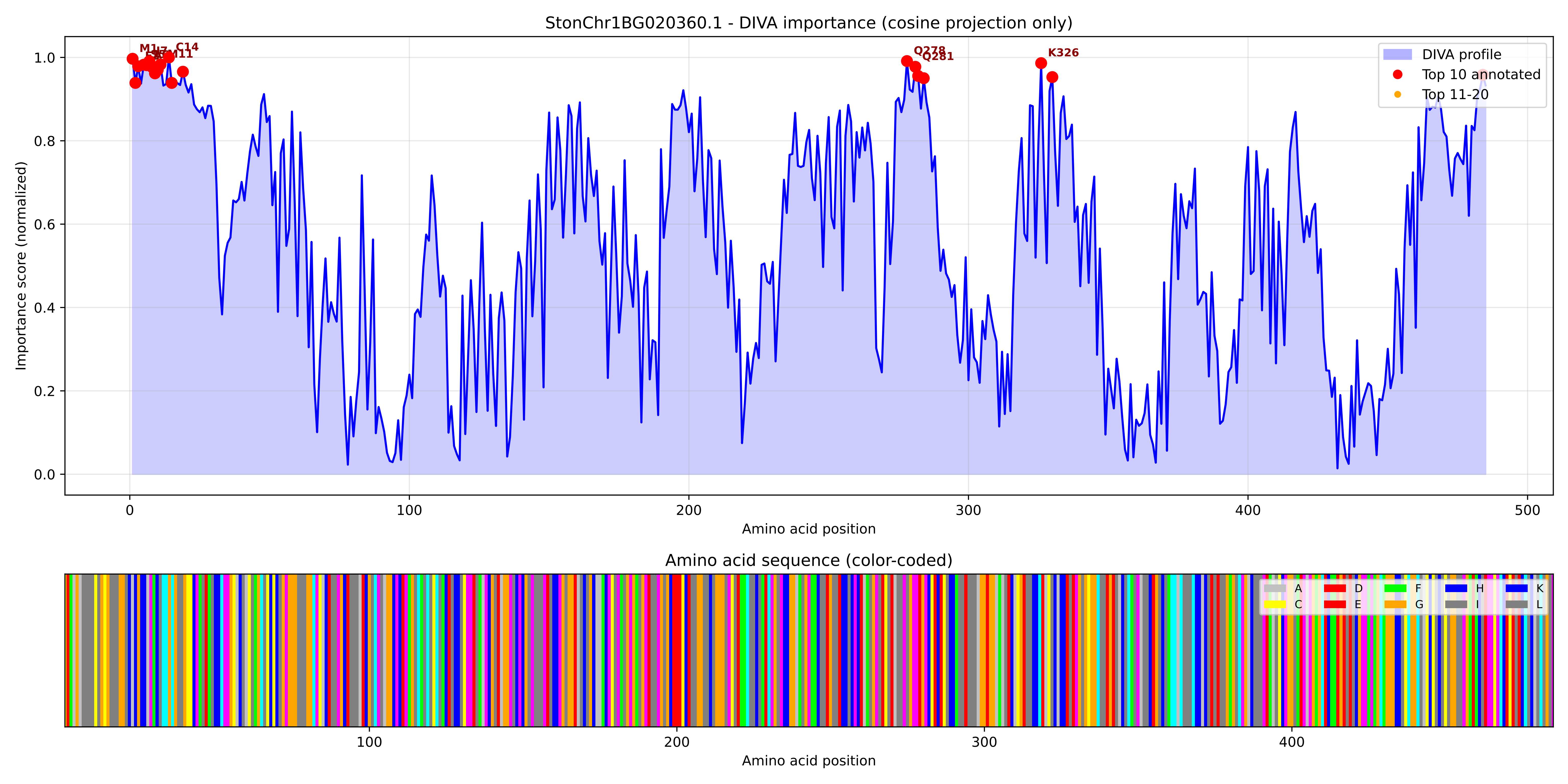

### StonChr1BG020360.1_importance.png

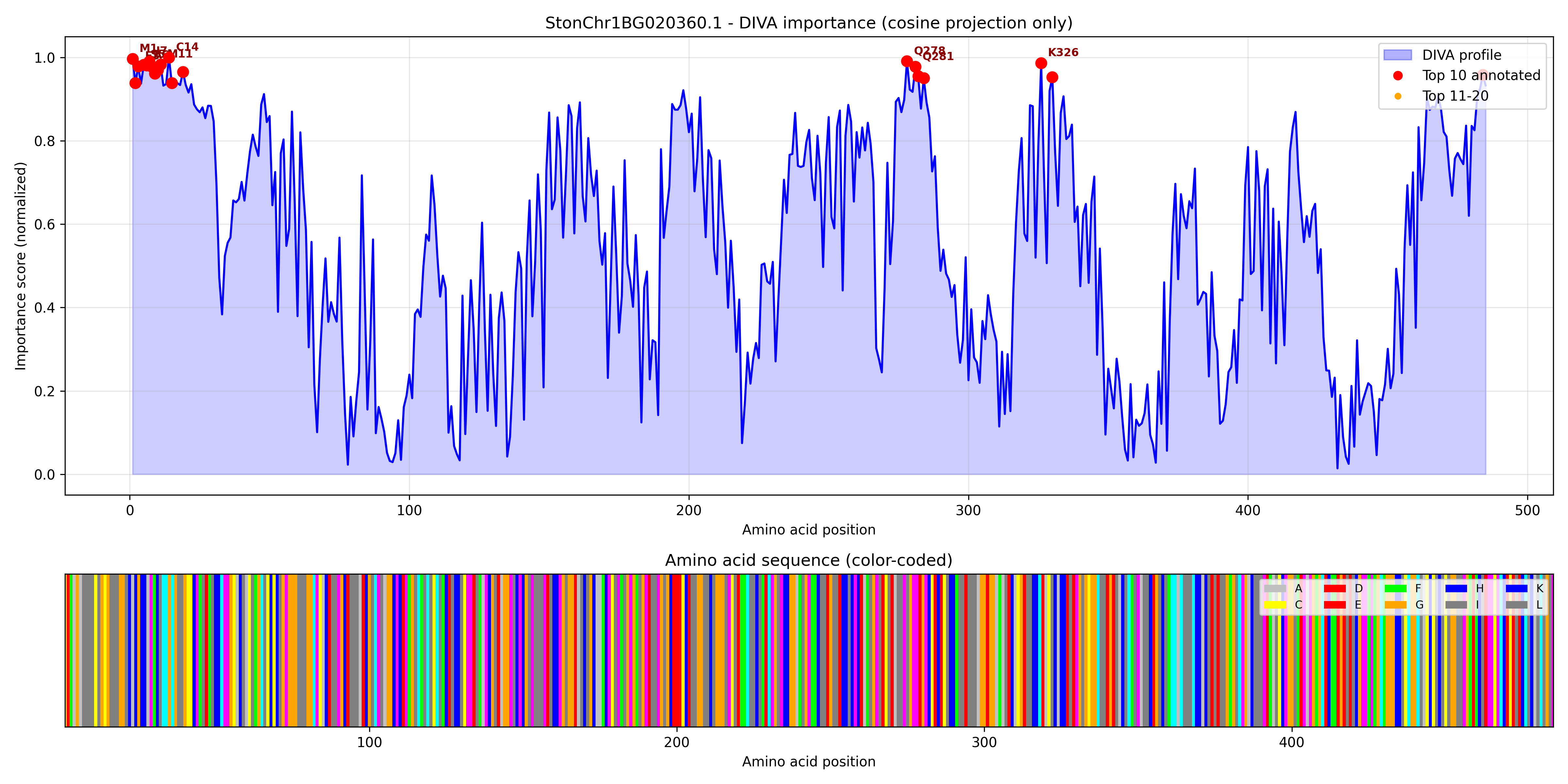

### StonChr1BG020600.1_importance.pdf

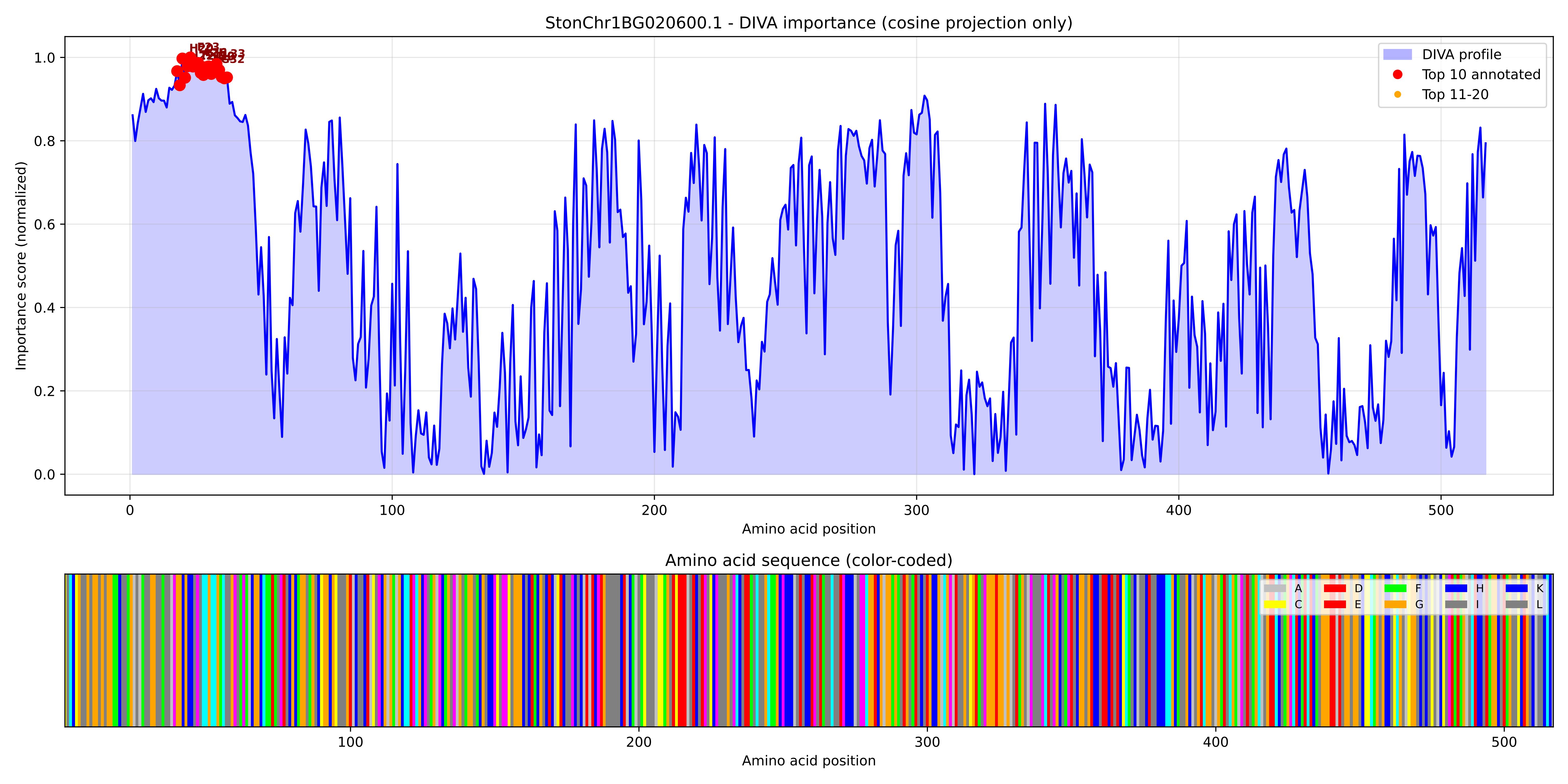

### StonChr1BG020600.1_importance.png

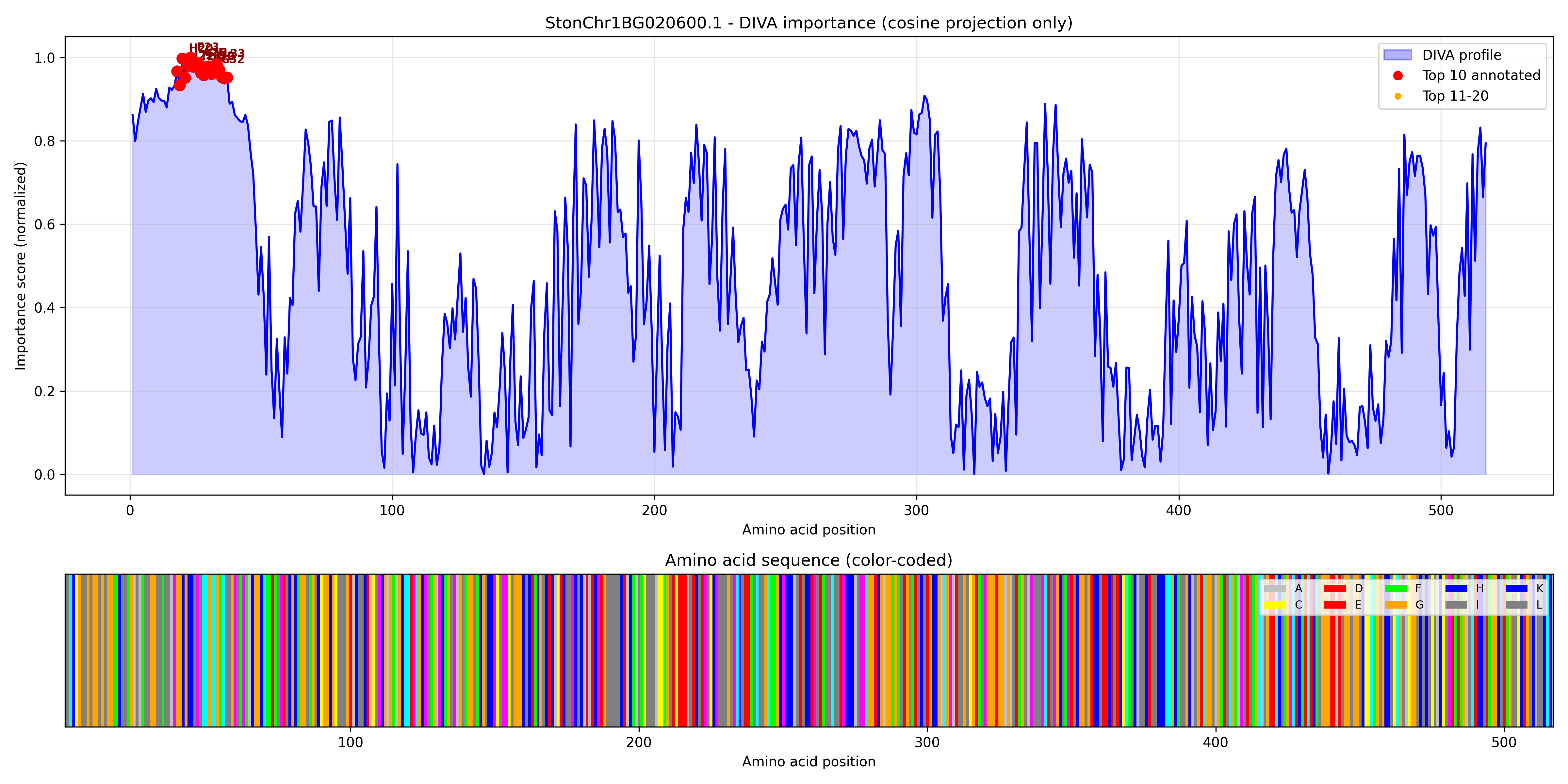

### StonChr1BG020720.1_importance.pdf

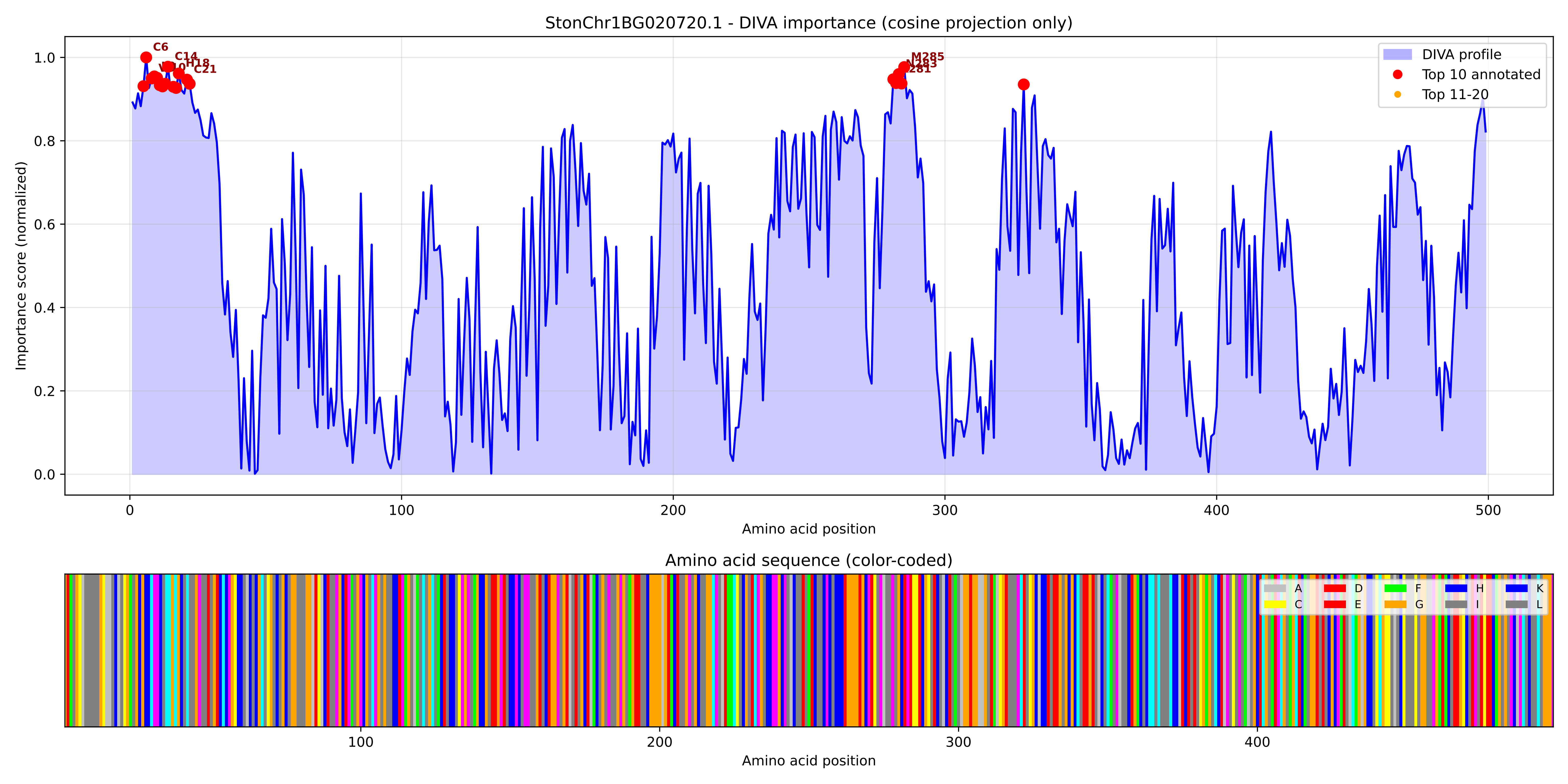

### StonChr1BG020720.1_importance.png

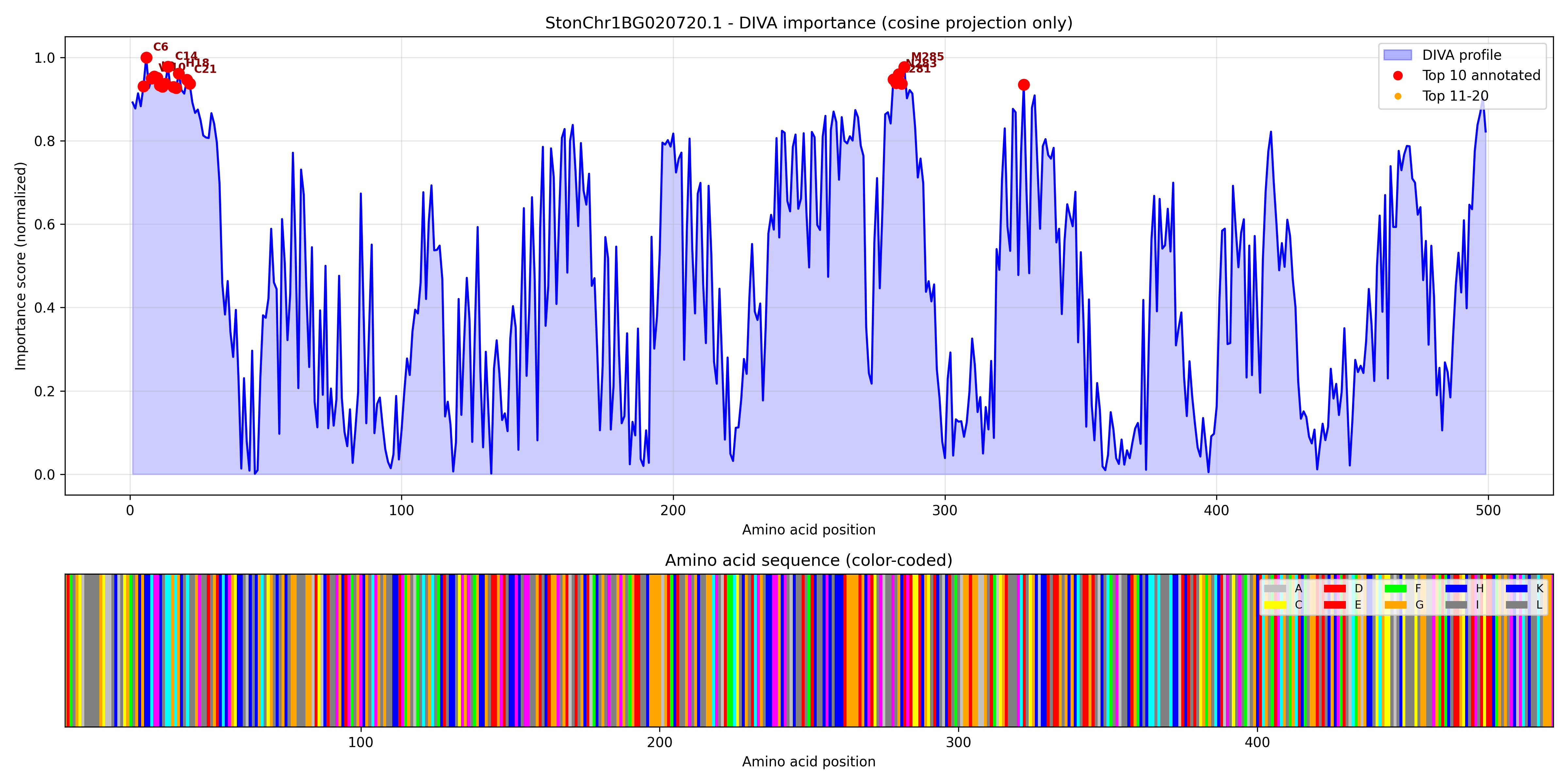

### StonChr1BG024990.1_importance.pdf

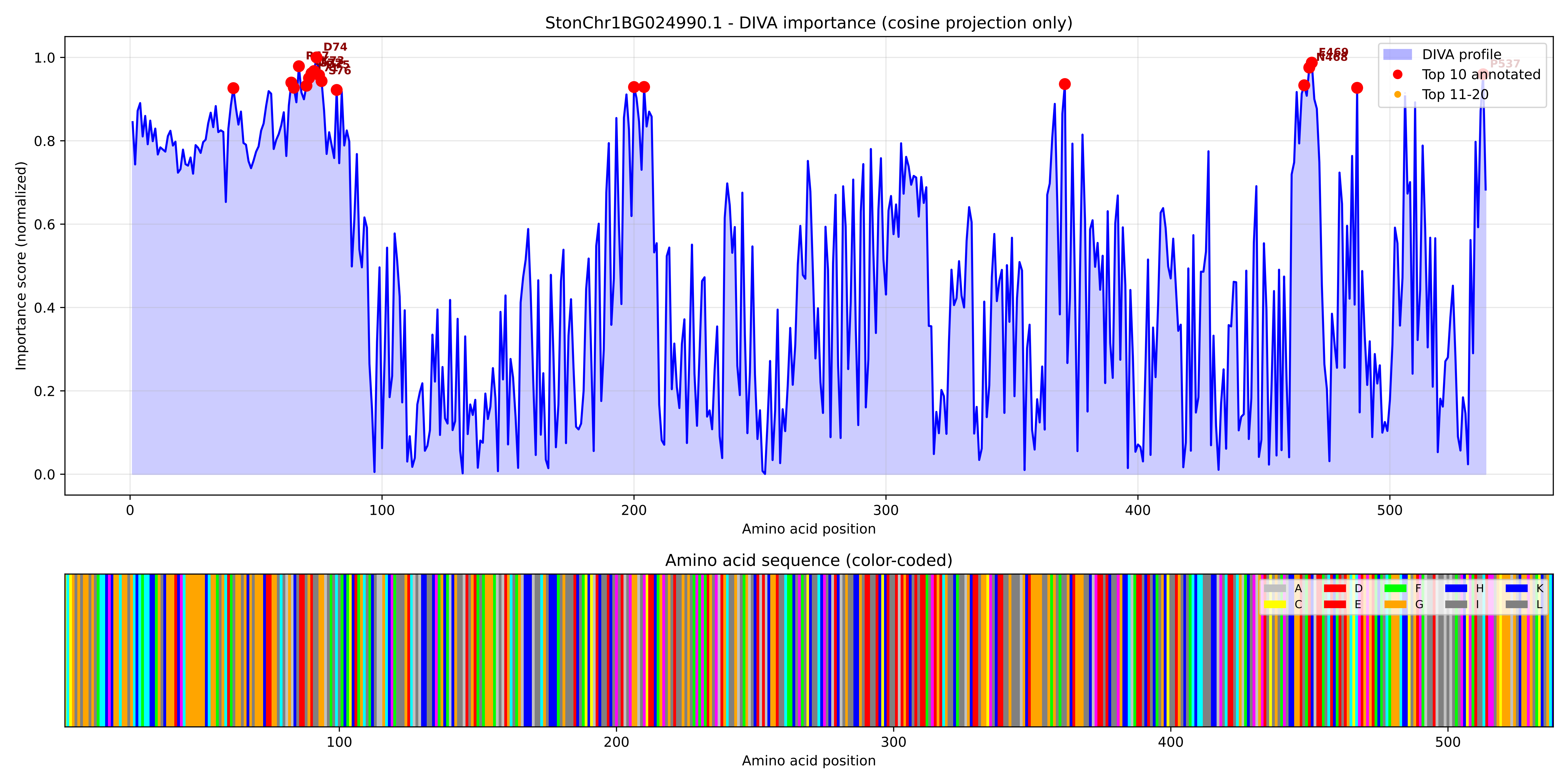

### StonChr1BG024990.1_importance.png

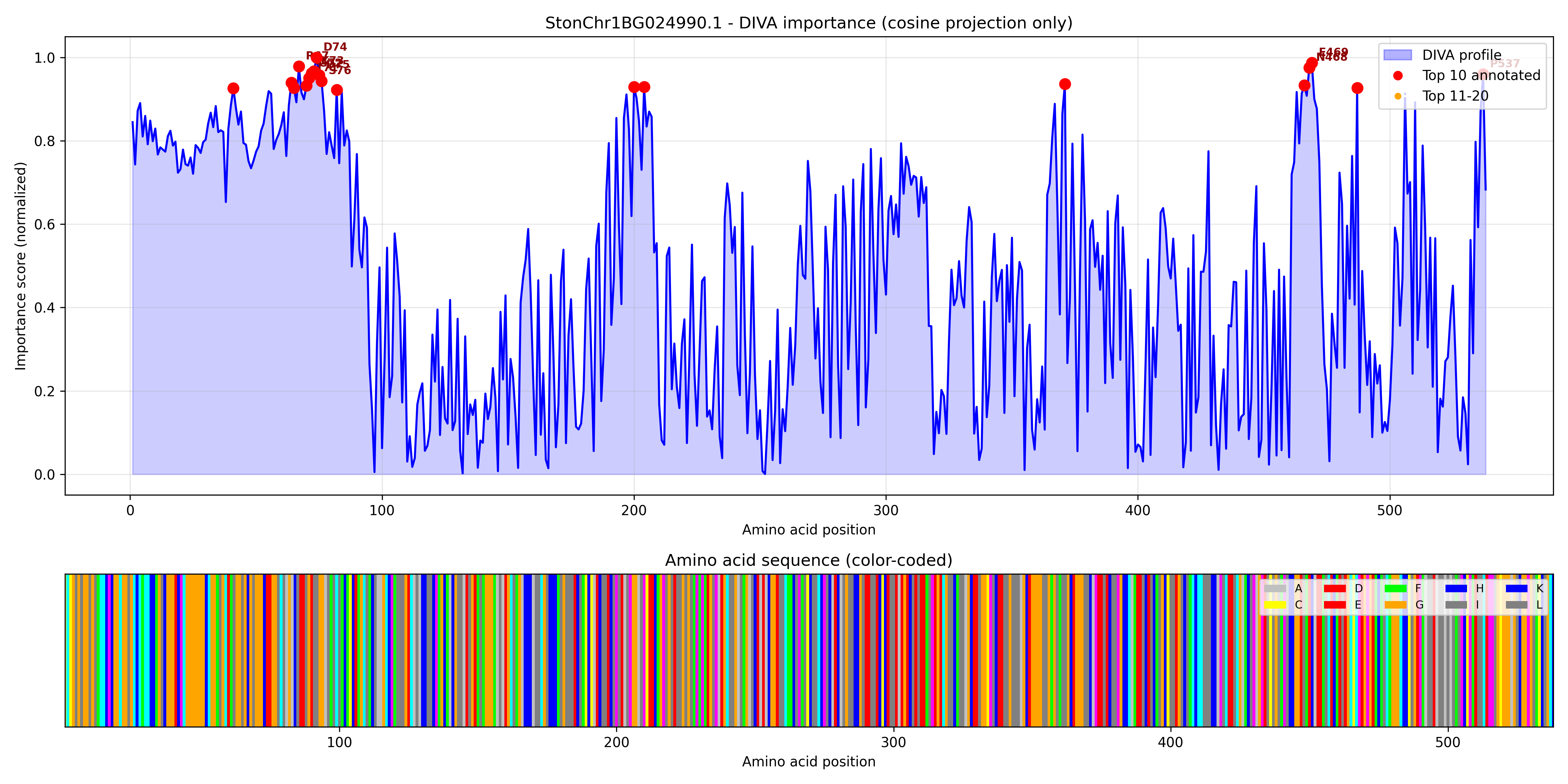

### StonChr1BG027090.1_importance.pdf

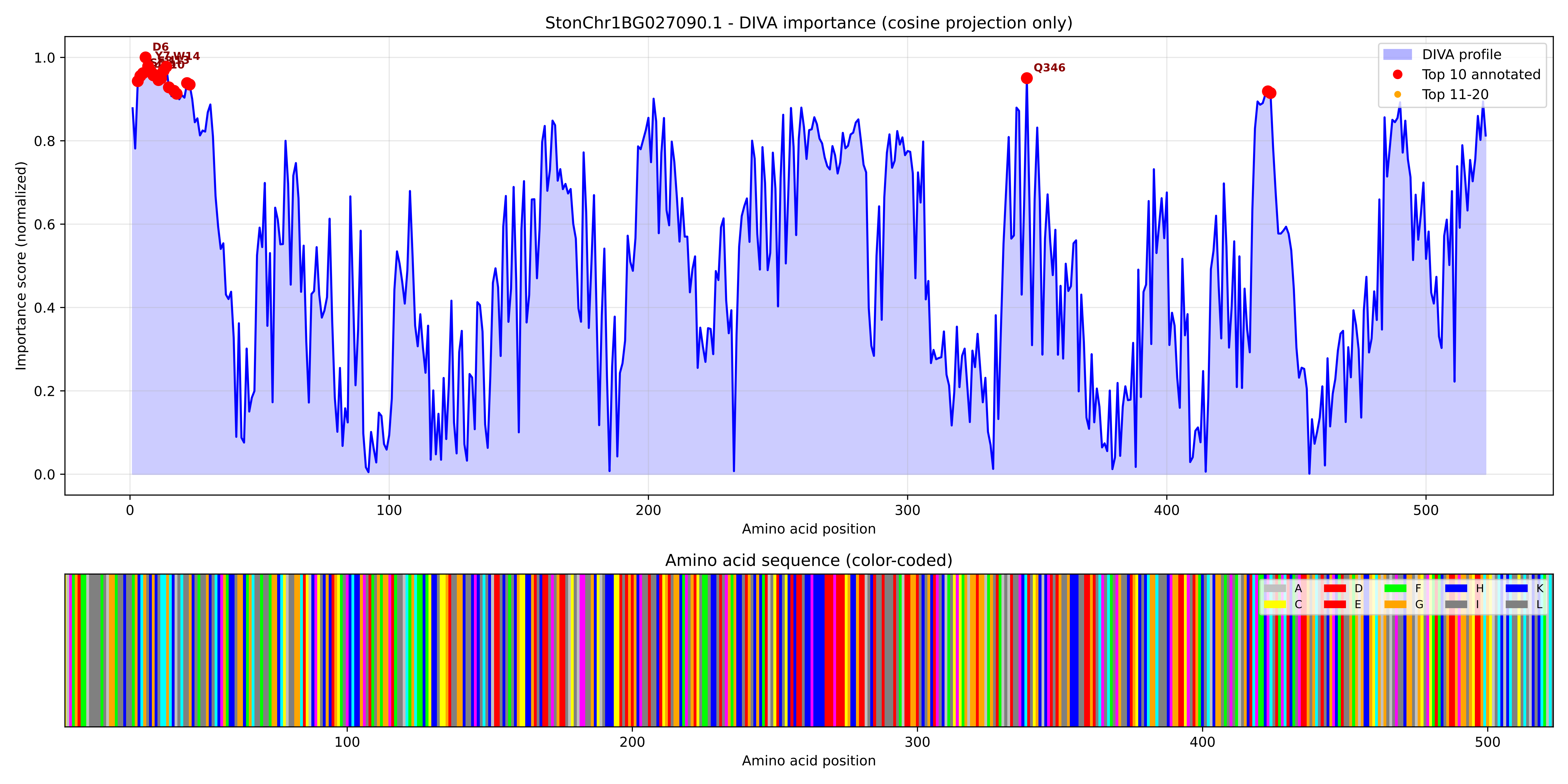

### StonChr1BG027090.1_importance.png

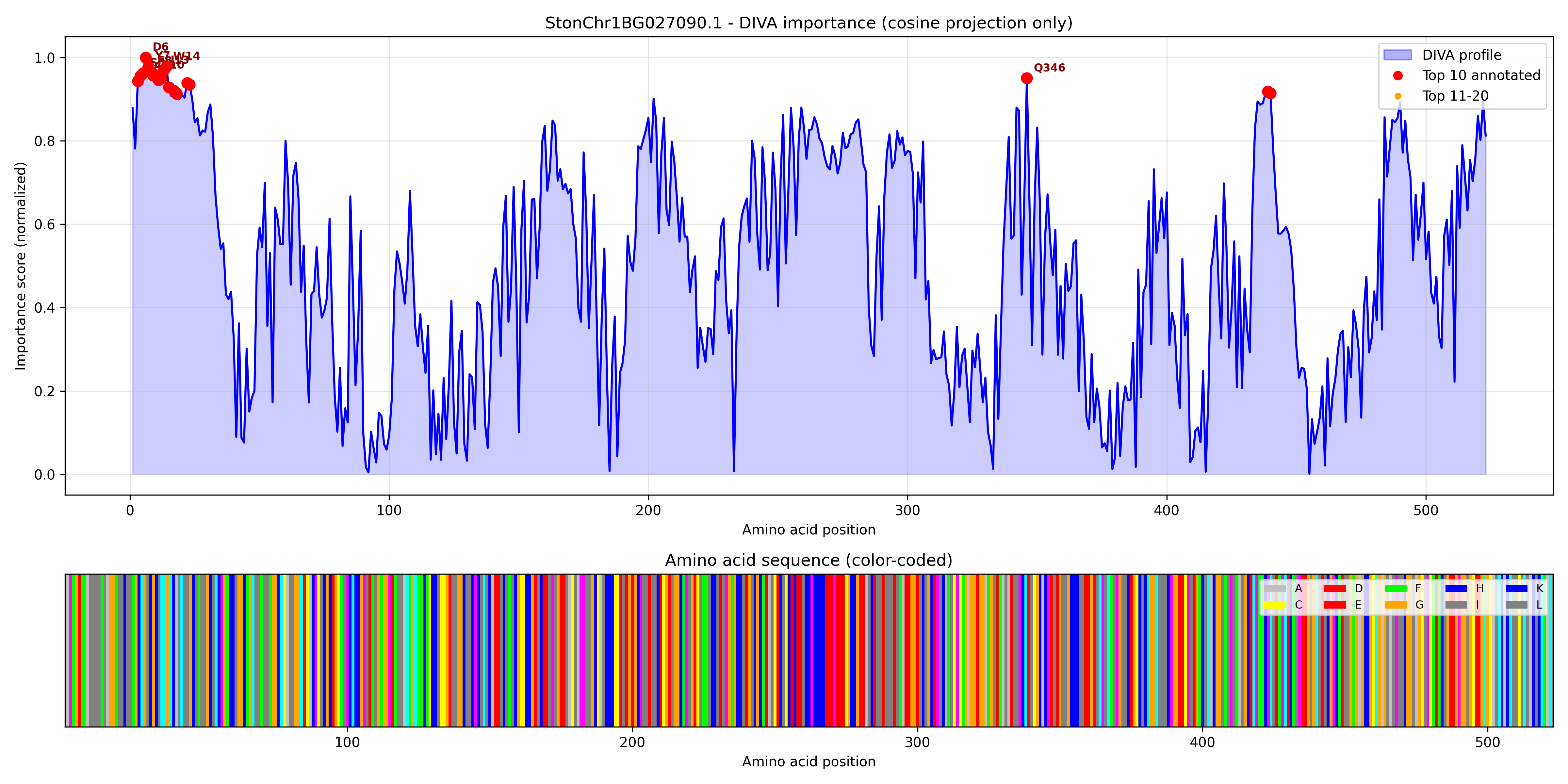

### StonChr1BG028350.1_importance.pdf

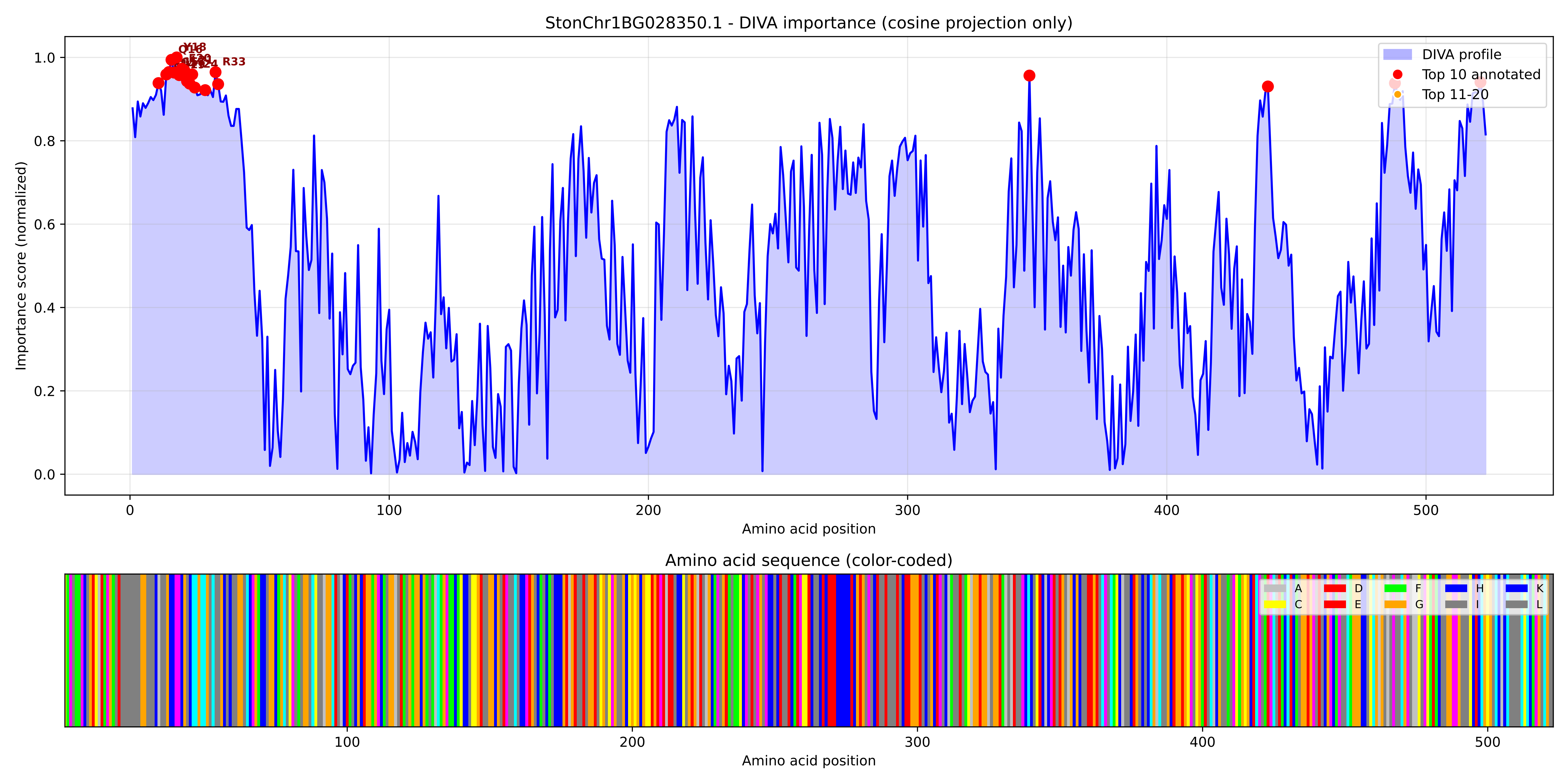

### StonChr1BG028350.1_importance.png

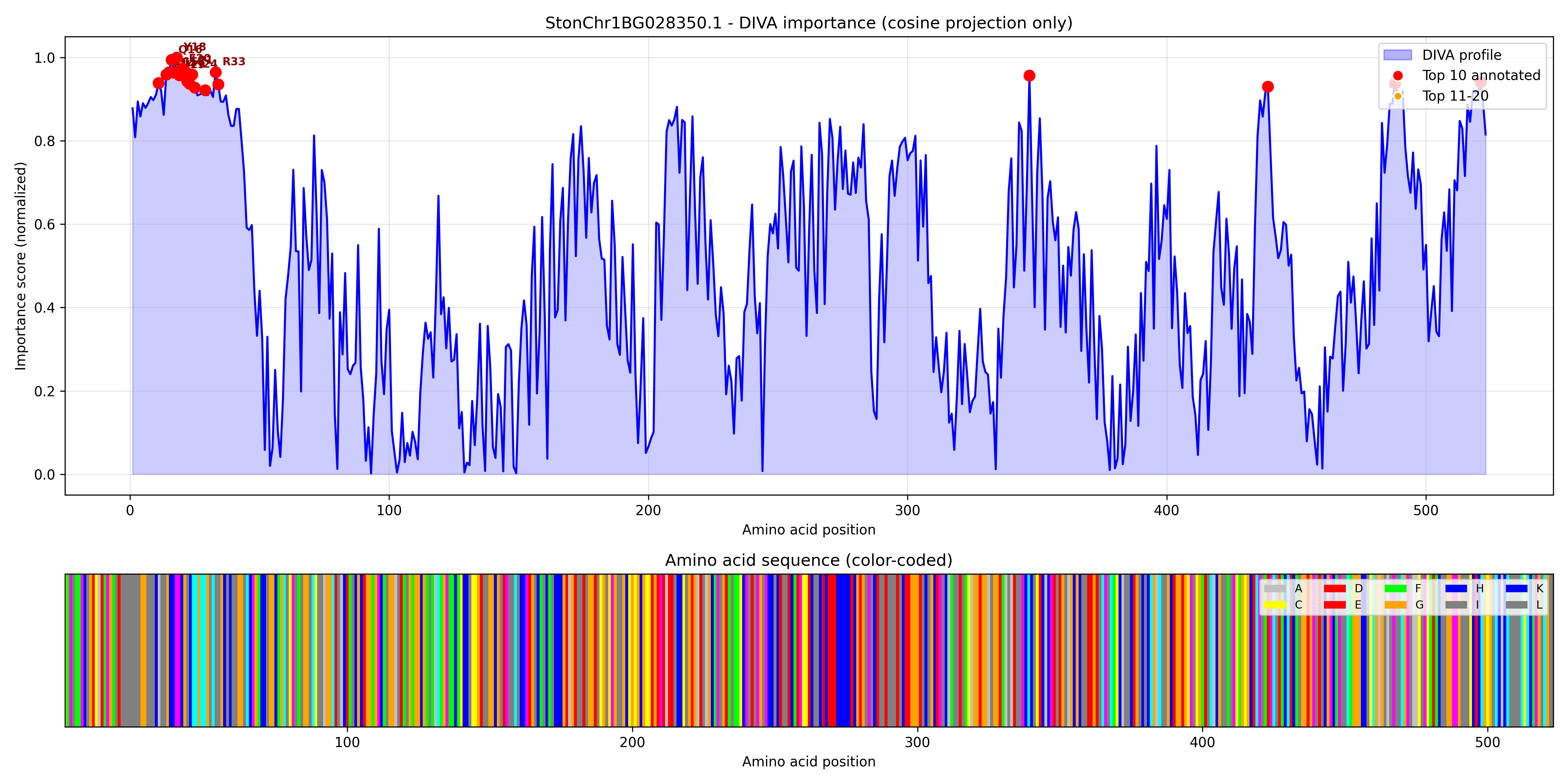

### StonChr1BG028380.1_importance.pdf

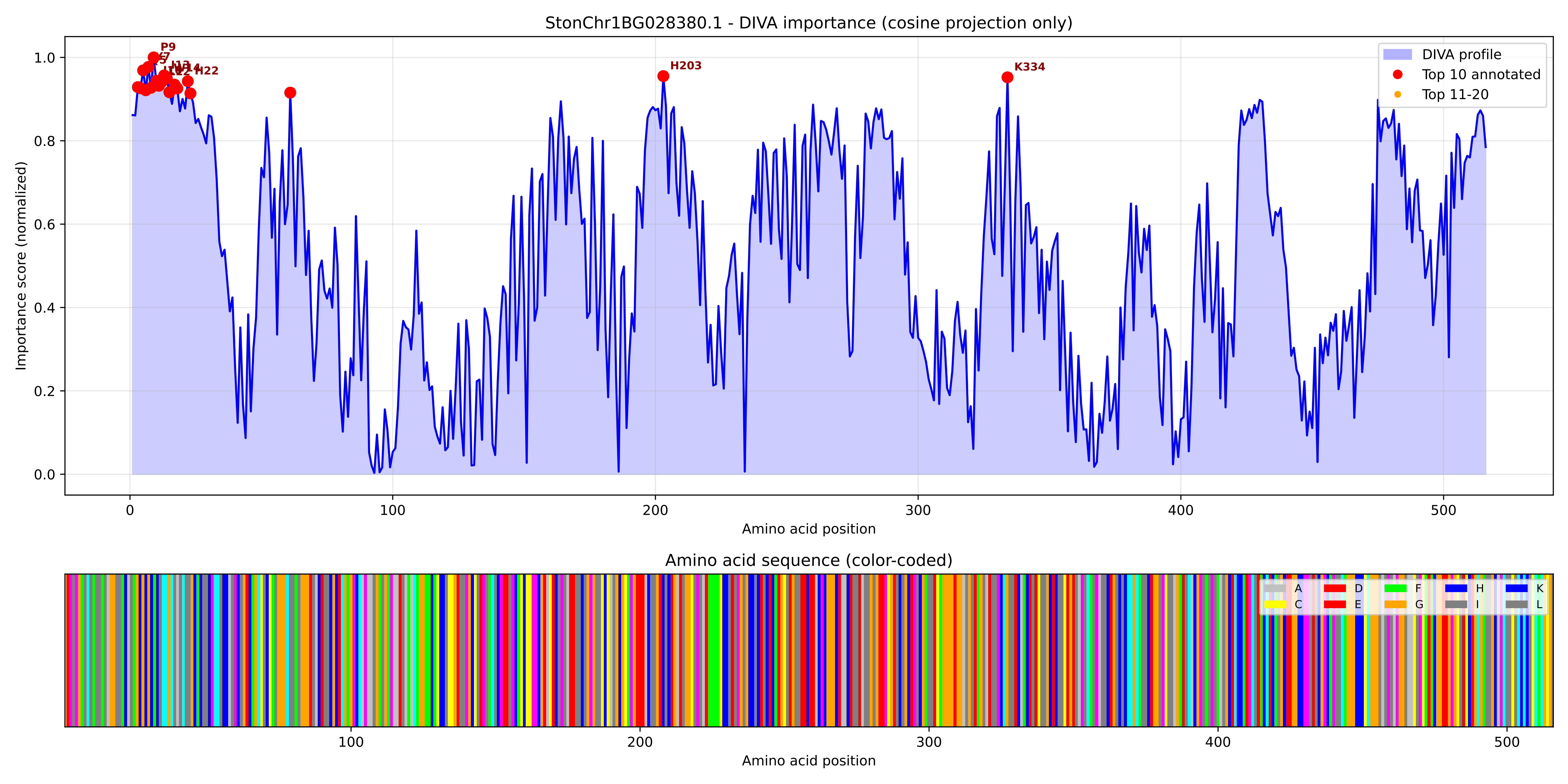

### StonChr1BG028380.1_importance.png

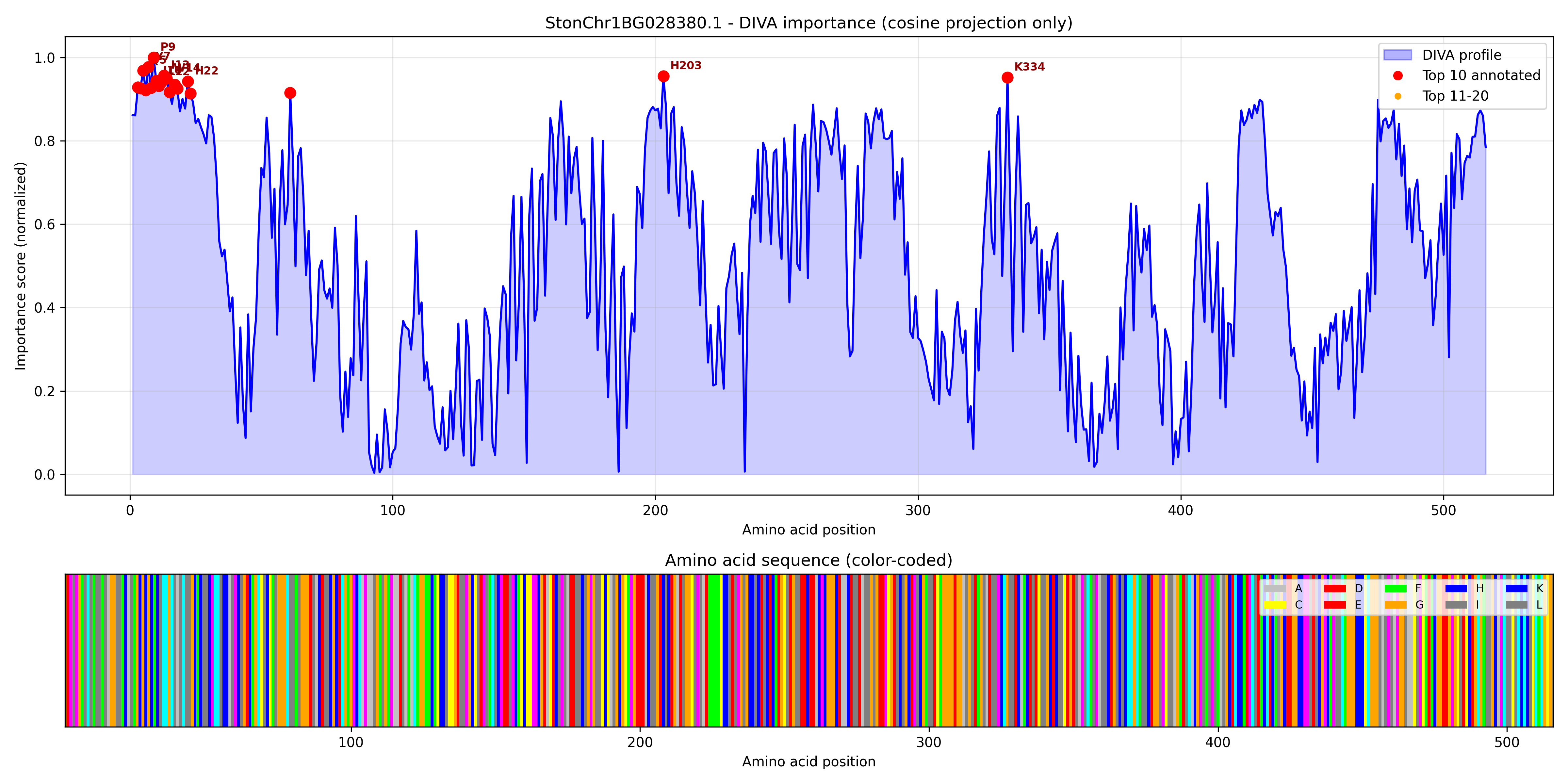

### StonChr1BG032350.1_importance.pdf

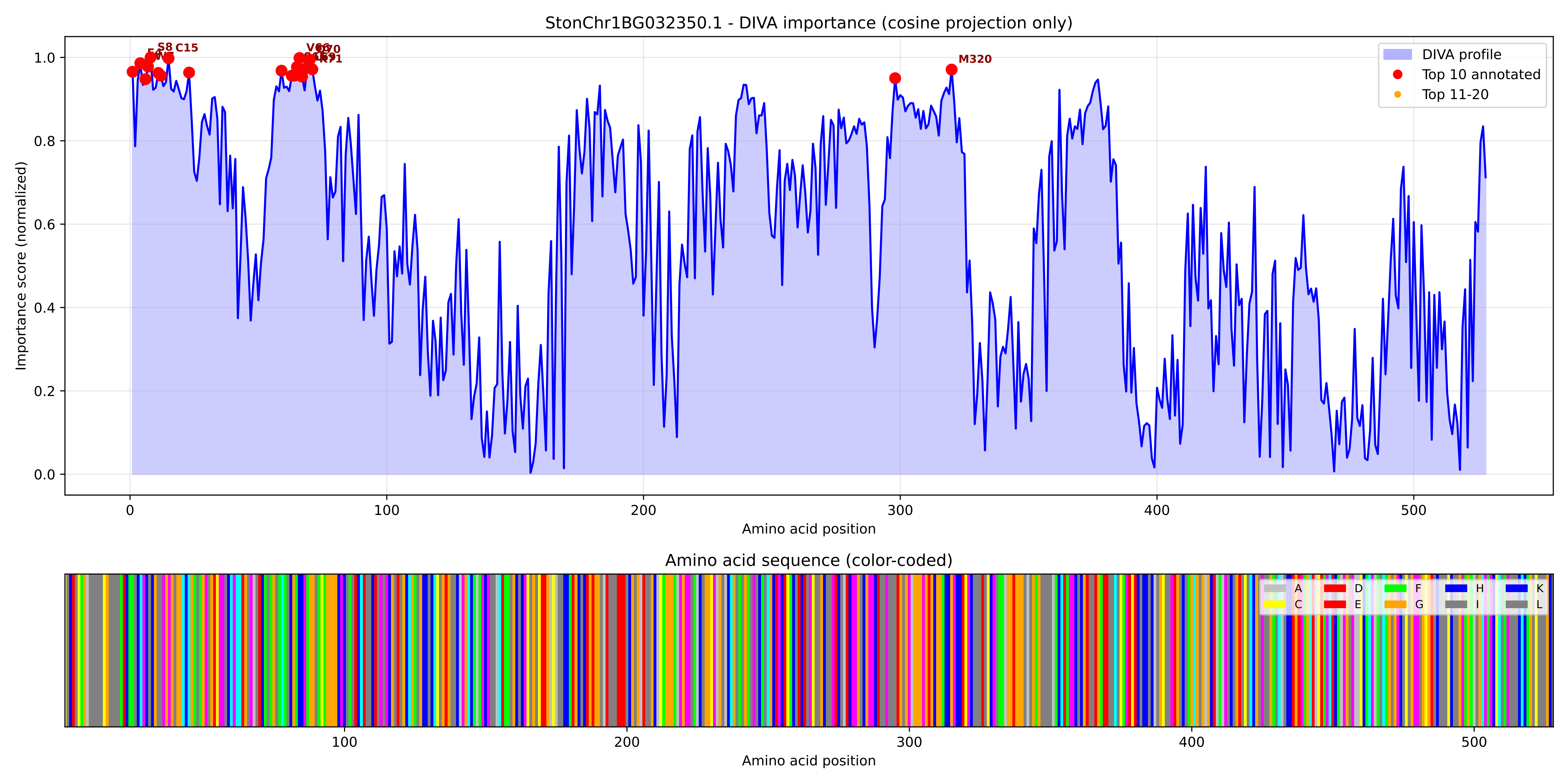

### StonChr1BG032350.1_importance.png

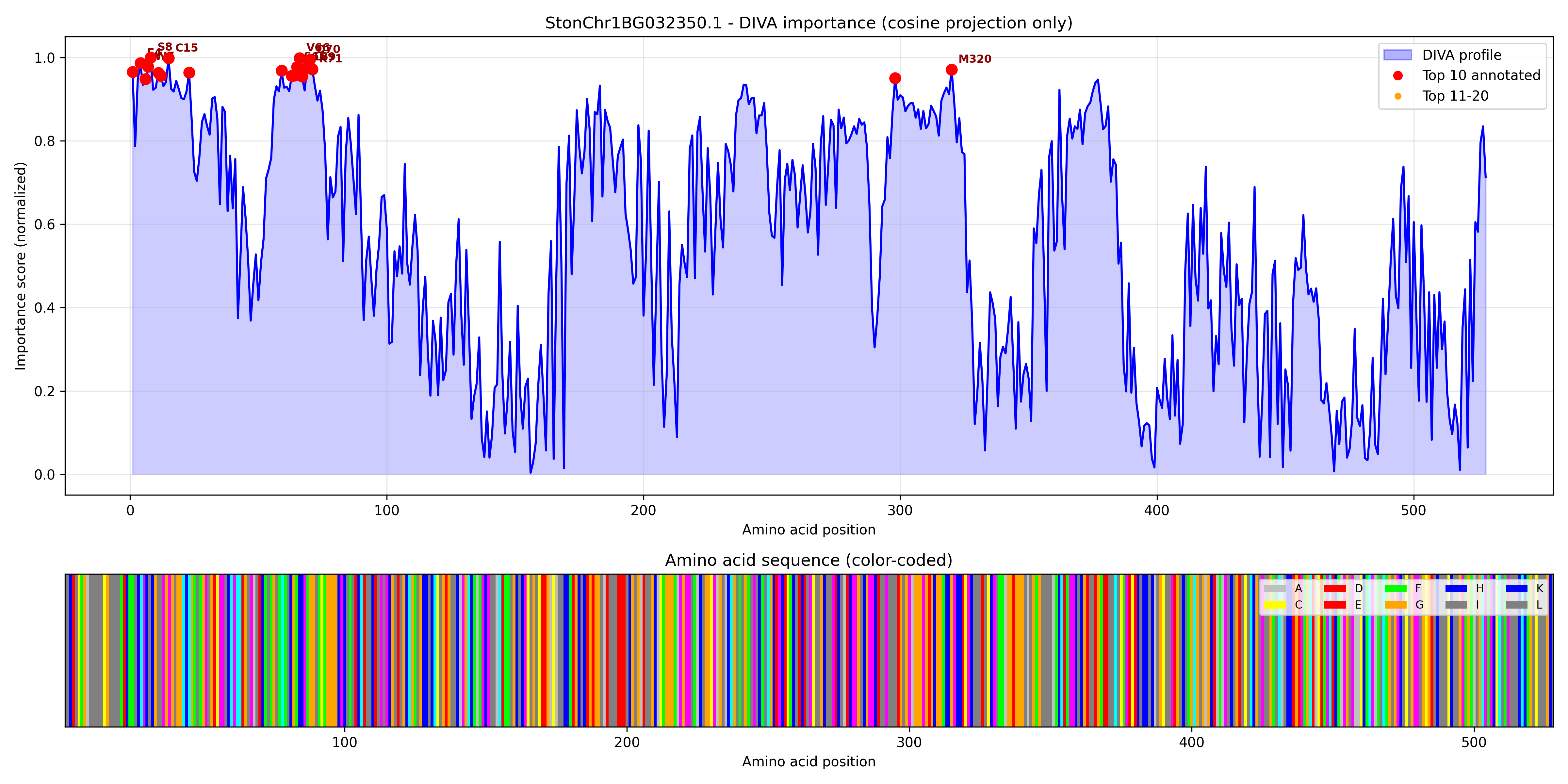

### StonChr1BG035030.1_importance.pdf

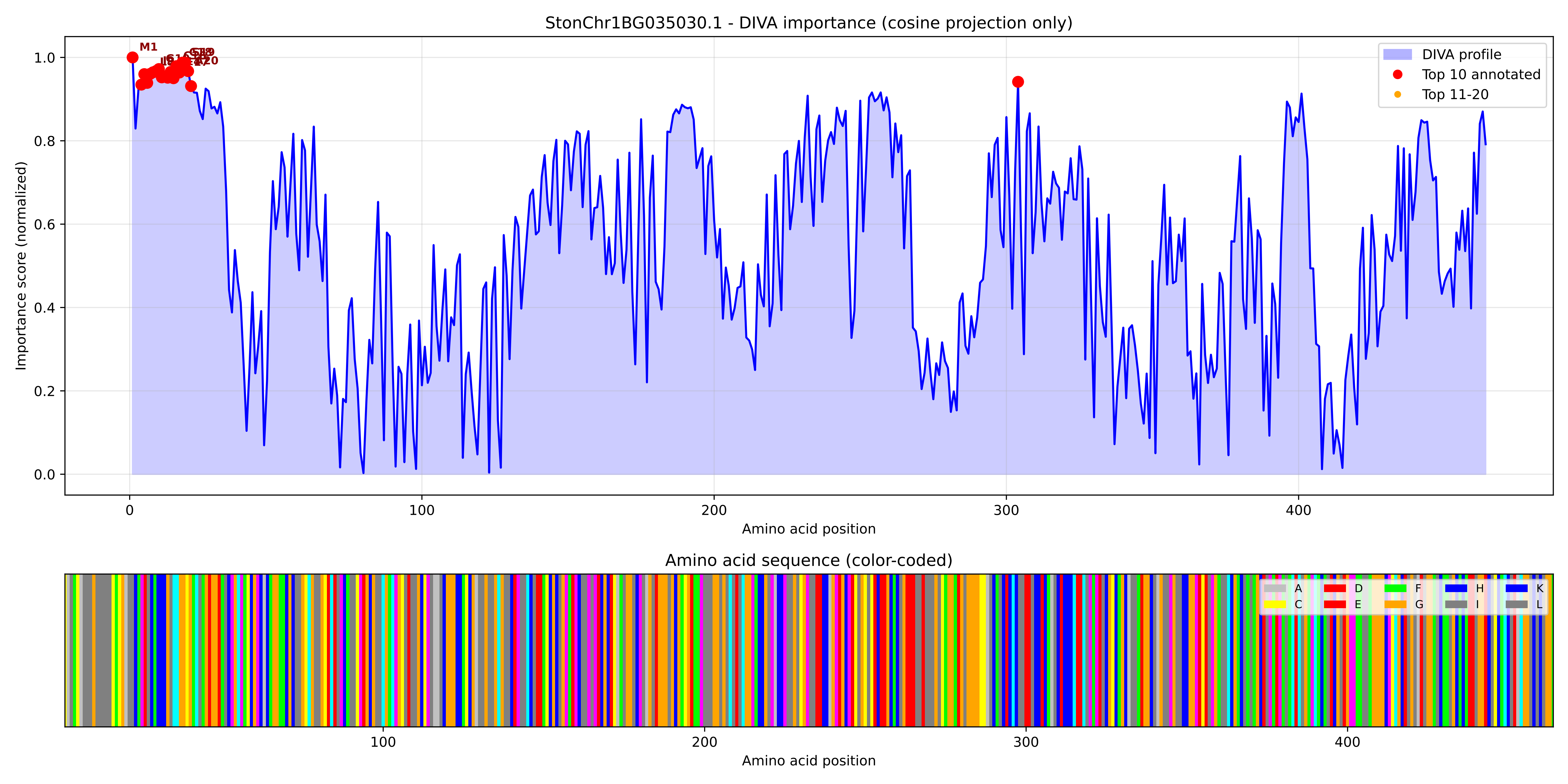

### StonChr1BG035030.1_importance.png

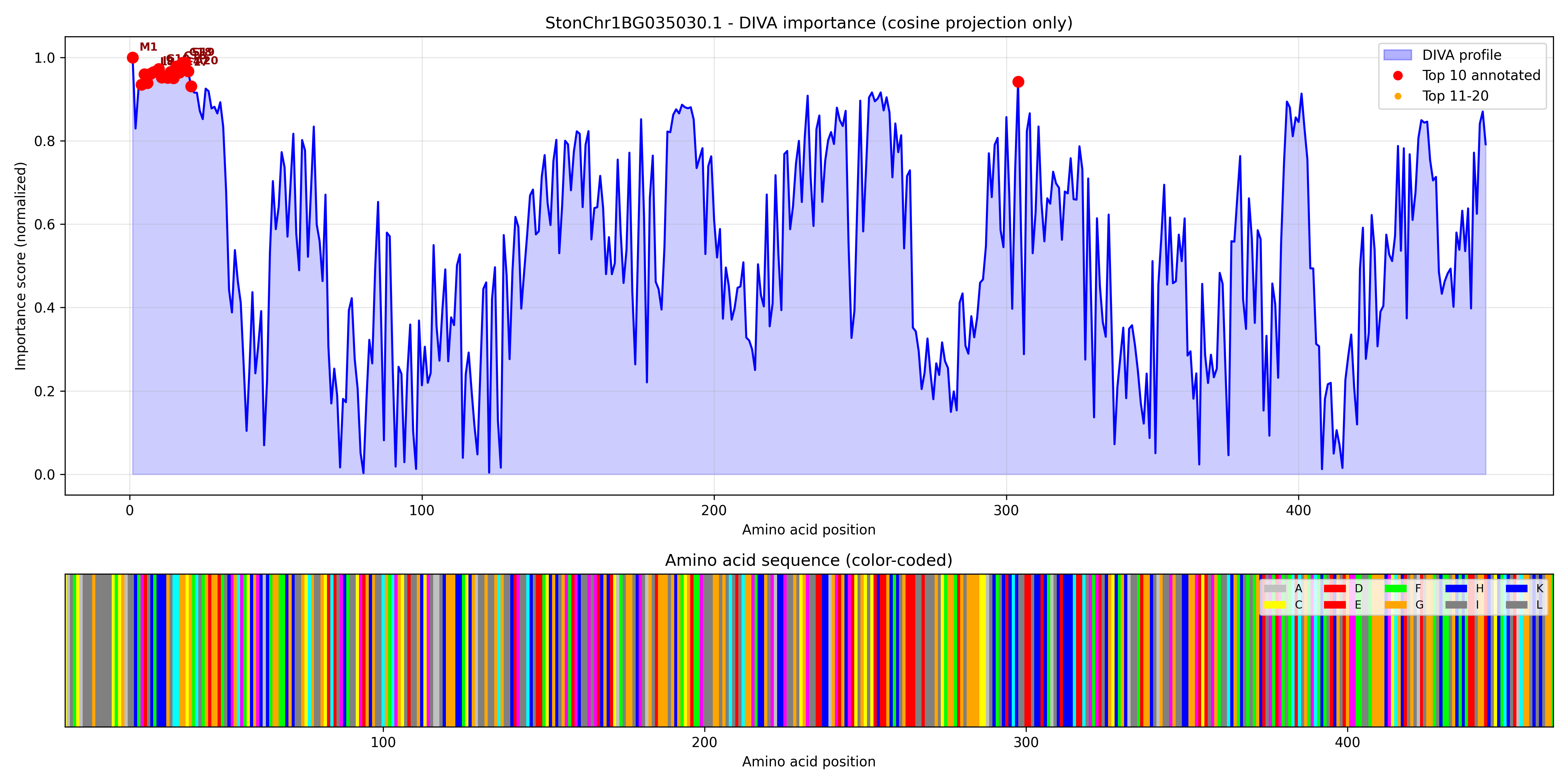

### StonChr1BG035310.1_importance.pdf

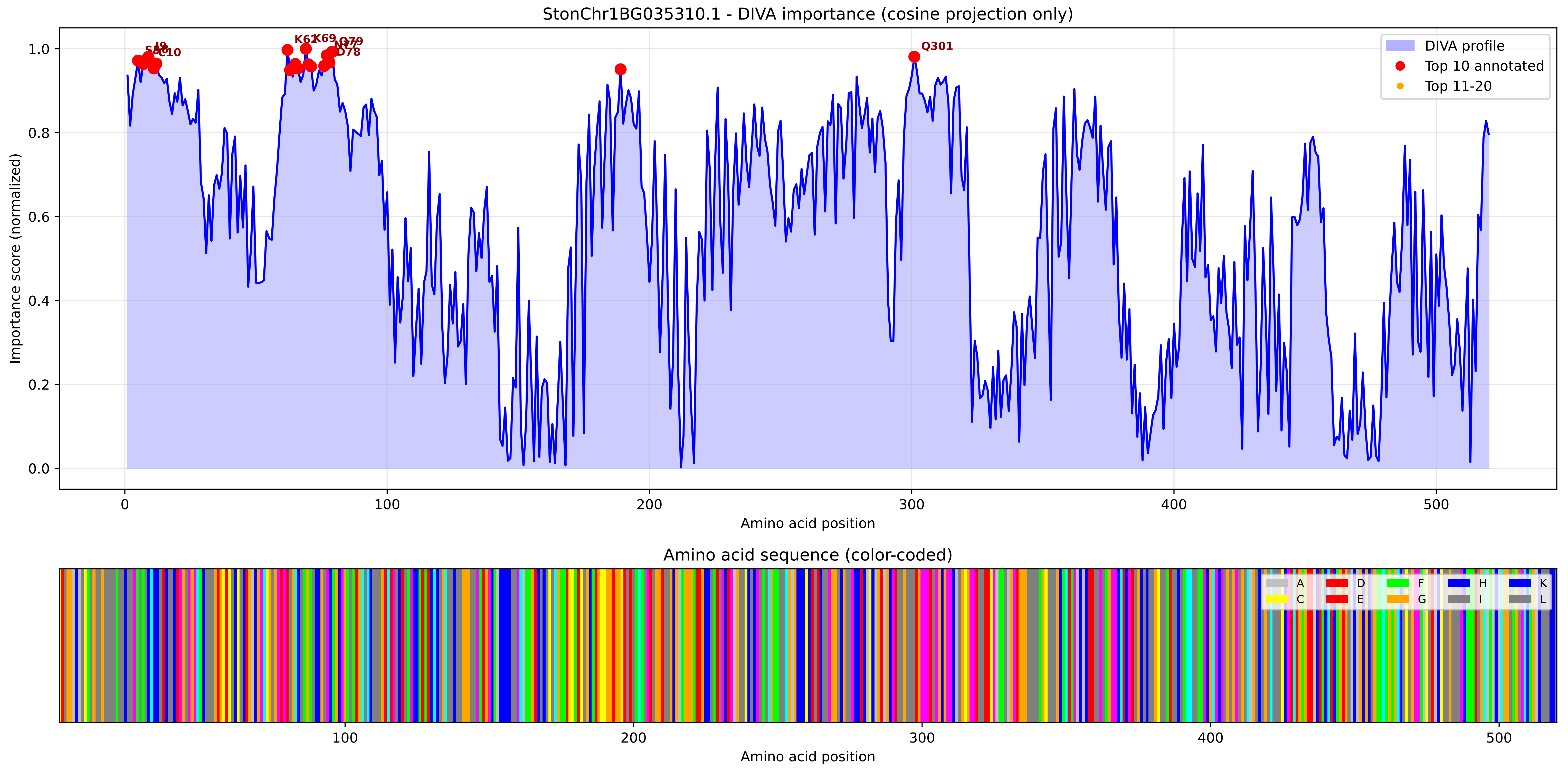

### StonChr1BG035310.1_importance.png

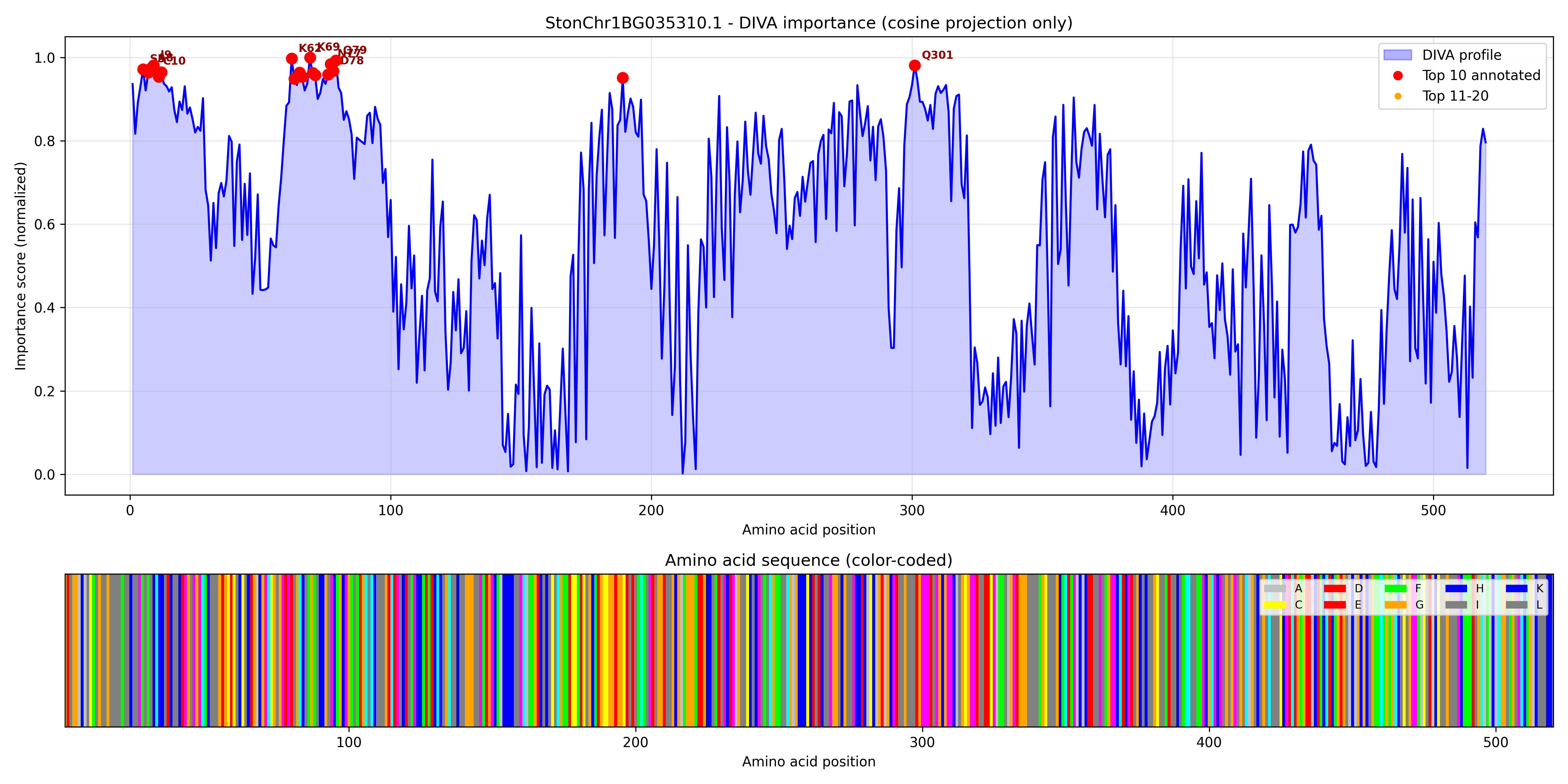

### StonChr1BG037600.1_importance.pdf

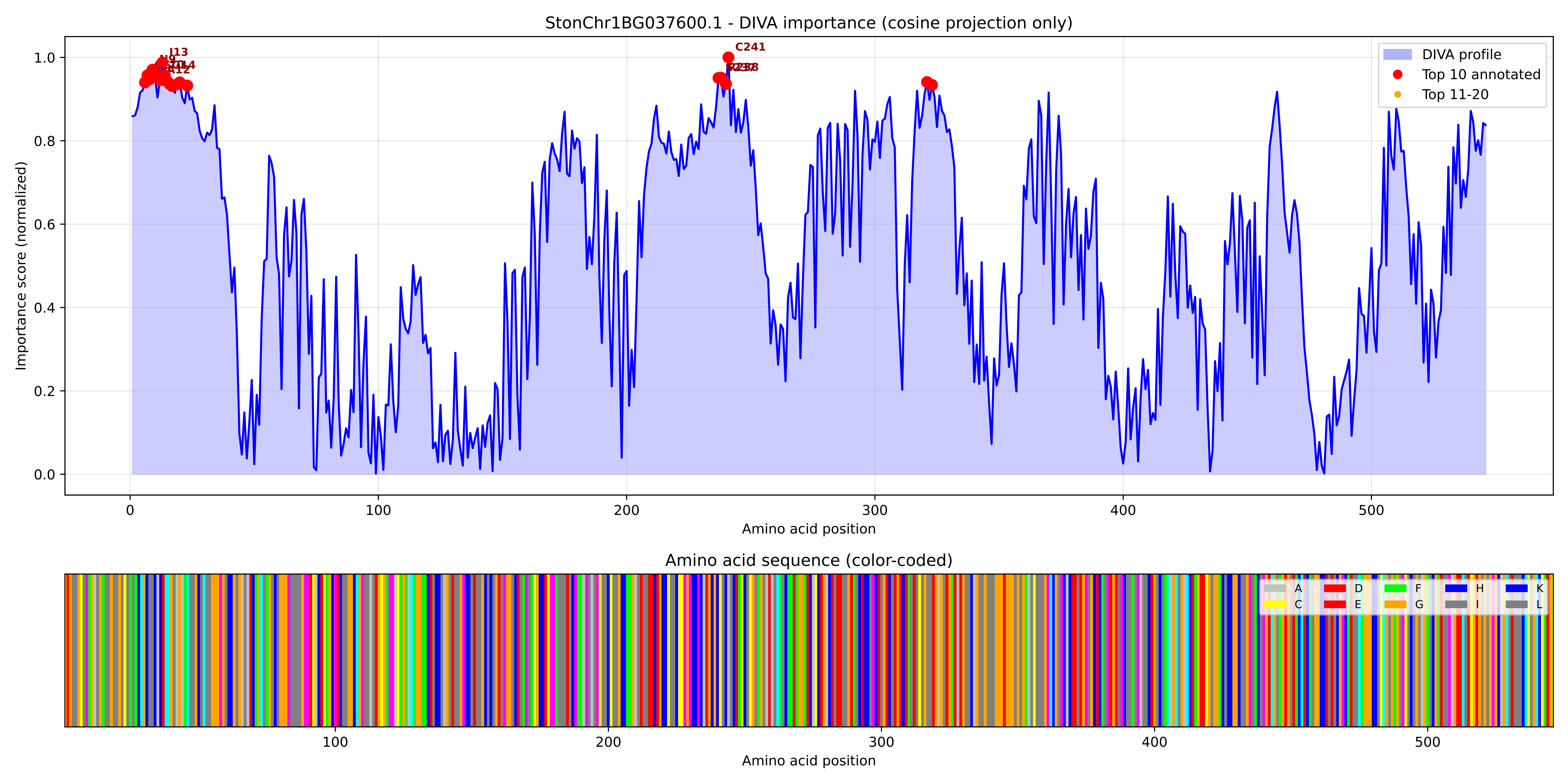

### StonChr1BG037600.1_importance.png

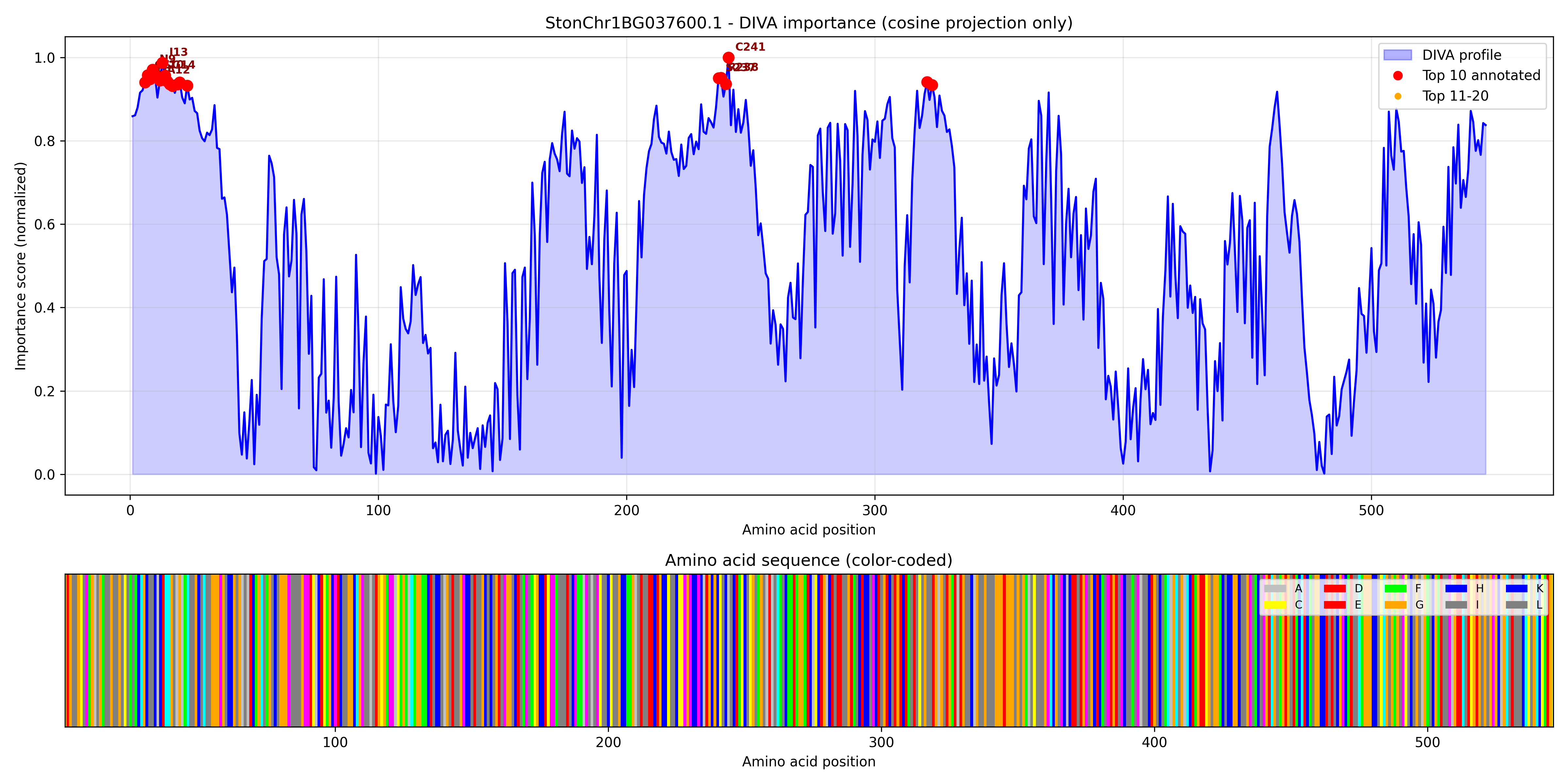

### StonChr1BG038920.1_importance.pdf

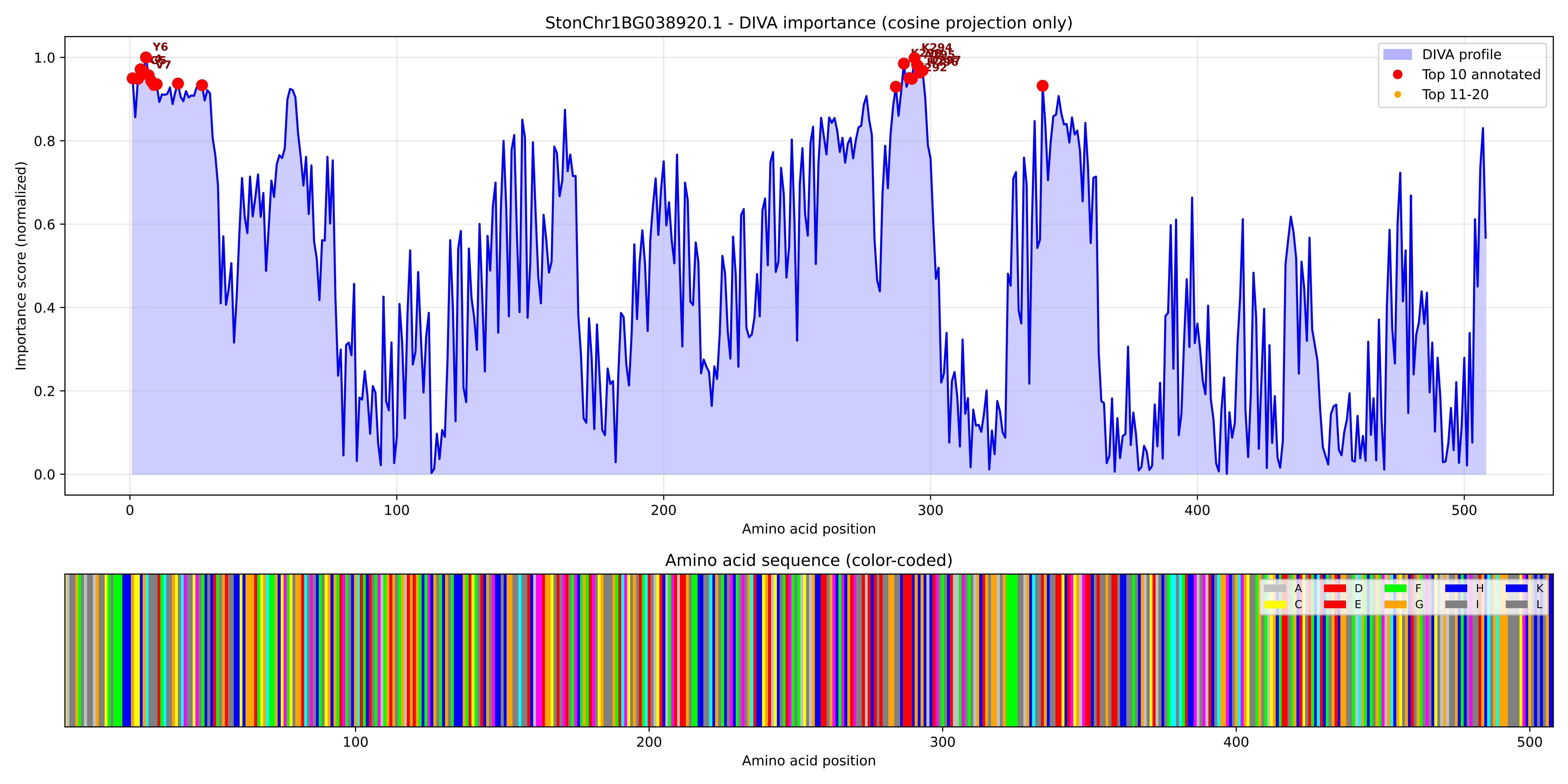

### StonChr1BG038920.1_importance.png

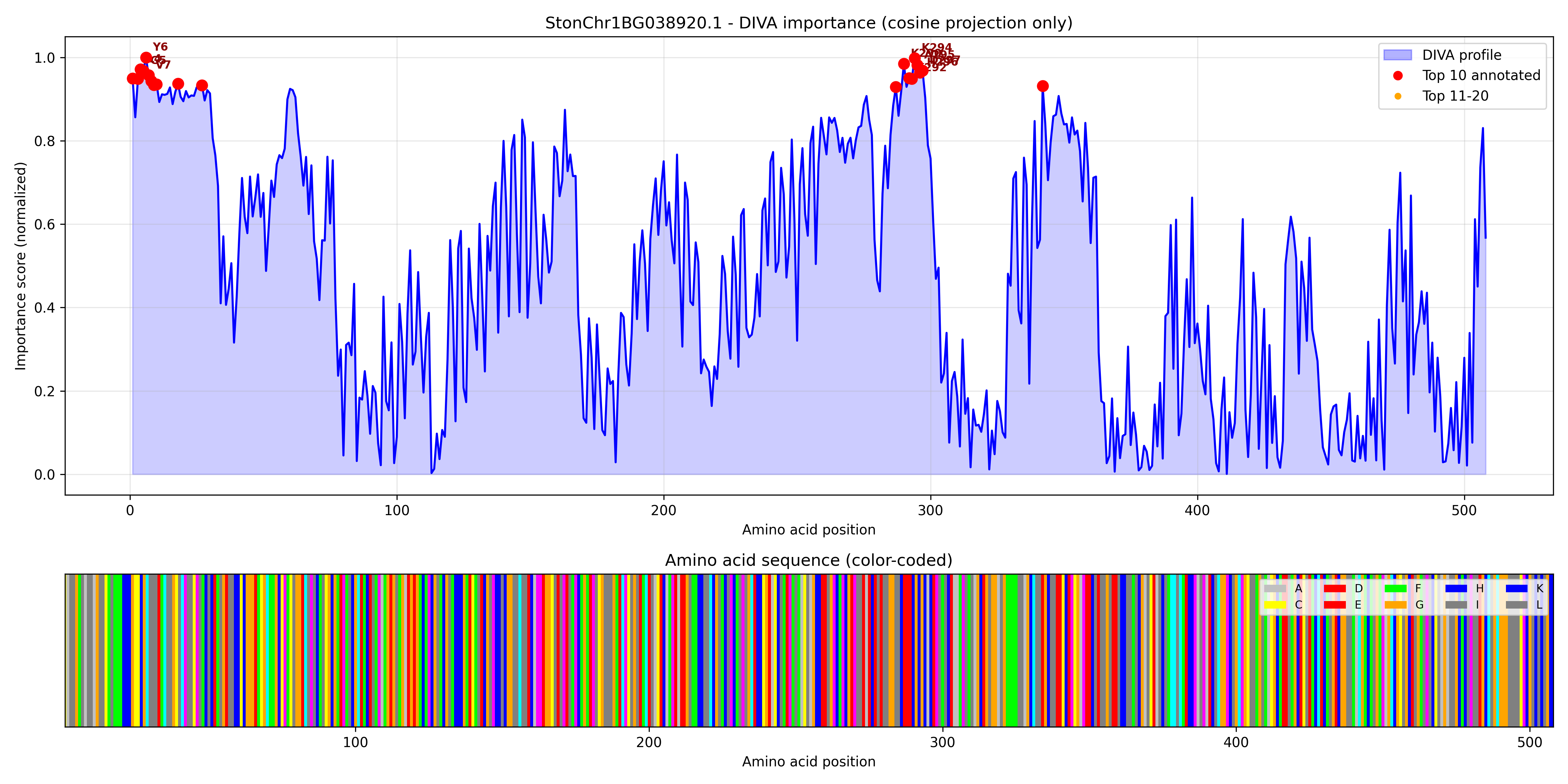

### StonChr2AG039470.1_importance.pdf

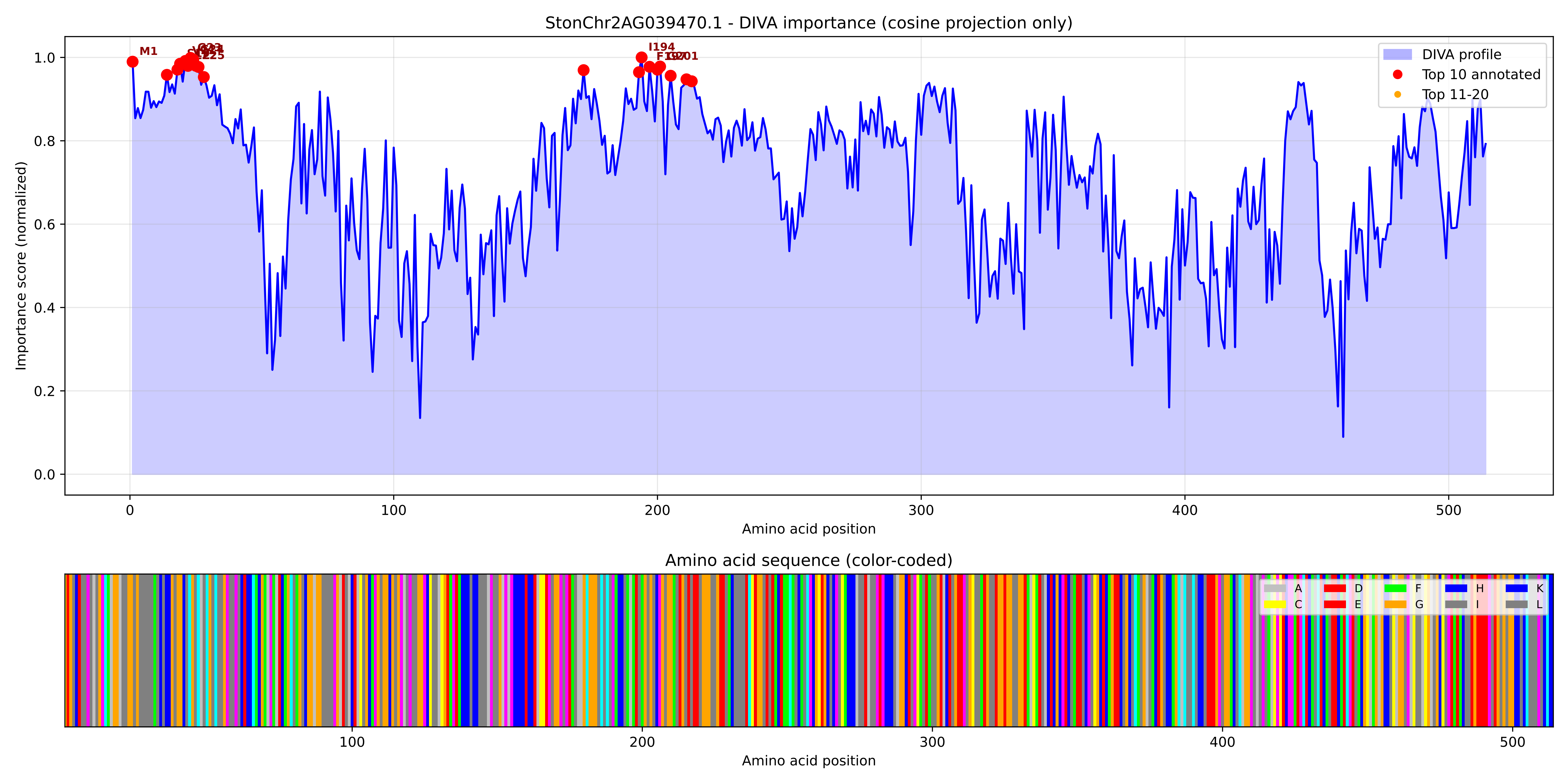

### StonChr2AG039470.1_importance.png

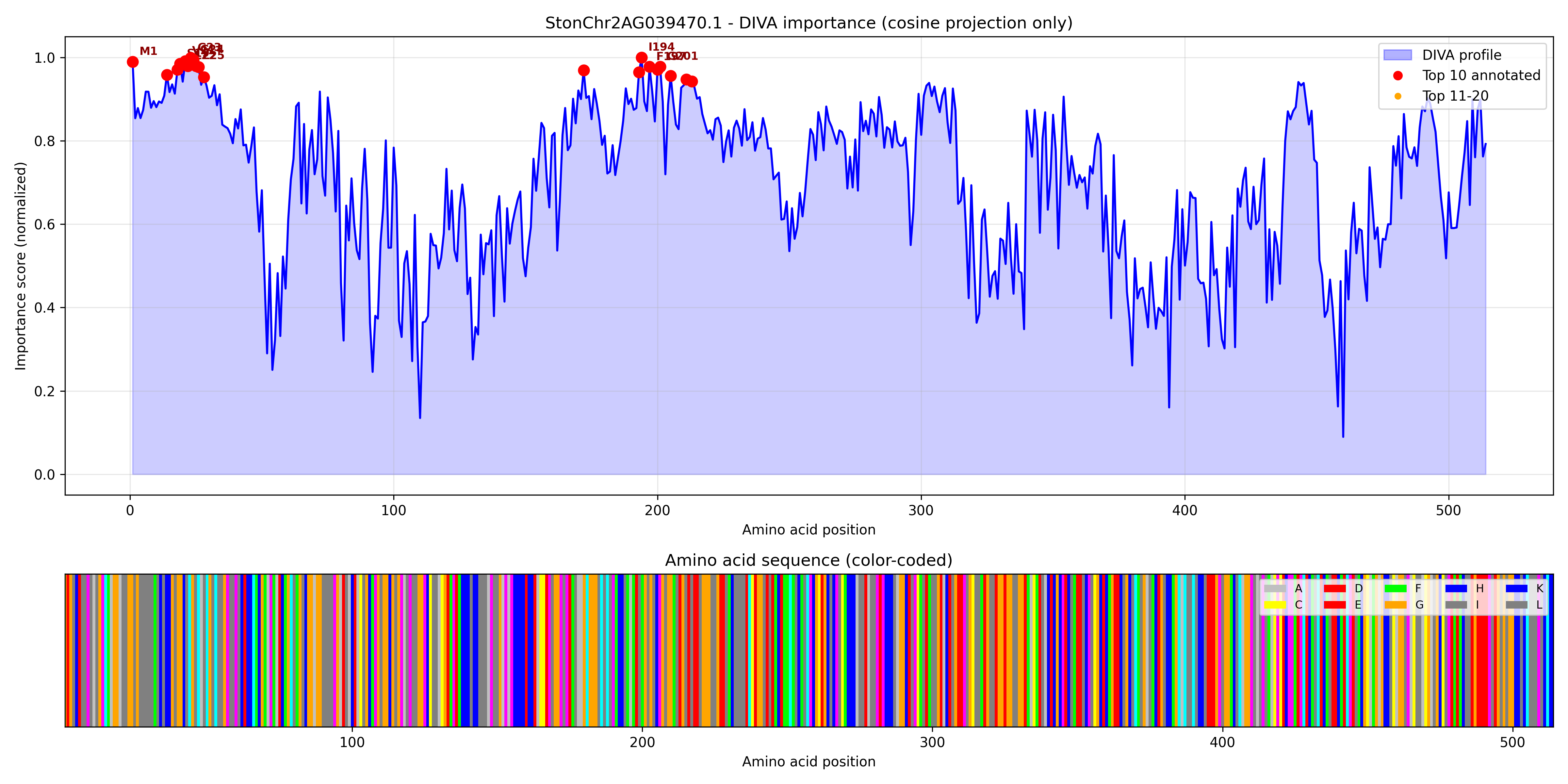

### StonChr2AG042630.1_importance.pdf

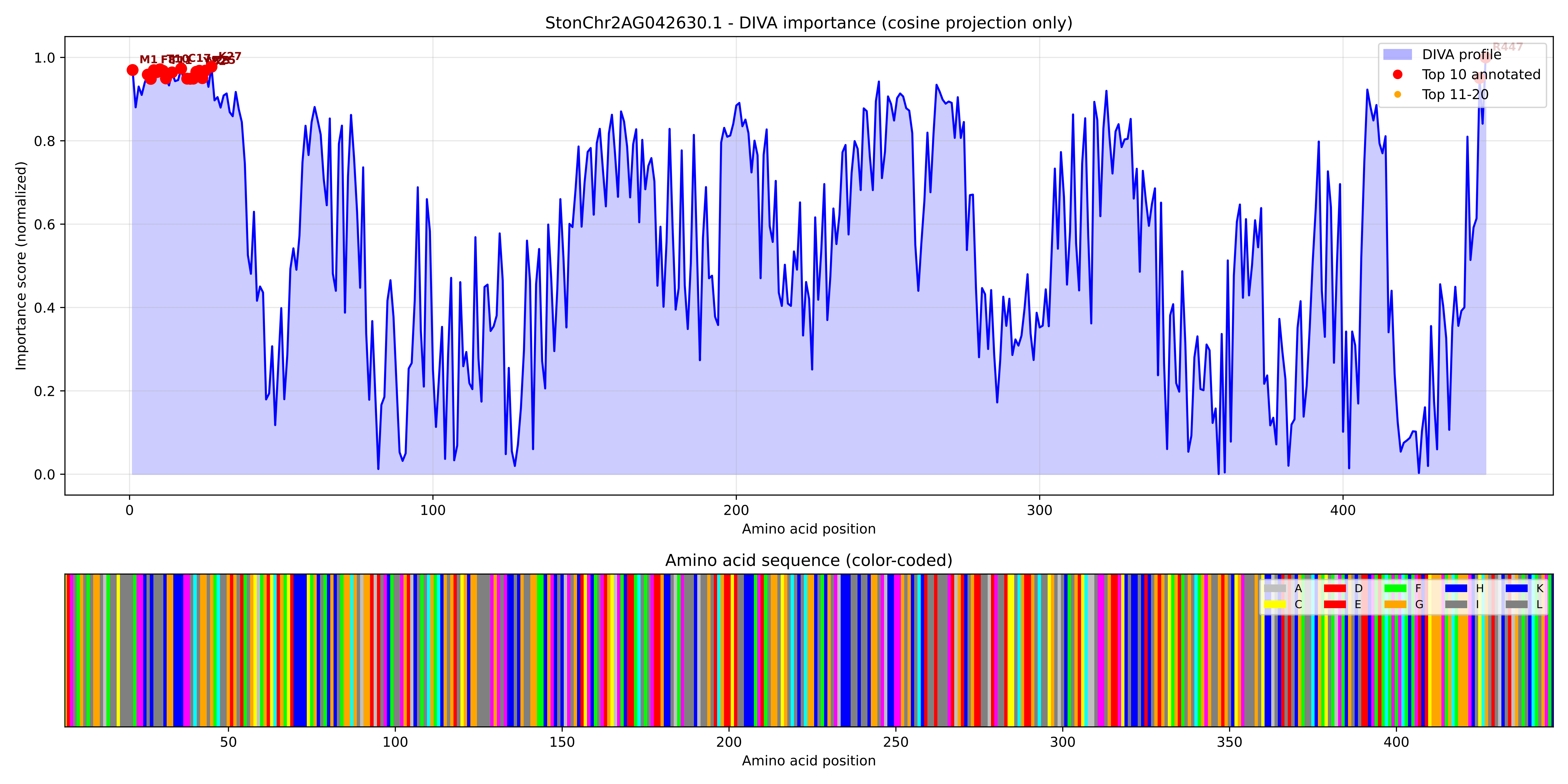

### StonChr2AG042630.1_importance.png

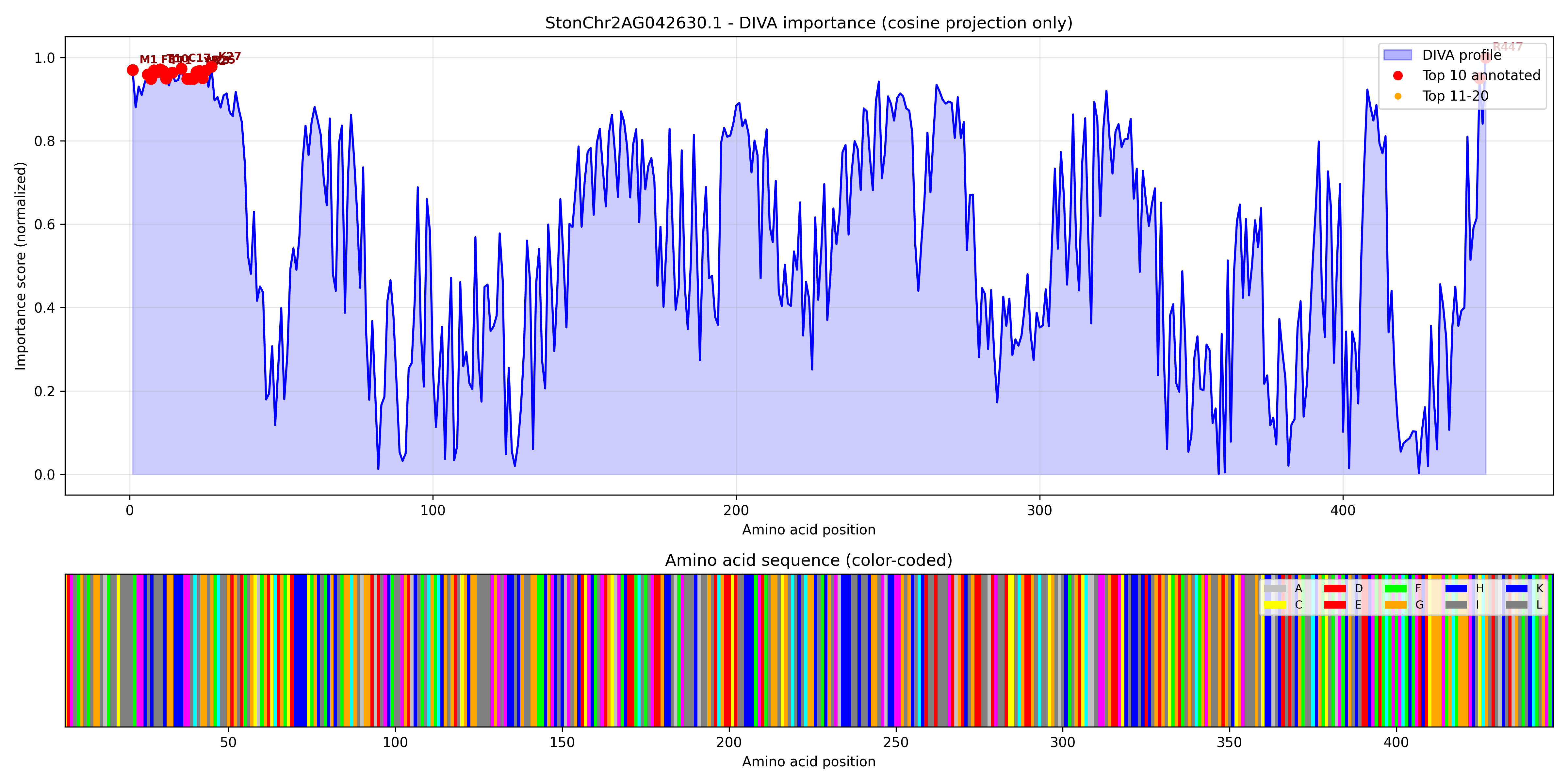

### StonChr2AG044080.1_importance.pdf

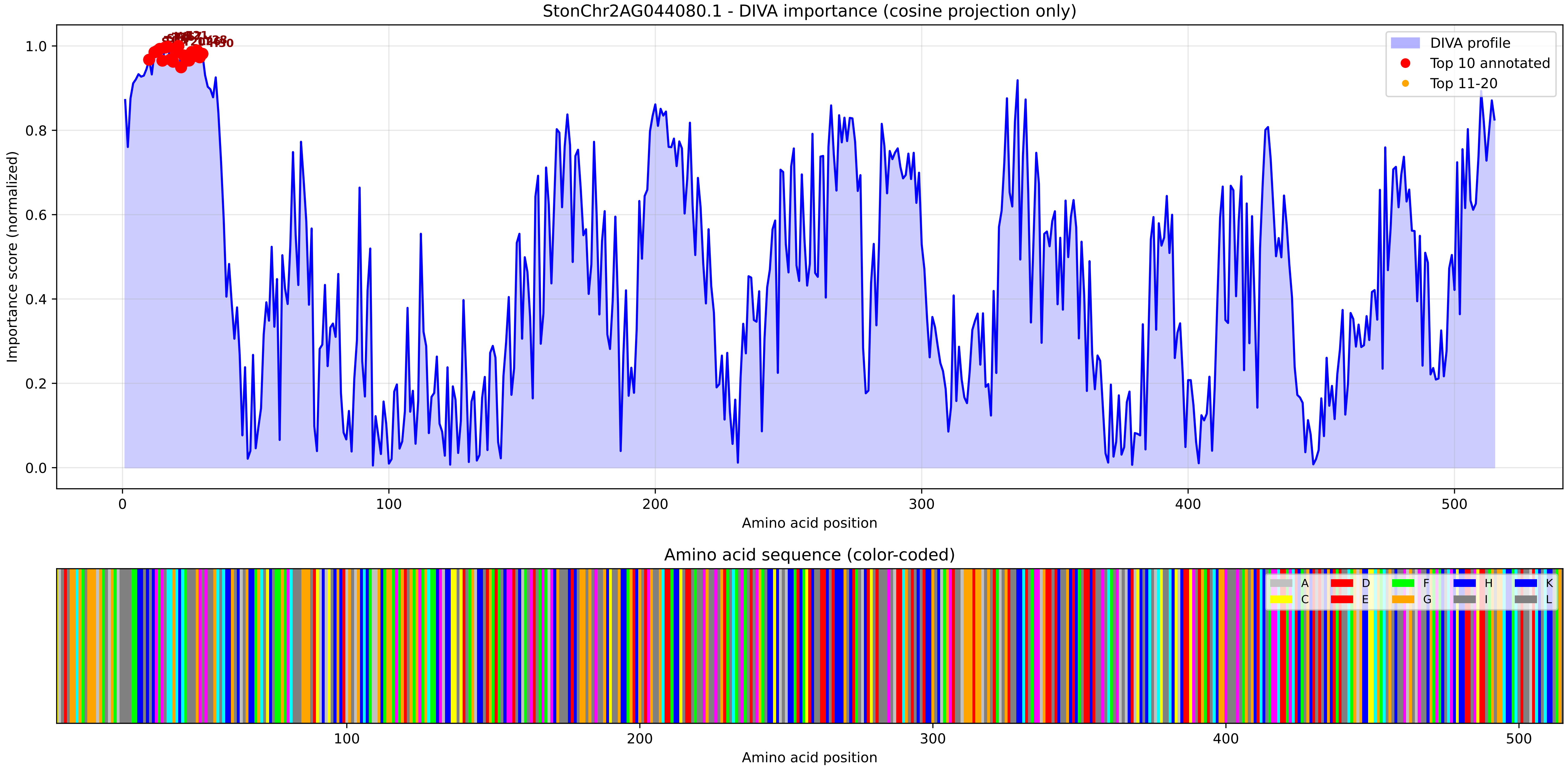

### StonChr2AG044080.1_importance.png

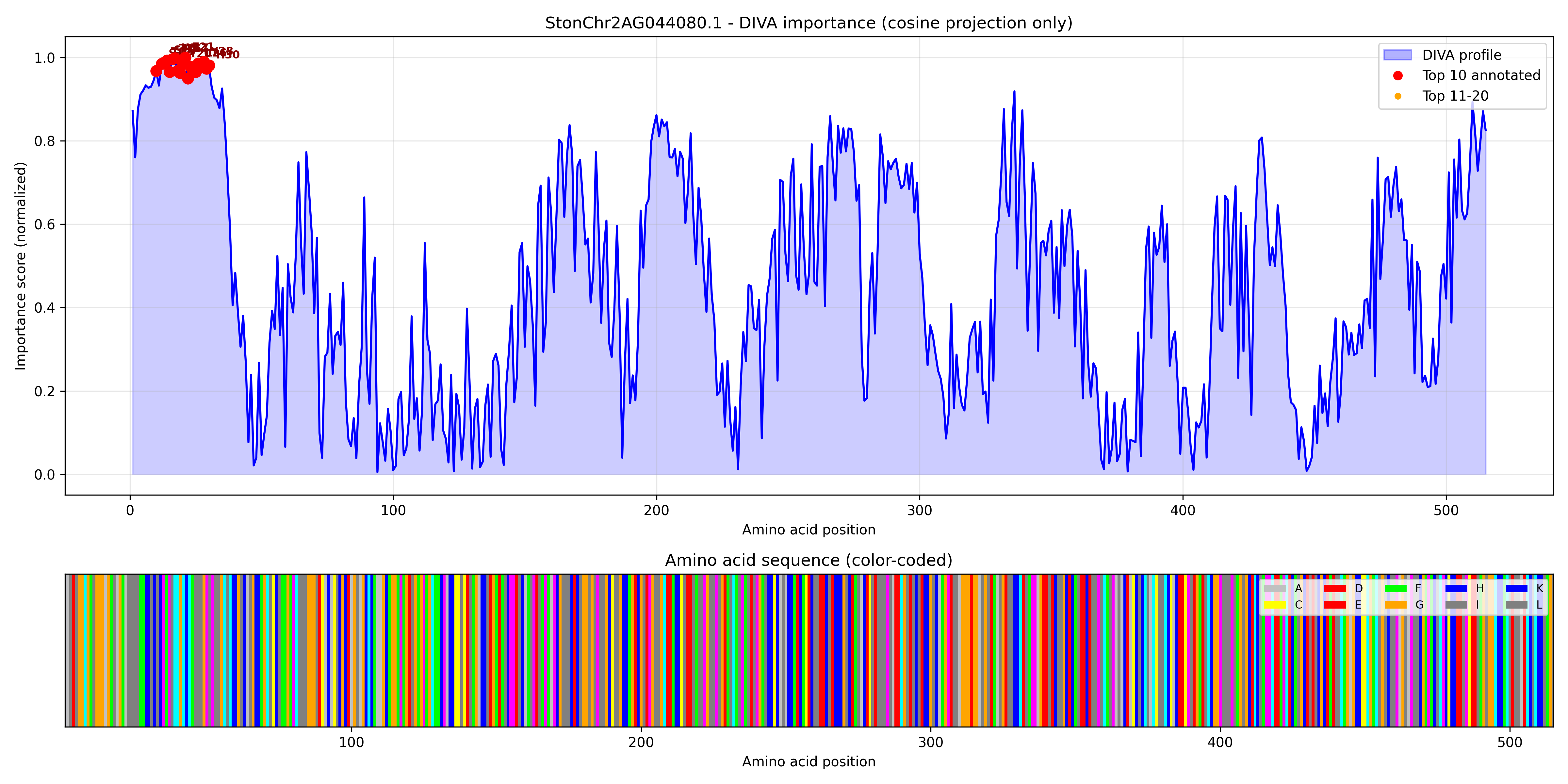

### StonChr2AG044120.1_importance.pdf

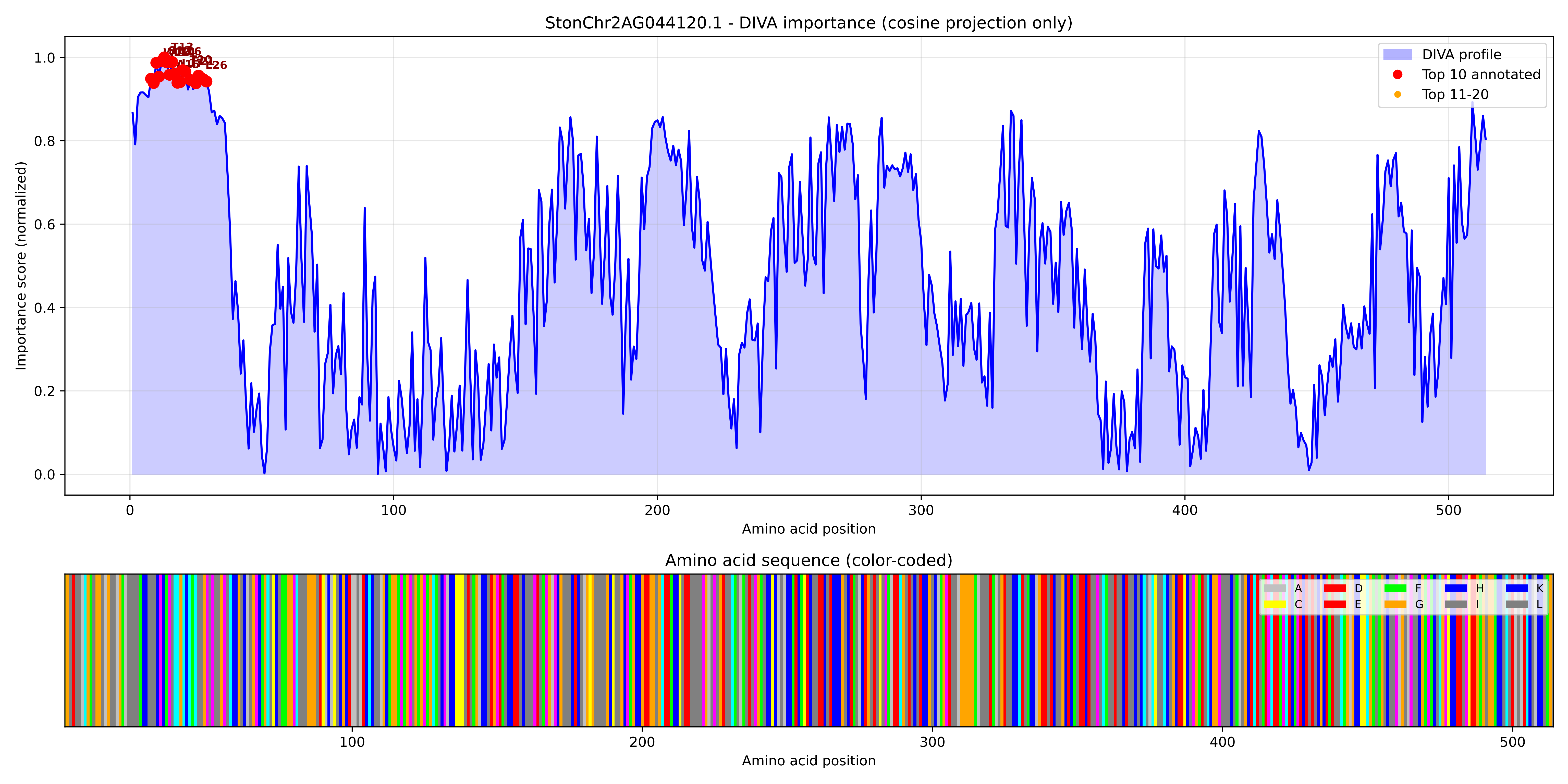

### StonChr2AG044120.1_importance.png

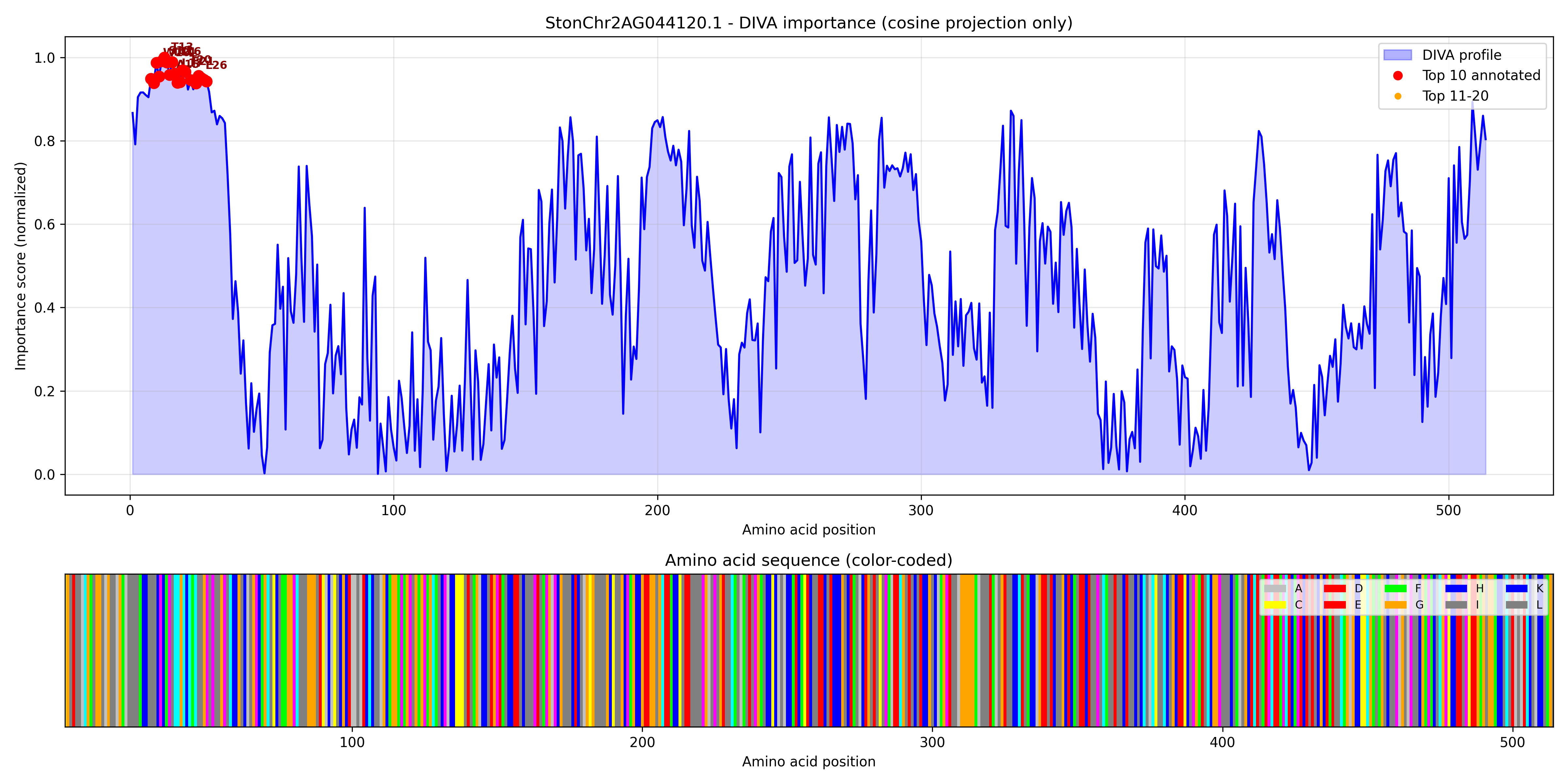

### StonChr2AG051160.1_importance.pdf

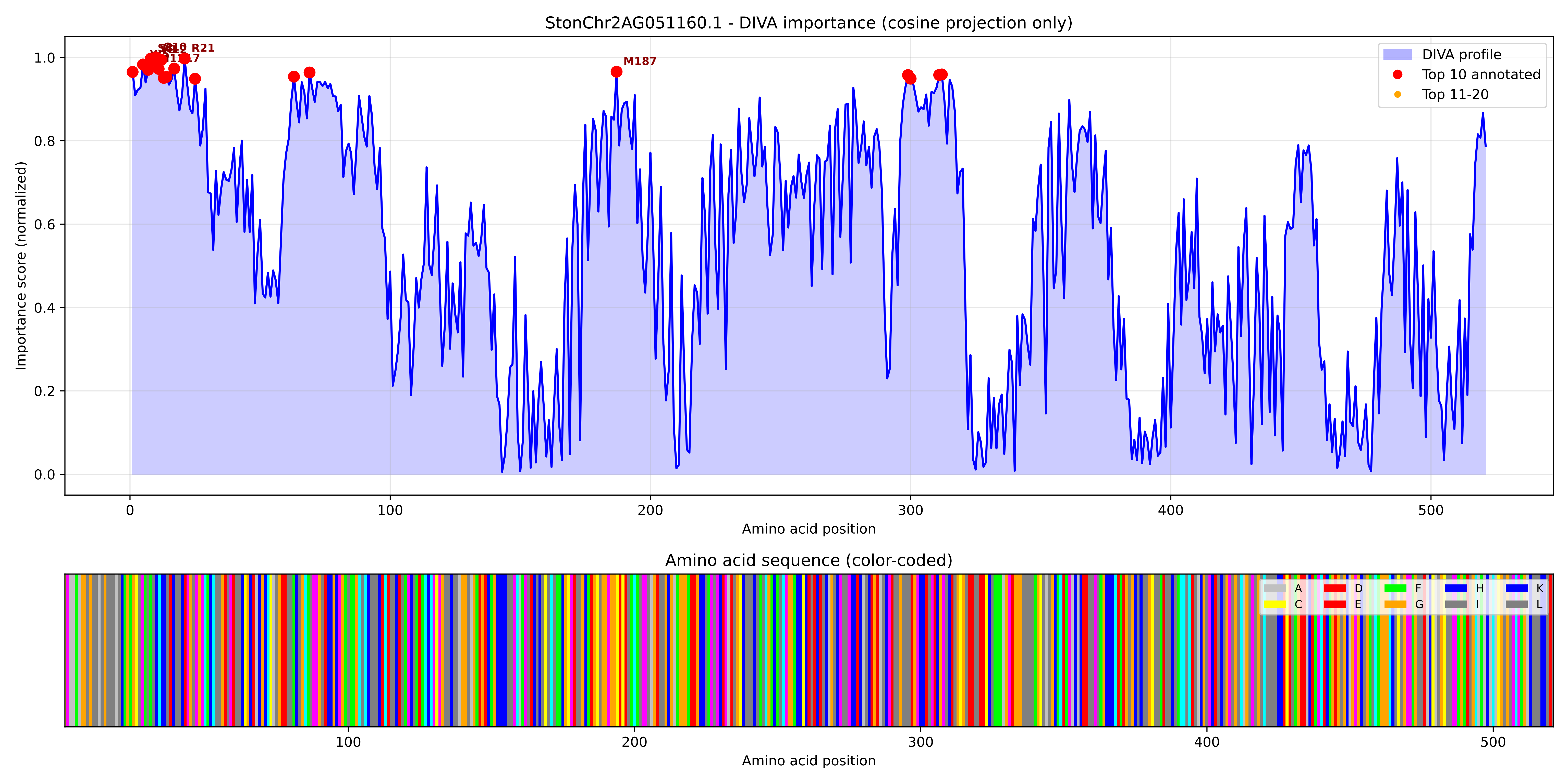

### StonChr2AG051160.1_importance.png

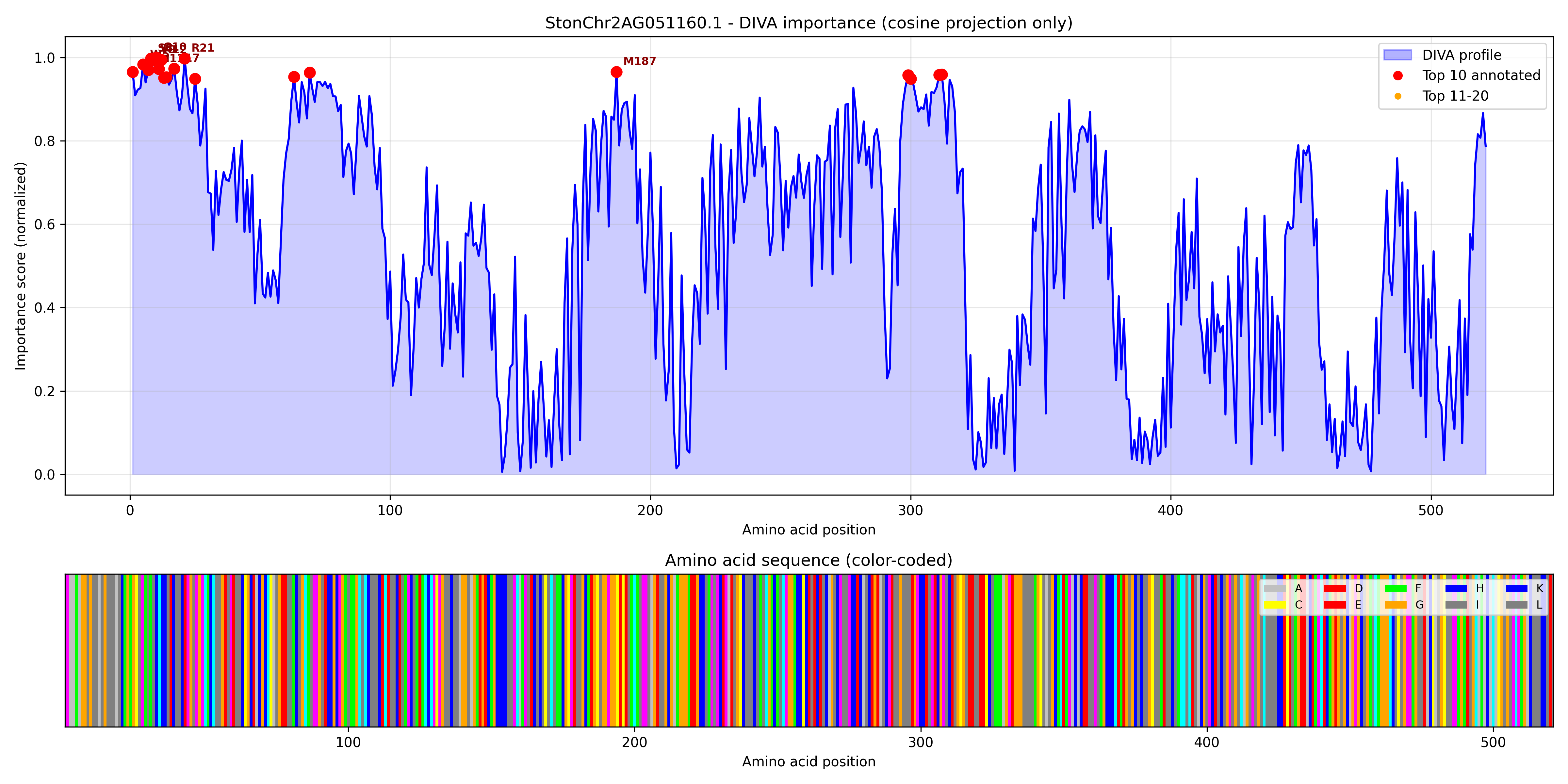

### StonChr2AG051170.1_importance.pdf

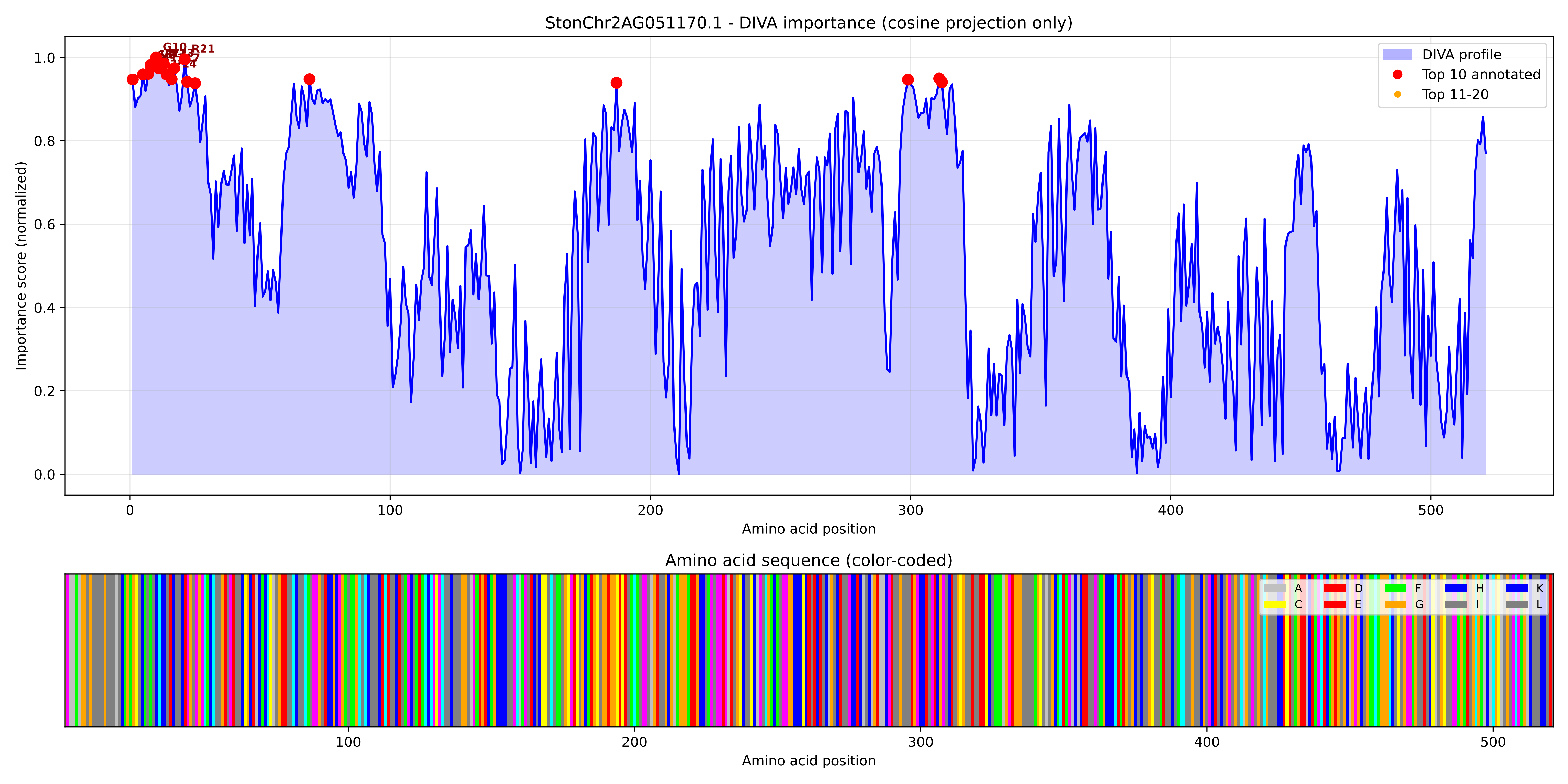

### StonChr2AG051170.1_importance.png

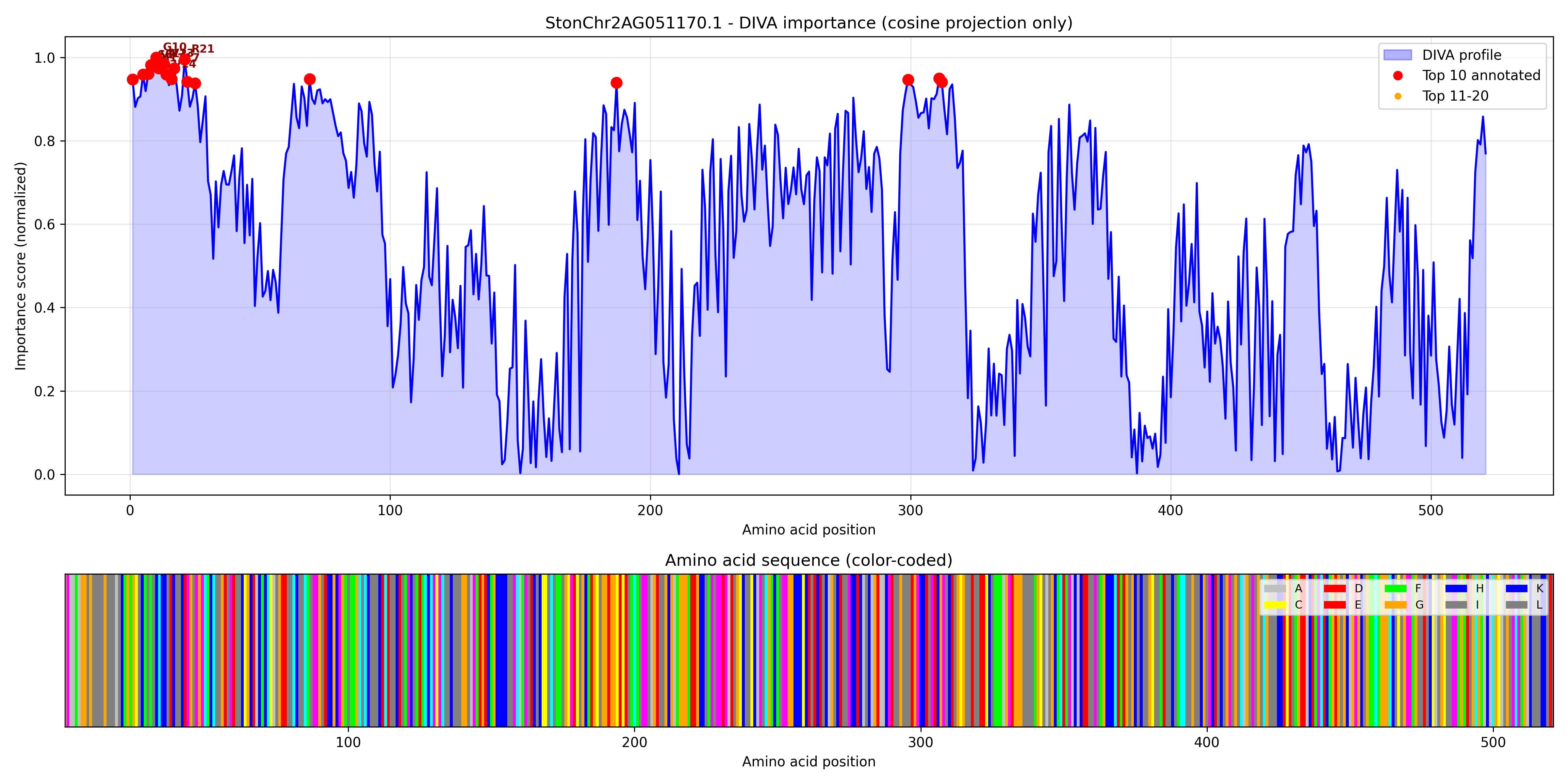

### StonChr2AG051590.1_importance.pdf

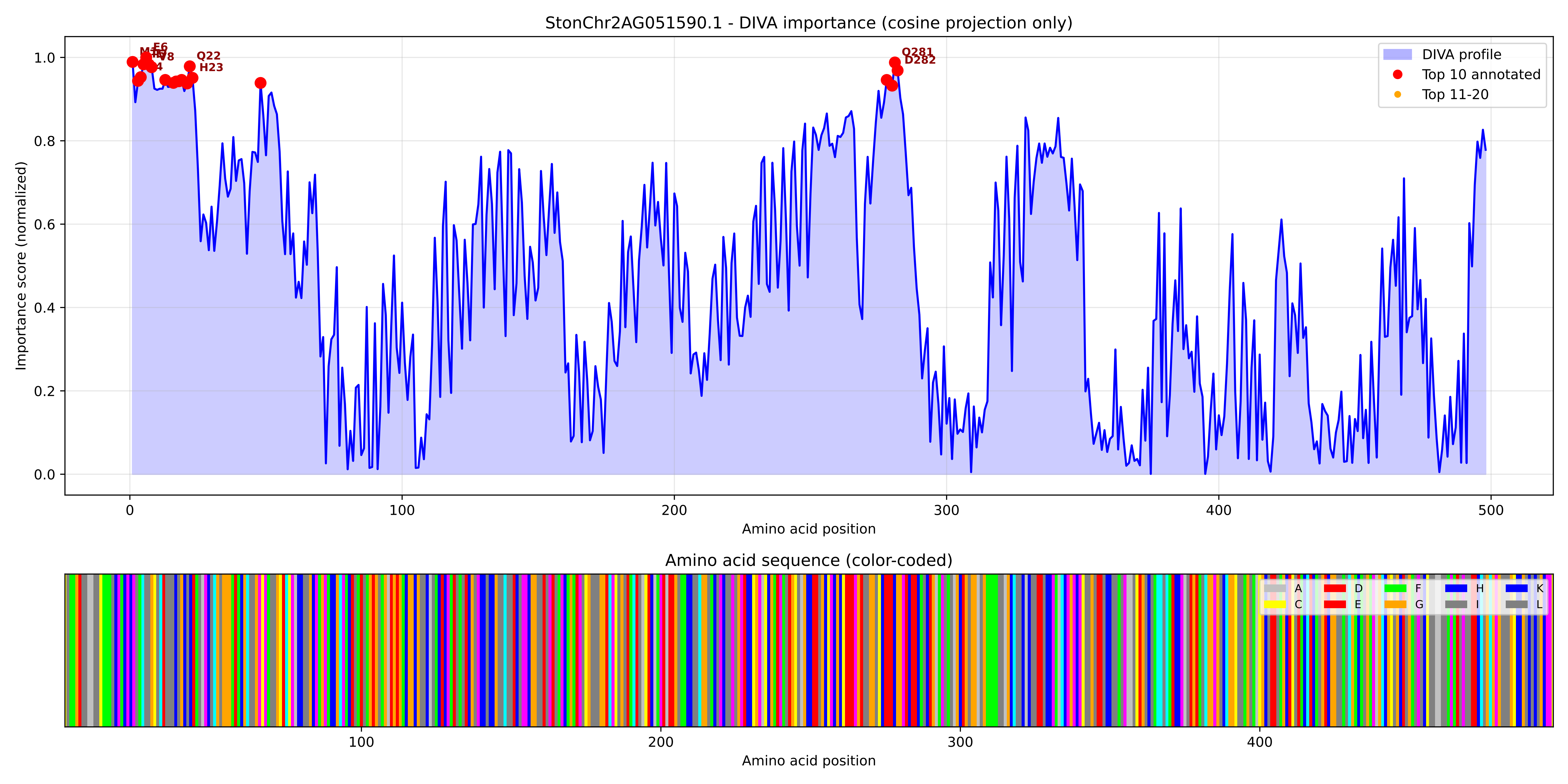

### StonChr2AG051590.1_importance.png

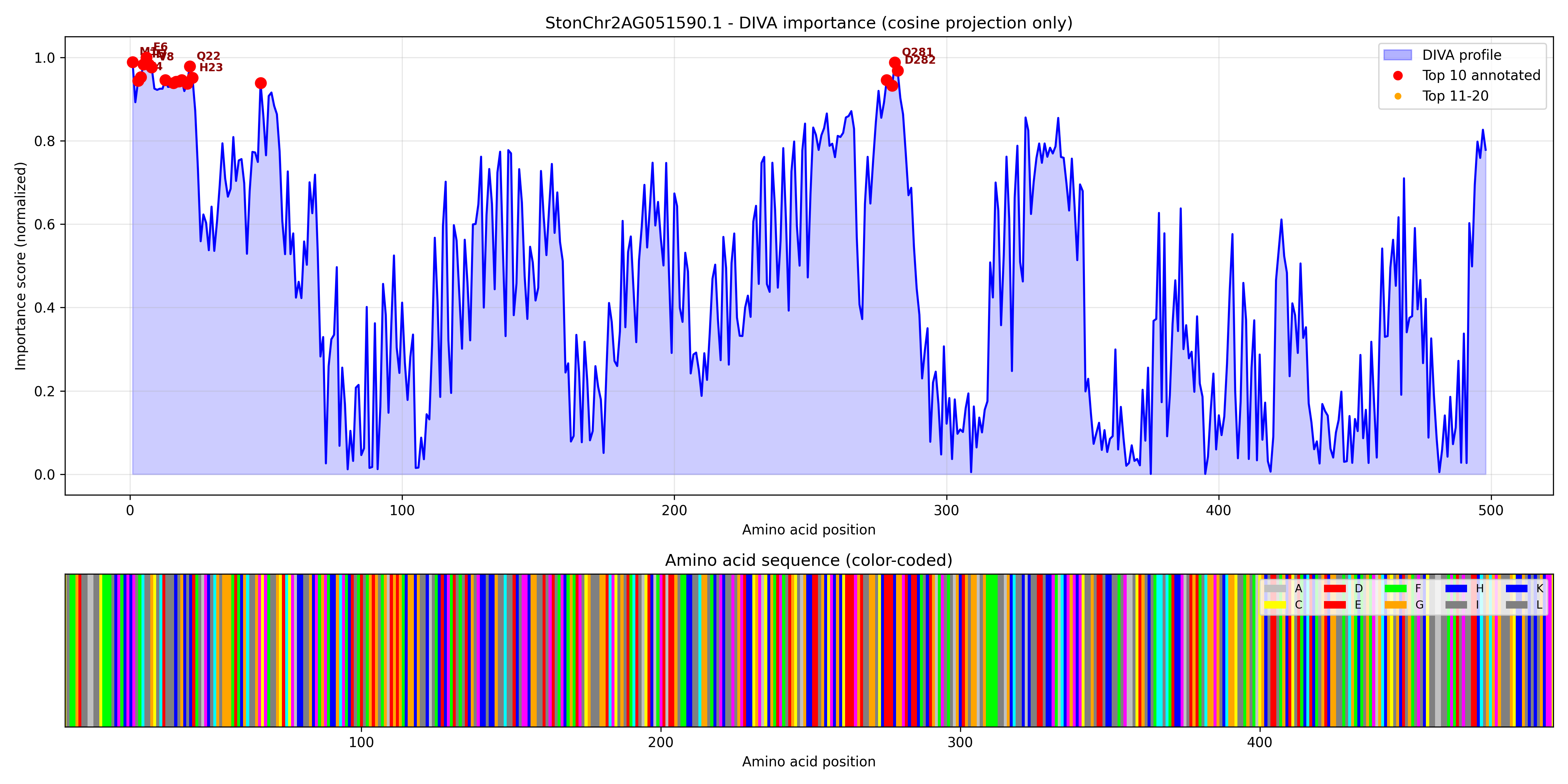

### StonChr2AG056020.1_importance.pdf

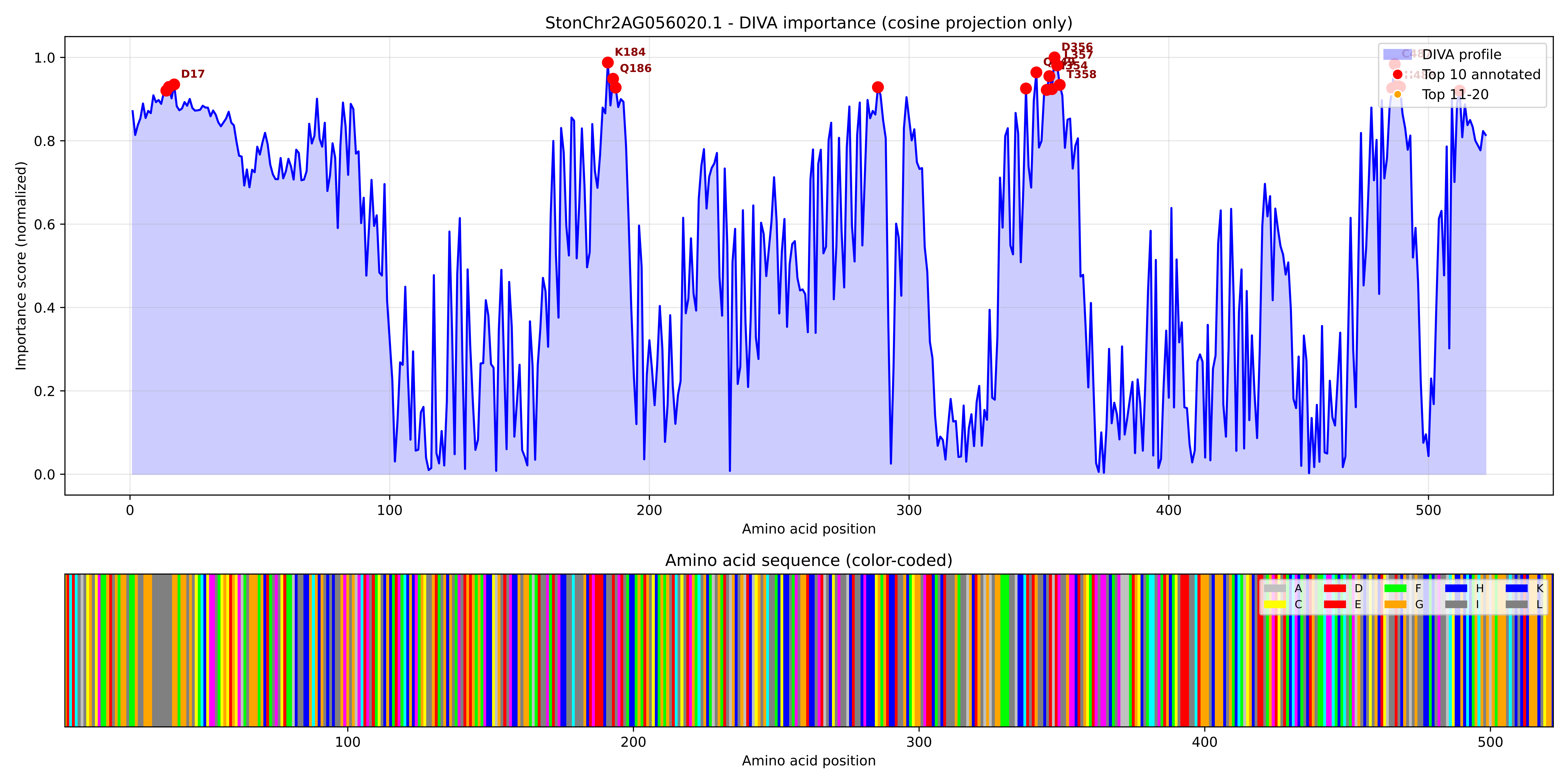

### StonChr2AG056020.1_importance.png

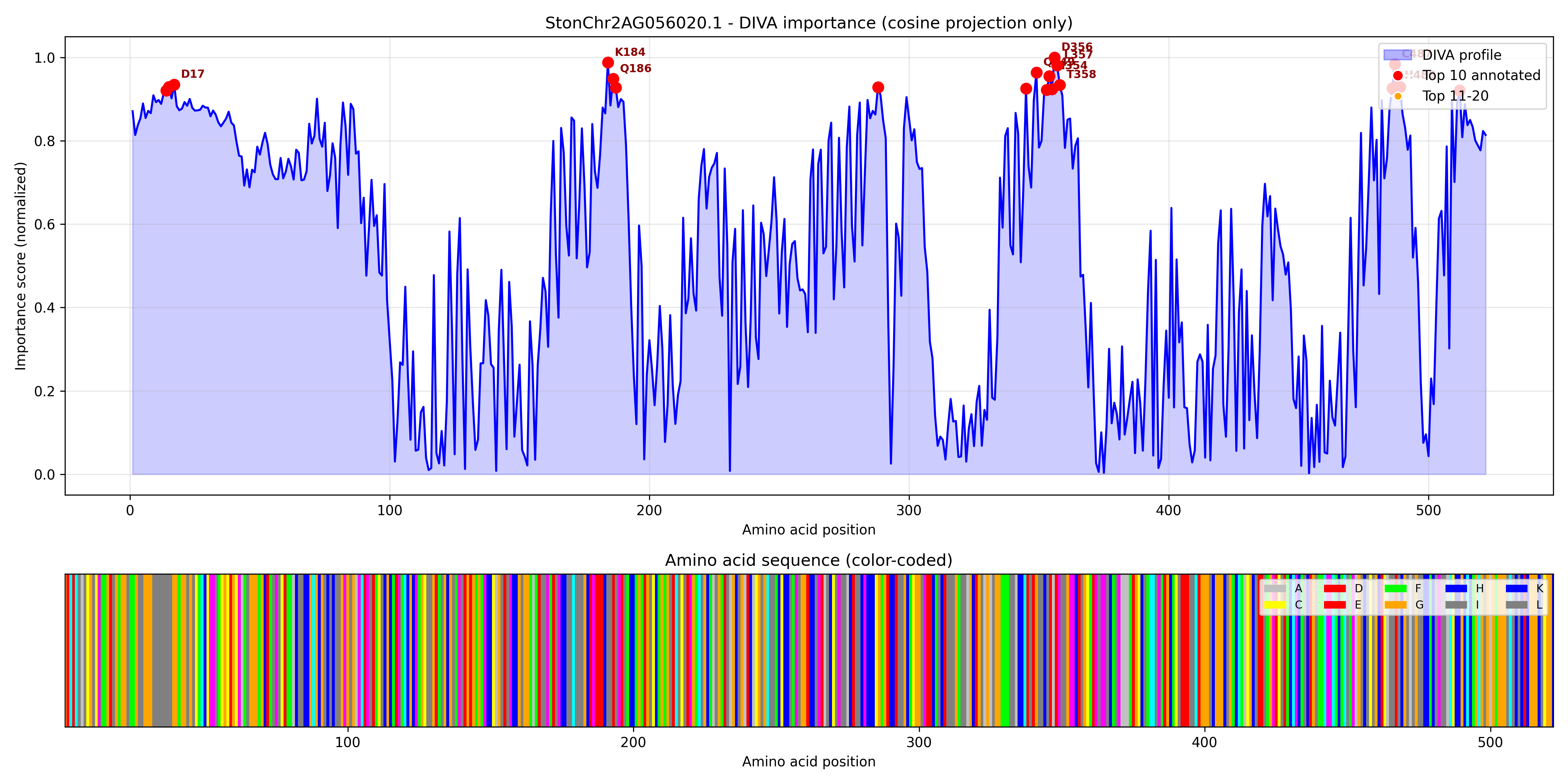

### StonChr2AG056390.1_importance.pdf

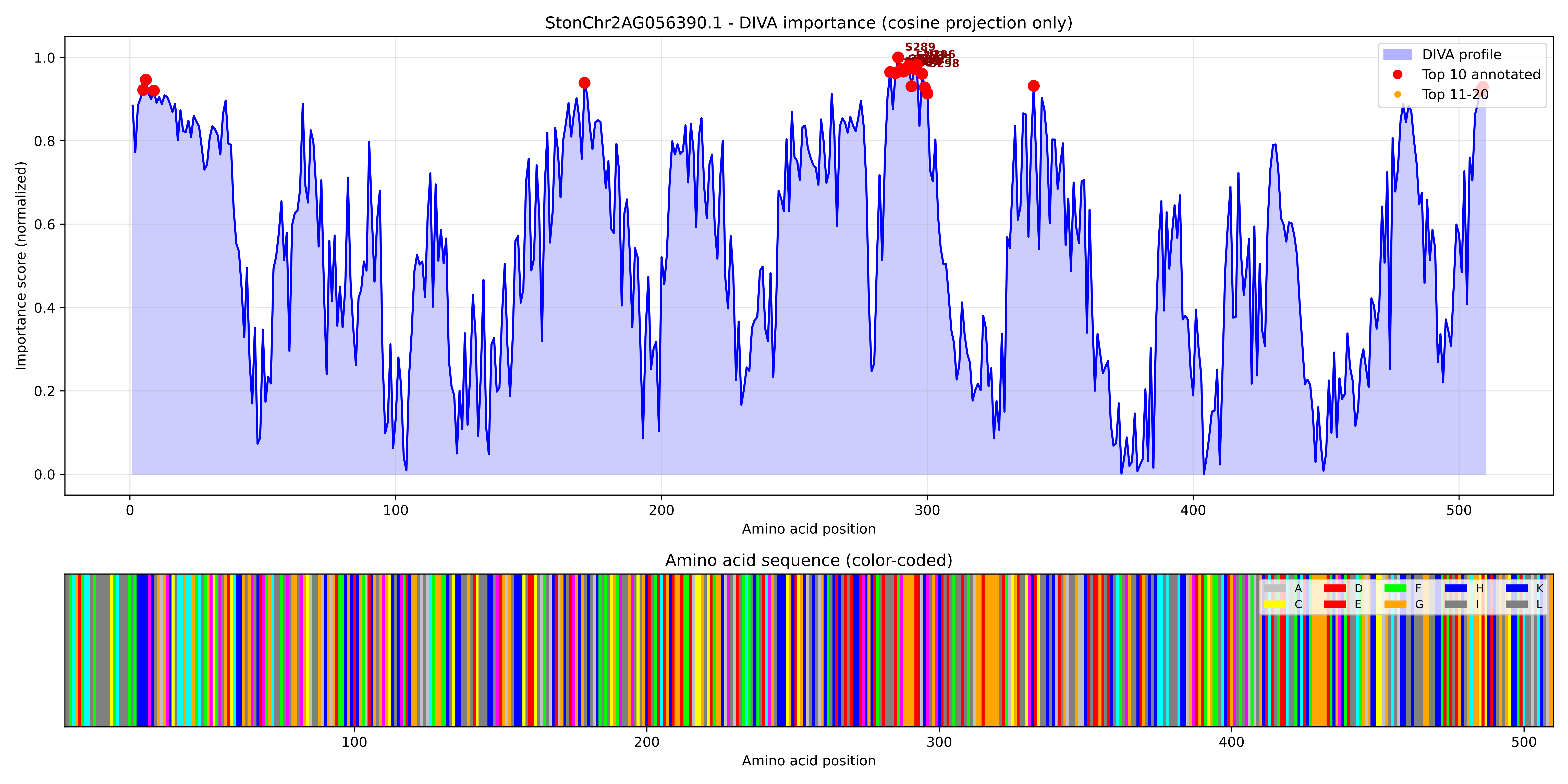

### StonChr2AG056390.1_importance.png

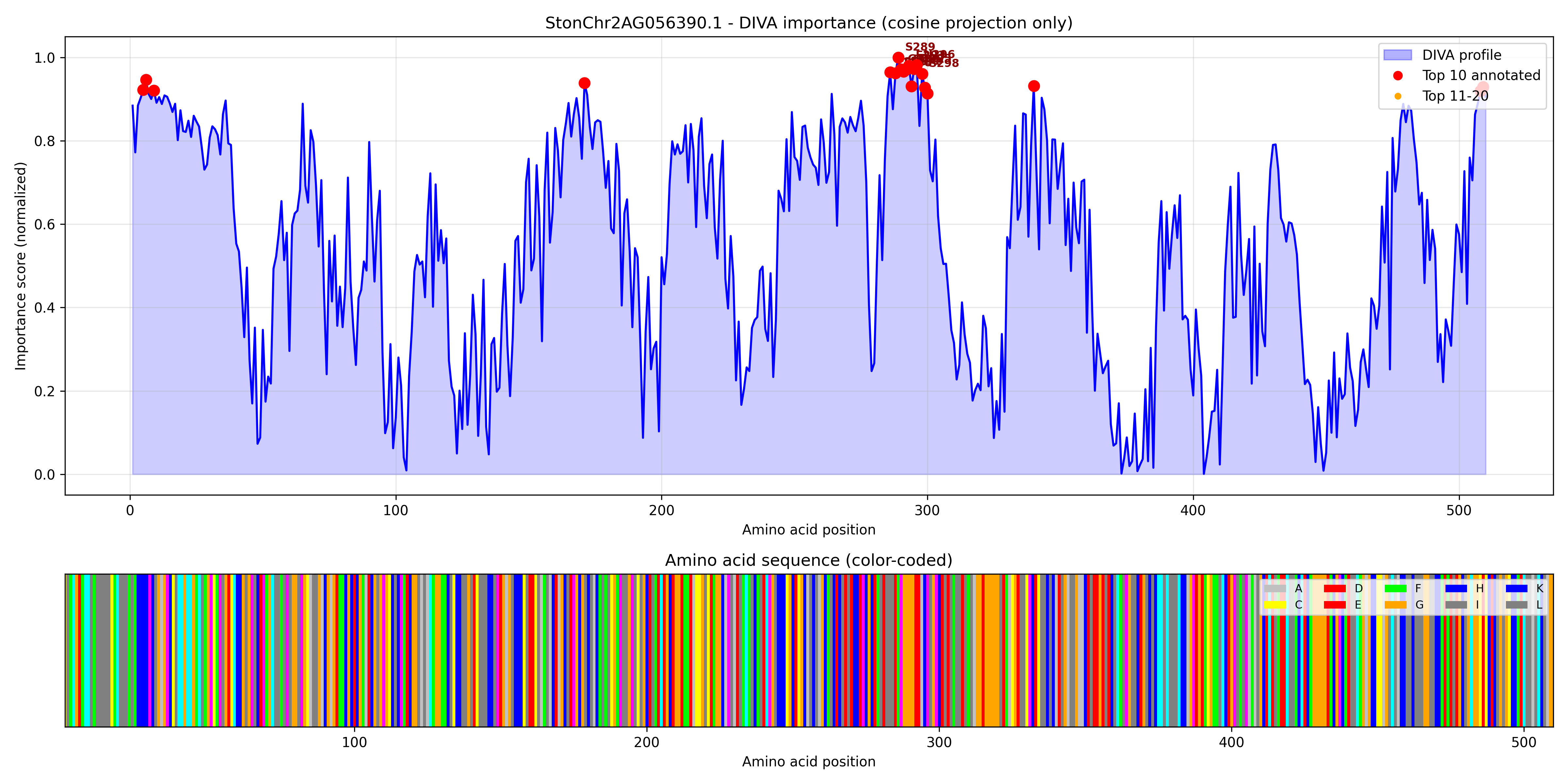

### StonChr2AG059180.1_importance.pdf

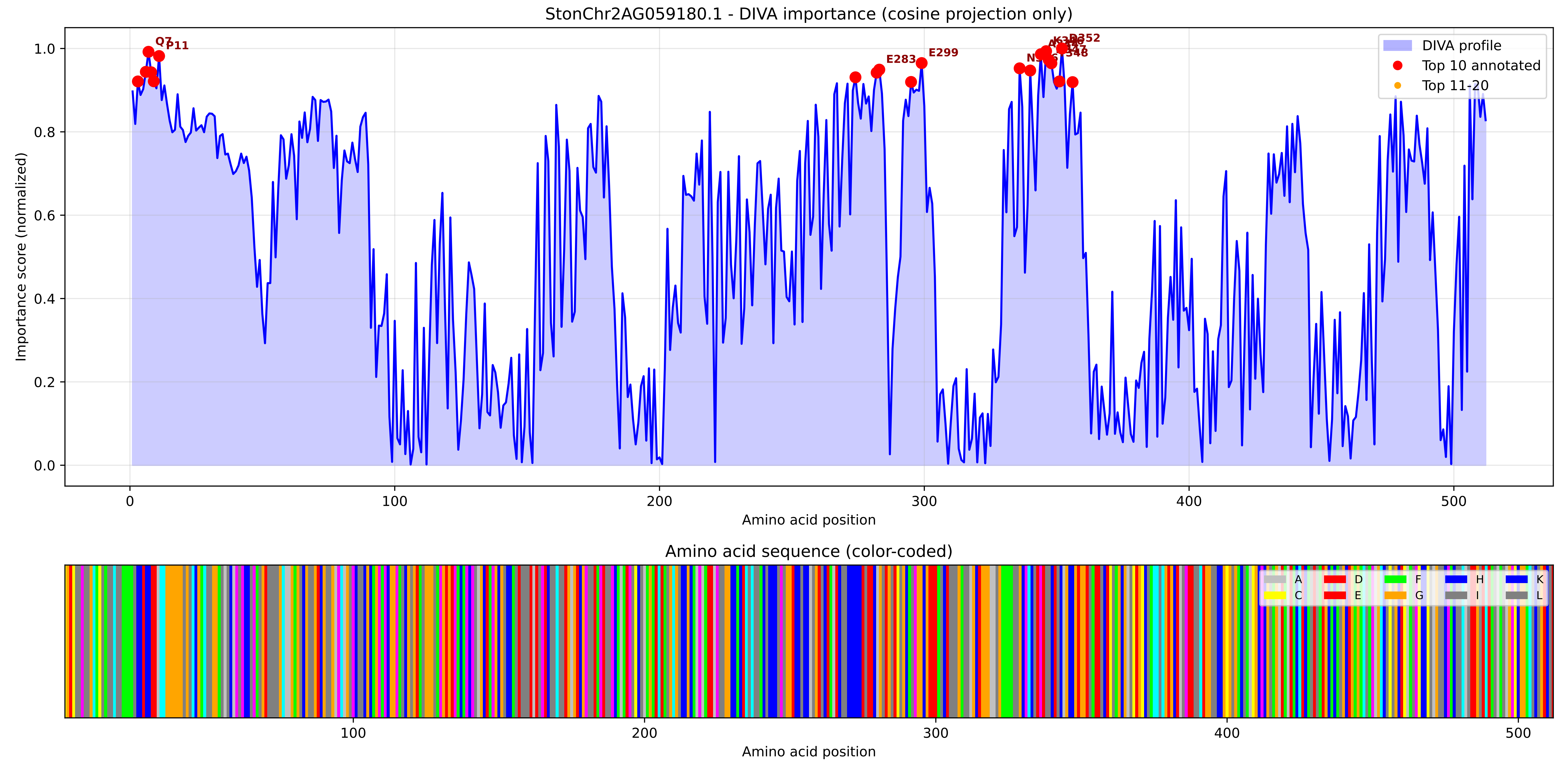

### StonChr2AG059180.1_importance.png

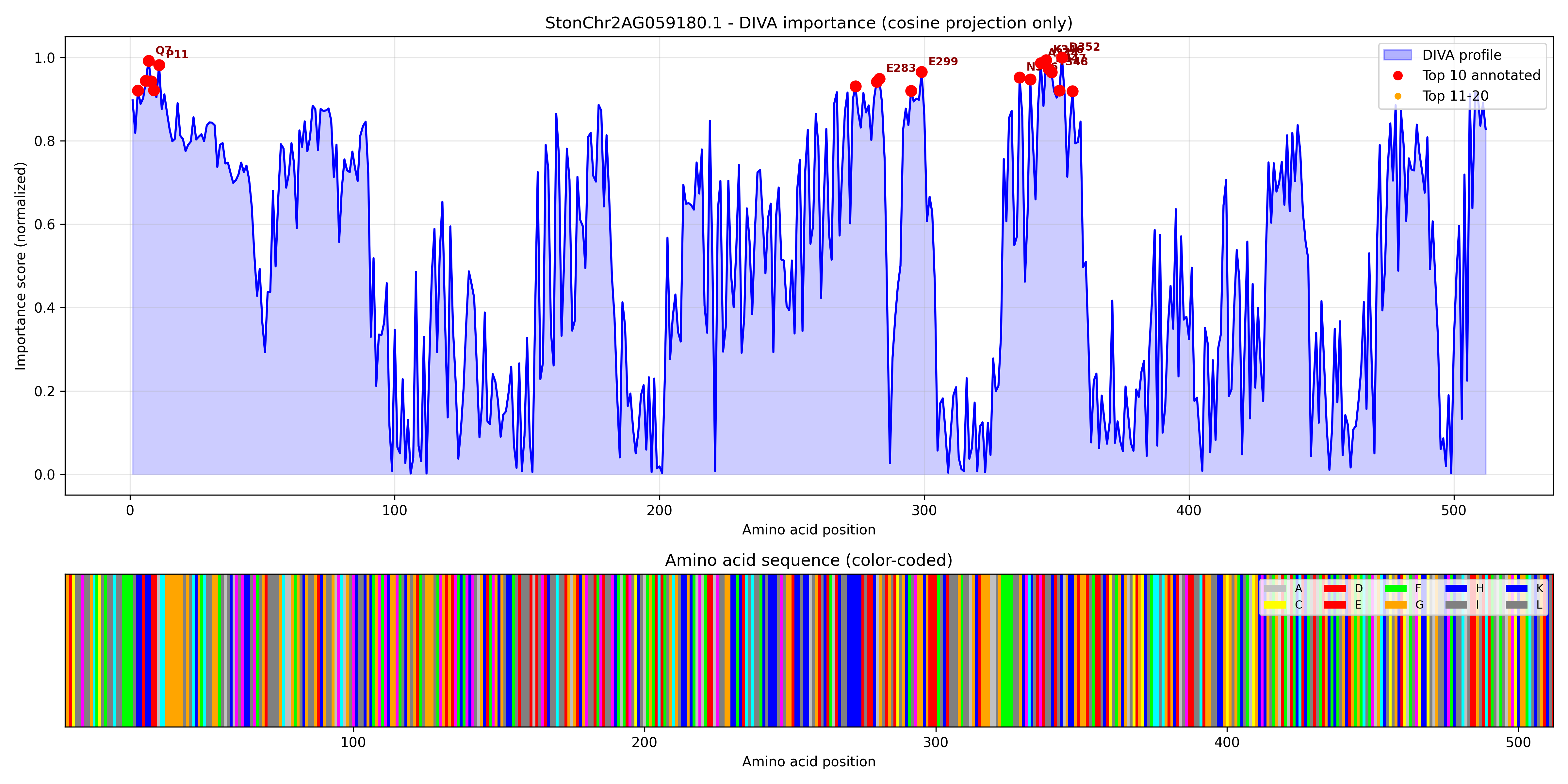

### StonChr2AG059190.1_importance.pdf

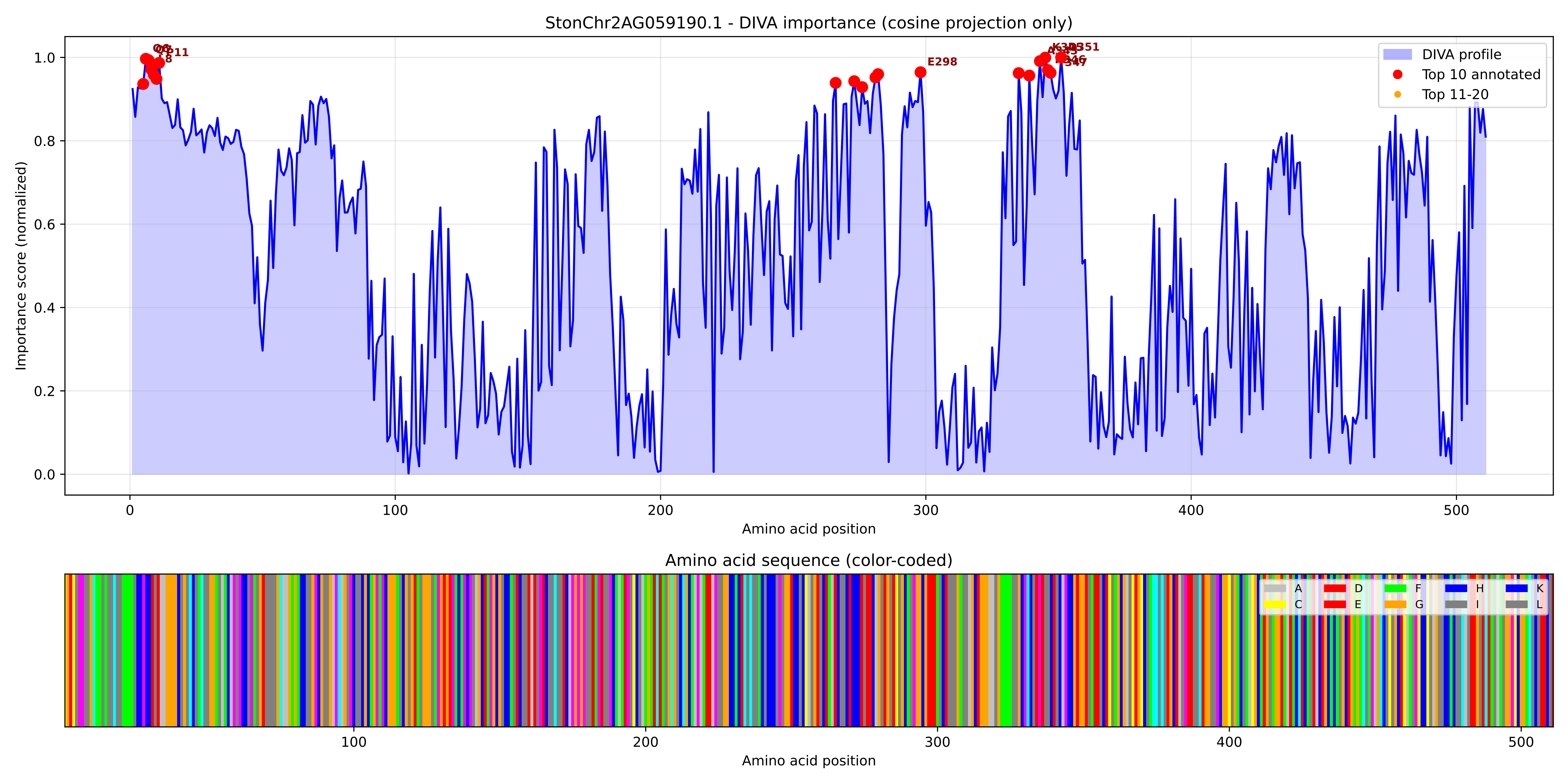

### StonChr2AG059190.1_importance.png

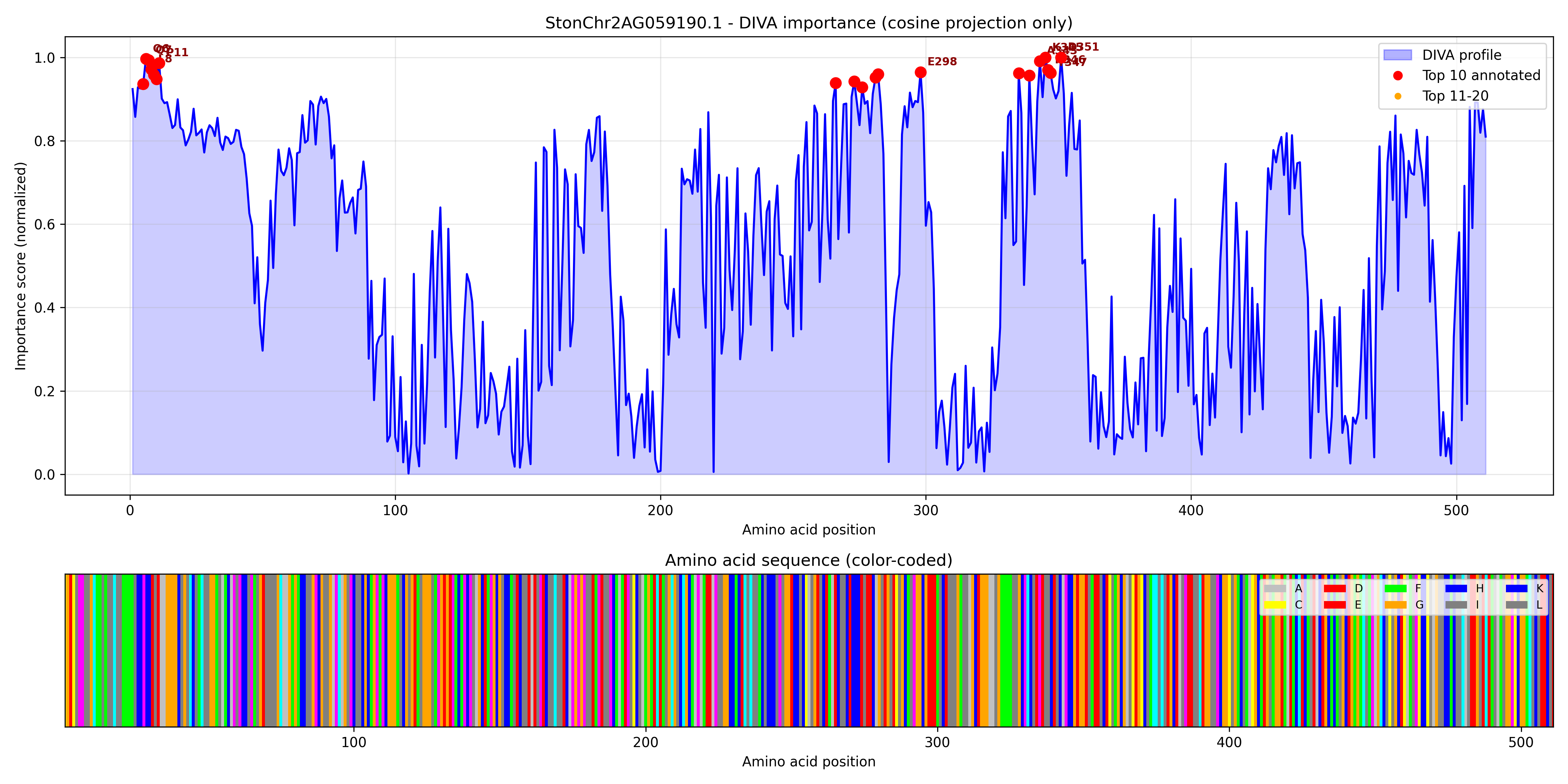

### StonChr2BG040140.1_importance.pdf

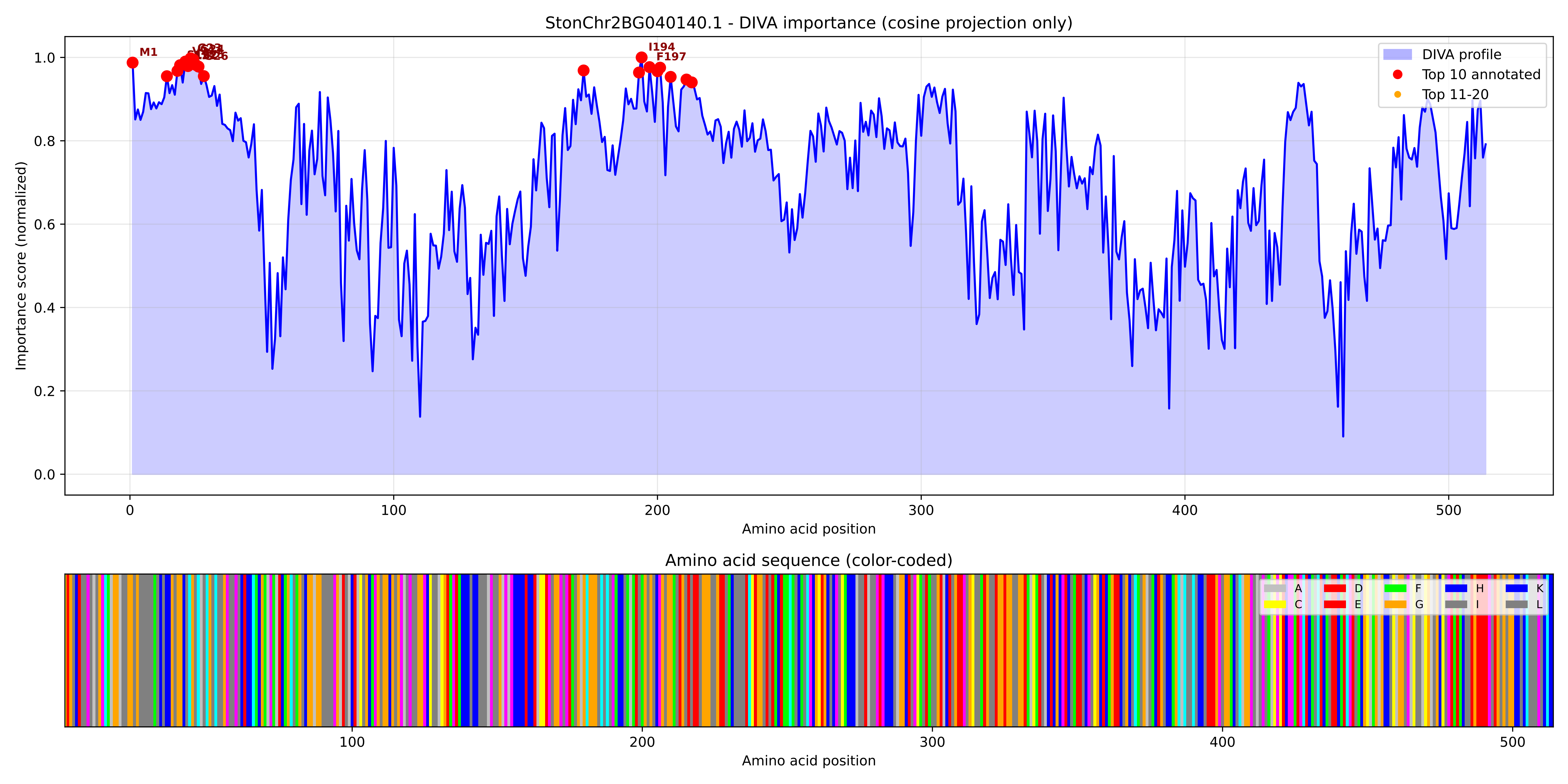

### StonChr2BG040140.1_importance.png

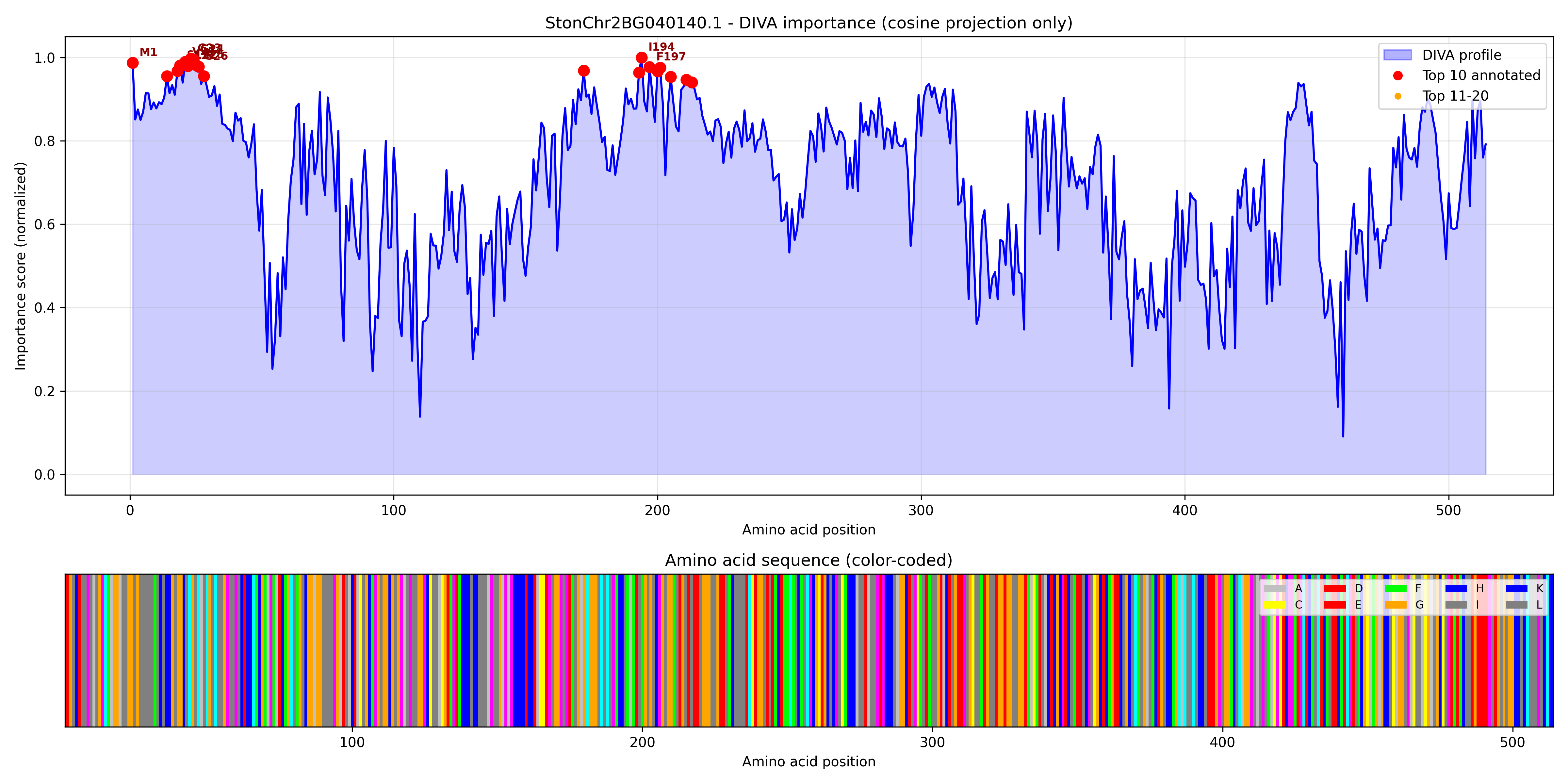

### StonChr2BG043800.1_importance.pdf

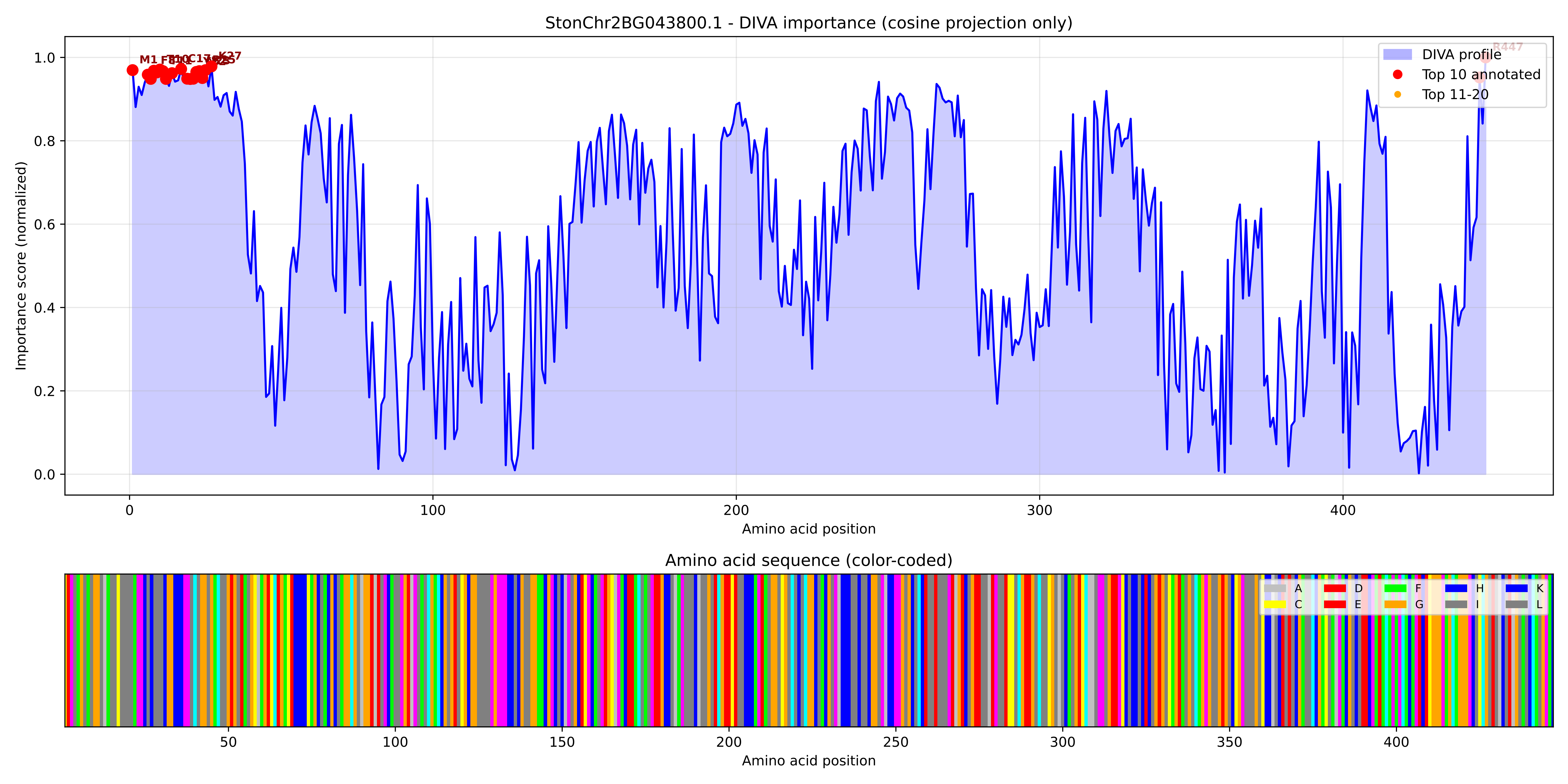

### StonChr2BG043800.1_importance.png

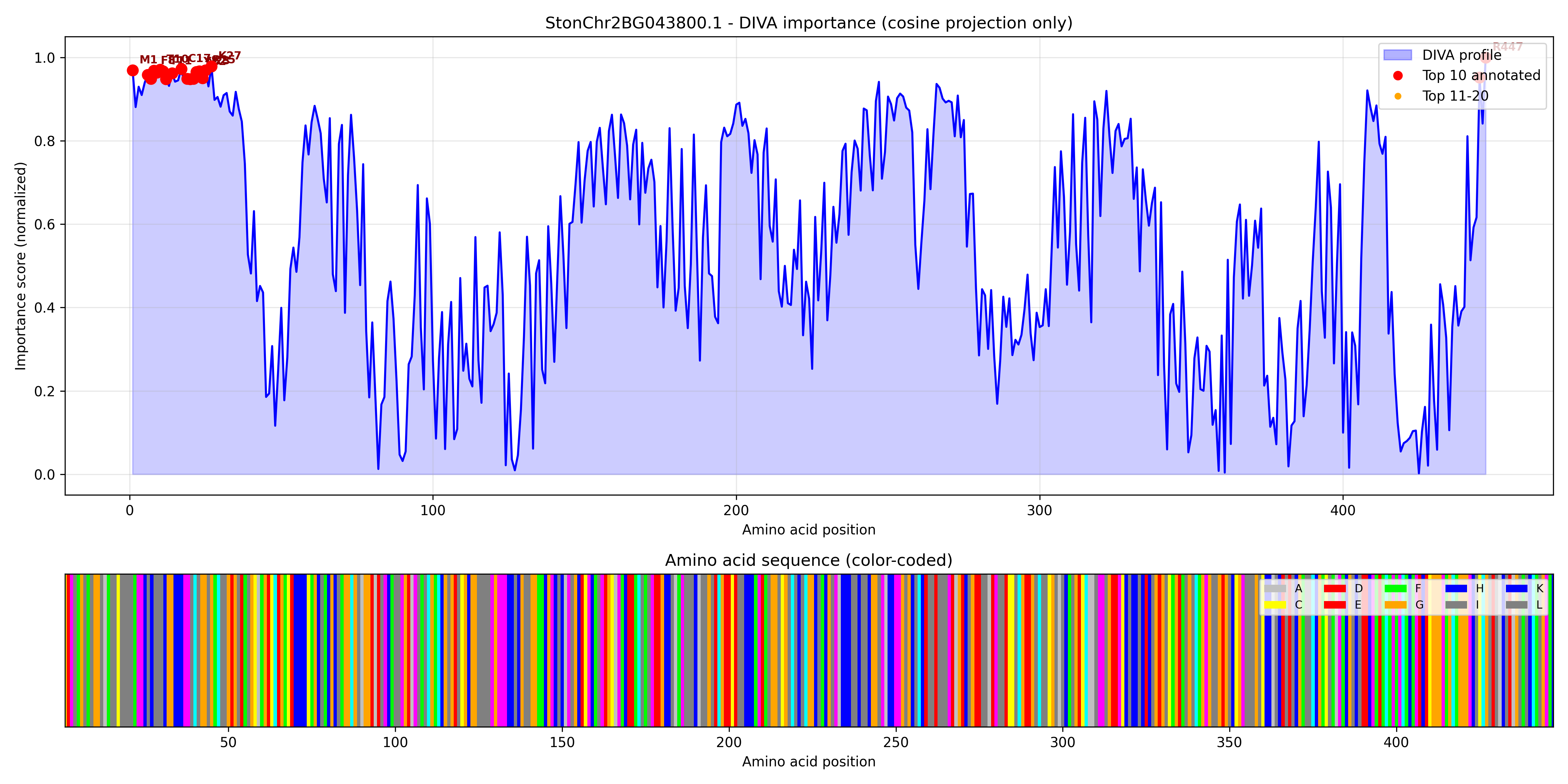
